# Mendelian Randomization for causal inference accounting for pleiotropy and sample structure using genome-wide summary statistics

**DOI:** 10.1101/2021.03.11.434915

**Authors:** Xianghong Hu, Jia Zhao, Zhixiang Lin, Yang Wang, Heng Peng, Hongyu Zhao, Xiang Wan, Can Yang

## Abstract

Mendelian Randomization (MR) is a valuable tool for inferring causal relationships among a wide range of traits using summary statistics from genome-wide association studies (GWASs). Existing summary-level MR methods often rely on strong assumptions, resulting in many false positive findings. To relax MR assumptions, ongoing research has been primarily focused on accounting for confounding due to pleiotropy. Here we show that sample structure is another major confounding factor, including population stratification, cryptic relatedness, and sample overlap. We propose a unified MR approach, MR-APSS, which (i) accounts for pleiotropy and sample structure simultaneously by leveraging genome-wide information; and (ii) allows to include more genetic variants with moderate effects as instrument variables (IVs) to improve statistical power without inflating type I errors. We first evaluated MR-APSS using comprehensive simulations and negative controls, and then applied MR-APSS to study the causal relationships among a collection of diverse complex traits. The results suggest that MR-APSS can better identify plausible causal relationships with high reliability. In particular, MR-APSS can perform well for highly polygenic traits, where the IV strengths tend to be relatively weak and existing summary-level MR methods for causal inference are vulnerable to confounding effects.

## Introduction

Inferring the causal relationship between a risk factor (exposure) and a phenotype of interest (outcome) is essential in biomedical research and social science [1]. Although randomized controlled trials (RCTs) are the gold standard for causal inference, RCTs can be very costly and sometimes even infeasible or unethical (e.g., random allocation to prenatal smoking) [2]. Mendelian randomization (MR) was introduced to mimic RCTs for causal inference in observational studies [3, 4]. Recently, MR analysis has drawn increasing attention [5] because it can take summary statistics from Genome-Wide Association Studies (GWASs) as input, including SNP effect size estimates and their standard errors, to investigate causal relationship among human complex traits.

MR is an instrumental variable (IV) method to infer the causal relationship between an exposure and an outcome, where genetic variants, e.g., single-nucleotide polymorphisms (SNPs), serve as IVs of the exposure [6, 7]. To eliminate the influence of confounding factors, conventional MR methods rely on strong assumptions, including **(A-I)** IVs are associated with the exposure; **(A-II)** IVs are independent of confounding factors; and **(A-III)** IVs only affect the outcome through the exposure. However, assumptions **(A-II)** and **(A-III)** are often not satisfied in practice due to confounding factors hidden in GWAS summary statistics, leading to false positive findings [5, 8]. To perform causal inference with genetic data, it is indispensable to distinguish two major confounding factors: pleiotropy [8] and sample structure [9, 10].

First, SNPs exhibit pervasive pleiotropic effects. Pleiotropy occurs when a genetic variant directly affects both exposure and outcome traits or indirectly through an intermediate phenotype [11]. Pleiotropy can induce trait association or genetic correlation in the absence of causality [11]. Due to the polygenicity of complex traits and linkage disequilibrium (LD) in the human genome, pleiotropic effects can widely spread across the whole genome [12]. Therefore, a substantial proportion of SNPs can carry pleiotropic effects and they fail to satisfy **(A-II)** and **(A-III)** on IVs in conventional MR methods.

Second, sample structure can lead to bias in SNP effect size estimates and introduce spurious trait associations. Here, sample structure encompasses population stratification, cryptic relatedness, and sample overlap in GWASs of the exposure and outcome traits. In the presence of population stratification and cryptic relatedness, SNPs can affect the outcome through sample structure and thus they violate assumptions **(A-II)** and **(A-III)** on IVs. Without correcting for sample structure, SNP effect size estimates can be severely biased, which may lead to misinterpretation on trait association and thus many false positive discoveries in causal inference. Sample overlap can also lead to spurious trait associations [13]. Although principal component analysis (PCA) [14] and linear mixed models (LMM) [15] are widely used to account for sample structure in GWASs, the results from LDSC [16] show that sample structure is often unsatisfactorily corrected in publicly available GWAS summary statistics.

To maximize the usage of publicly available GWAS summary statistics for causal inference, a number of summary-level MR methods have been developed, including Inverse Variance Weighted regression (IVW) [17], Egger [18], RAPS [19], dIVW [20], Weighted-median [21], Weighted-mode [22], MRMix [23], CML-MA [24], and CAUSE [25]. Despite these efforts, there are two major limitations in existing summary-level MR methods. First, most of them only use a small subset of SNPs passing the genome-wide significance (*p*-value ≤ 5 × 10^−8^) for causal inference. To account for pleiotropy (including correlated pleiotropy and uncorrelated pleiotropy [25]), it is challenging to fit a flexible model with limited information from genome-wide significant SNPs. Second, existing summary-level MR methods presume that PCA or LMM-based approaches have satisfactorily accounted for sample structure and thus they largely ignore the influence of sample structure in GWAS summary statistics. Due to the complexity of human genetics, sample structure driven by socioeconomic status [26] or geographic structure [27] may not be fully corrected by routine adjustment and it may remain as a major confounding factor hidden in GWAS summary statistics.

In this paper, we develop MR-APSS, a unified approach to MR Accounting for Pleiotropy and Sample Structure simultaneously. Specifically, we propose a foreground-background model to decompose the observed SNP effect sizes, where the background model accounts for confounding factors hidden in GWAS summary statistics, including correlated pleiotropy and sample structure, and the foreground model performs causal inference while accounting for uncorrelated pleiotropy. MR-APSS differs from existing methods in the following aspects. First, under the assumptions of LD score regression (LDSC) [16], the background model accounts for pleiotropy and sample structure using genome-wide summary statistics. In contrast, most summary-level MR methods only use SNPs passing the genome-wide significance (*p*-value ≤ 5 × 10^−8^). Second, MR-APSS allows us to include more SNPs without achieving the genome-wide significance as IVs to improve statistical power. With the pre-estimated background model, MR-APSS can inform whether an SNP belongs to the background component or the foreground component. Even in the presence of many invalid IVs, the type I error will not be inflated because only the foreground signals are used for causal inference. As more SNPs are included, the increasing amount of the foreground signal can improve the statistical power.

To demonstrate the effectiveness of MR-APSS, we have performed a comprehensive simulation study and analyzed 640 pairs of exposure and outcome traits from 26 GWASs. In the simulation study, we showed that MR-APSS still had satisfactory performance when the assumptions of IVs were violated. We examined MR-APSS on a wide spectrum of complex traits using GWAS summary statistics, including psychiatric/neurological disorders, social traits, anthropometric traits, cardiovascular traits, metabolic traits, and immune-related traits. Real data results indicate that pleiotropy and sample structure are two major confounding factors. By rigorous statistical modeling of these confounding factors, MR-APSS not only avoids many false positive findings but also improves the statistical power of MR. When inferring causal relationships among highly polygenic traits, such as psychiatric disorders and social traits, the strengths of IVs tend to be relatively weak and causal inference is vulnerable to confounding effects. Thus, existing MR methods will suffer from either low statistical power or inflated type I errors. The empirical results indicate that MR-APSS is particularly useful in this scenario because it accounts for confounding factors and allows for incorporating many IVs with moderate effects, demonstrating its advantage over existing MR methods.

## Results

### Overview of MR-APSS

Causality, pleiotropy, and sample structure are three major sources to induce correlation between GWAS estimates of exposure-outcome traits. To distinguish causality from correlation, it is indispensable to eliminate the possibility that correlation is induced by confounding factors, such as pleiotropy and sample structure (including population stratification, cryptic relatedness, and sample overlap).

MR-APSS takes GWAS summary statistics of exposure and outcome traits as its input and performs causal inference based on a proposed foreground-background model (see an overview in Fig. 1 and details in the Materials and Methods section). Under the assumptions of LDSC [16] (see details in SI Appendix, section 1.1), the background model can effectively account for confounding factors by disentangling pleiotropy (Fig. 1B) and sample structure (Fig. 1C). This is because the pleiotropic effects can be tagged by LD and the influence of sample structure is uncorrelated with LD [16]. In addition to the LDSC assumptions in the background model, we have made two key assumptions for causal inference. First, we assume that the correlated pleiotropy effects can be approximately characterized by the genetic correlation which can be estimated from genome-wide summary statistics. Second, we assume that the direct effect is independent of the instrument strength in our foreground model (known as the InSIDE condition). This is reasonable because correlated pleiotropy effects have been accounted for using genome-wide genetic correlation. By further accounting for selection bias [28] due to selection of IVs (see Materials and Methods section), the foreground model can use the classical causal diagram to perform causal inference (Fig. 1A). In summary, our method requires the LDSC assumptions for the background model and the InSIDE condition for the foreground model to relax assumptions **(A-II)** and **(A-III)**.

**Figure 1:**
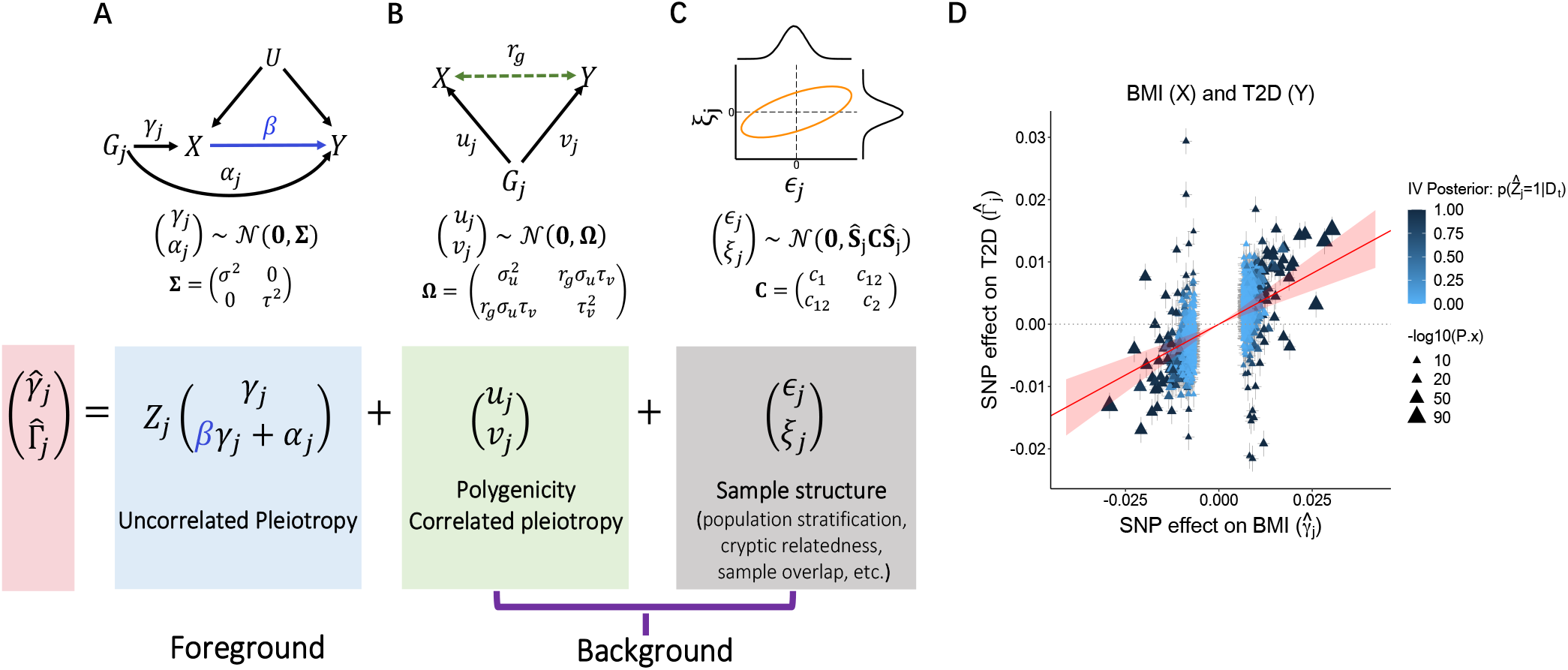
The MR-APSS approach. To infer the causal effect *β* between exposure *X* and outcome *Y*, MR-APSS uses a foreground-background model to characterize the estimated effects of SNPs *G_j_* on *X* and *Y* (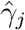 and 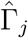) with standard errors 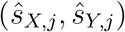, where the background model accounts for polygenicity, correlated pleiotropy (**B**) and sample structure (**C**), and the foreground model (**A**) aims to identify informative instruments and account for uncorrelated pleiotropy to perform causal inference. (**D**) We consider inferring the causal relationship between BMI and T2D as an illustrative example of MR-APSS. The estimated causal effect is indicated by a red line with its 95% confidence interval indicated by the shaded area in transparent red color. Triangles indicate the observed SNP effect sizes (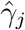 and 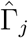). The color of triangles indicates the posterior of a valid IV, i.e., the posterior of an IV carrying the foreground signal (*Z_j_* = 1, dark blue) or not (*Z_j_* = 0, light blue).

### Compared methods

Because MR-APSS uses the GWAS summary statistics as its input, we mainly compare MR-APSS with nine summary-level MR methods and grouped them (including MR-APSS) into three groups based on their assumptions, including IVW from group 1; Egger, RAPS and dIVW from group 2; and Weighted-median, Weighted-mode, MRMix, CML-MA, CAUSE and MR-APSS from group 3 (see Table 1). We provide a review of them in SI Appendix, sections 2.1-2.2. We show theoretically that the IVW estimator and the dIVW estimator can be biased in the presence of pleiotropy and sample structure (SI Appendix, section 2.6). To establish a better connection with causal literature, we also provide a review of individual-level MR methods in SI Appendix, section 2.3 and Table S1. We conducted comparisons between summary-level MR methods and individual-level MR methods. Detailed results are provided in SI Appendix, sections 3.3 and 4.4, Figs. S1, S6, and S16-S21.

**Table 1:** Summary of ten summary-level MR methods

| Method | (A-II) | (A-III) | Key assumptions | Sample structure | Selection bias |
| --- | --- | --- | --- | --- | --- |
| IVW [17] | ✓ | ✓ | All IVs are valid; NOME. | × | × |
| Egger [18] | ✓ | × | InSIDE; NOME; Directional pleiotropy ($\mathbb{E}(\alpha_j) = \mu$). | × | × |
| RAPS [19] | ✓ | × | InSIDE; Balanced pleiotropy ($\alpha_j \sim \mathcal{N}(0, \tau^2)$). | × | × |
| dIVW [20] | ✓ | × | InSIDE; Balanced pleiotropy ($\alpha_j \sim \mathcal{N}(0, \tau^2)$). | × | ✓ |
| Weighted-median [21] | × | × | Majority valid; NOME. | × | × |
| Weighted-mode [22] | × | × | Plurality valid. | × | × |
| MRMix [23] | × | × | Plurality valid. | × | × |
| cML-MA [24] | × | × | Plurality valid. | × | × |
| CAUSE [25] | × | × | All IVs can be invalid, but majority of IVs should not be affected by correlated pleiotropy. | Sample overlap | × |
| **MR-APSS** | × | × | All IVs can be invalid; Assumptions of LDSC; InSIDE in the foreground model. | ✓ | ✓ |

IV: Instrumental Variable; Three IV assumptions: (A-I) IVs are associated with the exposure; (A-II) IVs are independent of confounders; and (A-III) IVs only affect the outcome through the exposure.

### Simulation studies

To evaluate MR-APSS in various scenarios and compare it with nine MR methods in Table 1, we first perform simulation studies under the MR-APSS model. After that, we investigate the robustness of MR-APSS in the presence of model misspecification.

For exposure and outcome traits, we used 47,049 SNPs on chromosomes 1 and 2 of 20,000 individuals of white British ancestry randomly drawn from the UK BioBank (UKBB). SNP effect sizes (*γ_j_, α_j_, u_j_, v_j_*) were generated from the relationship shown in Fig. 1 and Eq. [1] in Materials and Methods section. Based on real genotype data and simulated SNP effect sizes, we generated both traits and obtained summary statistics (see details in SI Appendix, section 3.1). The relationship shown in Fig. 1 is composed of the background signal and the foreground signal. For the background signal, polygenic effects (*u_j_,v_j_*) of all SNPs were normally distributed with variance components 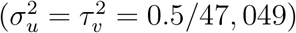, such that the heritabilities of both exposure *X* and outcome *Y* were specified at 0.5. The magnitudes of the error terms (*ϵ_j_*, *ξ_j_*) were determined by the fixed sample sizes of 20, 000. For the foreground signal, we randomly assigned 500 out of 47,049 SNPs as IVs. As the instrument strength (*γ_j_*) and the magnitude of the direct effect (*α_j_*) are given by variance components *σ*^2^ and *τ*^2^ (Fig. 1), we specified 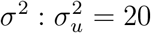 to mimic real data scenarios. We set 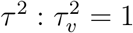, so the magnitude of the direct effects in the foreground model is the same as that of the polygenic effects.

We compared MR-APSS with nine MR methods, including IVW, dIVW, RAPS, MRMix, cML-MA, Egger, CAUSE, Weighted-median, and Weighted-mode. Note that the performance of MR methods depends on the selected IVs. Using a stringent criterion, fewer SNPs will be selected as IVs and MR methods tend to have lower power of detecting the causal effect and lower false positive rate. When more SNPs are included using a loose criterion, MR methods tend to have higher power but higer false positive rate because their model assumptions are more likely to be violated. To evaluate the performance of MR methods under null (*β* = 0), we used a stringent criterion (IV threshold *p* = 5 × 10^−6^) to select IVs for IVW, dIVW, RAPS, MRMix, cML-MA, Egger, Weighted-median, and Weighted-mode. For CAUSE, we used its default threshold *p* =1 × 10^−3^ to include IVs. For MR-APSS, we used *p* = 5 × 10^−4^. For all nine MR methods, we applied LD pruning (*r*^2^ = 0.01) to the selected IVs to ensure that they were nearly independent.

We first examined type I error control of different MR methods under null (*β* = 0) in the presence of genetic correlation induced by pleiotropy. We simulated data with genetic correlation but without correlation in estimation errors. Quantile-quantile plots of different MR methods are shown in Fig. 2A, 2B, 2E for genetic correlation *r_g_* = 0.2 (more results for different genetic correlations are given in SI Appendix, Fig. S2). Clearly, MR-APSS is the only method that produces well-calibrated *p*-values. To better examine how MR-APSS accounted for polygenicity and pleiotropy, we manually set the variance component of MR-APSS to zero, i.e., **Ω** = **0**. We denote this version of MR-APSS as MR-APSS (**Ω** = **0**). As shown in Fig. 2E, MR-APSS produced well-calibrated *p*-values while MR-APSS (**Ω** = **0**) produced overly inflated *p*-values. This suggests that variance component **Ω** plays a critical role in accounting for polygenicity and pleiotropy. We also noticed different performance of alternative MR methods (Fig. 2A, 2B). In the presence of non-zero genetic correlation, MR methods, such as IVW, dIVW, RAPS, MRMix, cML-MA, Weighted-median, and MR-APSS (**Ω** = **0**), tended to produce inflated *p*-values. Different from other MR methods, CAUSE produced very deflated *p*-values and thus CAUSE was very conservative in identifying causal effects.

**Figure 2:**
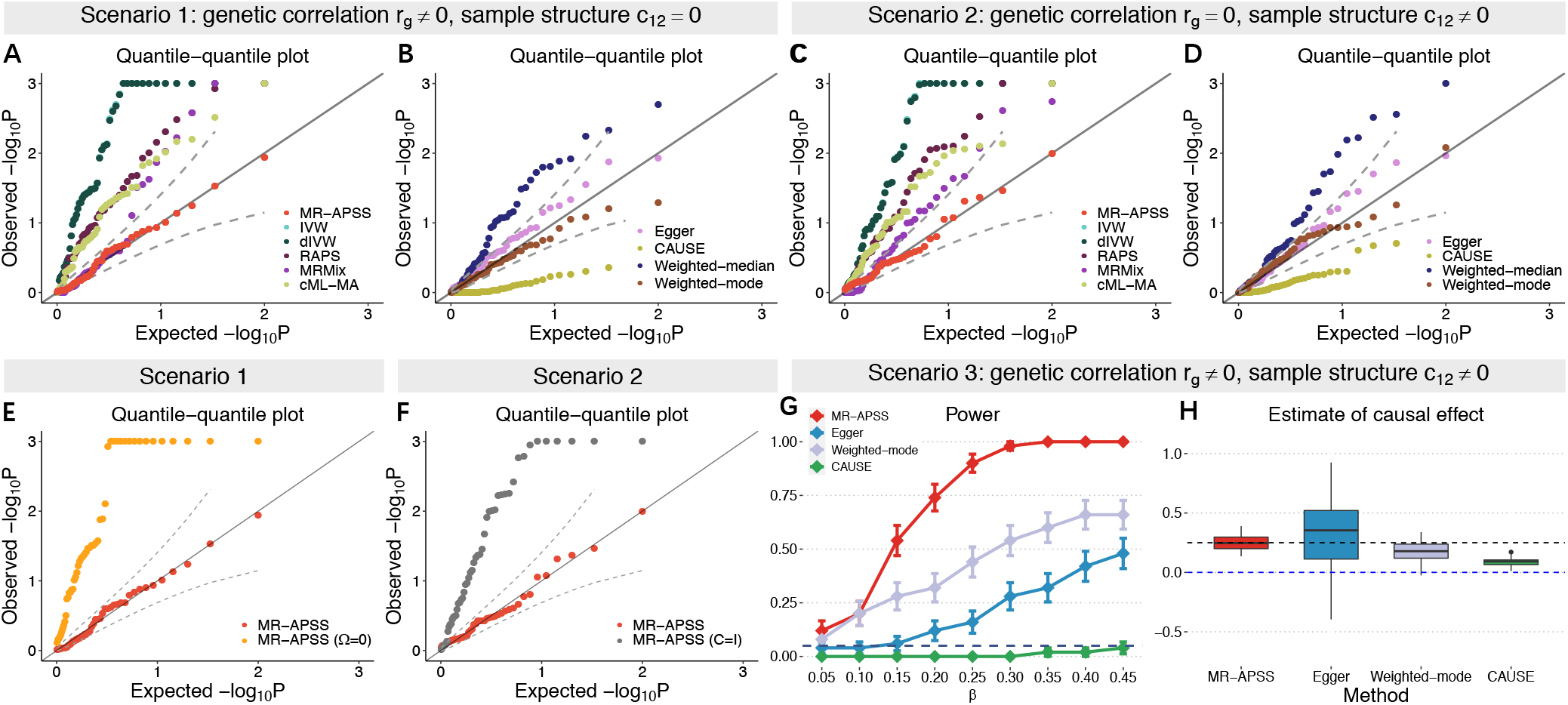
Comparison of ten summary-level MR methods on simulated data. (**A-F**) Quantile-quantile plots of – log_10_(*p*)-values from different methods under null simulations in the absence of causal effect (*β* = 0). Null simulations were performed under different scenarios: (**A, B, E**) Null simulations with genetic correlation (*r_g_* = 0.2) induced by pleiotropy, but without correlation in estimation errors (*c*_12_ = 0). (**C, D, F**) Null simulations in the presence of correlation in estimation errors (*c*_12_ = 0.15) due to sample structure, but in the absence of non-zero genetic correlation (*r_g_* = 0). Based on results in **A-D**, MR-APSS, Egger, Weighted-mode, and CAUSE do not provide overly inflated *p*-values. (**G,H**) Comparison of MR-APSS, Egger, Weighted-mode, and CAUSE under alternative simulations (*β* ≠ 0). (**G**) The power under the settings that the causal effect size *β* varied from 0.05 to 0.45. (**H**) Estimates of causal effect under the alternative simulations (*β* = 0.25). The results were summarized from 50 replications.

Next, we examined the type I error control under null (*β* = 0) in the presence of correlation between estimation errors due to sample structure. Specifically, we set genetic correlation *r_g_* = 0 and simply generated correlation of estimation errors (*c*_12_ = 0.15) using 10,000 overlapped samples in exposure and outcome studies (more results for different *c*_12_ are given in SI Appendix, Fig. S3). We notice that correlation between estimation errors can also be induced by population stratification and cryptic relatedness. To avoid unrealistic simulation of population stratification, we investigated this issue when we performed real data analysis. The quantile-quantile plots of different MR methods are shown in Fig. 2C, 2D, 2F. IVW, dIVW, RAPS, MRMix, cML-MA, and Weighted-median produced overly inflated *p*-values. These results indicate that correlation between estimator errors can be a major confounding factor leading to false positive findings. Again, CAUSE produced very deflated *p*-values. To see how MR-APSS accounts for correlation between estimation errors, we set **C** = **I**, i.e., *c*_1_ = *c*_2_ = 1 and *c*_12_ = 0. In such a way, MR-APSS was forced to ignore the correlation between estimation errors. We denote this version of MR-APSS as MR-APSS (**C** = **I**). As shown in Fig. 2F, MR-APSS (**C** = **I**) produced inflated *p*-values. In contrast, MR-APSS produced well-calibrated *p*-values. These results suggest that MR-APSS can satisfactorily account for correlation between estimation errors due to sample structure.

Finally, we examined the power of MR methods. As shown above, IVW, dIVW, RAPS, MRMix, cML-MA, and Weighted-median often produced overly inflated type I errors in the presence of either pleiotropy or sample structure. Hence, we only compared MR-APSS with Egger, Weighted-mode, and CAUSE. We simulated data with both genetic correlation (*r_g_* = 0.1) and correlation between estimation error (*c*_12_ = 0.1). We varied the causal effect size *β* from 0.05 to 0.45. MR-APSS was the overall winner in terms of power (Fig. 2 G). We further compared the estimation accuracy of the causal effects using MR-APSS, Egger, Weighted-mode, and CAUSE (Fig. 2 H). Consistent with the literature [29], we observed that Egger had a very large estimation error. As discussed in SI Appendix, section 2.4, CAUSE often misinterprets the causal effect as correlated pleiotropy, leading to underestimation of the true causal effect. Consistently, we observed that the estimate of Weighted-mode and CAUSE was biased to the null (*β* = 0). In the above simulations, the foreground-background variance ratio was fixed at *σ* : *σ_u_* = 20 : 1. We provide more results with different foreground-background variance ratios (*σ* : *σ_u_* ∈ {40, 10}) in SI Appendix, Figs. S4 and S5.

To evaluate the robustness of MR-APSS in the presence of model misspecification, we also conducted simulations with the CAUSE model. The main patterns of the performance of the ten MR methods largely remained the same. We provide details in SI Appendix, section 3.2, Figs. S6-S8.

### Real data analysis: negative control outcomes

To fairly examine the type I errors of MR methods, we use the negative control outcomes proposed by Sanderson et al. [9], where confounding factors (e.g., pleiotropy and sample structure) naturally exist. The traits that can serve as ideal negative control outcomes should satisfy two conditions. First, they should not be causally affected by any of the exposures considered. Second, the exposure and outcome traits could be affected by some unmeasured confounders, e.g., population stratification. Following the same way of Sanderson et al. [9] to choose negative control outcomes, we considered natural hair colors before greying (Hair color: black, Hair color: blonde, Hair color: light brown, Hair color: dark brown) and skin tanning ability (Tanning) from UKBB because they are largely determined at birth and they could be affected by sample structure.

We considered 26 exposure traits from UKBB and Genomics Consortiums (Details for the GWAS sources are given in SI Appendix, Table S2). These traits can be roughly divided into five categories, including psychiatric/neurological disorders, social traits, anthropometric traits, cardiometabolic traits, and immune-related traits. The data pre-processing steps for GWAS summary statistics are described in SI Appendix, section 4.1. The sample sizes of those GWASs range from 114,244 to 385,603, with a minimum of 15,954 for ASD and a maximum of 898,130 for T2D. Given the large sample sizes of GWASs, we used the genome-wide significance threshold 5 × 10^−8^ as the IV threshold for IVW, dIVW, RAPS, Egger, MRMix, CML-MA, Weighted-median, and Weighted-mode in real data analysis. This stringent criterion helps to exclude invalid IVs for these methods and thus reduce their false positive rates. Due to the stringent IV selection, we were not able to find enough SNPs (> 4) as IVs for four exposure traits, i.e., major depressive disorder (MDD), autism spectrum disorder (ASD), subject well-being (SWB), and the number of children Ever born (NEB). For CAUSE [25], we used its default *p*-value threshold *p* =1 × 10^−3^ to select IVs. For MR-APSS, we used 5 × 10^−5^ as the default IV threshold.

First, we applied MR-APSS and the nine summary-level MR methods to infer the causal effects between these 26 exposure traits and five negative control outcomes. To make the comparison fair, we focus on the results for 110 pairs where each method had sufficient IVs for MR analysis. Ideally, these *p*-values should be uniformly distributed between 0 and 1 under the null (*β* = 0). Fig. 3A shows the QQ-plots of – log_10_(*p*) values of the six methods (red dots).Clearly, MR-APSS and Weighted-mode produced well-calibrated *p*-values. IVW, dIVW, RAPS, MRMix, cML-MA, and Weighted-median produced overly inflated *p*-values, while Egger produced slightly inflated *p*-values. CAUSE produced deflated *p*-values in the beginning but inflated *p*-values later. We investigated the reasons why the five MR methods performed unsatisfactorily. As shown in Fig. 3B, we examined the estimates of two key parameters, *r_g_* and ci2, of our background model, where **r_g_** is the genetic correlation capturing the overall correlated pleiotropic effects and *c*_12_ captures the correlation of estimation errors due to sample structure (e.g., population stratification, cryptic relatedness, and sample overlap). Among the 110 exposure-outcome trait pairs, 81 trait pairs had nearly zero genetic correlation and 29 trait pairs had nonzero genetic correlation at the nominal level of 0.05 (marked by *). We also examined the correlation of estimation errors due to sample structure. Among the 110 trait pairs, 63 pairs had significant nonzero 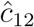 at the nominal level 0.05 (marked by *). To identify the major reason for the inflated *p*-values produced by the nine MR methods, we restricted ourselves to the 81 trait pairs whose genetic correlation was nearly zero. For these 81 pairs, we generated the QQ-plots of – log_10_(*p*) values of the ten MR methods (blue triangles in Fig. 3A). Clearly, IVW, dIVW, RAPS, MRMix, cML-MA, and Weighted-median still produced overly inflated *p*-values. Egger produced slightly better calibrated *p*-values. CAUSE produced deflated *p*-values in the beginning but inflated *p*-values later. We further restricted ourselves to trait pairs whose genetic correlation and correlation of estimation errors were both nearly zero. For these trait pairs (green diamond), MR-APSS, Weighted-mode RAPS, MRMix, cML-MA, Weighted-median, and Egger produced well-calibrated *p*-values. IVW and dIVW still produced inflated *p*-values. CAUSE produced very conservative *p* values. These results suggest that sample structure is another major confounding factor in addition to pleiotropy.

**Figure 3:**
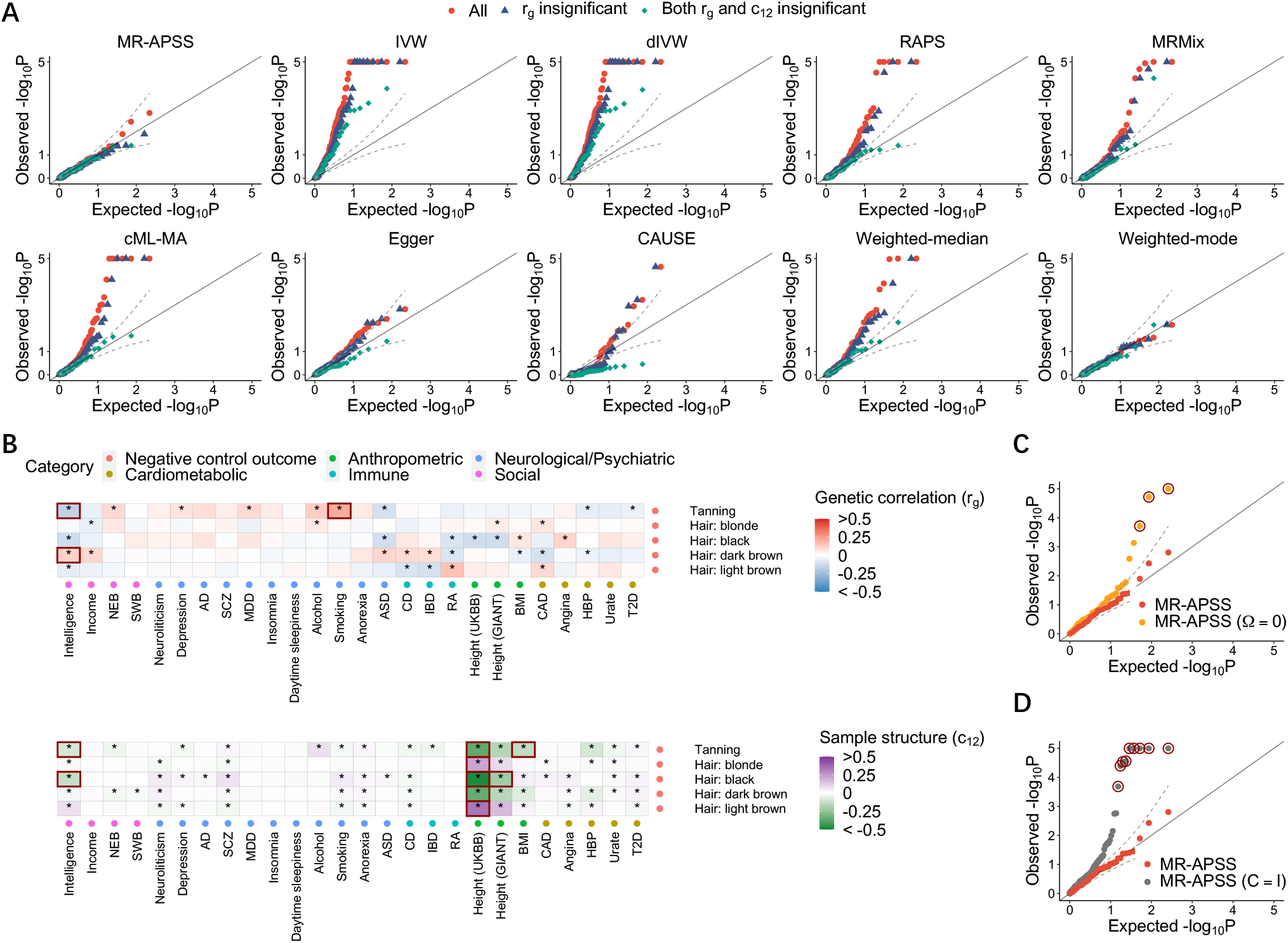
Evaluation of the type I error control of MR methods using negative control outcomes.(**A**) Quantile-quantile plots of – log_10_(*p*)-values from ten summary-level MR methods for causal inference between complex traits and negative control outcome. Red dots represent all 110 trait pairs tested by each method. Blue triangles represent the 81 trait pairs with insignificant genetic correlation at the nominal level of 0.05. Green diamonds represent the 29 trait pairs whose genetic correlation **r_g_** and *c*_12_ are both insignificant at the nominal level of 0.05. (**B**) Estimates of **r_g_** and *c*_12_ for trait pairs between 26 complex traits and five negative control outcomes. (**C**) Quantile-quantile plots of – log_10_(*p*)-values from MR-APSS, MR-APSS (**Ω** = **0**), and MR-APSS (**C** = **I**) for trait pairs between 26 complex traits and five negative control outcomes. The circled *p*-values correspond to the trait pairs marked by squares in (**B**), which are largely confounded by pleiotropy and sample structure.

It is worthwhile to mention that nonzero *c*_12_ can be induced by either population stratification or sample overlap. To see this, let us consider the relationship between Height (GIANT) [30] and Tanning from UKBB. Recall that parameters *c*_1_ and *c*_2_ capture the bias in estimation errors (*ϵ_j_*, *ξ_j_*) and parameter *c*_12_ captures their correlation (Fig. 1). By applying LDSC to estimate our background model, we obtained 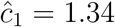 (s.e. = 0.022) for Height (GIANT) and 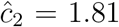 (s.e. = 0.023) for Tanning, respectively. These results indicate that the publicly released GWAS summary statistics are affected by confounding factors, such as population stratification. By applying LDSC, we obtained 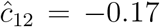 (s.e. = 0.011). As we know, the samples from GIANT do not overlap with UKBB [31]. Therefore, the nonzero 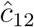 value should be mainly attributed to population stratification. As a comparison, we also considered Height (UKBB) [32] and Tanning from UKBB. By applying LDSC, we obtained 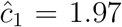 (s.e. = 0.040) for Height (UKBB), suggesting that the released GWAS summary statistics of Height (UKBB) might potentially suffer from population stratification. By applying LDSC, we obtained 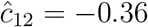 (s.e. = 0.014) for Height (UKBB) and Tanning (UKBB). Such a nonzero value could be attributed to both population stratification and sample overlap.

To better examine the role of MR-APSS in accounting for pleiotropy or sample structure, we applied MR-APSS but fixed **Ω** = **0** and **C** = **I**, respectively. We denote the two variations as MR-APSS (**Ω** = **0**) and MR-APSS (**C** = **I**), where MR-APSS (**Ω** = **0**) does not account for pleiotropy and MR-APSS (**C** = **I**) does not account for sample structure. As shown in 3C, both MR-APSS (**Ω** = **0**) and MR-APSS (**C** = **I**) reported inflated *p*-values. For example, based on Bonferroni correction, several trait pairs (marked with black circles in Fig. 3C) were falsely detected as causal by MR-APSS (**Ω** = **0**) and MR-APSS (**C** = **I**). As shown in Fig. 3B (marked by squares), their corresponding 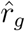 and 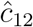 values were significantly different from zero. By using negative control outcomes, we show that MR-APSS can produce well-calibrated *p*-values by accounting for pleiotropy and sample structure.

### Inferring causal relationships among complex traits

To perform causal inference, we considered 26 complex traits from five categories including psychiatric/neurological disorders, social trait, anthropometric traits, cardiometabolic traits, and immune-related traits. Before applying MR methods, we examined the estimates of **r_g_** and *c*_12_ in the background model of MR-APSS for all 325 pairwise combination of the 26 traits. We found that genetic correlation (**r_g_**) of 198 pairs significantly differed from zero at the nominal level of 0.05 (marked by * in Fig. 4A). Among them, genetic correlation of 130 pairs remained to be significant after Bonferroni correction with *p* ≤ 0.05/325 (marked by ** in Fig. 4A). For the estimates of *c*_12_, 126 pairs had significant nonzero 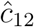 at the nominal level 0.05 (marked by * in Fig. 4B) and 76 pairs of them remained to be significantly different from zero after Bonferroni correction (marked by ** in Fig. 4B). Of note, 56 pairs of traits had significantly nonzero estimates of both 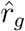 and 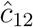 after Bonferroni correction. The above results suggest that both pleiotropy and sample structure are presented as major confounding factors for causal inference.

**Figure 4:**
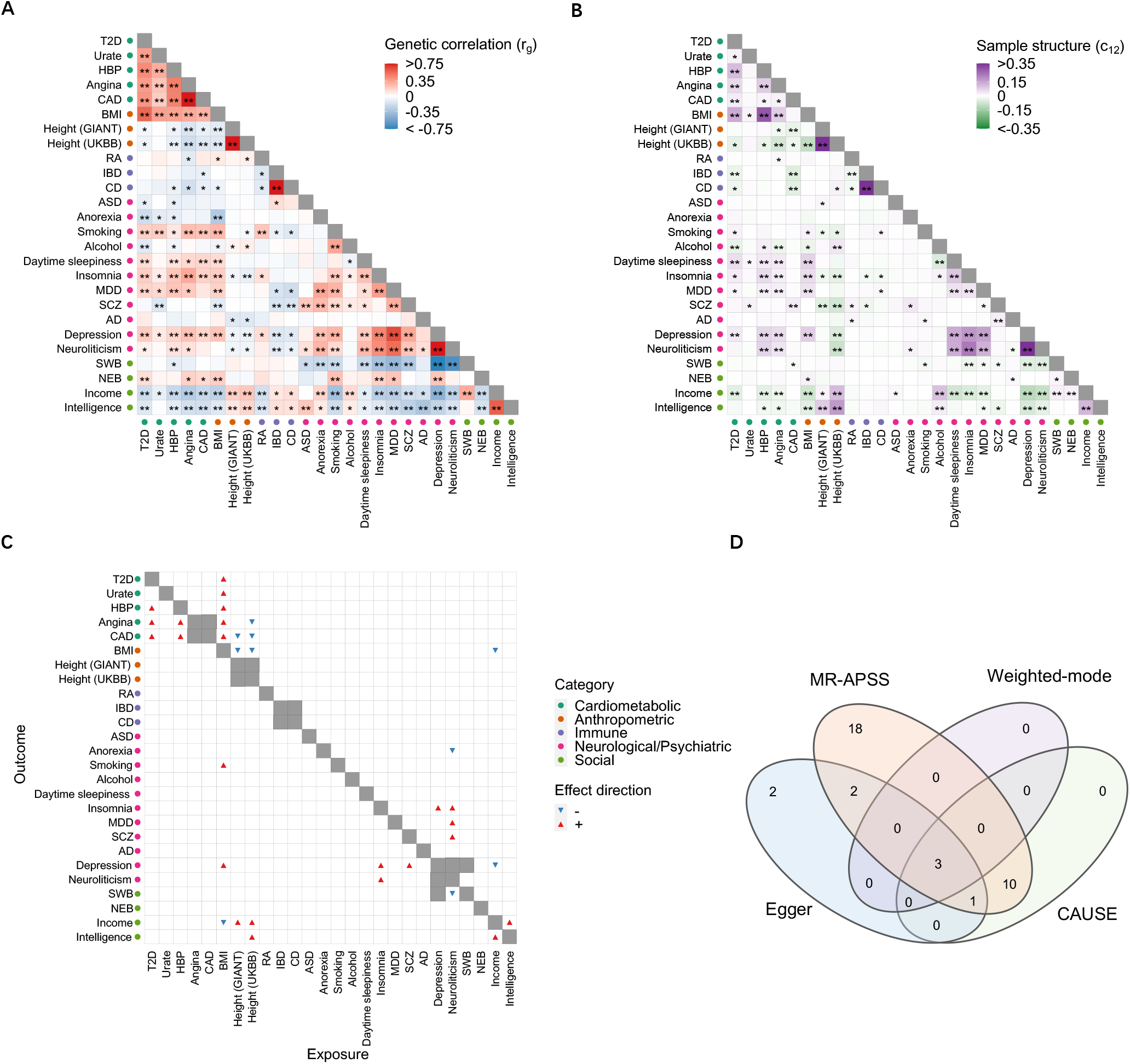
Application of MR-APSS to infer causal relationships between 26 complex traits. (**A**) Estimates of genetic correlation between 26 complex traits. Positive and negative estimates of genetic correlation 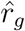 are indicated in red and blue, respectively. Trait pairs with significant 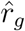 at the nominal level of 0.05 are marked by *. Trait pairs that remain to be significant after Bonferroni correction with *p* ≤ 0.05/325 are marked by **. (**B**) Estimates of *c*_12_ between 26 complex traits. Positive and negative estimates of *c*_12_ are shown in purple and green, respectively. Trait pairs with significant 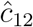 at the nominal level of 0.05 are marked by *. Trait pairs remain to be significant after Bonferroni correction with *p* ≤ 0.05/325 are marked by **. (**C**) Causal relationships detected by MR-APSS. The positive and negative estimates of causal effects of the exposure on the outcome are indicated by red up-pointing triangles and blue down-pointing triangles, respectively. (**D**) The Venn diagram shows the causal effects detected by MR-APSS, CAUSE, Egger, and Weighted-mode after Bonferroni correction.

We considered inferring the causal relationship between traits *X* and *Y* in both directions, i.e., *X* → *Y* (*X* as exposure and *Y* as outcome) and *Y* → *X* (*Y* as exposure and *X* as outcome). To avoid causal inference between two very similar phenotypes (e.g., Angina and CAD), we excluded several trait pairs which were marked in grey color as non-diagonal cells in Fig. 4C. Therefore, 640 trait pairs remained for MR tests in total. We applied MR-APSS to these trait pairs using IV threshold *p* = 5 × 10^−5^ and identified 34 significant causal relationships after Bonferroni correction (Fig. 4C, marked by triangles). As shown in Fig. 4A, many traits in social or neurological/psychiatric categories were observed to be genetically correlated with a wide range of complex traits from different categories. After accounting for pleiotropy and sample structure, the results from MR-APSS indicate that genetic correlation of many trait pairs should not be attributed to the causal effects. An example is Depression which was also genetically correlated with 18 complex traits from different categories, such as BMI (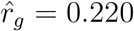, s.e. = 0.024) from Anthropometric category, Insomnia (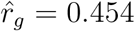, s.e. = 0.025), and SCZ (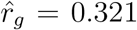, s.e. = 0.027) from neurological/psychiatric category. MR-APSS only confirmed the causal effect of Depression on Insomnia (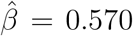, *p*-value = 4.38 × 10^−5^). Clearly, MR-APSS can serve as an effective tool to distinguish causality from genetic correlation.

As a comparison, we also applied the nine compared methods to infer the causal relationships for the 640 trait pairs. We used *p* = 5 × 10^−8^ as the IV selection threshold for IVW, dIVW, RAPS, Egger, MRMix, CML-MA, Weighted-median, and Weighted-mode, and *p* = 1 × 10^−3^ for CAUSE. For MR methods including IVW, dIVW, RAPS, Egger, MRMix, CML-MA, Weighted-median, and Weighted-mode, only 541 trait pairs were tested because 99 trait pairs had less than four SNPs as IVs. For CAUSE, all 640 trait pairs were included. A summary of the causal relationships detected by the nine compared methods are given in SI Appendix, Figs. S22-S30. RAPS reported 58 trait pairs with significant causal effects after Bonferroni correction. Among them, 24 trait pairs were considered insignificant by MR-APSS after Bonferroni correction. Notably, RAPS made a similar assumption with the foreground model of MR-APSS, however, it has no background model to account for pleiotropy and sample structure. To better understand the difference between RAPS and MR-APSS, we applied MR-APSS (**Ω** = **0**) or MR-APSS (**C** = **I**) to those trait pairs. The testing *p*-values of 18 trait pairs became significant based on Bonferroni correction. An example was BMI and Insomnia (SI Appendix, Table S3) with 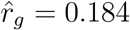 (s.e. = 0.025) and 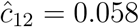 (s.e. = 0.010). RAPS produced 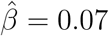 with *p*-value 3.04 × 10^−9^. Without accounting for pleiotropy or sample structure, MR-APSS (**Ω** = **0**) and MR-APSS (**C** = **I**) reported 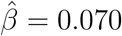 with *p*-value = 1.70 × 10^−7^ and 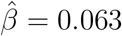 with *p*-value = 1.01 × 10^−4^, respectively. After accounting for both pleiotropy and sample structure, MR-APSS estimated causal effect between BMI and Insomnia as 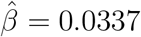 with *p*-value = 0.128. The results indicate that RAPS was likely affected by pleiotropy and sample structure.

Since IVW, dIVW, RAPS, MRMix, cML-MA, Weighted-median tended to have higher type I errors than the nominal level, we mainly compared statistical power of MR-APSS with Egger, CAUSE, and Weighted-mode (Fig. 4D). A complete list of causal relationship among these traits detected by MR-APSS, Egger, CAUSE, and Weighted-mode are summarized in SI Appendix, Table S4. Based on Bonferroni correction, MR-APSS detected 18 significant causal effects which were not reported by CAUSE, Egger, and Weighted-mode, showing higher statistical power of MR-APSS For example, MR-APSS detected significant causal effects of BMI on eight traits. Five of them were supported with evidence of causality from previous literature, including T2D [33], serum urate (Urate) [34], and three cardiovascular diseases (high blood pressure (HBP), Angina and CAD) [35]. For these five supported trait pairs, Egger only detected three significant causal relationships (BMI on CAD, T2D, and HBP), and CAUSE only detected three significant causal relationships (BMI on Urate, HBP, and T2D), and further Weighted-mode detected two significant causal relationships (BMI on T2D; BMI on HBP). In addition to the confirmed findings, MR-APSS detected significant causal effects of BMI on Depression (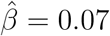, *p*-value = 2.09 × 10^−5^), ever smoked regularly (Smoking) (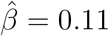, *p*-value = 1.36 × 10^−6^) and Income (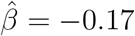, *p*-value = 1.83 × 10^−11^). Those findings are consistent with results from previous MR studies [36, 37, 38], suggesting that being overweight not only increases the risk of depression and tobacco dependence but also suffers from reduced income. Our results also revealed Neuroticism as an important health indicator especially for human psychiatric health. Neuroticism is one of the big five personality traits, characterized by negative emotional states including sadness, moodiness, and emotional instability. Higher neuroticism is associated with premature mortality and a wide range of mental illnesses or psychiatric disorders [31, 39]. There is growing evidence that neuroticism plays a causal role in psychiatric disorders, such as SCZ [40] and MDD [41]. Evidence from MR-APSS also supported the significant causal effect of Neuroticism on SCZ (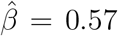, *p*-value = 7.02 × 10^−7^) and MDD (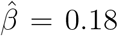, *p*-value = 2.06 × 10^−5^). None of the three methods, CAUSE, Egger and Weighted-mode detected significant causal effects of Neuroticism on MDD or SCZ. MR-APSS also revealed that Neuroticism could be causally linked to Insomnia (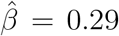, *p*-value = 2.7 × 10^−10^) and Anorexia (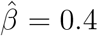, *p*-value = 6.90 × 10^−7^). Weighted-mode and Egger did not report these two cases, and CAUSE only detected a significant causal effect between Neuroticism and Insomnia (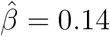, *p*-value = 3.89 × 10^−6^).

### Type I error control and statistical power with different IV thresholds

Existing summary-level MR methods select IVs based on a *p*-value threshold (or an equivalent *t* value). In this section, we would like to highlight the advantages of our method. Regarding the type I error control, our method is insensitive to the choice of threshold. Regarding the improvement of statistical power, our method prefers a loose threshold and we use *p*-value 5 × 10^−5^ as the default setting in real applications. More details regarding the default IV threshold in real applications is given in SI Appendix, section 4.3.

To examine the type I error control of MR-APSS when varying the IV thresholds, we varied the IV threshold from 5 × 10^−8^ to 5 × 10^−5^ when applying MR-APSS to infer the causal relationships between 26 complex traits and the five negative control outcomes. As more IVs involved with a looser IV threshold, the number of invalid IVs increases because they are prone to the violation of MR assumptions. However, most of IVs were detected by MR-APSS as invalid IVs (Fig. 5A). Since MR-APSS only uses the valid instrument strength in the foreground model for causal inference (*Z_j_* = 1), the type I error will not be inflated when more invalid IVs are included. As shown in Fig. 5B, the *p*-values from MR-APSS for trait pairs between 26 complex traits and five negative control outcomes remain well-calibrated at different IV thresholds. These results confirm that the type I error of MR-APSS is insensitive to the IV threshold. It is important to note that correction of the selection bias is a critical step to control type I errors in MR-APSS. Without accounting for the selection bias, the magnitude of the true effect of a selected SNP is largely overestimated and it tends to falsely contribute to the foreground signal (*Z_j_* = 1) for causal inference, thus produces false positives. To verify this, we modified MR-APSS to ignore selection bias and applied this modified version to the same trait pairs with negative control outcomes. Without accounting for the selection bias, the *p*-values produced by the MR-APSS model given in Eq. [6] become inflated (Fig. 5C). When the threshold varies from 5 × 10^−8^ to 5 × 10^−5^, the inflation of *p*-values becomes more severe because more SNPs will falsely contribute to the foreground signal. As a comparison, we ran other summary-level MR methods to the same trait pairs. The QQ-plots are shown in SI Appendix, Fig. S32. Clearly, *p*-values produced by most summary-level MR methods (except Weighted-mode) become more inflated when the IV threshold becomes less stringent.

**Figure 5:**
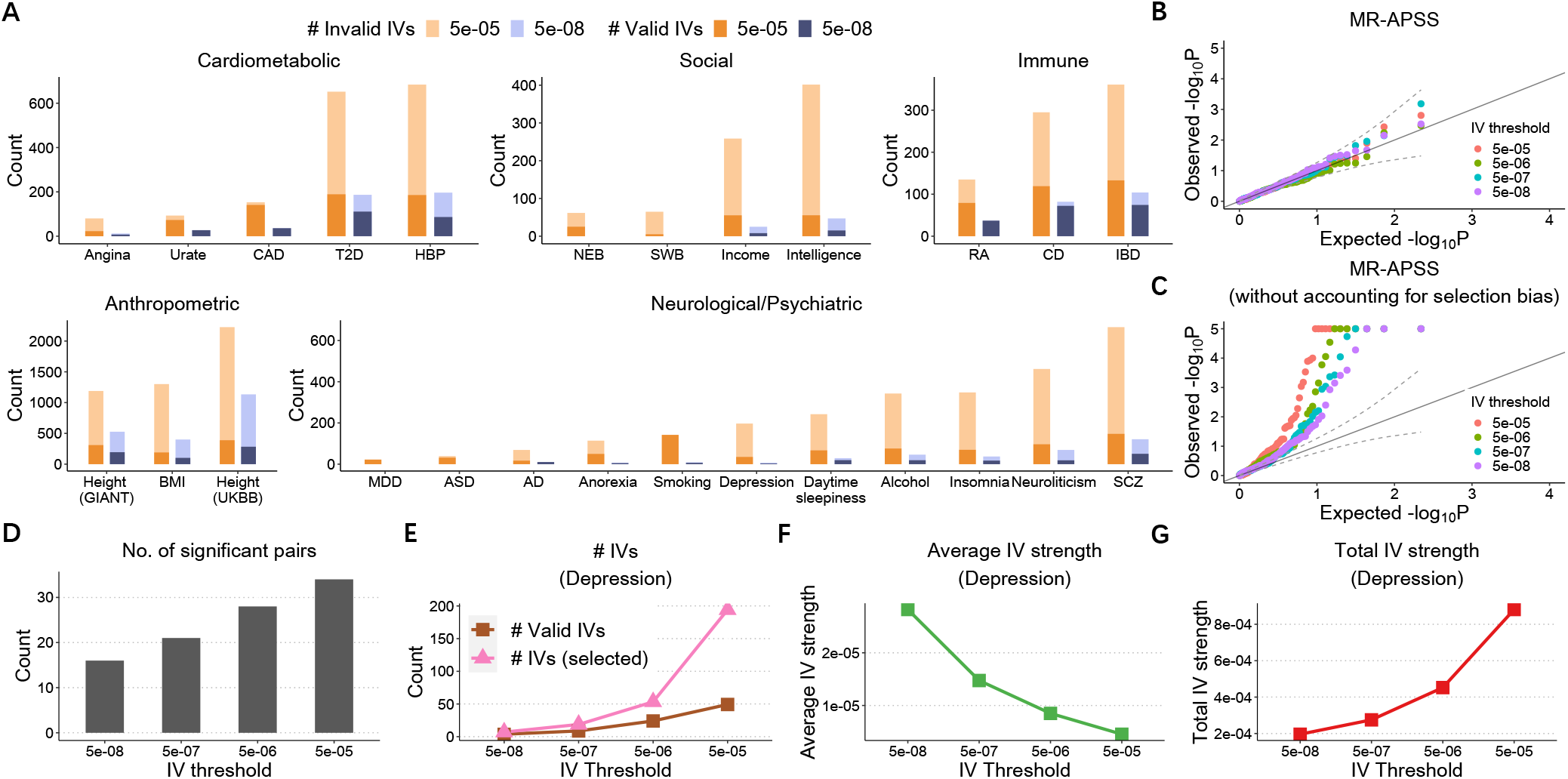
Evaluation of the performance of MR-APSS under different IV selection thresholds. (**A**) The average estimated number of valid IVs (dark color) and invalid IVs (light color) for traits from each category using IV thresholds *p* = 5 × 10^−5^ and *p* = 5 × 10^−8^. (**B**-**C**) Quantilequantile plots of – log_10_(*p*)-values from MR-APSS (**B**) and MR-APSS without accounting for selection bias (**C**) when applied to between 26 complex traits and five negative control outcomes. (**D**) The number of significant trait pairs between 26 complex traits identified by MR-APSS with different IV thresholds. (**E**-**G**): An illustrative examples of exposure: Depression. (**E**) The number of selected IVs *M_t_* at threshold *t* and the estimated number of valid IVs. (**F**) and (**G**) The estimated average and total IV strengths.

As *p*-values of MR-APSS are well-calibrated when the IV threshold varies from 5 × 10^−5^ to 5 × 10^−8^, we can examine the statistical power of MR-APSS with different IV thresholds. We applied MR-APSS to infer the causal relationships among 26 complex traits by varying the IV threshold at 5 × 10^−5^, 5 × 10^−6^, 5 × 10^−7^, and 5 × 10^−8^. In general, we find that the average IV strength (defined in Eq. [11]) decreases with the IV threshold becomes looser, and the total IV strength (defined in Eq. [12]) increases as more IVs are included in the analysis. We provide two concrete examples to illustrate these points (see details in SI Appendix, section 4.2, Fig S14). As a result, the statistical power of MR-APSS can be improved by including SNPs with moderate effects. These results are confirmed in Fig. 5D, where the number of significant pairs identified by MR-APSS increases from 16 to 34 when the IV threshold becomes looser from 5 × 10^−8^ to 5 × 10^−5^.

When investigating the causal relationship among 26 complex traits, the number of valid IVs as well as the total IV strength increased a lot by changing the IV threshold from 5 × 10^−8^ to 5 × 10^−5^ (Fig. 5A). We found that the social and neurological/psychiatric traits can benefit a lot from this property. Despite the large sample sizes for these traits, the number of IVs is too small to perform powerful MR analysis when using the IV threshold *p* = 5 × 10^−8^. For example, Depression only had a very small number of IVs using a stringent IV threshold *p* = 5 × 10^−8^. When the IV thresholds became looser, the number of selected IVs and the number of valid IVs increased a lot (Fig. 5E). Although the average IV strength decreased as IV threshold became looser (Fig. 5F), the total IV strength increased dramatically (Fig. 5G). We also observed that, due to the limited number of IVs using a stringent IV threshold *p* = 5 × 10^−8^, MR-APSS could not detect a significant causal effect of Depression on Insominia (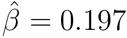, s.e. = 0.214, *p*-value = 0.358). By using a looser IV threshold, MR-APSS detected a significant causal relationship between Depression and Insomnia (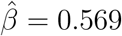, s.e. = 0.139, *p*-value = 4.38 × 10^−5^).

## Discussion

In this paper, we have developed a summary-level MR method, namely MR-APSS, to perform causal inference. To account for the confounding bias due to pleiotropy and sample structure, the background model of MR-APSS inherits the assumptions of LDSC. MR-APSS also assumes the InSIDE condition in the foreground model to infer the causal effect, i.e., *r_f_* = Corr(*γ_j_*, *α_j_*) = 0. In other words, we assume that the association between the exposure and the outcome should be induced by their causal relationship rather than *r_f_* after accounting for confounding factors (e.g., correlated pleiotropy and sample structure) in the background model. Although our method relies on this assumption to infer the causal effect, we can empirically check the influence of this assumption via the following sensitivity analysis. Specifically, we can evaluate how the estimated causal effect 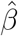 changes when Corr(*γ_j_, α_j_*) varies. In this way, users can obtain useful information about their inferred causal relationship under the perturbation of assumptions. We provide more details on sensitivity analysis in SI appendix, section 1.5, Fig. S13.

Besides the development of summary-level MR methods, we are aware of recent developments of individual-level MR methods, including sisVIVE [42], TSHT [43], GENIUS [44], GENIUS-MAWII [45], and MR-MiSTERI [46]. We believe that summary-level MR methods and individual-level MR methods are complementary to each other. On the one hand, summary-level methods relying on linear models only require marginal estimates and their standard errors. Therefore, they are widely applicable to screen causal relationship between an exposure and an outcome. This is important because the access to individual-level data may be restricted due to privacy protection [47]. On the other hand, individual-level methods can be more powerful than summary-level MR methods when individual-level data is accessible. First, individual-level MR methods can allow for a more flexible model to handle nonlinearity in causal inference. We are aware of several nonlinear MR methods using individual-level data [48, 49]. Unlike linear MR methods which approximate a population-averaged causal effect, the nonlinear MR methods estimate the localized average causal effects in each stratum of population using individual-level data. For example, a very recent MR study applies a nonlinear MR method to investigate whether a nonlinear model is a better fit for the relationship between diastolic blood pressure (DBP) and cardiovascular disease (CVD) [50]. Second, individual-level MR methods can utilize more information, which is only available in individual-level GWAS datasets. For example, the individual-level methods, GENIUS [44] and GENIUS-MAWII [45], require heteroscedasticity of the exposure but this kind of information is not available in GWAS summary statistics. We find that GENIUS and GENIUS-MAWII are robust in the presence of pleiotropy and sample structure. The estimation efficiency of GENIUS and GENIUS-MAWII depends on their IV strengths which are related to heteroscedasticity of the exposure. In this regard, GENIUS and GENIUS-MAWII relax classical MR assumptions by requiring heteroscedasticity of the exposure, while MR-APSS relaxes classical MR assumptions by imposing the LDSC assumptions in its background model and the InSIDE condition in its foreground model. Through simulation studies and real data analyses, we find GENIUS, GENIUS-MAWII and MR-APSS are quite complementary to each other. We provide more detailed results in SI Appendix, sections 2.3, 3.3 and 4.4. In summary, we believe that summary-level methods and individual-level MR methods are complementary to each other, and they jointly contribute to the MR literature for causal inference. Summary-level MR methods are often preferred for large-scale screening of causal relationships and individual-level MR methods can provide a closer examination for causal relationships of interest.

Similar to existing summary-level MR-methods, we consider linear models to perform causal inference even for binary traits. To have better interpretation of the causal effect estimates for binary traits, we show that the output from the observed 0-1 scale based on linear models can be transformed to the liability scale based on the probit models. We provide the details in SI Appendix, section 1.7.

Despite the improvement of MR-APSS over many existing MR methods, more research is needed for causal inference with genetic data. First, the background model is proposed to account for pleiotropy and sample structure hidden in GWASs of complex traits. The direct application of this model in some other contexts may not be suitable. For example, it is of great interest to infer the causal relationship between gene expression and complex diseases based on transcriptome-wide Mendelian randomization. However, it remains unclear what kind of signals should be considered as the background signals. The development of new statistical methods for transcriptome-wide Mendelian randomization is highly desirable. Second, multivariate Mendelian randomization (MVMR) is drawing more and more attention [51, 52]. As some risk factors are known to be related to a certain type of disease, it is more interesting to ask what other risk factors can be inferred conditioning on the known ones. We hope that MR-APSS can motivate more researchers to uncover more reliable causal relationships using rich genetic data resources.

## Materials and Methods

### The MR-APSS approach

MR-APSS takes GWAS summary statistics 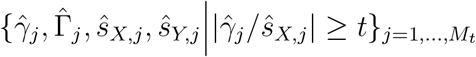 as input to perform causal inference, where 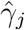 and 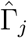 are the estimated *j*-th SNP’s effects on exposure *X* and outcome *Y*, respectively, and 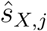, and 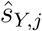 are their standard errors, 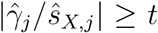 is the selection criterion to ensure that SNP *j* is associated with *X*, and *M_t_* is the number of SNPs selected as IVs using a threshold *t* of *z*-values. To infer the causal effect *β* of exposure *X* on outcome *Y*, we propose to decompose the observed SNP effect sizes into background and foreground signals (Fig. 1):

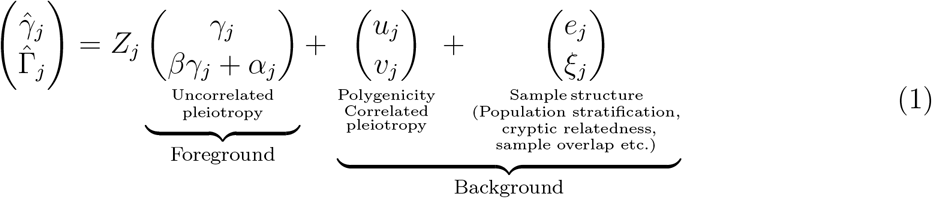

where *u_j_* and *v_j_* are the polygenic effects of SNP *j* on *X* and *Y*, *ϵ_j_* and *ξ_j_* are the estimation errors of SNP effect sizes, *γ_j_* is the remaining SNP effect on exposure *X* as the instrument strength, *α_j_* is the direct SNP effect on outcome *Y*, and *Z_j_* is a Bernoulli variable indicating whether SNP *j* has a foreground component (*Z_j_* = 1) or not (*Z_j_* = 0).

### The background model of MR-APSS

To model polygenic effects and their correlation induced by pleiotropy (Fig. 1b), we assume a variance component model

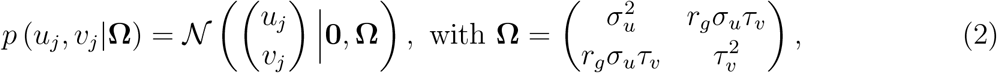

where (*u_j_, v_j_*) are random effects from a bivariate normal distribution with mean vector **0** and covariance matrix **Ω**, *r_g_* is the genetic correlation induced by pleiotropic effects between *X* and *Y*, and 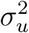 and 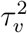 are the variance of polygenic effects on *X* and *Y*, respectively. To account for bias and correlation in estimation errors due to sample structure, we consider the following model:

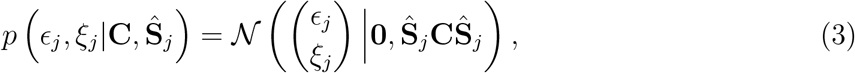

where 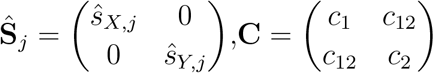, and the parameters *c*_1_ and *c*_2_ are used to adjust the bias in estimator errors and *c*_12_ accounts for the correlation between the estimation errors. In the presence of population stratification and cryptic relatedness, *c*_1_ and *c*_2_ will deviate from one (typically larger than one). Moreover, either population stratification or sample overlap can induce covariance between the estimation errors, resulting in nonzero *c*_12_.

Under the assumptions of LDSC [16], we can exploit the LD structure of human genome to account for confounding factors in the background model. Let 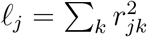 be the LD score of SNP *j*, where *r_jk_* is the correlation between SNP *j* and SNP *k*. The key idea to adjust LD effects is based on the fact: the true genetic effects are tagged by LD while the influence of sample structure is uncorrelated with LD. Then we show that our background model (*Z_j_* = 0) can be written as (see SI Appendix, section 1.1)

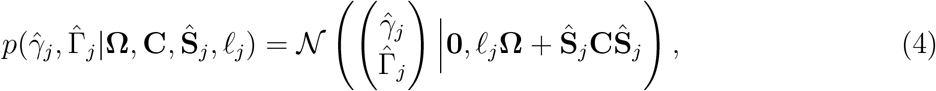

where pleiotropy and sample structure are captured by the first-order and zero-order terms of LD score, respectively. Therefore, both **Ω** and **C** in the background model are pre-estimated by LDSC using genome-wide summary statistics (see SI Appendix, section 1.4.1). As observed in real data analysis, pleiotropy and sample structure are two major confounding factors for causal inference. We provide more discussion about the asymptotic distribution of summary statistics after principal component adjustment in SI Appendix, section 1.9.

### The foreground model of MR-APSS

By accounting for confounding factors using the background model, we only need three mild assumptions on instrument strength *γ_j_* and direct effect *α_j_* to infer causal effect *β*, as shown in Fig. 1(a). First, there exist some nonzero values in {*γ_j_*}_*j*=1, …, *M_t_*_. Second, the strengths of instruments {*γ_j_*}_*j*=1, …, *M_t_*_ are independent of confounding factors. Third, the instrument strengths are independent of the direct effects (InSIDE condition), i.e., 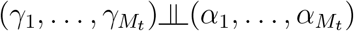. Although our assumptions seem similar to those of existing methods, they are only imposed to the foreground signal and thus they are much weaker than existing MR methods. Specifically, we assume that *γ_j_* and *α_j_* are normally distributed and independent of each other:

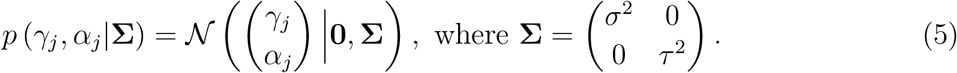

### The foreground-background model of MR-APSS

Now we combine the background model and the foreground model to characterize the observed SNP effect sizes 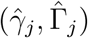. Let π_0_ = *p*(*Z_j_* = 1) be the probability that SNP *j* carries the foreground signal. Combining Eqs. [1,2,3,5] and integrating out *γ_j_*, *α_j_*, *u_j_*, *v_j_*, *ϵ_j_*, *ξ_j_*, and *Z_j_*, we have the following probabilistic model:

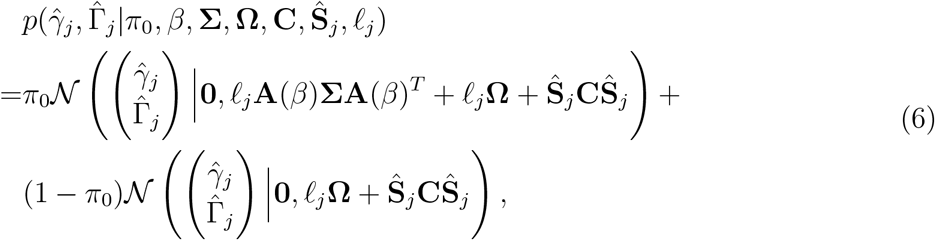

where 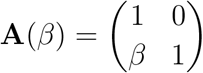. A detailed derivation for Eq. [6] is given in SI Appendix, section 1.2. The theoretical justification of the uniformity of the approximated distribution for 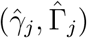 in Eq. [6] for *j* = 1, …, *M_t_* is given in SI Appendix, section 1.8.

### Accounting for selection bias in MR-APSS

Recall that SNPs are selected based on a *p*-value threshold or equivalently a threshold *t* of *z*-score, i.e., 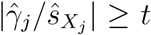. This selection process introduces non-ignorable bias, i.e., 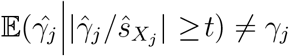, which has been known as winner’s curse in GWAS [53, 28]. To correct the selection bias in MR, we further take into account the selection condition 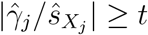. After some derivations (SI Appendix, section 1.3), model (6) becomes a mixture of truncated normal distributions:

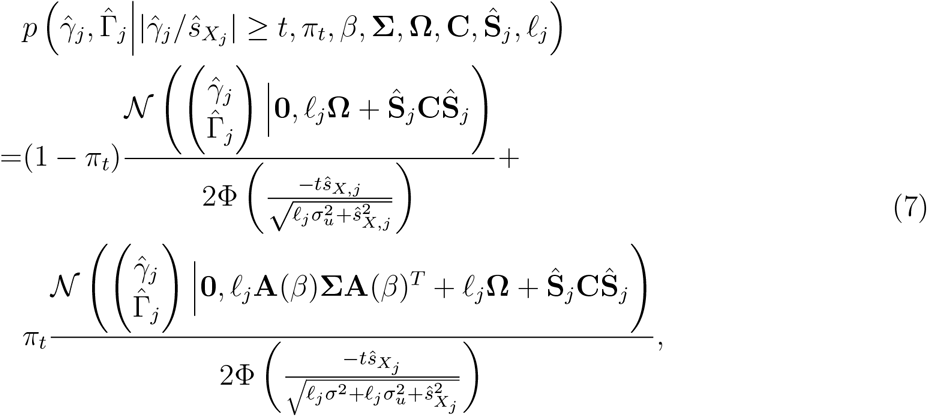

where 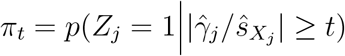 is the probability that the *j*-th SNP carries the foreground signal after selection.

### Parameter estimation and statistical inference

In MR-APSS, the parameters of 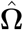 and 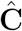 in the background model are estimated by LDSC using genome-wide summary statistics. Given 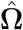 and 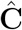, the log-likelihood function of the observed data 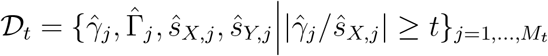 can be written as:

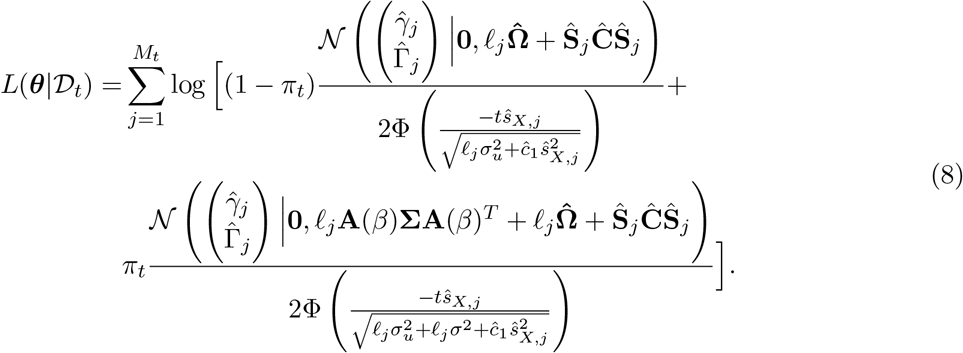

To obtain the maximum likelihood estimate of model parameters *θ* = {*β*, *π_t_*, **Σ**}, we then derive an efficient expectation-maximization (EM) algorithm (see details in SI Appendix, section 1.4.2). As a byproduct, we can estimate the numbers of valid IVs and invalid IVs as 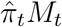 and 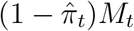, respectively.Real data results of the estimated numbers of valid and invalid IVs are shown in Fig. 5 A. The posterior of SNP *j* serving as a valid IV can be estimated as 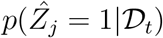, as shown in dark blue in Fig. 1D. The likelihood ratio test can be conducted to examine the existence of the causal effect. Considering the following hypothesis test:

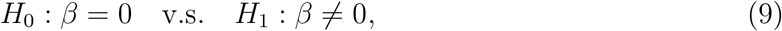

the likelihood-ratio test statistic (LRT) is given by

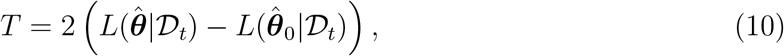

where 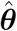 and 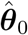 are the parameter estimates obtained under hypotheses *H*_1_ and *H*_0_, respectively. Under the null hypothesis *H*_0_, the test statistic *T* is asymptotically distributed as 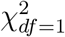 and its *p*-value can be obtained accordingly.

### IV strength

The performance of MR methods depend on the instrument strength. For MR-APSS, we define

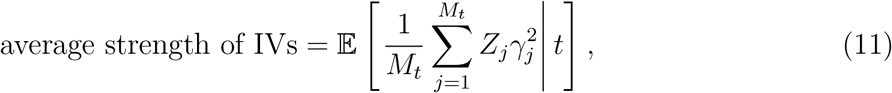

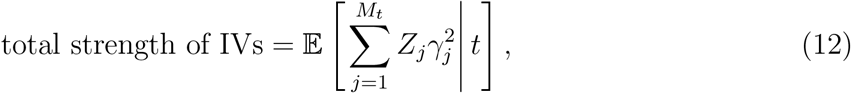

which measure the average/total IV strength for those *M_t_* SNPs with the selection criterion 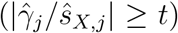. Given the observed summary statistics and the selection criterion *t*, we can use MR-APSS to obtain the posterior distributions of (*γ_j_*, *Z_j_*). Therefore, we can obtain the estimates of average IV strength and total IV strength defined in Eq. [11] and Eq. [12]. According to the above definitions, the average and total IV strengths depend on both the IV threshold and sample size. In general, we find that the average IV strength decreases when the IV threshold becomes looser, and the total IV strength increases as more IVs are included in the analysis. Our definitions of IV strengths for the MR-APSS model are closely connected to the IV strengths defined in MR literature (see details in SI Appendix, section 2.5).

## Data availability

All the GWAS summary statistics used in this paper are public available. The URLs for downloading the datasets are summarized in SI Appendix, Table S2.

## Code availability

The MR-APSS software is available at https://github.com/YangLabHKUST/MR-APSS.

## Acknowledgements

We thank the editor and two anonymous reviewers for their very detailed and constructive comments, which have greatly helped to improve our manuscript. We also thank Prof. Lin S. Chen, Prof. Lan Wang, Prof. Baolin Wu, Prof. Zhigang Bao, and Prof. Dong Xia for their helpful comments and insightful discussions. This work is supported in part by Hong Kong Research Grant Council [16307818, 16301419, 16308120, 12303618, 24301419, 14301120], Hong Kong Innovation and Technology Fund [PRP/029/19FX], the Hong Kong University of Science and Technology [startup grant R9405, Z0428 from the Big Data Institute], the Chinese University of Hong Kong direct grants [4053360, 4053423], the Chinese University of Hong Kong startup grant [4930181], the Chinese University of Hong Kong’s Project Impact Enhancement Fund (PIEF) and Science Faculty’s Collaborative Research Impact Matching Scheme (CRIMS), Key-Area Research and Development Program of Guangdong Province [2020B0101350001], and the Open Research Fund from Shenzhen Research Institute of Big Data [2019ORF01004]. The computational task for this work was partially performed using the X-GPU cluster supported by the RGC Collaborative Research Fund: C6021-19EF.

## References

[1] Kenneth J Rothman and Sander Greenland. Causation and causal inference in epidemiology. American journal of public health, 95(S1):S144–S150, 2005.

[2] Lars Bondemark and Sabine Ruf. Randomized controlled trial: the gold standard or an unobtainable fallacy? European Journal of Orthodontics, 37(5):457–461, 2015.

[3] George Davey Smith and Shah Ebrahim. ‘Mendelian randomization’: can genetic epidemiology contribute to understanding environmental determinants of disease? International Journal of Epidemiology, 32(1):1–22, 2003.

[4] George Davey Smith and Gibran Hemani. Mendelian randomization: genetic anchors for causal inference in epidemiological studies. Human Molecular Genetics, 23, 2014.

[5] Jean-Baptiste Pingault, Paul F O’reilly, Tabea Schoeler, George B Ploubidis, Frühling Rijsdijk, and Frank Dudbridge. Using genetic data to strengthen causal inference in observational research. Nature Reviews Genetics, 19(9):566–580, 2018.

[6] Stephen Burgess, Adam Butterworth, and Simon G Thompson. Mendelian randomization analysis with multiple genetic variants using summarized data. Genetic epidemiology, 37(7):658–665, 2013.

[7] Michael Baiocchi, Jing Cheng, and Dylan S Small. Instrumental variable methods for causal inference. Statistics in medicine, 33(13):2297–2340, 2014.

[8] Gibran Hemani, Jack Bowden, and George Davey Smith. Evaluating the potential role of pleiotropy in Mendelian randomization studies. Human Molecular Genetics, 27, 2018.

[9] Eleanor Sanderson, Tom G Richardson, Gibran Hemani, and George Davey Smith. The use of negative control outcomes in Mendelian Randomisation to detect potential population stratification or selection bias. International Journal of Epidemiology, 50(4):1350–1361, 2021.

[10] Hyun Min Kang, Jae Hoon Sul, Susan K Service, Noah A Zaitlen, Sit-yee Kong, Nelson B Freimer, Chiara Sabatti, and Eleazar Eskin. Variance component model to account for sample structure in genome-wide association studies. Nature Genetics, 42(4):348–354, 2010.

[11] Nadia Solovieff, Chris Cotsapas, Phil H Lee, Shaun M Purcell, and Jordan W Smoller. Pleiotropy in complex traits: challenges and strategies. Nature Reviews Genetics, 2013.

[12] Daniel M. Jordan, Marie Verbanck, and Ron Do. The Landscape of Pervasive Horizontal Pleiotropy in Human Genetic Variation is Driven by Extreme Polygenicity of Human Traits and Diseases. SSRN Electronic Journal, 2018.

[13] Stephen Burgess, Neil M. Davies, and Simon G. Thompson. Bias due to participant overlap in two-sample Mendelian randomization. Genetic Epidemiology, 40(7), 2016.

[14] Alkes L Price, Nick J Patterson, Robert M Plenge, Michael E Weinblatt, Nancy A Shadick, and David Reich. Principal components analysis corrects for stratification in genome-wide association studies. Nature Genetics, 38(8):904–909, 2006.

[15] Po-Ru Loh, George Tucker, Brendan K Bulik-Sullivan, Bjarni J Vilhjálmsson, Hilary K Finucane, Rany M Salem, Daniel I Chasman, Paul M Ridker, Benjamin M Neale, Bonnie Berger, Nick Patterson, and Alkes L Price. Efficient Bayesian mixed-model analysis increases association power in large cohorts. Nature Genetics, 47(3):284–290, 2015.

[16] Brendan K Bulik-Sullivan, Po-Ru Loh, Hilary K Finucane, Stephan Ripke, Jian Yang, Nick Patterson, Mark J Daly, Alkes L Price, and Benjamin M Neale. LD Score regression distinguishes confounding from polygenicity in genome-wide association studies. Nature genetics, 47(3):291–295, 2015.

[17] Vanessa Didelez and Nuala Sheehan. Mendelian randomization as an instrumental variable approach to causal inference. Statistical methods in medical research, 16(4):309–330, 2007.

[18] Jack Bowden, George Davey Smith, and Stephen Burgess. Mendelian randomization with invalid instruments: effect estimation and bias detection through Egger regression. International journal of epidemiology, 44(2):512–525, 2015.

[19] Qingyuan Zhao, Jingshu Wang, Gibran Hemani, Jack Bowden, and Dylan S Small. Statistical inference in two-sample summary-data mendelian randomization using robust adjusted profile score. The Annals of Statistics, 48(3):1742–1769, 2020.

[20] Ting Ye, Jun Shao, and Hyunseung Kang. Debiased inverse-variance weighted estimator in two-sample summary-data mendelian randomization. The Annals of Statistics, 49(4):2079–2100, 2021.

[21] Jack Bowden, George Davey Smith, Philip C Haycock, and Stephen Burgess. Consistent estimation in mendelian randomization with some invalid instruments using a weighted median estimator. Genetic epidemiology, 40(4):304–314, 2016.

[22] Fernando Pires Hartwig, George Davey Smith, and Jack Bowden. Robust inference in summary data mendelian randomization via the zero modal pleiotropy assumption. International Journal of Epidemiology, (6):6, 2017.

[23] Guanghao Qi and Nilanjan Chatterjee. Mendelian Randomization Analysis Using Mixture Models (MRMix) for genetic effect-size-distribution leads to robust estimation of causal effects. bioRxiv, page 367821, 2018.

[24] Haoran Xue, Xiaotong Shen, and Wei Pan. Constrained maximum likelihood-based Mendelian randomization robust to both correlated and uncorrelated pleiotropic effects. The American Journal of Human Genetics, 108(7):1251–1269, 2021.

[25] Jean Morrison, Nicholas Knoblauch, Joseph H. Marcus, Matthew Stephens, and Xin He. Mendelian randomization accounting for correlated and uncorrelated pleiotropic effects using genome-wide summary statistics. Nature Genetics, 2020.

[26] Abdel Abdellaoui, David Hugh-Jones, Loic Yengo, Kathryn E. Kemper, and Peter M. Visscher. Genetic correlates of social stratification in great britain. Nature Human Behaviour, 3(12), 2019.

[27] Simon Haworth, Ruth Mitchell, Laura Corbin, Kaitlin H. Wade, Tom Dudding, Ashley Budu-Aggrey, David Carslake, Gibran Hemani, Lavinia Paternoster, and George Davey and Smith. Apparent latent structure within the UK Biobank sample has implications for epidemiological analysis. Nature Communications, 10(1), 2019.

[28] Sebastian Zollner and Jonathan K Pritchard. Overcoming the winner’s curse: Estimating penetrance parameters from case-control data. American Journal of Human Genetics, 80(4):605–615, 2007.

[29] Stephen Burgess and Simon G. Thompson. Interpreting findings from Mendelian randomization using the MR-Egger method. European Journal of Epidemiology, 2017.

[30] Andrew R Wood, Tonu Esko, Jian Yang, Sailaja Vedantam, Tune H Pers, Stefan Gustafsson, Audrey Y Chu, Karol Estrada, Zoltán Kutalik, Najaf Amin, et al. Defining the role of common variation in the genomic and biological architecture of adult human height. Nature genetics, 46(11):1173–1186, 2014.

[31] Loic Yengo, Julia Sidorenko, Kathryn E Kemper, Zhili Zheng, Andrew R Wood, Michael N Weedon, Timothy M Frayling, Joel Hirschhorn, Jian Yang, Peter M Visscher, and the GIANT Consortium. Meta-analysis of genome-wide association studies for height and body mass index in 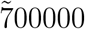 individuals of European ancestry. Human Molecular Genetics, 27(20):3641–3649, 08 2018.

[32] Kyoko Watanabe, Sven Stringer, Oleksandr Frei, Maša UmićevićMirkov, Christiaan de Leeuw, Tinca J. C. Polderman, Sophie van der Sluis, Ole A. Andreassen, Benjamin M. Neale, and Danielle Posthuma. A global overview of pleiotropy and genetic architecture in complex traits. Nature Genetics, 51(9):1339–1348, 2019.

[33] Michael L Ganz, Neil Wintfeld, Qian Li, Veronica Alas, Jakob Langer, and Mette Hammer. The association of body mass index with the risk of type 2 diabetes: a case–control study nested in an electronic health records system in the united states. Diabetology and Metabolic Syndrome, 6(1):50, 2014.

[34] Kentaro Tanaka, Soshiro Ogata, Haruka Tanaka, Kayoko Omura, Chika Honda, Osaka Twin Research Group, and Kazuo Hayakawa. The relationship between body mass index and uric acid: a study on japanese adult twins. Environmental health and preventive medicine, 20(5):347–353, 09 2015.

[35] Sadiya S. Khan, Hongyan Ning, John T. Wilkins, Norrina Allen, Mercedes Carnethon, Jarett D. Berry, Ranya N. Sweis, and Donald M. Lloyd-Jones. Association of Body Mass Index With Lifetime Risk of Cardiovascular Disease and Compression of Morbidity. JAMA Cardiology, 2018.

[36] Amy E Taylor, Rebecca C Richmond, Teemu Palviainen, Anu Loukola, Robyn E Wootton, Jaakko Kaprio, Caroline L Relton, George Davey Smith, and Marcus R Munafò. The effect of body mass index on smoking behaviour and nicotine metabolism: a Mendelian randomization study. Human molecular genetics, 28(8):1322–1330, 2019.

[37] Jessica Tyrrell, Anwar Mulugeta, Andrew R Wood, Ang Zhou, Robin N Beaumont, Marcus A Tuke, Samuel E Jones, Katherine S Ruth, Hanieh Yaghootkar, Seth Sharp, et al. Using genetics to understand the causal influence of higher bmi on depression. International journal of epidemiology, 48(3):834–848, 2019.

[38] Jessica Tyrrell, Samuel E Jones, Robin Beaumont, Christina M Astley, Rebecca Lovell, Hanieh Yaghootkar, Marcus Tuke, Katherine S Ruth, Rachel M Freathy, and Joel N Hirschhorn. Height, body mass index, and socioeconomic status: mendelian randomisation study in UK Biobank. Bmj British Medical Journal, page i582, 2016.

[39] Lahey and B. Benjamin. Public health significance of neuroticism. American Psychologist, 64(4):241, 2009.

[40] Jim Van Os and Peter B. Jones. Neuroticism as a risk factor for schizophrenia. Psychological Medicine, 31(6):1129–34, 2001.

[41] Anne Farmer, Kate Redman, Tanya Harris, Arshad Mahmood, Stephanie Sadler, Andrea Pickering, and Peter McGuffin. Neuroticism, extraversion, life events and depression: The Cardiff Depression Study. The British Journal of Psychiatry, 181(2):118–122, 2002.

[42] Hyunseung Kang, Anru Zhang, T Tony Cai, and Dylan S Small. Instrumental variables estimation with some invalid instruments and its application to mendelian randomization. Journal of the American statistical Association, 111(513):132–144, 2016.

[43] Zijian Guo, Hyunseung Kang, T Tony Cai, and Dylan S Small. Confidence intervals for causal effects with invalid instruments by using two-stage hard thresholding with voting. Journal of the Royal Statistical Society: Series B (Statistical Methodology), 80(4):793–815, 2018.

[44] Eric Tchetgen Tchetgen, BaoLuo Sun, and Stefan Walter. The GENIUS approach to robust Mendelian randomization inference. Statistical Science, 36(3):443–464, 2021.

[45] Ting Ye, Zhonghua Liu, Baoluo Sun, and Eric Tchetgen Tchetgen. GENIUS-MAWII: For Robust Mendelian Randomization with Many Weak Invalid Instruments. arXiv preprint arXiv:2107.06238, 2021.

[46] Zhonghua Liu, Ting Ye, Baoluo Sun, Mary Schooling, and Eric Tchetgen Tchetgen. On Mendelian Randomization Mixed-Scale Treatment Effect Robust Identification (MR MiSTERI) and Estimation for Causal Inference. arXiv preprint arXiv:2009.14484, 2020.

[47] Xiang Zhu and Matthew Stephens. Bayesian large-scale multiple regression with summary statistics from genome-wide association studies. The Annals of Applied Statistics, 11(3):1561–1592, 2017.

[48] Stephen Burgess, Neil M Davies, and Simon G Thompson. Instrumental variable analysis with a nonlinear exposure–outcome relationship. Epidemiology (Cambridge, Mass.), 25(6):877, 2014.

[49] James R Staley and Stephen Burgess. Semiparametric methods for estimation of a nonlinear exposure-outcome relationship using instrumental variables with application to mendelian randomization. Genetic epidemiology, 41(4):341–352, 2017.

[50] Marios Arvanitis, Guanghao Qi, Deepak L Bhatt, Wendy S Post, Nilanjan Chatterjee, Alexis Battle, and John W McEvoy. Linear and nonlinear mendelian randomization analyses of the association between diastolic blood pressure and cardiovascular events: the J-curve revisited. Circulation, 143(9):895–906, 2021.

[51] Eleanor Sanderson, George Davey Smith, Frank Windmeijer, and Jack Bowden. An examination of multivariable Mendelian randomization in the single-sample and two-sample summary data settings. International journal of epidemiology, 48(3):713–727, 2019.

[52] Verena Zuber, Johanna Maria Colijn, Caroline Klaver, and Stephen Burgess. Selecting likely causal risk factors from high-throughput experiments using multivariable mendelian randomization. Nature communications, 11(1):1–11, 2020.

[53] John Ferguson, Judy H Cho, Can Yang, and Hongyu Zhao. Empirical Bayes Correction for the Winner’s Curse in Genetic Association Studies. Genetic Epidemiology, 37(1):60–68, 2013.

