## Supplementary notes, figures and tables for "Mendelian Randomization for causal inference accounting for pleiotropy and sample structure using genome-wide summary statistics"

4.4 Evaluation of the performance of individual-level MR methods in real data analysis 48
**Figure** S8: Type I error control of 14 MR methods on inferring causal effects under CAUSE model in the presence of sample structure under CAUSE model. 57
**Figure** S23: Causal relationships between 26 complex traits detected by dIVW. . 72
**Figure** S24: Causal relationships between 26 complex traits detected by RAPS. . 73
**Figure** S25: Causal relationships between 26 complex traits detected by Egger. . 74
**Figure** S26: Causal relationships between 26 complex traits detected by MRMix. 75
**Figure** S27: Causal relationships between 26 complex traits detected by cML-MA. 76
**Table** S4: The 36 trait pairs detected by MR-APSS, Egger, CAUSE or Weighted-mode 86

### SI Materials and Methods

### 1 The MR-APSS approach

#### 1.1 Derivation of the background model of MR-APSS

Let $(\hat{\gamma}_j, \hat{\Gamma}_j)$ be the GWAS estimates of SNP $j$ for exposure $X$ and outcome $Y$. Under the assumptions of LDSC, we will derive that the background model can be written as

$$p\left(\hat{\gamma}_j, \hat{\Gamma}_j | \mathbf{\Omega}, \mathbf{C}, \hat{\mathbf{S}}_j, \ell_j\right) = \mathcal{N}\left(\begin{pmatrix}\hat{\gamma}_j \\ \hat{\Gamma}_j\end{pmatrix} \middle| \mathbf{0}, \ell_j \mathbf{\Omega} + \hat{\mathbf{S}}_j \mathbf{C} \hat{\mathbf{S}}_j\right), \quad \text{(S.1)}$$

where $\mathbf{\Omega}$ is the variance component of polygenic effects $(u_j, v_j)$, $\ell_j = \sum_k r_{jk}^2$ is the LD score of SNP $j$, $r_{jk}$ is the correlation between SNP $j$ and SNP $k$, $\hat{\mathbf{S}}_j = \begin{pmatrix}\hat{s}_{X,j} & 0 \\ 0 & \hat{s}_{Y,j}\end{pmatrix}$, $\mathbf{C} = \begin{pmatrix}c_1 & c_{12} \\ c_{12} & c_2\end{pmatrix}$, and $\hat{\mathbf{S}}_j \mathbf{C} \hat{\mathbf{S}}_j$ is the variance component for the GWAS estimation errors $(\epsilon_j, \xi_j)$ in the presence of sample structure (e.g., population stratification, cryptic relatedness, and sample overlap).

##### 1.1.1 Statistical Model

Let $N_1$ and $N_2$ be the GWAS sample sizes of two studies for exposure $X$ and outcome $Y$. Following LDSC, we consider the population structure by a mixture of two sub-populations (sub-population 1 and sub-population 2) of equal proportion with the following genetic drift model: (i) Let $\mathbf{G}_1 = \{G_{1,ij}\} \in \mathbb{R}^{N_1 \times M}$ and $\mathbf{G}_2 = \{G_{2,ij}\} \in \mathbb{R}^{N_2 \times M}$ be the standardized genotype matrices for exposure $X$ and exposure $Y$, respectively, where $M$ is the number of SNPs in the genome. For individual $i$ in sub-population 1, we have $\mathbb{E}(G_{1,ij} | i \in \text{ sub-pop 1}) = \mathbb{E}(G_{2,ij} | i \in \text{ sub-pop 1}) = f_j$. For individual $i$ in sub-population 2, we have $\mathbb{E}(G_{1,ij} | i \in \text{ sub-pop 2}) = \mathbb{E}(G_{2,ij} | i \in \text{ sub-pop 2}) = -f_j$. We also have $\text{Var}(G_{1,ij}) = \text{Var}(G_{2,ij}) = 1$ for all $j$ because of standardization; (ii) The genetic drift term $f_j \sim N(0, F_{st})$ for all $j$, and $\text{Cov}(f_j, f_k) = 0$ for all $j \neq k$; (iii) Let $\ell_{j,1}$ and $\ell_{j,2}$ be the LD score of SNP $j$ in sub-populations 1 and 2, respectively. We assume $\ell_{j,1} \approx \ell_{j,2} = \ell_j$ for all $j$. This assumption may be questionable when sub-population 1 and sub-population 2 differ a lot (e.g., when they are from different continents). However, as discussed in the LDSC paper [7], this assumption is reasonable when we are interested in modeling population stratification after principal components adjustment in GWAS where samples are from non-admixed populations.

With the above genetic drift model, we consider the following individual-level background model:

$$\mathbf{x} = \mathbf{G}_1 \mathbf{u} + \mathbf{s}_1 + \mathbf{e}_1, \quad \mathbf{y} = \mathbf{G}_2 \mathbf{v} + \mathbf{s}_2 + \mathbf{e}_2, \quad \text{(S.2)}$$

where $\mathbf{x} = \{x_i\}_{i=1,\ldots,N_1}$ is an $N_1 \times 1$ phenotype vector for exposure $X$, $\mathbf{y} = \{y_i\}_{i=1,\ldots,N_2}$ is an $N_2 \times 1$ phenotype vector for outcome $Y$, $\mathbf{u} = \{u_j\}_{j=1,\ldots,M}$ and $\mathbf{v} = \{v_j\}_{j=1,\ldots,M}$ are $M \times 1$ vectors of polygenic effects, $\mathbf{e}_1 = \{e_{1,i}\}_{i=1,\ldots,N_1}$ and $\mathbf{e}_2 = \{e_{2,i}\}_{i=1,\ldots,N_2}$ are the vectors of independent noises, and $\mathbf{s}_1 = \{s_{1,i}\}_{i=1,\ldots,N_1}$ and $\mathbf{s}_2 = \{s_{2,i}\}_{i=1,\ldots,N_2}$ are the environmental stratification terms defined by

$$s_{1,i} = \begin{cases} \sigma_s, & i \in \text{sub-population 1} \\ -\sigma_s, & i \in \text{sub-population 2} \end{cases}, \quad i = 1, \ldots, N_1,$$

and

$$s_{2,i} = \begin{cases} \sigma_s, & i \in \text{sub-population}\,1 \\ -\sigma_s, & i \in \text{sub-population}\,2 \end{cases}, \quad i = 1, \dots, N_2,$$

where $\sigma_s$ is the mean phenotype difference between sub-population 1 and sub-population 2. Please note that the zero-mean assumption on the environmental stratification terms $\mathbf{s}_1$ and $\mathbf{s}_2$ is not required in our model. The background model of MR-APSS can be estimated by LDSC even though the environmental stratification terms have non-zero mean. This is because the influence of population stratification enters our model through the variance term rather than the mean term. We assume random effects to characterize the polygenic effects,

$$\begin{pmatrix} u_j \\ v_j \end{pmatrix} \sim \mathcal{N}\left(\mathbf{0}, \mathbf{\Omega}\right), \text{ where } \mathbf{\Omega} = \begin{pmatrix} \sigma_u^2 & r_g\sigma_u\tau_v \\ r_g\sigma_u\tau_v & \tau_v^2 \end{pmatrix}, \quad j = 1, \dots, M.$$

The noise terms $(\mathbf{e}_1, \mathbf{e}_2)$ are assumed to have expectations $\mathbb{E}[\mathbf{e}_1] = \mathbb{E}[\mathbf{e}_2] = \mathbf{0}$, and variances $\text{Var}(\mathbf{e}_1) = \sigma_{e_1}^2\mathbf{I}$ and $\text{Var}(\mathbf{e}_2) = \sigma_{e_2}^2\mathbf{I}$. Here we set $\sigma_{e_1}^2 = (1 - M\sigma_u^2 - \sigma_s^2)$ and $\sigma_{e_2}^2 = (1 - M\tau_v^2 - \sigma_s^2)$ to assure that phenotype variances equal one, i.e., $M\sigma_u^2 + \sigma_s^2 + \sigma_{e_1}^2 = 1$ and $M\tau_v^2 + \sigma_s^2 + \sigma_{e_2}^2 = 1$. We assume that the noise terms of different samples are independent. To account for correlation due to sample overlapping, the noise terms for $N_s$ overlapped samples are assumed to be correlated, i.e., $\text{Cov}(e_{1,i}, e_{2,i}) = \rho_e$, where $i$ is the index of overlapped samples.

###### 1.1.2 Summary statistics

Let $\mathbf{G}_{1,j}$ and $\mathbf{G}_{2,j}$ be the $j$-th column of the standardized genotype matrices $\mathbf{G}_1$ and $\mathbf{G}_2$. The GWAS estimates for the $j$-th variant $\hat{\gamma}_j$ and $\hat{\Gamma}_j$ can be obtained, respectively by

$$\hat{\gamma}_j = \frac{\mathbf{G}_{1,j}^T\mathbf{x}}{\mathbf{G}_{1,j}^T\mathbf{G}_{1,j}} = \frac{\mathbf{G}_{1,j}^T\mathbf{x}}{N_1}, \quad \hat{\Gamma}_j = \frac{\mathbf{G}_{2,j}^T\mathbf{y}}{\mathbf{G}_{2,j}^T\mathbf{G}_{2,j}} = \frac{\mathbf{G}_{2,j}^T\mathbf{y}}{N_2}. \tag{S.3}$$

Because a single SNP only explains little phenotypic variance due to polygenicity, the standard errors can be well approximated as

$$\hat{s}_{X,j} \approx \frac{1}{\sqrt{N_1}}, \quad \hat{s}_{Y,j} \approx \frac{1}{\sqrt{N_2}}. \tag{S.4}$$

We then calculate the $z$-scores as

$$z_{X,j} \approx \frac{1}{\sqrt{N_1}}\mathbf{G}_{1,j}^T\mathbf{x}, \quad z_{Y,j} \approx \frac{1}{\sqrt{N_2}}\mathbf{G}_{2,j}^T\mathbf{y}.$$

Given the estimates of effect sizes in Eq. [S.3], we have

$$\mathbb{E}(\hat{\gamma}_j|\mathbf{u}) = \mathbb{E}\left(\frac{\mathbf{G}_{1,j}^T(\mathbf{G}_1\mathbf{u} + \mathbf{s}_1 + \mathbf{e}_1)}{N_1}|\mathbf{u}\right) = \frac{\mathbb{E}\left(\mathbf{G}_{1,j}^T\mathbf{G}_1\mathbf{u}\right)}{N_1} = \sum_k r_{jk}u_k,$$

where $r_{jk} = \mathbb{E}(G_{1,ij}G_{1,ik})$ is the correlation between SNP $j$ and SNP $k$. Similarly, we have

$$\mathbb{E}(\hat{\Gamma}_j|\mathbf{v}) = \sum_k r_{jk}v_k.$$

By taking expectations over $\mathbf{u}$ and $\mathbf{v}$, we have

$$\mathbb{E}(\hat{\gamma}_j) = 0, \quad \mathbb{E}(\hat{\Gamma}_j) = 0. \tag{S.5}$$

Furthermore, we can express the GWAS estimates as

$$\begin{aligned}\hat{\gamma}_j &= \tilde{\gamma}_j + \epsilon_j,\\ \hat{\Gamma}_j &= \Gamma_j + \xi_j,\end{aligned}$$

where $\tilde{\gamma}_j = \sum_k r_{j,k}u_k$ and $\Gamma_j = \sum_k r_{j,k}v_k$ represent the true marginal effects of SNP $j$ on $X$ and $Y$, $\epsilon_j$ and $\xi_j$ are the estimation errors due to the sampling variation and confounding biases from sample structure.

###### 1.1.3 Derivation of the variance component of the background model

Using the results of single trait LDSC [7], the expected values of $z^2_{X,j}$ and $z^2_{Y,j}$ can be written as

$$\mathbb{E}(z^2_{X,j}) = \frac{N_1}{M}h_1^2\ell_j + \underbrace{1 + N_1F_{ST}(h_1^2F_{ST} + \sigma_s^2)}_{c_1},$$

$$\mathbb{E}(z^2_{Y,j}) = \frac{N_2}{M}h_2^2\ell_j + \underbrace{1 + N_2F_{ST}(h_2^2F_{ST} + \sigma_s^2)}_{c_2}.$$

Using the bivariate LD score regression, the expected value of $z_{X,j}z_{Y,j}$ can be written as [39],

$$\mathbb{E}(z_{X,j}z_{Y,j}) = \frac{\sqrt{N_1N_2}}{M}\rho_g\ell_j + \underbrace{\frac{N_s(\rho_g + \rho_e)}{\sqrt{N_1N_2}} + \rho_gF_{ST}^2\sqrt{N_1N_2} + \sqrt{N_1N_2}F_{ST}\sigma_s^2}_{c_{12}},$$

where $h_1^2$ and $h_2^2$ are heritabilities of $X$ and $Y$, $\rho_g$ is the genetic covariance between $X$ and $Y$, $c_1 \geq 1$ and $c_2 \geq 1$ in the presence of population stratification ($F_{ST} \neq 0$), and $c_{12} \neq 0$ in the presence of either population stratification (i.e., $F_{ST} \neq 0$) or sample overlap (i.e., $N_s \neq 0$). With the above results, we can obtain

$$\mathrm{Var}\begin{pmatrix}\hat{\gamma}_j\\ \hat{\Gamma}_j\end{pmatrix} = \frac{1}{M}\begin{pmatrix}h_1^2 & \rho_g\\ \rho_g & h_2^2\end{pmatrix}\ell_j + \begin{pmatrix}c_1\hat{s}^2_{X,j} & c_{12}\hat{s}_{X,j}\hat{s}_{Y,j}\\ c_{12}\hat{s}_{X,j}\hat{s}_{Y,j} & c_2\hat{s}^2_{Y,j}\end{pmatrix} = \ell_j\mathbf{\Omega} + \hat{\mathbf{S}}_j\mathbf{C}\hat{\mathbf{S}}_j, \tag{S.6}$$

where $\mathbf{\Omega} = \begin{pmatrix}\sigma_u^2 & r_g\sigma_u\tau_v\\ r_g\sigma_u\tau_v & \tau_v^2\end{pmatrix} = \frac{1}{M}\begin{pmatrix}h_1^2 & \rho_g\\ \rho_g & h_2^2\end{pmatrix}$ and $\hat{\mathbf{S}}_j = \begin{pmatrix}\hat{s}_{X,j} & 0\\ 0 & \hat{s}_{Y,j}\end{pmatrix}$. From Eq. [S.6], the polygenic effects and their correlation are tagged by the slope of the LD score $\ell_j$ and the influence of sample structure is captured by the intercept term. Considering the large sample size of modern GWASs, the estimation errors can be assumed asymptotically normally distributed. Combining Eqs. [S.5] and [S.6], we can obtain the background model given in Eq. [S.1].

We note that our model can incorporate covariates. To see this, we can extend model (S.2) to incorporate covariates:

$$\mathbf{x} = \mathbf{W}_1\mathbf{b}_{\text{cov,x}} + \mathbf{G}_1\mathbf{u} + \mathbf{s}_1 + \mathbf{e}_1, \qquad \mathbf{y} = \mathbf{W}_2\mathbf{b}_{\text{cov,y}} + \mathbf{G}_2\mathbf{v} + \mathbf{s}_2 + \mathbf{e}_2, \tag{S.7}$$

where $\mathbf{W}_1$ and $\mathbf{W}_2$ are the two matrices of covariates, $\mathbf{b}_{\text{cov,x}}$ and $\mathbf{b}_{\text{cov,y}}$ are the vectors of covariate effects. Now we define projection matrices $\mathbf{P}_1 = \mathbf{I} - \mathbf{W}_1(\mathbf{W}_1^T\mathbf{W}_1)^{-1}\mathbf{W}_1^T$ and $\mathbf{P}_2 = \mathbf{I} - \mathbf{W}_2(\mathbf{W}_2^T\mathbf{W}_2)^{-1}\mathbf{W}_2^T$. We can transform model (S.2) as following:

$$\mathbf{P}_1\mathbf{x} = \mathbf{P}_1\mathbf{G}_1\mathbf{u} + \mathbf{P}_1\mathbf{s}_1 + \mathbf{P}_1\mathbf{e}_1, \qquad \mathbf{P}_2\mathbf{y} = \mathbf{P}_2\mathbf{G}_2\mathbf{v} + \mathbf{P}_2\mathbf{s}_2 + \mathbf{P}_2\mathbf{e}_2. \tag{S.8}$$

By working with projected genotypes and phenotypes, model (S.7) is reduced to the same form of model (S.2) without covariates.

In summary, the background model of MR-APSS inherits the assumptions of LDSC to account for the confounding bias due to pleiotropy and sample structure. First, SNP effect sizes are assumed to be random effects, which allows the variance and covariance of SNP effects to be captured by the slope of LDSC, i.e., the coefficients of LD score $\ell_j$. Second, the rows of individual-level genotype matrices are assumed to be drawn i.i.d. from some distributions. This helps us to bypass the difficulty when individual-level GWAS data are inaccessible. Third, LDSC assumes the confounding bias from population stratification and overlapped samples is nearly constant across SNPs, such that their influence can be well captured by the intercept terms of LDSC. The first assumption and the third assumption allow us to distinguish genetic effects (polygenicity and correlated pleiotropy) from confounding bias due to sample structure. With these assumptions, we can estimate the parameters in the background model using genome-wide summary statistics. We have closely investigated the summary-statistics-based methods for estimating heritability and genetic correlation [40], including LDSC [7], GNOVA [23], and HDL [27]. Both simulation studies and real data analysis results suggest that the LDSC assumptions can provide a robust estimation of genetic correlation based on summary-level data as long as the reference genomes offer a matched LD estimation. In this paper, we mainly focus on causal inference in European ancestry. The reference genomes (e.g., from 1000 Genomes Project) are known to provide accurate LD estimation for European ancestry. Thus, MR-APSS can provide robust results even in the presence of model mis-specification.

#### 1.2 Derivation of the foreground-background model of MR-APSS

In this section, we derive the foreground-background model of MR-APSS given by Eq. [6] in the main text. We begin with the individual-level foreground-background model,

$$\mathbf{x} = \mathbf{G}_1(\mathbf{Z}\boldsymbol{\gamma} + \mathbf{u}) + \mathbf{s}_1 + \mathbf{e}_1, \quad \mathbf{y} = \mathbf{G}_2\left(\mathbf{Z}(\beta\boldsymbol{\gamma} + \boldsymbol{\alpha}) + \mathbf{v}\right) + \mathbf{s}_2 + \mathbf{e}_2, \tag{S.9}$$

where $\mathbf{x} = \{x_i\}_{i=1,\ldots,N_1}$ is an $N_1 \times 1$ phenotype vector for exposure $X$, $\mathbf{y} = \{y_i\}_{i=1,\ldots,N_2}$ is an $N_2 \times 1$ phenotype vector for outcome $Y$, $\mathbf{Z}$ is an $M \times M$ diagonal matrix where the $j$-th diagonal entry $Z_j \sim \text{Bern}(\pi_0)$ indicates that the SNP $j$ has a foreground signal ($Z_j = 1$) or not ($Z_j = 0$) with $\pi_0 = p(Z_j = 1)$, $\boldsymbol{\gamma} = \{\gamma_j\}_{j=1,\ldots,M}$ and $\boldsymbol{\alpha} = \{\alpha_j\}_{j=1,\ldots,M}$ are vectors collecting the instrument strengths and direct effects of the $M$ SNPs. We adopt the same assumptions as the background model above for $\mathbf{G}_1, \mathbf{G}_2, \mathbf{e}_1, \mathbf{e}_2, \mathbf{u}$, and $\mathbf{v}$. Additionally, we assume that $\gamma_j$ and $\alpha_j$ are normally distributed and independent of each other,

$$\begin{pmatrix} \gamma_j \\ \alpha_j \end{pmatrix} \sim \mathcal{N}\left(\mathbf{0}, \boldsymbol{\Sigma}\right), \text{ where } \boldsymbol{\Sigma} = \begin{pmatrix} \sigma^2 & 0 \\ 0 & \tau^2 \end{pmatrix}.$$

To assure the phenotype variances equal one, we require that $\sigma^2_{e_1}$ and $\sigma^2_{e_2}$ satisfy $M\pi_0\sigma^2 + M\sigma^2_u + \sigma^2_s + \sigma^2_{e_1} = 1$, and $M\pi_0(\tau^2 + \beta^2\sigma^2) + M\tau^2_v + \sigma^2_s + \sigma^2_{e_2} = 1$.

In MR-APSS, we adopt normality assumptions on the background effects $(\mathbf{u}, \mathbf{v})$ and the foreground effects $(\boldsymbol{\gamma}, \boldsymbol{\alpha})$. Specifically, $(\mathbf{u}, \mathbf{v})$ are random effects that capture the polygenicity of complex traits. Although many genome-wide significant variants have been identified in the early stage of GWASs, these variants can only explain a small fraction of phenotypic variance of complex traits, such as height, BMI, and T2D. This phenomenon was referred to as "missing heritability" [25]. Yang et al. [36] proposed a linear-mixed-model-based approach, where the random effects were assumed to be normal and the heritability was estimated using the restricted maximum likelihood (REML) approach. This seminal paper shows that the majority of heritability is not missing but jointly contributed by many variants with small effects known as polygenic effects. Nowadays, the polygenicity of complex traits is well accepted by the scientific community [32]. Regarding the distributions of $(\boldsymbol{\gamma}, \boldsymbol{\alpha})$, we made a normal assumption when building our MR-APSS model. Accumulating evidence from analyzing large-scale genetic data has implied that the normal distribution for $(\boldsymbol{\gamma}, \boldsymbol{\alpha})$ is a simple but very effective assumption for characterizing the effects deviating from the polygenic effects. An example includes Regression with Summary Statistics (RSS) [45, 46], where the large effects deviating from the polygenic effects are also well characterized by a normal distribution. Their comprehensive real data results also suggest that the normal distribution of SNP effect sizes is an effective assumption. In the MR literature, the normal assumption on the effect sizes is also commonly adopted. Examples include MRMix [30] and RAPS [41].

Let $\tilde{\gamma}_j$ and $\Gamma_j$ are the true marginal effects of SNP $j$ on $X$ and $Y$, respectively. Based on model in Eq. [S.9], we can obtain the estimated marginal effects of SNP $j$ and their standard errors by,

$$\begin{aligned} \hat{\gamma}_j &= \frac{\mathbf{G}_{1,j}^T \mathbf{x}}{N_1}, \quad \hat{s}_{X,j} \approx \frac{1}{\sqrt{N_1}}, \\ \hat{\Gamma}_j &= \frac{\mathbf{G}_{2,j}^T \mathbf{y}}{N_2}, \quad \hat{s}_{Y,j} \approx \frac{1}{\sqrt{N_2}}. \end{aligned} \tag{S.10}$$

Note that we only use the summary statistics for a subset of $M_t$ independent IVs for causal inference in our analysis, i.e., $\{\hat{\gamma}_j, \hat{\Gamma}_j, \hat{s}_{X,j}, \hat{s}_{Y,j} \| \hat{\gamma}_j / \hat{s}_{X,j}| > t\}_{j=1,\dots,M_t}$, which are obtained by PLINK clumping ($r^2 < 0.001$, 1Mb). After LD clumping, the IVs become representatives for the corresponding LD regions. To simplify the derivation, we assume that the $j$-th IV and the SNPs in its local LD region share the same indicator $Z_j$. We then have the following approximation:

$$\begin{aligned} \mathbb{E}(\hat{\gamma}_j | \mathbf{u}, \boldsymbol{\gamma}, Z_j) &\approx \sum_k r_{jk}(Z_j \gamma_k + u_k), \\ \mathbb{E}(\hat{\Gamma}_j | \mathbf{v}, \boldsymbol{\alpha}, Z_j) &\approx \sum_k r_{jk}(Z_j(\beta \gamma_k + \alpha_k) + v_k). \end{aligned}$$

Under the foreground-background model, we express $(\hat{\gamma}_j, \hat{\Gamma}_j)$ as

$$\begin{aligned} \hat{\gamma}_j &= \tilde{\gamma}_j + \epsilon_j, \\ \hat{\Gamma}_j &= \Gamma_j + \xi_j, \end{aligned} \tag{S.11}$$

where $\tilde{\gamma}_j = \sum_k r_{jk}(Z_j \gamma_k + u_k)$ and $\Gamma_j = \sum_k r_{jk}(Z_j(\beta \gamma_k + \alpha_k) + v_k)$ are the underlying true marginal effects of SNP $j$ on $X$ and $Y$, $\epsilon_j$ and $\xi_j$ are the estimation errors which capture the effects of sampling variation and confounding biases due to sample structure. As the GWAS sample size is large enough, we assume that $\epsilon_j$ and $\xi_j$ follow a normal distribution,

$$p(\hat{\gamma}_j, \hat{\Gamma}_j | Z_j, \mathbf{u}, \boldsymbol{\gamma}, \mathbf{v}, \boldsymbol{\alpha}, \mathbf{C}, \hat{\mathbf{S}}_j, \ell_j) = \mathcal{N}\left(\begin{pmatrix}\hat{\gamma}_j \\ \hat{\Gamma}_j\end{pmatrix} \Big| \begin{pmatrix}\sum_k r_{jk}(Z_j \gamma_k + u_k) \\ \sum_k r_{jk}(Z_j(\beta\gamma_k + \alpha_k) + v_k)\end{pmatrix}, \hat{\mathbf{S}}_j \mathbf{C} \hat{\mathbf{S}}_j\right). \tag{S.12}$$

By integrating out $u_k, v_k, \gamma_k, \alpha_k$, and $Z_j$ in Eq. [S.12], we obtain the foreground-background model of MR-APSS,

$$\begin{aligned}
&p(\hat{\gamma}_j, \hat{\Gamma}_j | \pi_0, \beta, \boldsymbol{\Sigma}, \boldsymbol{\Omega}, \mathbf{C}, \hat{\mathbf{S}}_j, \ell_j) \\
=&\pi_0 \mathcal{N}\left(\begin{pmatrix}\hat{\gamma}_j \\ \hat{\Gamma}_j\end{pmatrix} \Big| \mathbf{0}, \ell_j \mathbf{A}(\beta) \boldsymbol{\Sigma} \mathbf{A}(\beta)^T + \ell_j \boldsymbol{\Omega} + \hat{\mathbf{S}}_j \mathbf{C} \hat{\mathbf{S}}_j\right) + (1-\pi_0)\mathcal{N}\left(\begin{pmatrix}\hat{\gamma}_j \\ \hat{\Gamma}_j\end{pmatrix} \Big| \mathbf{0}, \ell_j \boldsymbol{\Omega} + \hat{\mathbf{S}}_j \mathbf{C} \hat{\mathbf{S}}_j\right),
\end{aligned}$$

where $\mathbf{A}(\beta) = \begin{pmatrix}1 & 0 \\ \beta & 1\end{pmatrix}$.

#### 1.3 Accounting for selection bias in MR-APSS

We derive the probabilistic model given in Eq. [7] in the main text. To account for bias due to the IV selection, we modify model (6) in the main text by conditioning on the selection operation $|\hat{\gamma}_j/\hat{s}_{X,j}| > t$, where $t$ is a $z$-score threshold corresponding to a $p$-value threshold for the IV selection. Thus, we have

$$\begin{aligned}
&p\left(\hat{\gamma}_j, \hat{\Gamma}_j \Big| |\hat{\gamma}_j/\hat{s}_{X,j}| > t\right) \\
=&p\left(Z_j = 1 \Big| |\hat{\gamma}_j/\hat{s}_{X,j}| > t\right) p\left(\hat{\gamma}_j, \hat{\Gamma}_j \Big| Z_j = 1, |\hat{\gamma}_j/\hat{s}_{X,j}| > t\right) + \\
&p\left(Z_j = 0 \Big| |\hat{\gamma}_j/\hat{s}_{X,j}| > t\right) p\left(\hat{\gamma}_j, \hat{\Gamma}_j \Big| Z_j = 0, |\hat{\gamma}_j/\hat{s}_{X,j}| > t\right) \\
=&\pi_t \frac{p\left(\hat{\gamma}_j, \hat{\Gamma}_j | Z_j = 1\right)}{p\left(|\hat{\gamma}_j/\hat{s}_{X,j}| > t \Big| Z_j = 1\right)} + (1-\pi_t)\frac{p\left(\hat{\gamma}_j, \hat{\Gamma}_j \Big| Z_j = 0\right)}{p\left(|\hat{\gamma}_j/\hat{s}_{X,j}| > t \Big| Z_j = 0\right)} \\
=&\pi_t \frac{\mathcal{N}\left(\begin{pmatrix}\hat{\gamma}_j \\ \hat{\Gamma}_j\end{pmatrix} \Big| \mathbf{0}, \ell_j \mathbf{A}(\beta) \boldsymbol{\Sigma} \mathbf{A}(\beta)^T + \ell_j \boldsymbol{\Omega} + \hat{\mathbf{S}}_j \mathbf{C} \hat{\mathbf{S}}_j\right)}{2\Phi\left(\frac{-t\hat{s}_{X,j}}{\sqrt{\ell_j \sigma^2 + \ell_j \sigma_u^2 + c_1 \hat{s}_{X,j}^2}}\right)} + (1-\pi_t)\frac{\mathcal{N}\left(\begin{pmatrix}\hat{\gamma}_j \\ \hat{\Gamma}_j\end{pmatrix} \Big| \mathbf{0}, \ell_j \boldsymbol{\Omega} + \hat{\mathbf{S}}_j \mathbf{C} \hat{\mathbf{S}}_j\right)}{2\Phi\left(\frac{-t\hat{s}_{X,j}}{\sqrt{\ell_j \sigma_u^2 + c_1 \hat{s}_{X,j}^2}}\right)},
\end{aligned}$$

where $\pi_t = p\left(Z_j = 1 \Big| |\hat{\gamma}_j/\hat{s}_{X,j}| > t\right)$ is the probability that the $j$-th IV carries the foreground signal after selection and $\Phi(\cdot)$ is the standard normal cumulative distribution function. From the third line to the fourth line, we used the foreground component and background component

$$\begin{aligned}
p(\hat{\gamma}_j, \hat{\Gamma}_j | Z_j = 1) &= \mathcal{N}\left(\begin{pmatrix}\hat{\gamma}_j \\ \hat{\Gamma}_j\end{pmatrix} \Big| \mathbf{0}, \ell_j \mathbf{A}(\beta) \boldsymbol{\Sigma} \mathbf{A}(\beta)^T + \ell_j \boldsymbol{\Omega} + \hat{\mathbf{S}}_j \mathbf{C} \hat{\mathbf{S}}_j\right), \\
p(\hat{\gamma}_j, \hat{\Gamma}_j | Z_j = 0) &= \mathcal{N}\left(\begin{pmatrix}\hat{\gamma}_j \\ \hat{\Gamma}_j\end{pmatrix} \Big| \mathbf{0}, \ell_j \boldsymbol{\Omega} + \hat{\mathbf{S}}_j \mathbf{C} \hat{\mathbf{S}}_j\right).
\end{aligned}$$

#### 1.4 Parameter Estimation

##### 1.4.1 Estimation of $\mathbf{\Omega}$ and $\mathbf{C}$ in the background model

We use LDSC to estimate the matrices $\mathbf{\Omega}$ and $\mathbf{C}$ in the background model of MR-APSS, where genome-wide summary statistics are taken as inputs. Based on Eq. [S.6], we then construct the estimates of $\mathbf{\Omega}$ and $\mathbf{C}$ by:

$$\hat{\mathbf{\Omega}} = \frac{1}{M}\begin{pmatrix} \hat{h}_1^2 & \hat{\rho}_g \\ \hat{\rho}_g & \hat{h}_2^2 \end{pmatrix},$$

and

$$\hat{\mathbf{C}} = \begin{pmatrix} \hat{c}_1 & \hat{c}_{12} \\ \hat{c}_{12} & \hat{c}_2 \end{pmatrix},$$

where $\hat{h}_1^2$ and $\hat{h}_2^2$ are the heritability estimates from the slopes of single-trait LD score regressions for $X$ and $Y$, $\hat{c}_1$ and $\hat{c}_2$ are the intercepts estimated from single-trait LD score regression for $X$ and $Y$, $\hat{\rho}_g$ is the estimate of genetic covariance, and $\hat{c}_{12}$ is the intercept estimate from bivariate LD score regression, respectively.

Regarding the estimation of $\mathbf{\Omega}$ and $\mathbf{C}$, there are two important questions to be addressed. First, MR-APSS is built upon the proposed foreground-background model. Since $\hat{\mathbf{\Omega}}$ is estimated based on summary statistics across the whole genome, the foreground signals are also included to estimate $\mathbf{\Omega}$. Does $\hat{\mathbf{\Omega}}$ over-estimate the magnitude of the invalid signals of the background component and thus lead to reduced statistical power? Second, MR-APSS assumes that the estimation uncertainty of $\hat{\mathbf{\Omega}}$ and $\hat{\mathbf{C}}$ can be ignored. To what extent does the estimation uncertainty of $\hat{\mathbf{\Omega}}$ and $\hat{\mathbf{C}}$ affect causal inference? Next, we provide more evidence to address these two questions.

**(1). Evaluation of the influence of overestimation of $\mathbf{\Omega}$ on the power of MR-APSS**

We believe that the over-estimation should be minor due to the polygenic nature of human genetics: the genome-wide significant SNPs often can only explain a very small proportion of heritability, and thus the inclusion of those SNPs can only contribute a tiny amount of overestimation. As we have shown before, the $p$-values of MR-APSS were well-calibrated (nearly uniformly distributed between 0 and 1) in both null simulations and real data analysis with negative control outcomes, suggesting the small amount of over-estimation is ignorable under null. Then a remaining concern is whether the over-estimation would reduce the power of MR-APSS under alternatives. To illustrate this, we manually fixed the background component $\hat{\mathbf{\Omega}}$ and $\hat{\mathbf{C}}$ at their ground truth (denoted as MR-APSS (fix background at its truth)) and compared it with MR-APSS. The comparison of the two methods in terms of the power is shown in Fig. S12, suggesting that the overestimation of $\mathbf{\Omega}$ lead to a minor decrease in power. As shown in the comprehensive simulations and real-data analysis, MR-APSS can still provide high statistical power.

**(2). Evaluation of the influence of the estimation uncertainty in $\hat{\mathbf{\Omega}}$ and $\hat{\mathbf{C}}$ on $p$-values from MR-APSS**

We have shown that MR-APSS produced calibrated $p$-values in the null simulations under various settings. Here we conducted the simulation under alternative to evaluate the influence of the estimation uncertainty in $\hat{\mathbf{\Omega}}$ and $\hat{\mathbf{C}}$ on $p$-values from MR-APSS. We compared the $p$-values from MR-APSS with/without accounting for uncertainty in $\hat{\mathbf{\Omega}}$ and $\hat{\mathbf{C}}$, denoted as MR-APSS (account for uncertainty in $\hat{\mathbf{\Omega}}$ and $\hat{\mathbf{C}}$) and MR-APSS, respectively.

We used a block-wise jackknife approach to measure the uncertainty in $\hat{\beta}$ due to estimation error of $\hat{\mathbf{\Omega}}$ and $\hat{\mathbf{C}}$ using genome-wide summary statistics. Specifically, we divided the genome-wide SNPs into $n = 22$ blocks and then applied a delete-one-block procedure to estimate $\mathbf{\Omega}$ and $\mathbf{C}$. As such, we obtained $n = 22$ pairs of estimated $\hat{\mathbf{\Omega}}$ and $\hat{\mathbf{C}}$. After that, we applied MR-APSS using these estimates and obtained $n = 22$ estimated $\hat{\beta}$ which were regarded as the delete-one-block estimates of $\beta$. Based on these estimates, we then calculated the jackknife Standard Error (SE) which accounts for the uncertainty in $\hat{\beta}$ due to estimation of $\mathbf{\Omega}$ and $\mathbf{C}$. We denoted this standard error as $\mathrm{SE}_0(\hat{\beta})$. As a conservative estimation of the standard error, we defined the total standard error of $\hat{\beta}$ accounting for the uncertainty in $\hat{\mathbf{\Omega}}$, $\hat{\mathbf{C}}$, and the model fitting as

$$\mathrm{SE}_{Total}(\hat{\beta}) = \sqrt{\mathrm{SE}_0(\hat{\beta})^2 + \mathrm{SE}(\hat{\beta})^2},$$

where $\mathrm{SE}(\hat{\beta})$ is the standard error of $\beta$ from MR-APSS (without accounting for uncertainty in $\hat{\mathbf{\Omega}}$ and $\hat{\mathbf{C}}$).

From simulation results shown in Fig. S11, we found that the inference $p$-values obtained by MR-APSS (accounting for uncertainty in $\hat{\mathbf{\Omega}}$ and $\hat{\mathbf{C}}$) and MR-APSS agreed well with each other. Our results suggest that the influence of estimation uncertainty in $\hat{\mathbf{\Omega}}$ and $\hat{\mathbf{C}}$ on $p$-values obtained from MR-APSS was ignorable.

##### 1.4.2 The Variational EM algorithm

We derive a variational EM algorithm to obtain the estimates of the unknown parameters $\boldsymbol{\theta} = (\beta, \pi_t, \sigma^2, \tau^2)$ by maximizing the log-likelihood function given in Eq. [8] of the main text. We denote $\hat{\boldsymbol{\gamma}} = \{\hat{\gamma}_j\}_{j=1,\dots,M_t}$, $\hat{\mathbf{\Gamma}} = \{\hat{\Gamma}_j\}_{j=1,\dots,M_t}$, $\boldsymbol{\gamma} = \{\gamma_j\}_{j=1,\dots,M_t}$, $\boldsymbol{\alpha} = \{\alpha_j\}_{j=1,\dots,M_t}$, and $\mathbf{Z} = \{Z_j\}_{j=1,\dots,M_t}$. By treating $\boldsymbol{\gamma}, \boldsymbol{\alpha}$, and $\mathbf{Z}$ as latent variables, the complete data likelihood can be obtained as following:

$$\begin{aligned}
& p\left(\hat{\boldsymbol{\gamma}}, \hat{\mathbf{\Gamma}}, \boldsymbol{\gamma}, \boldsymbol{\alpha}, \mathbf{Z} \middle| \boldsymbol{\theta}, t, M_t\right) \\
=& \prod_{j=1}^{M_t} p\left(\hat{\gamma}_j, \hat{\Gamma}_j \middle| \gamma_j, \alpha_j, Z_j, \boldsymbol{\theta}, |\hat{\gamma}_j/\hat{s}_{X,j}| > t) \cdot p(\gamma_j, \alpha_j \middle| Z_j, \boldsymbol{\theta}, |\hat{\gamma}_j/\hat{s}_{X,j}| > t\right) \cdot p\left(Z_j \middle| \boldsymbol{\theta}, |\hat{\gamma}_j/\hat{s}_{X,j}| > t\right) \\
=& \prod_{j=1}^{M_t} \frac{p\left(\hat{\gamma}_j, \hat{\Gamma}_j \middle| \gamma_j, \alpha_j, Z_j, \boldsymbol{\theta}\right)}{p\left(|\hat{\gamma}_j/\hat{s}_{X,j}| > t \middle| \gamma_j, \alpha_j, Z_j, \boldsymbol{\theta}\right)} \cdot \frac{p\left(|\hat{\gamma}_j/\hat{s}_{X,j}| > t \middle| \gamma_j, \alpha_j, Z_j\right) p\left(\gamma_j, \alpha_j \middle| Z_j, \boldsymbol{\theta}\right)}{p\left(|\hat{\gamma}_j/\hat{s}_{X,j}| > t \middle| Z_j, \boldsymbol{\theta}\right)} \cdot p\left(Z_j \middle| |\hat{\gamma}_j/\hat{s}_{X,j}| > t, \boldsymbol{\theta}\right) \\
=& \prod_{j=1}^{M_t} \frac{p\left(\hat{\gamma}_j, \hat{\Gamma}_j \middle| \gamma_j, \alpha_j, Z_j, \boldsymbol{\theta}\right) p\left(\gamma_j, \alpha_j \middle| Z_j, \boldsymbol{\theta}\right)}{p\left(|\hat{\gamma}_j/\hat{s}_{X,j}| > t \middle| Z_j, \boldsymbol{\theta}\right)} \cdot p\left(Z_j \middle| |\hat{\gamma}_j/\hat{s}_{X,j}| > t, \boldsymbol{\theta}\right) \\
=& \prod_{j=1}^{M_t} \frac{\mathcal{N}\left(\begin{pmatrix} \hat{\gamma}_j \\ \hat{\Gamma}_j \end{pmatrix} | Z_j \mathbf{A}(\beta) \begin{pmatrix} \gamma_j \\ \alpha_j \end{pmatrix}, \ell_j \hat{\mathbf{\Omega}} + \hat{\mathbf{S}}_j \hat{\mathbf{C}} \hat{\mathbf{S}}_j\right) \mathcal{N}\left(\begin{pmatrix} \gamma_j \\ \alpha_j \end{pmatrix} | \mathbf{0}, \ell_j \mathbf{\Sigma}\right)}{\left(2\Phi\left(\frac{-t\hat{s}_{X,j}}{\sqrt{\ell_j \hat{\sigma}_u^2 + \hat{c}_1 \hat{s}_{X,j}^2}}\right)\right)^{1-Z_j} \left(2\Phi\left(\frac{-t\hat{s}_{X,j}}{\sqrt{\ell_j \sigma^2 + \ell_j \hat{\sigma}_u^2 + \hat{c}_1 \hat{s}_{X,j}^2}}\right)\right)^{Z_j}} \pi_t^{Z_j} \pi_t^{1-Z_j}.
\end{aligned} \tag{S.13}$$

Here we use $p(\gamma_j, \alpha_j|\ell_j, \mathbf{\Sigma}) = \mathcal{N}\left(\begin{pmatrix}\gamma_j\\ \alpha_j\end{pmatrix} \Big| \mathbf{0}, \ell_j\mathbf{\Sigma}\right)$ after accounting for LD.

Given Eq. [S.13], the complete data log-likelihood can be written as:

$$\begin{aligned}
&\log p(\hat{\boldsymbol{\gamma}}, \hat{\mathbf{\Gamma}}, \boldsymbol{\gamma}, \boldsymbol{\alpha}, \mathbf{Z}\Big|\boldsymbol{\theta}, t, M_t)\\
=&\sum_{j=1}^{M_t} \log \mathcal{N}\left(\begin{pmatrix}\hat{\gamma}_j\\ \hat{\Gamma}_j\end{pmatrix} \mid Z_j\mathbf{A}(\beta)\begin{pmatrix}\gamma_j\\ \alpha_j\end{pmatrix}, \ell_j\hat{\mathbf{\Omega}} + \hat{\mathbf{S}}_j\hat{\mathbf{C}}\hat{\mathbf{S}}_j\right) +\\
&\sum_{j=1}^{M_t} \log \mathcal{N}\left(\begin{pmatrix}\gamma_j\\ \alpha_j\end{pmatrix} \mid \mathbf{0}, \ell_j\mathbf{\Sigma}\right) + \sum_{j=1}^{M_t} Z_j \log \pi_t + (1-Z_j)\log(1-\pi_t) -\\
&\sum_{j=1}^{M_t} Z_j \log\left(2\Phi\left(\frac{-t\hat{s}_{X,j}}{\sqrt{\ell_j\sigma^2 + \ell_j\hat{\sigma}_u^2 + \hat{c}_1\hat{s}_{X,j}^2}}\right)\right) - (1-Z_j)\log\left(2\Phi\left(\frac{-t\hat{s}_{X,j}}{\sqrt{\ell_j\hat{\sigma}_u^2 + \hat{c}_1\hat{s}_{X,j}^2}}\right)\right)\\
=&\sum_{j=1}^{M_t} -\frac{1}{2}\log\det(\hat{\mathbf{S}}_j\hat{\mathbf{C}}\hat{\mathbf{S}}_j + \ell_j\hat{\mathbf{\Omega}}) -\\
&\sum_{j=1}^{M_t} \frac{1}{2}\left\{\begin{pmatrix}\hat{\gamma}_j\\ \hat{\Gamma}_j\end{pmatrix} - Z_j\mathbf{A}(\beta)\begin{pmatrix}\gamma_j\\ \alpha_j\end{pmatrix}\right\}^T (\hat{\mathbf{S}}_j\hat{\mathbf{C}}\hat{\mathbf{S}}_j + \ell_j\hat{\mathbf{\Omega}})^{-1}\left\{\begin{pmatrix}\hat{\gamma}_j\\ \hat{\Gamma}_j\end{pmatrix} - Z_j\mathbf{A}(\beta)\begin{pmatrix}\gamma_j\\ \alpha_j\end{pmatrix}\right\} +\\
&\sum_{j=1}^{M_t} -\frac{1}{2}\log\det(\ell_j\mathbf{\Sigma}) - \frac{1}{2}\begin{pmatrix}\gamma_j\\ \alpha_j\end{pmatrix}^T (\ell_j\mathbf{\Sigma})^{-1}\begin{pmatrix}\gamma_j\\ \alpha_j\end{pmatrix} + \sum_{j=1}^{M_t} Z_j \log \pi_t + (1-Z_j)\log(1-\pi_t) -\\
&\sum_{j=1}^{M_t} Z_j \log\left(2\Phi\left(\frac{-t\hat{s}_{X,j}}{\sqrt{\ell_j\sigma^2 + \ell_j\hat{\sigma}_u^2 + \hat{c}_1\hat{s}_{X,j}^2}}\right)\right) - (1-Z_j)\log\left(2\Phi\left(\frac{-t\hat{s}_{X,j}}{\sqrt{\ell_j\hat{\sigma}_u^2 + \hat{c}_1\hat{s}_{X,j}^2}}\right)\right) + \text{constant}.
\end{aligned}$$

Let $q(\boldsymbol{\gamma}, \boldsymbol{\alpha}, \mathbf{Z})$ be a variational distribution. The logarithm of the marginal likelihood can be written as

$$\begin{aligned}
&\log p(\hat{\boldsymbol{\gamma}}, \hat{\mathbf{\Gamma}}|\boldsymbol{\theta}, t, M_t)\\
=&\mathbb{E}_{q(\boldsymbol{\gamma},\boldsymbol{\alpha},\mathbf{Z})}(\log p(\hat{\boldsymbol{\gamma}}, \hat{\mathbf{\Gamma}}|\boldsymbol{\theta}, t, M_t))\\
=&\mathbb{E}_{q(\boldsymbol{\gamma},\boldsymbol{\alpha},\mathbf{Z})}\left(\log \frac{p(\hat{\boldsymbol{\gamma}}, \hat{\mathbf{\Gamma}}, \boldsymbol{\gamma}, \boldsymbol{\alpha}, \mathbf{Z}|\boldsymbol{\theta}, t, M_t)}{p(\boldsymbol{\gamma}, \boldsymbol{\alpha}, \mathbf{Z}\big|\hat{\boldsymbol{\gamma}}, \hat{\mathbf{\Gamma}}, \boldsymbol{\theta}, t, M_t)}\right)\\
=&\mathbb{E}_{q(\boldsymbol{\gamma},\boldsymbol{\alpha},\mathbf{Z})}\left(\log \frac{p(\hat{\boldsymbol{\gamma}}, \hat{\mathbf{\Gamma}}, \boldsymbol{\gamma}, \boldsymbol{\alpha}, \mathbf{Z}|\boldsymbol{\theta}, t, M_t)}{q(\boldsymbol{\gamma}, \boldsymbol{\alpha}, \mathbf{Z})} - \log \frac{p(\boldsymbol{\gamma}, \boldsymbol{\alpha}, \mathbf{Z}\big|\hat{\boldsymbol{\gamma}}, \hat{\mathbf{\Gamma}}, \boldsymbol{\theta}, t, M_t)}{q(\boldsymbol{\gamma}, \boldsymbol{\alpha}, \mathbf{Z})}\right)\\
=&\mathcal{L}(q; \boldsymbol{\theta}, t, M_t) + \mathrm{D}_{\mathrm{KL}}\left(q(\boldsymbol{\gamma}, \boldsymbol{\alpha}, \mathbf{Z})||p(\boldsymbol{\gamma}, \boldsymbol{\alpha}, \mathbf{Z}\big|\hat{\boldsymbol{\gamma}}, \hat{\mathbf{\Gamma}}, \boldsymbol{\theta}, t, M_t)\right),
\end{aligned}$$

where

$$\mathcal{L}(q; \boldsymbol{\theta}, t, M_t) = \mathbb{E}_{q(\boldsymbol{\gamma},\boldsymbol{\alpha},\mathbf{Z})}\left(\log \frac{p(\hat{\boldsymbol{\gamma}}, \hat{\mathbf{\Gamma}}, \boldsymbol{\gamma}, \boldsymbol{\alpha}, \mathbf{Z}|\boldsymbol{\theta}, t, M_t)}{q(\boldsymbol{\gamma}, \boldsymbol{\alpha}, \mathbf{Z})}\right),$$

$$\mathrm{D}_{\mathrm{KL}}\left(q(\boldsymbol{\gamma}, \boldsymbol{\alpha}, \mathbf{Z})||p(\boldsymbol{\gamma}, \boldsymbol{\alpha}, \mathbf{Z}\big|\hat{\boldsymbol{\gamma}}, \hat{\mathbf{\Gamma}}, \boldsymbol{\theta}, t, M_t)\right) = -\mathbb{E}_{q(\boldsymbol{\gamma},\boldsymbol{\alpha},\mathbf{Z})}\left(\log \frac{p(\boldsymbol{\gamma}, \boldsymbol{\alpha}, \mathbf{Z}\big|\hat{\boldsymbol{\gamma}}, \hat{\mathbf{\Gamma}}, \boldsymbol{\theta}, t, M_t)}{q(\boldsymbol{\gamma}, \boldsymbol{\alpha}, \mathbf{Z})}\right).$$

Since the Kullback-Leibler (KL) divergence $\mathrm{D}_{\mathrm{KL}}\left(q(\boldsymbol{\gamma}, \boldsymbol{\alpha}, \mathbf{Z})||p(\boldsymbol{\gamma}, \boldsymbol{\alpha}, \mathbf{Z}\big|\hat{\boldsymbol{\gamma}}, \hat{\mathbf{\Gamma}}, \boldsymbol{\theta}, t, M_t)\right)$ is non-negative, $\mathcal{L}(q; \boldsymbol{\theta}, t, M_t)$ is the evidence lower bound (ELBO) of the marginal log-likelihood $\log p(\hat{\boldsymbol{\gamma}}, \hat{\mathbf{\Gamma}}|\boldsymbol{\theta}, t, M_t)$. Thus, maximization of $\mathcal{L}(q; \boldsymbol{\theta}, t, M_t)$ w.r.t. variational distribution $q$ and parameter $\boldsymbol{\theta}$ follows the EM framework: in the E-step, variational distribution $q$ is updated to approximate the true posterior; in the M-step, parameters in $\boldsymbol{\theta}$ are optimized to increase the ELBO.

**E-step.** To make it feasible for evaluation of the lower bound $\mathcal{L}(q; \boldsymbol{\theta}, t, M_t)$, we adopt the mean-field assumption that the variational distribution $q(\boldsymbol{\gamma}, \boldsymbol{\alpha}, \mathbf{Z})$ can be factorized as:

$$q(\boldsymbol{\gamma}, \boldsymbol{\alpha}, \mathbf{Z}) = \prod_{j=1}^{M_t} q(\gamma_j, \alpha_j, Z_j) = \prod_{j=1}^{M_t} q(\gamma_j, \alpha_j | Z_j) q(Z_j). \tag{S.14}$$

Noting that $Z_j$ is a binary variable, we define

$$q(Z_j) = \omega_j^{Z_j} (1 - \omega_j)^{(1-Z_j)}, \quad \text{where } \omega_j = q(Z_j = 1). \tag{S.15}$$

Based on the mean-field approximation, we can derive the optimal solutions for the $q$ distribution in Eq. [S.14] at each step. We first obtain the optimal solution for $q(\gamma_j, \alpha_j | Z_j)$, for $j = 1, \dots, M_t$. Given $Z_j = 1$, we have

$$\log q(\gamma_j, \alpha_j | Z_j = 1) = \mathbb{E}_{q_{-j}} \left( \log p(\hat{\boldsymbol{\gamma}}, \hat{\boldsymbol{\Gamma}}, \boldsymbol{\gamma}, \boldsymbol{\alpha}, \mathbf{Z} | \boldsymbol{\theta}, t, M_t) \right) + \text{constant},$$

where $\mathbb{E}_{q_{-j}}$ denotes the expectation w.r.t. the $q$ distribution over $(\boldsymbol{\gamma},\ \boldsymbol{\alpha})$ except $(\gamma_j, \alpha_j)$, conditioning on $Z_j = 1$. Thus, we have

$$\begin{aligned} &\log q(\gamma_j, \alpha_j | Z_j = 1) \\ &= -\frac{1}{2} \left\{ \begin{pmatrix} \hat{\gamma}_j \\ \hat{\Gamma}_j \end{pmatrix} - \mathbf{A}(\beta) \begin{pmatrix} \gamma_j \\ \alpha_j \end{pmatrix} \right\}^T (\hat{\mathbf{S}}_j \hat{\mathbf{C}} \hat{\mathbf{S}}_j + \ell_j \hat{\boldsymbol{\Omega}})^{-1} \left\{ \begin{pmatrix} \hat{\gamma}_j \\ \hat{\Gamma}_j \end{pmatrix} - \mathbf{A}(\beta) \begin{pmatrix} \gamma_j \\ \alpha_j \end{pmatrix} \right\} \\ &\quad - \frac{1}{2} \begin{pmatrix} \gamma_j \\ \alpha_j \end{pmatrix}^T \ell_j^{-1} \boldsymbol{\Sigma}^{-1} \begin{pmatrix} \gamma_j \\ \alpha_j \end{pmatrix} + \text{constant}. \end{aligned}$$

We observe that the right hand side of the above expression is a quadratic function of $\gamma_j$ and $\alpha_j$, and we can identify $q(\gamma_j, \alpha_j | Z_j = 1)$ as a Gaussian distribution:

$$q(\gamma_j, \alpha_j | Z_j = 1) = \mathcal{N} \left( \begin{pmatrix} \gamma_j \\ \alpha_j \end{pmatrix} \Big| \boldsymbol{\mu}_j, \boldsymbol{\Lambda}_j^{-1} \right), \tag{S.16}$$

where

$$\begin{aligned} \boldsymbol{\Lambda}_j &= \mathbf{A}(\beta)^T (\hat{\mathbf{S}}_j \hat{\mathbf{C}} \hat{\mathbf{S}}_j + \ell_j \hat{\boldsymbol{\Omega}})^{-1} \mathbf{A}(\beta) + \ell_j^{-1} \boldsymbol{\Sigma}^{-1}, \\ \boldsymbol{\mu}_j &= \boldsymbol{\Lambda}_j^{-1} \mathbf{A}(\beta)^T (\hat{\mathbf{S}}_j \hat{\mathbf{C}} \hat{\mathbf{S}}_j + \ell_j \hat{\boldsymbol{\Omega}})^{-1} \begin{pmatrix} \hat{\gamma}_j \\ \hat{\Gamma}_j \end{pmatrix}. \end{aligned}$$

Similarly, the optimal solution for $q(\gamma_j, \alpha_j | Z_j = 0)$ is given by

$$\log q(\gamma_j, \alpha_j | Z_j = 0) = -\frac{1}{2} \begin{pmatrix} \gamma_j \\ \alpha_j \end{pmatrix}^T \ell_j^{-1} \boldsymbol{\Sigma}^{-1} \begin{pmatrix} \gamma_j \\ \alpha_j \end{pmatrix} + \text{constant}.$$

Thus, we have

$$q(\gamma_j, \alpha_j | Z_j = 0) = \mathcal{N} \left( \begin{pmatrix} \gamma_j \\ \alpha_j \end{pmatrix} \Big| \mathbf{0}, \ell_j \boldsymbol{\Sigma} \right). \tag{S.17}$$

Combining Eqs. [S.14], [S.15], [S.16], and [S.17], we have

$$q(\gamma_j, \alpha_j, Z_j) = \left[\omega_j \mathcal{N}\left(\begin{pmatrix}\gamma_j \\ \alpha_j\end{pmatrix} \middle| \boldsymbol{\mu}_j, \boldsymbol{\Lambda}_j^{-1}\right)\right]^{Z_j} \left[(1-\omega_j)\mathcal{N}\left(\begin{pmatrix}\gamma_j \\ \alpha_j\end{pmatrix} \middle| \mathbf{0}, \ell_j \boldsymbol{\Sigma}\right)\right]^{1-Z_j}.$$

Once the variational distribution $q(\gamma_j, \alpha_j, Z_j)$ is obtained, we can evaluate the ELBO $\mathcal{L}(q; \boldsymbol{\theta}, t, M_t)$:

$$\mathcal{L}(q; \boldsymbol{\theta}, t, M_t) = \mathbb{E}_q \log p(\hat{\boldsymbol{\gamma}}, \hat{\boldsymbol{\Gamma}}, \boldsymbol{\gamma}, \boldsymbol{\alpha}, \mathbf{Z} | \boldsymbol{\theta}, t, M_t) - \mathbb{E}_q \log q(\boldsymbol{\gamma}, \boldsymbol{\alpha}, \mathbf{Z}), \tag{S.18}$$

where

$$\begin{aligned}
&\mathbb{E}_q \log p(\hat{\boldsymbol{\gamma}}, \hat{\boldsymbol{\Gamma}}, \boldsymbol{\gamma}, \boldsymbol{\alpha}, \mathbf{Z} \Big| \boldsymbol{\theta}, t, M_t) \\
=&\sum_{j=1}^{M_t} \omega_j \boldsymbol{\mu}_j^T \mathbf{A}(\beta)^T (\hat{\mathbf{S}}_j \hat{\mathbf{C}} \hat{\mathbf{S}}_j + \ell_j \hat{\boldsymbol{\Omega}})^{-1} \begin{pmatrix}\hat{\gamma}_j \\ \hat{\Gamma}_j\end{pmatrix} - \\
&\sum_{j=1}^{M_t} \frac{1}{2} \omega_j \operatorname{Tr}\left[\mathbf{A}(\beta)^T (\hat{\mathbf{S}}_j \hat{\mathbf{C}} \hat{\mathbf{S}}_j + \ell_j \hat{\boldsymbol{\Omega}})^{-1} \mathbf{A}(\beta) (\boldsymbol{\Lambda}_j^{-1} + \boldsymbol{\mu}_j \boldsymbol{\mu}_j^T)\right] + \\
&\sum_{j=1}^{M_t} -\frac{1}{2} \log\det(\ell_j \boldsymbol{\Sigma}) - \frac{1}{2} \omega_j \ell_j^{-1} \boldsymbol{\mu}_j^T \boldsymbol{\Sigma}^{-1} \boldsymbol{\mu}_j - \frac{1}{2} \operatorname{Tr}\left[\omega_j \ell_j^{-1} \boldsymbol{\Sigma}^{-1} \boldsymbol{\Lambda}_j^{-1}\right] - (1-\omega_j) + \\
&\sum_{j=1}^{M_t} \omega_j \log \pi_t + (1-\omega_j) \log(1-\pi_t) - \\
&\sum_{j=1}^{M_t} \omega_j \log\left(2\Phi\left(\frac{-t\hat{s}_{X,j}}{\sqrt{\ell_j \sigma^2 + \ell_j \hat{\sigma}_u^2 + \hat{c}_1 \hat{s}_{X,j}^2}}\right)\right) - (1-\omega_j) \log\left(2\Phi\left(\frac{-t\hat{s}_{X,j}}{\sqrt{\ell_j \hat{\sigma}_u^2 + \hat{c}_1 \hat{s}_{X,j}^2}}\right)\right) + \text{constant},
\end{aligned}$$

and

$$\begin{aligned}
&- \mathbb{E}_q \log q(\boldsymbol{\gamma}, \boldsymbol{\alpha}, \mathbf{Z}) \\
=&\sum_{j=1}^{M_t} \frac{1}{2} \omega_j \log\det(\boldsymbol{\Lambda}_j^{-1}) + \frac{1}{2}(1-\omega_j) \log\det(\ell_j \boldsymbol{\Sigma}) - \omega_j \log \omega_j - (1-\omega_j) \log(1-\omega_j).
\end{aligned}$$

By maximizing $\mathcal{L}(q; \boldsymbol{\theta}, t, M_t)$ w.r.t. $\omega_j$, we obtain

$$\omega_j = \frac{1}{1 + \exp(-\mathbf{b}_j)},$$

where

$$\mathbf{b}_j = \frac{1}{2} \boldsymbol{\mu}_j^T \boldsymbol{\Lambda}_j \boldsymbol{\mu}_j + \log \frac{\pi_t}{1-\pi_t} + \frac{1}{2} \log \frac{\det(\boldsymbol{\Lambda}_j^{-1})}{\det(\ell_j \boldsymbol{\Sigma})} - \log \frac{\Phi\left(\frac{-t\hat{s}_{X,j}}{\sqrt{\ell_j \sigma^2 + \ell_j \hat{\sigma}_u^2 + \hat{c}_1 \hat{s}_{X,j}^2}}\right)}{\Phi\left(\frac{-t\hat{s}_{X,j}}{\sqrt{\ell_j \hat{\sigma}_u^2 + \hat{c}_1 \hat{s}_{X,j}^2}}\right)}.$$

**M-step.** We derive the updating equations for parameters $\beta$, $\pi_t$, $\tau^2$, and $\sigma^2$. We first derive the updating equation for $\beta$. The terms in $\mathcal{L}(q; \boldsymbol{\theta}, t, M_t)$ involving $\beta$ are

$$\begin{aligned}
\mathcal{L}(\beta) =&\sum_{j=1}^{M_t} \omega_j \boldsymbol{\mu}_j^T \mathbf{A}(\beta)^T (\hat{\mathbf{S}}_j \hat{\mathbf{C}} \hat{\mathbf{S}}_j + \ell_j \hat{\boldsymbol{\Omega}})^{-1} \begin{pmatrix}\hat{\gamma}_j \\ \hat{\Gamma}_j\end{pmatrix} \\
&\sum_{j=1}^{M_t} -\frac{1}{2} \omega_j Tr\left[\mathbf{A}(\beta)^T (\hat{\mathbf{S}}_j \hat{\mathbf{C}} \hat{\mathbf{S}}_j + \ell_j \hat{\boldsymbol{\Omega}})^{-1} \mathbf{A}(\beta) (\boldsymbol{\Lambda}_j^{-1} + \boldsymbol{\mu}_j \boldsymbol{\mu}_j^T)\right].
\end{aligned}$$

Here we write $\mathbf{A}(\beta) = \begin{pmatrix} 1 & 0 \\ \beta & 1 \end{pmatrix}$ as $\mathbf{A}(\beta) = \mathbf{I}_2 + \beta \mathbf{V}_1$, where $\mathbf{I}_2 = \begin{pmatrix} 1 & 0 \\ 0 & 1 \end{pmatrix}$, and $\mathbf{V}_1 = \begin{pmatrix} 0 & 0 \\ 1 & 0 \end{pmatrix}$. Taking the derivative of $\mathcal{L}(\beta)$ w.r.t. $\beta$ and setting it to zero, the updating equation for $\beta$ is given as

$$\beta = \frac{\sum_{j=1}^{M_t} \omega_j \boldsymbol{\mu}_j^T \mathbf{V}_1^T (\hat{\mathbf{S}}_j \hat{\mathbf{C}} \hat{\mathbf{S}}_j + \ell_j \hat{\boldsymbol{\Omega}})^{-1} \begin{pmatrix} \hat{\gamma}_j \\ \hat{\Gamma}_j \end{pmatrix} - \omega_j \operatorname{Tr}(\mathbf{V}_1^T (\hat{\mathbf{S}}_j \hat{\mathbf{C}} \hat{\mathbf{S}}_j + \ell_j \hat{\boldsymbol{\Omega}})^{-1} (\boldsymbol{\Lambda}_j^{-1} + \boldsymbol{\mu}_j \boldsymbol{\mu}_j^T))}{\sum_{j=1}^{M_t} \omega_j \operatorname{Tr}\left[\mathbf{V}_1^T (\hat{\mathbf{S}}_j \hat{\mathbf{C}} \hat{\mathbf{S}}_j + \ell_j \hat{\boldsymbol{\Omega}})^{-1} \mathbf{V}_1 (\boldsymbol{\Lambda}_j^{-1} + \boldsymbol{\mu}_j \boldsymbol{\mu}_j^T)\right])}. \tag{S.19}$$

We next derive the updating equation for $\pi_t$. The terms in $\mathcal{L}(q; \boldsymbol{\theta}, t, M_t)$ involving $\pi_t$ are

$$\mathcal{L}(\pi_t) = \sum_{j=1}^{M_t} \omega_j \log \pi_t + \sum_{j=1}^{M_t} (1 - \omega_j) \log(1 - \pi_t).$$

By setting the derivative of $\mathcal{L}(\pi_t)$ w.r.t. $\pi_t$ to zero, we obtain

$$\pi_t = \frac{\sum_{j=1}^{M_t} \omega_j}{M_t}. \tag{S.20}$$

We then derive the updating equation for $\tau^2$. Denote $\boldsymbol{\mu}_j = (\mu_{\gamma_j}, \mu_{\alpha_j})$ and the diagonal elements in $\boldsymbol{\Lambda}_j^{-1}$ by $(\sigma_{\gamma_j}^2, \sigma_{\alpha_j}^2)$. The terms in $\mathcal{L}(q; \boldsymbol{\theta}, t, M_t)$ involving $\tau^2$ are given as

$$\mathcal{L}(\tau^2) = -\frac{1}{2} \sum_{j=1}^{M_t} \omega_j \log \tau^2 - \frac{1}{2} \sum_{j=1}^{M_t} \omega_j \frac{\mu_{\alpha_j}^2 + \sigma_{\alpha_j}^2}{\ell_j \tau^2}.$$

Therefore, we obtain the updating equation for $\tau^2$ as

$$\tau^2 = \frac{\sum_{j=1}^{M_t} \omega_j (\mu_{\alpha_j}^2 + \sigma_{\alpha_j}^2)/\ell_j}{\sum_{j=1}^{M_t} \omega_j}.$$

Finally, we derive the update for $\sigma^2$. The terms in $\mathcal{L}(q; \boldsymbol{\theta}, t, M_t)$ involving $\sigma^2$ are

$$\mathcal{L}(\sigma^2) = \sum_{j=1}^{M_t} -\frac{1}{2} \omega_j \log \sigma^2 - \frac{1}{2} \omega_j \frac{\mu_{\gamma_j}^2 + \sigma_{\gamma_j}^2}{l_j \sigma^2} - \omega_j \log \left( 2\Phi \left( \frac{-t \hat{s}_{X,j}}{\sqrt{\ell_j \sigma^2 + \ell_j \hat{\sigma}_u^2 + \hat{c}_1 \hat{s}_{X,j}^2}} \right) \right).$$

If $t = 0$, we directly set the derivative of $\mathcal{L}(\sigma^2)$ w.r.t. $\sigma^2$ to zero and obtain the update for $\sigma^2$:

$$\sigma^2 = \frac{\sum_{j=1}^{M_t} \omega_j (\mu_{\gamma_j}^2 + \sigma_{\gamma_j}^2)/\ell_j}{\sum_{j=1}^{M_t} \omega_j}.$$

If $t \neq 0$, direct maximization of $\mathcal{L}(\sigma^2)$ is intractable because of the normalization terms in the truncated Gaussians distributions. Instead, we can obtain a tractable lower bound for $\mathcal{L}(\sigma^2)$:

$$\mathcal{L}(\sigma^2) \geq -\sum_{j=1}^{M_t} \frac{1}{2} \frac{\omega_j}{\sigma^{2(old)}} (\sigma^2 - \sigma^{2(old)}) - \sum_{j=1}^{M_t} \frac{1}{2} \omega_j \frac{\mu_{\gamma_j}^2 + \sigma_{\gamma_j}^2}{\ell_j \sigma^2} -$$

$$\sum_{j=1}^{M_t} \frac{1}{2} \omega_j t \ell_j \hat{s}_{X,j} \frac{\phi\left( \frac{-t \hat{s}_{X,j}}{\sqrt{l_j \sigma^{2(old)} + \ell_j \hat{\sigma}_u^2 + \hat{c}_1 \hat{s}_{X,j}^2}} \right)}{\Phi\left( \frac{-t \hat{s}_{X,j}}{\sqrt{\ell_j \sigma^{2(old)} + \ell_j \hat{\sigma}_u^2 + \hat{c}_1 \hat{s}_{X,j}^2}} \right)} (\ell_j \sigma^{2(old)} + \ell_j \hat{\sigma}_u^2 + \hat{c}_1 \hat{s}_{X,j}^2)^{-\frac{3}{2}} (\sigma^2 - \sigma^{2(old)}),$$

where $\sigma^{2(old)}$ is the estimate of $\sigma^2$ from the previous step. To obtain the tractable lower bound, we use the facts that -log $\sigma^2$ and -log $\left(2\Phi\left(\frac{-t\hat{s}_{X,j}}{\sqrt{\ell_j\sigma^2+\ell_j\hat{\sigma}_u^2+\hat{c}_1\hat{s}_{X,j}^2}}\right)\right)$ are concave w.r.t. $\sigma^2$. Then we can maximize the tractable lower bound w.r.t. $\sigma^2$ to obtain the update for $\sigma^2$ as

$$\sigma^2 = \sqrt{\frac{\sum_{j=1}^{M_t}\omega_j(\mu_{\gamma_j}^2+\sigma_{\gamma_j}^2)/\ell_j}{\sum_{j=1}^{M_t}\frac{\omega_j}{\sigma^{2(old)}}+\sum_{j=1}^{M_t}\omega_j t\ell_j\hat{s}_{X,j}\frac{\phi\left(\frac{-t\hat{s}_{X,j}}{\sqrt{\ell_j\sigma^{2(old)}+\ell_j\hat{\sigma}_u^2+\hat{c}_1\hat{s}_{X,j}^2}}\right)}{\Phi\left(\frac{-t\hat{s}_{X,j}}{\sqrt{\ell_j\sigma^{2(old)}+\ell_j\hat{\sigma}_u^2+\hat{c}_1\hat{s}_{X,j}^2}}\right)}(\ell_j\sigma^{2(old)}+\ell_j\hat{\sigma}_u^2+\hat{c}_1\hat{s}_{X,j}^2)^{-\frac{3}{2}}}}.$$

#### 1.5 Sensitivity Analysis

In the foreground model of MR-APSS, we assume that the direct effect $\alpha_j$ is independent of the IV strength $\gamma_j$, i.e., $r_f = \text{Corr}(\gamma_j, \alpha_j) = 0$ (the subscript $f$ refers to the foreground model). In other words, we assume that the association between the exposure and the outcome is induced by their causal relationship rather than $r_f$ after accounting for confounding factors in the background model, such as correlated pleiotropy and sample structure. Although our method relies on this assumption to infer the causal effect, we can empirically check the influence of this assumption via the following sensitivity analysis. We can check how $\hat{\beta}$ changes when $r_f$ is fixed at different values. Let's consider a real example for BMI and T2D where MR-APSS reported the causal effect between BMI and T2D $\hat{\beta} = 0.328$ with $p$-value $= 6.77 \times 10^{-9}$. We first set $\beta = 0$ and obtained the foreground correlation as $\hat{r}_f = 0.330$. This means that the foreground correlation is at most 0.330 even in the absence of causal effects. Then we varied $r_f$ at a grid of values $r_f \in \{0, 0.033, \ldots, 0.330\}$ and then re-estimated $\beta$. As shown in supplementary Fig. S13, the estimated causal effect $\hat{\beta}$ varied as $r_f$ was set to different values. Clearly, the results of sensitivity analysis indicate that the causal effect between BMI and T2D remained to be significant as long as $r_f < 0.198$.

In this analysis, we need to estimate parameters $(\beta, \pi_t, \mathbf{\Sigma})$ when we assume that the IV strength $(\gamma_j)$ and the direct effect $(\alpha_j)$ are not independent and the correlation parameter $r_f$ is set to a non-zero value $(r_f \neq 0)$. The only change to the EM algorithm of MR-APSS (see SI Appendix, section 1.4.2) is the update function for $\mathbf{\Sigma}$. We now derive the update function for $\mathbf{\Sigma}$ when $r_f$ is set to be none-zero. We rewrite $\mathbf{\Sigma}$ as

$$\mathbf{\Sigma} = \begin{pmatrix}\sigma' & 0\\ 0 & \tau'\end{pmatrix}\begin{pmatrix}1 & r_f\\ r_f & 1\end{pmatrix}\begin{pmatrix}\sigma' & 0\\ 0 & \tau'\end{pmatrix} = \mathbf{\Sigma}_0\mathbf{R}\mathbf{\Sigma}_0, \tag{S.21}$$

where $\mathbf{\Sigma}_0 = \begin{pmatrix}\sigma' & 0\\ 0 & \tau'\end{pmatrix}$ and $\mathbf{R} = \begin{pmatrix}1 & r_f\\ r_f & 1\end{pmatrix}$. Because $\mathbf{R} = \begin{pmatrix}1 & r_f\\ r_f & 1\end{pmatrix}$ is fixed and known, we only need to obtain the update function for $\mathbf{\Sigma}_0$.

Recall that terms in $\mathcal{L}(q;\boldsymbol{\theta},t,M_t)$ given in Eq. [S.18] involving $\boldsymbol{\Sigma}$ are

$$\mathcal{L}(\boldsymbol{\Sigma}) = \sum_{j=1}^{M_t} -\frac{1}{2}\log\det(\ell_j\boldsymbol{\Sigma}) - \frac{1}{2}\omega_j\ell_j^{-1}\boldsymbol{\mu}_j^T\boldsymbol{\Sigma}^{-1}\boldsymbol{\mu}_j - \frac{1}{2}\operatorname{Tr}\left[\omega_j\ell_j^{-1}\boldsymbol{\Sigma}^{-1}\boldsymbol{\Lambda}_j^{-1}\right]+$$

$$\sum_{j=1}^{M_t}\omega_j\log\left(2\Phi\left(\frac{-t\hat{s}_{X,j}}{\sqrt{\ell_j\sigma^2+\ell_j\hat{\sigma}_u^2+\hat{c}_1\hat{s}_{X,j}^2}}\right)\right) - (1-\omega_j)\log\left(2\Phi\left(\frac{-t\hat{s}_{X,j}}{\sqrt{\ell_j\hat{\sigma}_u^2+\hat{c}_1\hat{s}_{X,j}^2}}\right)\right) + \text{constant}.$$

This function can be bounded by

$$\mathcal{L}(\boldsymbol{\Sigma}) \geq \sum_{j=1}^{M_t} -\frac{1}{2}\omega_j\log\det(\boldsymbol{\Sigma}^{(old)}) - \frac{1}{2}\omega_j\operatorname{Tr}\left(\boldsymbol{\Sigma}^{-(old)}(\boldsymbol{\Sigma}-\boldsymbol{\Sigma}^{(old)})\right) -$$

$$\sum_{j=1}^{M_t}\frac{1}{2}\omega_j\ell_j^{-1}\boldsymbol{\mu}_j^T\boldsymbol{\Sigma}^{-1}\boldsymbol{\mu}_j - \frac{1}{2}\operatorname{Tr}\left[\omega_j\ell_j^{-1}\boldsymbol{\Sigma}^{-1}\boldsymbol{\Lambda}_j^{-1}\right]-$$

$$\sum_{j=1}^{M_t}\frac{1}{2}\omega_j\ell_j t\hat{s}_{X,j}\frac{\phi\left(\frac{-t\hat{s}_{X,j}}{\sqrt{\ell_j\sigma^2+\ell_j\hat{\sigma}_u^2+\hat{c}_1\hat{s}_{X,j}^2}}\right)}{\Phi\left(\frac{-t\hat{s}_{X,j}}{\sqrt{\ell_j\sigma^2+\ell_j\hat{\sigma}_u^2+\hat{c}_1\hat{s}_{X,j}^2}}\right)}(\ell_j\sigma^2+\ell_j\hat{\sigma}_u^2+\hat{c}_1\hat{s}_{X,j}^2)^{-3/2}\mathbf{b}_1^T(\boldsymbol{\Sigma}-\boldsymbol{\Sigma}^{(old)})\mathbf{b}_1,$$

where $\mathbf{b}_1^T = \begin{pmatrix}1 & 0\end{pmatrix}$ and $\boldsymbol{\Sigma}^{(old)}$ is the estimate of $\boldsymbol{\Sigma}$ from the previous step. By taking the derivative of $\mathcal{L}(\boldsymbol{\Sigma})$ with respect to $\boldsymbol{\Sigma}_0$ and setting it to zero, we have

$$\sum_{j=1}^{M_t} -\omega_j\boldsymbol{\Sigma}^{-(old)}\boldsymbol{\Sigma}_0\mathbf{R} + \sum_{j=1}^{M_t}\omega_j\ell_j^{-1}\boldsymbol{\Sigma}_0^{-1}\mathbf{R}^{-1}\boldsymbol{\Sigma}_0^{-1}(\boldsymbol{\mu}_j\boldsymbol{\mu}_j^T+\boldsymbol{\Lambda}_j^{-1})\boldsymbol{\Sigma}_0^{-1}-$$

$$\sum_{j=1}^{M_t}\omega_j\ell_j t\hat{s}_{X,j}\frac{\phi\left(\frac{-t\hat{s}_{X,j}}{\sqrt{\ell_j\sigma^2+\ell_j\hat{\sigma}_u^2+\hat{c}_1\hat{s}_{X,j}^2}}\right)}{\Phi\left(\frac{-t\hat{s}_{X,j}}{\sqrt{\ell_j\sigma^2+\ell_j\hat{\sigma}_u^2+\hat{c}_1\hat{s}_{X,j}^2}}\right)}(\ell_j\sigma^2+\ell_j\hat{\sigma}_u^2+\hat{c}_1\hat{s}_{X,j}^2)^{-3/2}\mathbf{b}_1\mathbf{b}_1^T\boldsymbol{\Sigma}_0\mathbf{R} = 0.$$

Denote $\mathbf{B}_1 = \sum_{j=1}^{M_t}\omega_j\boldsymbol{\Sigma}^{-(old)} + \sum_{j=1}^{M_t}\omega_j\ell_j t\hat{s}_{X,j}\frac{\phi\left(\frac{-t\hat{s}_{X,j}}{\sqrt{\ell_j\sigma^2+\ell_j\hat{\sigma}_u^2+\hat{c}_1\hat{s}_{X,j}^2}}\right)}{\Phi\left(\frac{-t\hat{s}_{X,j}}{\sqrt{\ell_j\sigma^2+\ell_j\hat{\sigma}_u^2+\hat{c}_1\hat{s}_{X,j}^2}}\right)}(\ell_j\sigma^2+\ell_j\hat{\sigma}_u^2+\hat{c}_1\hat{s}_{X,j}^2)^{-3/2}\mathbf{b}_1\mathbf{b}_1^T$, and $\mathbf{B}_2 = \sum_{j=1}^{M_t}\omega_j\ell_j^{-1}(\boldsymbol{\mu}_j\boldsymbol{\mu}_j^T+\boldsymbol{\Lambda}_j^{-1})$. We can rewrite the above equation as following

$$\mathbf{B}_1 = \boldsymbol{\Sigma}_0^{-1}\mathbf{R}^{-1}\boldsymbol{\Sigma}_0^{-1}\mathbf{B}_2\boldsymbol{\Sigma}_0^{-1}\mathbf{R}^{-1}\boldsymbol{\Sigma}_0^{-1}.$$

To solve the above equation, we use the following lemma from matrix computation [42]: Given two positive definite matrices $\mathbf{A}$ and $\mathbf{B}$, they are related with the matrix equation $\mathbf{B} = \mathbf{X}^{-1}\mathbf{A}\mathbf{X}^{-1}$. Then $\mathbf{Y} = \mathbf{L}^{-T}(\mathbf{L}^T\mathbf{A}\mathbf{L})^{1/2}\mathbf{L}^{-1}$ is the unique positive definite solution to the matrix equation, where $\mathbf{L}$ is the Cholesky factor of $\mathbf{B}$. By applying this lemma, we have

$$\boldsymbol{\Sigma}_0\mathbf{R}\boldsymbol{\Sigma}_0 = \mathbf{L_1}^{-T}(\mathbf{L_1}^T\mathbf{B}_2\mathbf{L_1})^{1/2}\mathbf{L_1}^{-1}, \tag{S.22}$$

where $\mathbf{L_1}$ is the Cholesky factor of $\mathbf{B}_1$ satisfying $\mathbf{B}_1 = \mathbf{L_1}\mathbf{L_1}^T$. To obtain $\boldsymbol{\Sigma}_0$, we apply the above lemma to Eq. [S.22] again and then obtain

$$\boldsymbol{\Sigma}_0 = \mathbf{L_R}^{-T}(\mathbf{L_R}^T(\mathbf{L_1}^{-T}(\mathbf{L_1}^T\mathbf{B}_2\mathbf{L_1})^{1/2}\mathbf{L_1}^{-1})\mathbf{L_R})^{1/2}\mathbf{L_R}^{-1},$$

where $\mathbf{L_R}$ is the Cholesky factor of $\mathbf{R}$ satisfying $\mathbf{R} = \mathbf{L_R}\mathbf{L_R}^T$. Then we set the non-diagonal elements of $\boldsymbol{\Sigma}_0$ to zero and update $\boldsymbol{\Sigma}$ using Eq. [S.21].

#### 1.6 Adjustment of bias due to LD clumping

Besides the bias due to the $p$-value thresholding, we are aware of the selection bias due to the LD clumping procedure. This is because SNPs with smaller $p$-values are selected as IVs in the LD clumping procedure. As shown in Fig. S10 (Left panel), the median of $p$-values from IVs after LD clumping is smaller than that of IVs before LD clumping. To account for the bias due to LD clumping, we propose the following adjustment on the $p$-value threshold in MR-APSS:

$$p\text{-value threshold} \leftarrow \text{IV threshold} \times \min\left(\frac{\text{median}(p\text{-values}_{\text{after}})}{\text{median}(p\text{-values}_{\text{before}})}, 1\right),$$

where $p$-values$_{\text{before}}$ and $p$-values$_{\text{after}}$ correspond to the $p$-values from IVs before LD clumping and after LD clumping. As shown in the formula, we adjust the IV threshold by the ratio of the median of $p$-values.

To examine the effectiveness of the adjustment, we compared the results of MR-APSS with the adjusted $p$-value threshold and its non-adjusted version. As we know, selection bias can lead to over-estimation of foreground signals and thus more invalid IVs will be falsely detected as valid IVs, resulting in an inflated type I error rate. Therefore, we first examined the ability of the two methods (MR-APSS with / without $p$-value threshold adjustment) on the detection of effective IVs. As shown in Fig. S10 (Right panel), we observed a reasonable increase of effective IVs as the IV threshold became looser using MR-APSS with the adjusted threshold. However, the number of effective IVs detected by MR-APSS without threshold adjustment increased sharply as the IV threshold became relaxed. We next examined the $p$-values from causal inference provided by the two methods. As shown in Fig. S9, the $p$-values provided by MR-APSS without threshold adjustment were inflated. Meanwhile, based on our proposed threshold adjustment, the $p$-values were uniformly distributed without inflation, indicating the effectiveness of the adjustment.

#### 1.7 Binary traits

Similar to existing summary-level MR-methods, we consider linear models to perform causal inference even for binary traits. There are two major reasons. First, most of the released GWAS summary statistics are obtained under linear models [22]. As long as the case-control ratio is not extremely unbalanced, linear models are known to work well when they are applied to binary traits (0-1 observed scale) in GWASs [22, 43]. This is because a linear model can be viewed as a first-order approximation to the liability model [20, 44]. Second, the effect sizes or heritability estimated using linear models can be transformed to the liability scale [20, 44, 8] or odds ratio [10, 28].

To have better interpretation of the causal effect estimates for binary traits, here we show that the analysis result of traits in the observed 0-1 scale based on linear models can be transformed to the liability scale based on the probit model. Specifically, we consider the following three cases with a binary exposure or a binary outcome: (a) a continuous exposure and a binary outcome; (b) a binary exposure and a continuous outcome; and (c) a binary exposure and a binary outcome. We show that the causal effect estimate obtained with linear models is still interpretable for these three cases.

Case (a): a continuous exposure ($X$) and a binary outcome ($Y$).

For a continuous exposure $X$, we consider a linear model to relate genotypes with phenotypes:

$$x_i = b_{0,\mathrm{x}} + \mathbf{W}_i^T \mathbf{b}_{\mathrm{cov,x}} + \mathbf{G}_i^T(\mathbf{Z}\boldsymbol{\gamma} + \mathbf{u}) + e_{1i}, \quad \text{(S.23)}$$

where $x_i$ is the $i$-th individual's phenotypic value of the exposure, $\mathbf{G}_i$ is an $M \times 1$ genotype vector,$\mathbf{W}_i$ is the covariate vector, and $\mathbf{b}^b_{\mathrm{cov,x}}$ is the corresponding coefficient vector. $\mathbf{Z}$ is an $M \times M$ diagonal matrix with the $j$-th entry $Z_j \in \{0, 1\}$ indicating whether the $j$-th SNP is an effective IV with a foreground effect, $\boldsymbol{\gamma} = \{\gamma_j\}_{j=1,\ldots,M}$ is a vector of the instrument strength, $\mathbf{u} = \{u_j\}_{j=1,\ldots,M}$ is a vector of the polygenic effects on $X$, and $e_{1i}$ is the independent noise term.

For a binary outcome trait $Y$, we consider the following probit model (which is also known as the liability model in genetics [20]):

$$p\,(y_i = 1|\mathbf{G}_i, \mathbf{W}_i) = \Phi\big(b^b_{0,\mathrm{y}} + \mathbf{W}_i^T \mathbf{b}^b_{\mathrm{cov,y}} + \beta^b x_i + \mathbf{G}_i^T \mathbf{Z}\boldsymbol{\alpha}^b + \mathbf{G}_i^T \mathbf{v}'^b\big), \quad \text{(S.24)}$$

where $y_i \in \{0, 1\}$ is the phenotypic value of the $i$-th individual, $b^b_{0,\mathrm{y}}$ is the intercept term, $\mathbf{b}^b_{\mathrm{cov,y}}$ is the corresponding coefficient vector of covariates, $\beta^b$ represents the causal effect of $X$ on $Y$ in the liability scale, $\mathbf{v}'^b = \{v'^b_j\}_{j=1,\ldots,M}$ is an $M \times 1$ vector of SNP effect sizes, $\boldsymbol{\alpha}^b = \{\alpha^b_j\}_{j=1,\ldots,M}$ is a vector of direct effects, $\Phi(\cdot)$ is the cumulative distribution function of standard normal distribution, and the superscript $b$ denotes the coefficient of a binary trait in the liability scale.

Plugging Eq. [S.23] into Eq. [S.24], we have

$$p\,(y_i = 1|\mathbf{G}_i, \mathbf{W}_i) = \Phi\big(b^b_0 + \mathbf{W}_i^T \mathbf{b}^b_{\mathrm{cov}} + \mathbf{G}_i^T \mathbf{Z}(\beta^b\boldsymbol{\gamma} + \boldsymbol{\alpha}^b) + \mathbf{G}_i \mathbf{v}^b + \beta^b e_{1i}\big),$$

where $b^b_0 = b_{0,\mathrm{x}}\beta + b_{0,\mathrm{y}}$, $\mathbf{b}^b_{\mathrm{cov}} = \mathbf{b}_{\mathrm{cov,x}}\beta + \mathbf{b}_{\mathrm{cov,y}}$, and $\mathbf{v}^b = \beta^b\mathbf{u} + \mathbf{v}'^b$ which represents the polygenic effects of genotypes on $Y$ in the liability scale.

To make the notation simple, we rewrite the above model as

$$p\,(y_i = 1|\mathbf{G}_i, \mathbf{W}_i) = \Phi\big(b^b_0 + \mathbf{W}_i \mathbf{b}^b_{\mathrm{cov}} + \mathbf{G}_i\boldsymbol{\gamma}^b + \beta^b e_{1i}\big),$$

where $\boldsymbol{\gamma}^b = \mathbf{Z}(\beta^b\boldsymbol{\gamma} + \boldsymbol{\alpha}^b) + \mathbf{v}^b$ is an $M \times 1$ vector collecting the genetic effects on $Y$ in the liability scale. We denote the $j$-th element of $\boldsymbol{\gamma}^b$ as

$$\Gamma^b_j = Z_j(\beta^b\gamma_j + \alpha^b_j) + v^b_j.$$

With the above preparation, we can apply the known results in [20, 44] (e.g., see Eq. [76] in Text S3 of [44]) to obtain a linear approximation of $p\,(y_i = 1|\mathbf{G}_i, \mathbf{W}_i)$ as

$$p\,(y_i = 1|\mathbf{G}_i, \mathbf{W}_i) \approx k_2 + \frac{k_2(1-k_2)\phi(b^b_0)}{K_2(1-K_2)}\big(\mathbf{W}_i\mathbf{b}^b_{\mathrm{cov}} + \mathbf{G}_i\boldsymbol{\gamma}^b + \beta^b e_{1i}\big),$$

where $k_2$ is the case proportion in the ascertained case-control sample, $K_2$ is the case proportion in the population, and $\phi(\cdot)$ is the normal density function. This implies that the effect sizes estimated by linear regression, $\Gamma_j$, can be transformed into the liability scale $\Gamma^b_j$ by

$$\Gamma^b_j = \frac{K_2(1-K_2)}{k_2(1-k_2)\phi(b^b_0)}\Gamma_j.$$

Consequently, we have

$$Z_j(\beta^b\gamma_j + \alpha_j^b) + v_j^b = \frac{K_2(1-K_2)}{k_2(1-k_2)\phi(b_0^b)}(Z_j(\beta\gamma_j + \alpha_j) + v_j),$$

where $\beta^b$ is the causal effect in the liability scale and $\beta$ is the causal effect obtained by the linear model. Therefore, we can first obtain the causal effect with linear models and then transform it back to the liability scale

$$\beta^b = \frac{K_2(1-K_2)}{k_2(1-k_2)\phi(b_0^b)}\beta.$$

In our paper, we perform hypothesis testing ($H_0$: $\beta = 0$ vs $H_A$: $\beta \neq 0$) to examine the significance of causal relationship. The testing result can be directly applied to examine whether the causal effect exists in the liability scale ($H_0$: $\beta^b = 0$ vs $H_A$: $\beta^b \neq 0$).

Case (b): a binary exposure and a continuous outcome

For a binary exposure $X$, we again consider the following probit model:

$$p\left(x_i = 1|\mathbf{G}_i, \mathbf{W}_i\right) = \Phi\big(b_{0,\text{x}}^b + \mathbf{W}_i^T\mathbf{b}_{\text{cov,x}}^b + \mathbf{G}_i^T(\mathbf{Z}\boldsymbol{\gamma}^b + \mathbf{u}^b)\big), \tag{S.25}$$

where $b_{0,\text{x}}^b$ is the intercept term, $\mathbf{b}_{\text{cov,x}}^b$ is the coefficient vector of covariates, $\boldsymbol{\gamma}^b = \{\gamma_j^b\}_{j=1,\ldots,M}$ is an $M \times 1$ vector of foreground effects, $\mathbf{u}^b = \{u_j^b\}_{j=1,\ldots,M}$ is an $M \times 1$ vector of background effects, and the superscript $b$ denotes the coefficient of a binary trait in the liability scale.

For a continuous outcome $Y$, we consider the following linear model,

$$y_i = b_{0,\text{y}} + \mathbf{W}_i^T\mathbf{b}_{\text{cov,y}} + \beta x_i + \mathbf{G}_i^T(\mathbf{Z}\boldsymbol{\alpha} + \mathbf{v}') + e_{2i}, \tag{S.26}$$

where $b_{0,\text{y}}$ is the intercept term, $\mathbf{b}_{\text{cov,y}}$ is the corresponding coefficient vector of covariates, $\mathbf{v} = \{v_j\}_{j=1,\ldots,M}$ is an $M \times 1$ vector of background effects, $\boldsymbol{\alpha} = \{\alpha_j\}_{j=1,\ldots,M}$ is the vector of direct effect on $y_i$, and $\beta$ is the causal effect of interest.

With the above preparation, we can apply the known results in [20, 44] to obtain an approximation of $p\left(x_i = 1|\mathbf{G}_i, \mathbf{W}_i\right)$ as

$$p\left(x_i = 1|\mathbf{G}_i, \mathbf{W}_i\right) = \mathbb{E}(x_i|\mathbf{G}_i, \mathbf{W}_i) \approx k_1 + \frac{k_1(1-k_1)\phi(b_{0,\text{x}}^b)}{K_1(1-K_1)}\big(\mathbf{W}_i^T\mathbf{b}_{\text{cov,x}}^b + \mathbf{G}_i^T\mathbf{Z}\boldsymbol{\gamma}^b + \mathbf{G}_i^T\mathbf{u}^b\big),$$

where $k_1$ is the case proportion in the ascertained case-control sample of the exposure, $K_1$ is the case proportion in the population of the exposure, and $\phi(\cdot)$ is the normal density function. The above equation suggests that we can obtain a linear approximation for the binary trait $X$,

$$x_i \approx b_{0,\text{x}} + \mathbf{W}_i^T\mathbf{b}_{\text{cov,x}} + \mathbf{G}_i^T\mathbf{Z}\boldsymbol{\gamma} + \mathbf{G}_i^T\mathbf{u}, \tag{S.27}$$

where $b_{0,\text{x}} = k_1, \mathbf{b}_{\text{cov,x}} = \frac{k_1(1-k_1)\phi(b_{0,\text{x}}^b)}{K_1(1-K_1)}\mathbf{b}_{\text{cov,x}}^b, \boldsymbol{\gamma} = \frac{k_1(1-k_1)\phi(b_{0,\text{x}}^b)}{K_1(1-K_1)}\boldsymbol{\gamma}^b$, and $\mathbf{u} = \frac{k_1(1-k_1)\phi(b_{0,\text{x}}^b)}{K_1(1-K_1)}\mathbf{u}^b$. Note that $\boldsymbol{\gamma}$ and $\mathbf{u}$ are vectors of effects in linear scale.

Plugging Eq. [S.27] into Eq. [S.26], we have

$$y_i = b_0 + \mathbf{W}_i^T\mathbf{b}_{\text{cov}} + \mathbf{G}_i^T(\mathbf{Z}(\beta\boldsymbol{\gamma} + \boldsymbol{\alpha}) + \mathbf{v}) + e_{2i},$$

where $b_0 = b_{0,\text{y}} + \beta b_{0,\text{x}}$, $\mathbf{b}_{\text{cov}} = \mathbf{b}_{\text{cov,y}} + \beta\mathbf{b}_{\text{cov,x}}$, and $\mathbf{v} = \beta\mathbf{u} + \mathbf{v}'$ which represents the vector of polygenic effects of genotypes on $Y$. We denote the $j$-th element of $\boldsymbol{\gamma}$, the $j$-th element of $\mathbf{u}$, and the $j$-th element of $\mathbf{v}$ as $\gamma_j$, $u_j$, and $v_j$, respectively.

To make the notation simple, we rewrite the above model as

$$y_i = b_0 + \mathbf{W}_i^T \mathbf{b}_{\text{cov}} + \mathbf{G}_i^T \mathbf{\Gamma} + e_{2i},$$

where $\mathbf{\Gamma} = \mathbf{Z}(\beta\boldsymbol{\gamma} + \boldsymbol{\alpha}) + \mathbf{v}$. The $j$-th element of $\mathbf{\Gamma}$ can be expressed as

$$\Gamma_j = Z_j(\beta\gamma_j + \alpha_j) + v_j.$$

This implies that we can obtain a good approximation of the causal effect $\beta$ using linear models for binary exposure $X$ and continuous outcome $Y$.

Case (c): a binary exposure and a binary outcome

Similarly, for a binary exposure $X$, we consider a probit model given in Eq. [S.25]. We then apply the known results in [20, 44] to obtain a linear approximation for the exposure which is given in Eq. [S.27]. For a binary outcome $Y$, we consider the probit model given in Eq. [S.24].

Plugging Eq. [S.27] into Eq. [S.24], we have

$$p\,(y_i = 1|\mathbf{G}_i, \mathbf{W}_i) = \Phi\big(b_0^b + \mathbf{W}_i^T \mathbf{b}_{\text{cov}}^b + \mathbf{G}_i^T \mathbf{Z}(\beta^b\boldsymbol{\gamma} + \boldsymbol{\alpha}^b) + \mathbf{G}_i \mathbf{v}^b\big),$$

where $b_0^b = b_{0,\text{y}}^b + \beta^b b_{0,\text{x}}$ is the intercept term, $\mathbf{b}_{\text{cov}}^b = \mathbf{b}_{\text{cov,y}}^b + \beta^b \mathbf{b}_{\text{cov,x}}$ is the coefficient vector of covariates, and $\mathbf{v}^b = \beta^b \mathbf{u} + \mathbf{v}'^b$ represents the vector of polygenic effects on $Y$ in the liability scale.

Again, we can rewrite the above model as

$$p\,(y_i = 1|\mathbf{G}_i, \mathbf{W}_i) = \Phi\big(b_0^b + \mathbf{W}_i^T \mathbf{b}_{\text{cov}}^b + \mathbf{G}_i \mathbf{\Gamma}^b\big),$$

where $\mathbf{\Gamma}^b = \mathbf{Z}(\beta^b\boldsymbol{\gamma} + \boldsymbol{\alpha}^b) + \mathbf{v}^b$ is an $M \times 1$ vector collecting the genetic effects on $Y$ in the liability scale. We denote the $j$-th element of $\mathbf{\Gamma}^b$ as

$$\Gamma_j^b = Z_j(\beta^b\gamma_j + \alpha_j^b) + v_j^b.$$

Now, we apply the known results in [20, 44] to obtain a linear approximation of $p\,(y_i = 1|\mathbf{G}_i, \mathbf{W}_i)$ as

$$p\,(y_i = 1|\mathbf{G}_i, \mathbf{W}_i) \approx k_2 + \frac{k_2(1-k_2)\phi(b_0^b)}{K_2(1-K_2)}\big(\mathbf{W}_i \mathbf{b}_{\text{cov}}^b + \mathbf{G}_i \mathbf{\Gamma}^b\big),$$

where $k_2$ is the case proportion in the ascertained case-control sample of the outcome, $K_2$ is the case proportion in the population of the outcome, and $\phi(\cdot)$ is the normal density function. This implies that the effect sizes estimated by linear regression, $\Gamma_j$, can be transformed into the liability scale $\Gamma_j^b$ by

$$\Gamma_j^b = \frac{K_2(1-K_2)}{k_2(1-k_2)\phi(b_0^b)}\Gamma_j.$$

Consequently, we have

$$Z_j(\beta^b\gamma_j + \alpha_j^b) + v_j^b = \frac{K_2(1-K_2)}{k_2(1-k_2)\phi(b_0^b)}(Z_j(\beta\gamma_j + \alpha_j) + v_j),$$

where $\beta^b$ is the causal effect in the liability scale and $\beta$ is the causal effect obtained by the linear model. Therefore, we can first obtain the causal effect with linear models and then transform it back to the liability scale

$$\beta^b = \frac{K_2(1-K_2)}{k_2(1-k_2)\phi(b_0^b)}\beta.$$

From the above derivation, we can conclude that the causal effect estimate obtained with linear models is still interpretable for the three cases.

#### 1.8 Theoretical justification of the uniformity of the approximated distribution for GWAS summary statistics

Based on the Berry-Essen theorem [4, 9], we can show that the uniformity of the approximated distribution of the summary statistics $\hat{\gamma}_j$ and $\hat{\Gamma}_j$ for all $j$ can be guaranteed when the third absolute moment of phenotype and the genotype variables are bounded by finite values. Given the true marginal effect of SNP $j$ on the exposure, denoted as $\tilde{\gamma}_j, j = 1, \ldots, M$, we first provide detailed proof for the uniformity of the normality approximation of the conditional distribution of $\hat{\gamma}_j|\tilde{\gamma}_j$ for all $j$ (see proposition 1 below). We next show that $\hat{\gamma}_j$ follows a two-component Gaussian mixtures uniformly for all $j$ under the MR-APSS model (Main text, Eq. [1]). Analogously, we can obtain the uniformity of the approximation of distribution of $\hat{\Gamma}_j$ and the uniformity of the approximation of the joint distribution of $(\hat{\gamma}_j, \hat{\Gamma}_j)$ for all $j$ (Main text, Eq. [6]).

**(a). Uniform normal approximation of $\hat{\gamma}_j|\tilde{\gamma}_j$ for all $j$**

Given the true marginal effect of SNP $j$ on the exposure, denoted as $\tilde{\gamma}_j, j = 1, \ldots, M$, we first derive the uniform normal approximation of the conditional distribution for $\hat{\gamma}_j|\tilde{\gamma}_j$.

**Proposition 1 (Uniformity of normal approximation of** $\hat{\gamma}_j|\tilde{\gamma}_j$ for all $j$**).** Under the model given in Eq. [S.9] and summary statistics $(\hat{\gamma}_j,\ \hat{s}_{X,j})$ given in Eq. [S.10], and if $\mathbb{E}(|x_i|^3) \leq C_1 < \infty$ and $\mathbb{E}(|G_{1,ij}|^3) \leq C_2 < \infty$ for any $i, j$, then the conditional distribution of $\hat{\gamma}_j|\tilde{\gamma}_j, j = 1, \ldots, M$, uniformly in distribution converges to a normal distribution, i.e.

$$\frac{(\hat{\gamma}_j - \tilde{\gamma}_j)}{\hat{s}_{X,j}} \xrightarrow{d} \mathcal{N}(0, 1), \text{uniformly for } j = 1, \ldots, M,$$

*Proof.* By Eq. [S.10], $\hat{s}_{X,j}$ is replaced by $\sqrt{1/N_1}$. Let's denote the cumulative distribution function (cdf) of $\sqrt{N_1}(\hat{\gamma}_j - \tilde{\gamma}_j)$ by $F_{N_1,j}(\cdot)$ and denote the cdf of the standard normal distribution by $\Phi(\cdot)$.

Given $\hat{\gamma}_j = \frac{\mathbf{G}_{1,j}^T \mathbf{x}}{\mathbf{G}_{1,j}^T \mathbf{G}_{1,j}}$ and $\frac{\mathbf{G}_{1,j}^T \mathbf{G}_{1,j}}{N_1} = \frac{\sum_{i=1}^{N_1} \mathbf{G}_{1,ij}^2}{N_1} = 1$, we have

$$\begin{aligned}
\sqrt{N_1}(\hat{\gamma}_j - \tilde{\gamma}_j) &= \sqrt{N_1}\left(\frac{\mathbf{G}_{1,j}^T \mathbf{x}}{N_1} - \frac{\mathbf{G}_{1,j}^T \mathbf{G}_{1,j}}{N_1}\tilde{\gamma}_j\right) \\
&= \frac{1}{\sqrt{N_1}}\mathbf{G}_{1,j}^T(\mathbf{x} - \mathbf{G}_{1,j}\tilde{\gamma}_j) \\
&= \frac{1}{\sqrt{N_1}}\mathbf{G}_{1,j}^T \mathbf{e}_j \\
&= \frac{1}{\sqrt{N_1}}\sum_{i=1}^{N_1} G_{1,ij} e_{ij} \\
&\triangleq \frac{1}{\sqrt{N_1}}\sum_{i=1}^{N_1} \zeta_{ij},
\end{aligned}$$

where $\zeta_{ij} = G_{1,ij} e_{ij}$. Given $\mathbb{E}(G_{1,ij}) = 0$, $\mathrm{Var}(G_{1,ij}) = 1$, $\mathbb{E}(e_{ij}) = 0$, and $\mathrm{Var}(e_{ij}) \approx 1$, we have $\mathbb{E}(\zeta_{ij}) = 0$ and $\mathbb{E}(\zeta_{ij}^2) = \mathbb{E}(G_{1,ij}^2)\,\mathbb{E}(e_{ij}^2) \approx 1$, for any $i, j$. Because $\mathbb{E}(|G_{1,ij}|^3) < C_1 < \infty$, and $\mathbb{E}(|e_{ij}|^3) = \mathbb{E}(|x_i - G_{1,ij}\tilde{\gamma}_j|^3) \leq 4(\mathbb{E}(|x_i|^3) - \mathbb{E}(|G_{1,ij}\tilde{\gamma}_j|^3)) = 4\,\mathbb{E}(|x_i|^3) - 4\,\mathbb{E}(|G_{1,ij}|^3)\,\mathbb{E}(|\tilde{\gamma}_j|^3) \leq 4C_1 - 4C_2 * r_3 \leq \infty$, where $\mathbb{E}(|\tilde{\gamma}_j|^3) = r_3 < \infty$, we have $\mathbb{E}\,|\zeta_{ij}^*|^3 = \mathbb{E}(|G_{1,ij}|^3)\,\mathbb{E}(|e_{ij}|^3) \leq C_1(4C_1 - 4C_2 * r_3) = C^* < \infty$, for any $i, j$.

According to the Berry-Essen theorem [4, 9], we have

$$\sup_x |F_{N_1,j}(x) - \Phi(x)| \leq C_0\psi_0, \text{ for } j = 1, \ldots, M,$$

where $C_0$ is an absolute constant, and $\psi_0 = N_1^{-3/2} \sum_{i=1}^{N_1} \mathbb{E}\,|\zeta_{ij}|^3 < \frac{1}{\sqrt{N_1}} C^*$.

It means that, for any $j$, we have

$$\sup_x |F_{N_1,j}(x) - \Phi(x)| \leq \frac{C_0}{\sqrt{N_1}} C^* \to 0, \text{ as } N_1 \to \infty,$$

Hence, we can obtain

$$\sqrt{N_1}(\hat{\gamma}_j - \tilde{\gamma}_j) \xrightarrow{d} \mathcal{N}(0, 1) \text{ uniformly for } j = 1, \ldots, M.$$

We thus obtain a uniform normal approximation of the conditional distribution of $\hat{\gamma}_j|\tilde{\gamma}_j \sim \mathcal{N}(\tilde{\gamma}_j, \hat{s}^2_{X,j})$ where $\hat{s}_{X,j} = \frac{1}{\sqrt{N_1}}$. □

**(b). Uniform approximation of the distribution of $\hat{\gamma}_j$**

We first consider the case that SNPs are independent of each other (i.e., there is no LD effects between SNPs). Based on our model assumption in Eq. [S.9], we have

$$\tilde{\gamma}_j = Z_j\gamma_j + u_j,$$

and

$$\tilde{\gamma}_j \sim \pi_0\mathcal{N}(0, \sigma^2 + \sigma_u^2) + (1 - \pi_0)\mathcal{N}(0, \sigma_u^2).$$

Let $F_{\hat{\gamma}_j}(x) = p(\hat{\gamma}_j \leq x)$ be the cumulative distribution of $\hat{\gamma}_j$, we have

$$\begin{aligned} F_{\hat{\gamma}_j}(x) =& p(\hat{\gamma}_j \leq x) \\ =& p(\sqrt{N_1}(\hat{\gamma}_j - \tilde{\gamma}_j) \leq \sqrt{N_1}(x - \tilde{\gamma}_j)) \\ =& \mathbb{E}\left[p(\sqrt{N_1}(\hat{\gamma}_j - \tilde{\gamma}_j) \leq \sqrt{N_1}(x - \tilde{\gamma}_j)|\tilde{\gamma}_j)\right] \\ =& \mathbb{E}\left[F_{N_1,j}(\sqrt{N_1}(x - \tilde{\gamma}_j))\right] \\ =& \mathbb{E}\left[F_{N_1,j}((x - \tilde{\gamma}_j)/\hat{s}_{X,j})\right]. \end{aligned}$$

As shown above, we have obtained

$$\sup_x |F_{N_1,j}(x) - \Phi(x)| \leq \frac{C_0}{\sqrt{N_1}} C^* \to 0, \text{ as } N_1 \to \infty, \text{for any } j.$$

Therefore,

$$\begin{aligned} &\sup_x \left| F_{\hat{\gamma}_j}(x) - \mathbb{E}\left[\Phi((x - \tilde{\gamma}_j)/\hat{s}_{X,j})\right] \right| \\ =& \sup_x \left|\mathbb{E}\left[F_{N_1,j}((x - \tilde{\gamma}_j)/\hat{s}_{X,j})\right] - \mathbb{E}\left[\Phi((x - \tilde{\gamma}_j)/\hat{s}_{X,j})\right]\right| \\ \leq& \mathbb{E}\left[\sup_x |F_{N_1,j}((x - \tilde{\gamma}_j)/\hat{s}_{X,j}) - \Phi((x - \tilde{\gamma}_j)/\hat{s}_{X,j})|\right] \\ \leq& \frac{C_0}{\sqrt{N_1}} C^* \to 0, \text{ as } N_1 \to \infty, \text{for any } j. \end{aligned}$$

Consequently, we obtain that the approximated distribution of $\hat{\gamma}_j$, i.e. $F_{\hat{\gamma}_j}(x)$, uniformly converges in distribution to $\mathbb{E}\left[\Phi((x - \tilde{\gamma}_j)/\hat{s}_{X,j})\right]$ for any $j = 1, \ldots, M$.

Now, we derive the closed form of the approximated distribution $\mathbb{E}\left[\Phi((x-\tilde{\gamma}_j)/\hat{s}_{X,j})\right]$, which is given by

$$
\begin{aligned}
&\mathbb{E}\left[\Phi((x-\tilde{\gamma}_j)/\hat{s}_{X,j})\right] \\
&= \int_{-\infty}^{+\infty}\int_{-\infty}^{x} \frac{1}{\sqrt{2\pi \hat{s}_{X,j}^2}} \exp\left\{-\frac{(t-\tilde{\gamma}_j)^2}{2\hat{s}_{X,j}^2}\right\} dt \left(\frac{\pi_0}{\sqrt{2\pi(\sigma^2+\sigma_u^2)}} \exp\left\{-\frac{\tilde{\gamma}_j^2}{2(\sigma^2+\sigma_u^2)}\right\} + \frac{1-\pi_0}{\sqrt{2\pi\sigma_u^2}} \exp\left\{-\frac{\tilde{\gamma}_j^2}{2\sigma_u^2}\right\}\right) d\tilde{\gamma}_j \\
&= \int_{-\infty}^{x}\int_{-\infty}^{+\infty} \frac{\pi_0}{2\pi\sqrt{\hat{s}_{X,j}^2(\sigma^2+\sigma_u^2)}} \exp\left\{-\frac{(t-\tilde{\gamma}_j)^2}{2\hat{s}_{X,j}^2} - \frac{\tilde{\gamma}_j^2}{2(\sigma^2+\sigma_u^2)}\right\} + \frac{1-\pi_0}{2\pi\sqrt{\hat{s}_{X,j}^2\sigma_u^2}} \exp\left\{-\frac{(t-\tilde{\gamma}_j)^2}{2\hat{s}_{X,j}^2} + \frac{\tilde{\gamma}_j^2}{2\sigma_u^2}\right\} d\tilde{\gamma}_j dt.
\end{aligned}
\tag{S.28}
$$

The term in the first exponent within the integrals of above equation can be simplified as

$$
\begin{aligned}
\frac{(t-\tilde{\gamma}_j)^2}{\hat{s}_{X,j}^2} + \frac{\tilde{\gamma}_j^2}{(\sigma^2+\sigma_u^2)} &= \left(\frac{1}{\hat{s}_{X,j}^2} + \frac{1}{\sigma^2+\sigma_u^2}\right)\tilde{\gamma}_j^2 + \frac{t^2}{\hat{s}_{X,j}^2} - \frac{2\tilde{\gamma}_j t}{\hat{s}_{X,j}^2} \\
&= \left(\frac{1}{\hat{s}_{X,j}^2} + \frac{1}{\sigma^2+\sigma_u^2}\right)\left(\tilde{\gamma}_j - \frac{\frac{t}{\hat{s}_{X,j}^2}}{\frac{1}{\hat{s}_{X,j}^2} + \frac{1}{\sigma^2+\sigma_u^2}}\right)^2 + \frac{t^2}{\sigma^2+\sigma_u^2+\hat{s}_{X,j}^2}.
\end{aligned}
$$

As a result, we have

$$
\begin{aligned}
&\int_{-\infty}^{+\infty} \frac{\pi_0}{2\pi\sqrt{\hat{s}_{X,j}^2(\sigma^2+\sigma_u^2)}} \exp\left\{-\frac{(t-\tilde{\gamma}_j)^2}{2\hat{s}_{X,j}^2} - \frac{\tilde{\gamma}_j^2}{2(\sigma^2+\sigma_u^2)}\right\} d\tilde{\gamma}_j \\
=& \frac{\pi_0}{2\pi\sqrt{\hat{s}_{X,j}^2(\sigma^2+\sigma_u^2)}} \exp\left\{-\frac{t^2}{2(\sigma^2+\sigma_u^2+\hat{s}_{X,j}^2)}\right\} \int_{-\infty}^{+\infty} \exp\left\{-\left(\frac{1}{2\hat{s}_{X,j}^2} + \frac{1}{2(\sigma^2+\sigma_u^2)}\right)\left(\tilde{\gamma}_j - \frac{\frac{t}{\hat{s}_{X,j}^2}}{\frac{1}{\hat{s}_{X,j}^2} + \frac{1}{\sigma^2+\sigma_u^2}}\right)^2\right\} d\tilde{\gamma}_j \\
=& \frac{\pi_0}{2\pi\sqrt{\hat{s}_{X,j}^2(\sigma^2+\sigma_u^2)}} \exp\left\{-\frac{t^2}{2(\sigma^2+\sigma_u^2+\hat{s}_{X,j}^2)}\right\} \frac{\sqrt{2\pi}}{\sqrt{\frac{1}{\hat{s}_{X,j}^2} + \frac{1}{\sigma^2+\sigma_u^2}}} \\
=& \frac{\pi_0}{\sqrt{2\pi(\sigma^2+\sigma_u^2+\hat{s}_{X,j}^2)}} \exp\left\{-\frac{t^2}{2(\sigma^2+\sigma_u^2+\hat{s}_{X,j}^2)}\right\}.
\end{aligned}
$$

Similarly, we can compute

$$
\begin{aligned}
&\int_{-\infty}^{+\infty} \frac{(1-\pi_0)}{2\pi\sqrt{\hat{s}_{X,j}^2\sigma_u^2}} \exp\left\{-\frac{(t-\tilde{\gamma}_j)^2}{2\hat{s}_{X,j}^2} - \frac{\tilde{\gamma}_j^2}{2\sigma_u^2}\right\} d\tilde{\gamma}_j \\
=& \frac{1-\pi_0}{\sqrt{2\pi(\sigma_u^2+\hat{s}_{X,j}^2)}} \exp\left\{-\frac{t^2}{2(\sigma_u^2+\hat{s}_{X,j}^2)}\right\}.
\end{aligned}
$$

Consequently, we have

$$
\begin{aligned}
\mathbb{E}\left[\Phi((x-\tilde{\gamma}_j)/\hat{s}_{X,j})\right] = \int_{-\infty}^{x} & \frac{\pi_0}{\sqrt{2\pi(\sigma^2+\sigma_u^2+\hat{s}_{X,j}^2)}} \exp\left\{-\frac{t^2}{2(\sigma^2+\sigma_u^2+\hat{s}_{X,j}^2)}\right\} + \\
& \frac{1-\pi_0}{\sqrt{2\pi(\sigma_u^2+\hat{s}_{X,j}^2)}} \exp\left\{-\frac{t^2}{2(\sigma_u^2+\hat{s}_{X,j}^2)}\right\} dt,
\end{aligned}
$$

which gives

$$\hat{\gamma}_j \sim \pi_0 \mathcal{N}(0, \sigma^2 + \sigma_u^2 + \hat{s}_{X,j}^2) + (1-\pi_0)\mathcal{N}(0, \sigma_u^2 + \hat{s}_{X,j}^2), \text{ for } j = 1, \ldots M.$$

Next, we consider the case that SNPs are in LD. Note that we only use the summary statistics for a subset of $M_t$ independent IVs from the genome-wide SNPs for causal inference in MR-APSS analysis, i.e., $\{\hat{\gamma}_j, \hat{\Gamma}_j, \hat{s}_{X,j}, \hat{s}_{Y,j} \Big| |\hat{\gamma}_j / \hat{s}_{X,j}| > t\}_{j=1,\ldots,M_t}$, which are obtained by LD clumping ($r^2 < 0.001$, 1Mb). In the presence of LD, we have $\tilde{\gamma}_j = \sum_k r_{jk}(Z_j \gamma_k + u_k)$ (Eq. [S.11]). The marginal distribution of $\tilde{\gamma}_j$ for the $j$-th SNP is given by $\tilde{\gamma}_j \sim \pi_0 \mathcal{N}(0, \ell_j \sigma^2 + \ell_j \sigma_u^2) + (1-\pi_0)\mathcal{N}(0, \ell_j \sigma_u^2)$, where $\ell_j = \sum_{k=1}^{M} r_{jk}^2$ with $r_{jk}$ denotes the correlation between SNP $j$ and $k$. Given the uniform approximation of the conditional distribution $\hat{\gamma}_j | \tilde{\gamma}_j$, we can obtain uniform mixture Gaussian approximations for those $M_t$ independent SNPs:

$$\hat{\gamma}_j \sim \pi_0 \mathcal{N}(0, \ell_j \sigma^2 + \ell_j \sigma_u^2 + \hat{s}_{X,j}^2) + (1-\pi_0)\mathcal{N}(0, \ell_j \sigma_u^2 + \hat{s}_{X,j}^2), \text{ for } j = 1, \ldots, M_t.$$

Similarly, we can obtain the uniformity of the approximation of distribution of $\hat{\Gamma}_j$, i.e. $\hat{\Gamma}_j \sim \pi_0 \mathcal{N}(0, \ell_j \beta^2 \sigma^2 + \ell_j \tau^2 + \ell_j \sigma_v^2 + \hat{s}_{Y,j}^2) + (1-\pi_0)\mathcal{N}(0, \ell_j \sigma_v^2 + \hat{s}_{Y,j}^2)$, for $j = 1, \ldots, M_t$., and the uniformity of the approximation of the joint distribution of $(\hat{\gamma}_j, \hat{\Gamma}_j)$ given in Eq. [6] of the main text for all $j = 1, \ldots, M_t$.

#### 1.9 Discussion on the asymptotic normality of GWAS summary statistics after PC adjustment

PC adjustment is a standard approach to accounting for population stratification in GWAS data analysis [29]. To our best knowledge, the distribution of summary statistics after PC adjustment has not been rigorously established in the literature. In this section, we provide a justification on the asymptotic normality of GWAS summary statistics after PC adjustment. To avoid confusion, we will first discuss how PC adjustment is applied in the GWAS context and then provide our justification. For clarity, we use notations different from our main content.

Let us begin our discussion with the following linear model:

$$y = Z\alpha + \epsilon, \tag{S.29}$$

where $y$ is an $n \times 1$ vector of phenotypic values, $Z$ is an $n \times p$ standardized genotype matrix (whose column has mean zero and variance $1/p$), $\alpha$ is a $p \times 1$ vector of SNP effect sizes, and $\epsilon$ is an $n \times 1$ vector of independent errors that is distributed as $N(0, \sigma_\epsilon^2 I_n)$ with $I_n$ being the $n$-dimensional identity matrix. To generate the summary statistics, the following simple linear model is often used in GWAS:

$$y = Z_j a_j + \xi,$$

where only one SNP is considered at a time. The summary statistics are obtained as

$$\hat{a}_j = (Z_j^T Z_j)^{-1} Z_j^T y, \quad \text{s.e.}(\hat{a}_j) = \sqrt{\sigma_j^2 (Z_j^T Z_j)^{-1}}. \tag{S.30}$$

This approach is often referred to as marginal screening in the statistical community [14]. In the early days of GWAS, people have found that the summary statistics given by Eq. [S.30] are largely confounded by population stratification [29].

To account for population stratification, a few PCs are calculated from the standardized genotype matrix $Z$ and PC scores are included as covariates [29]. Let $Z = \hat{U}\hat{\Lambda}\hat{V}^T$ be the singular value decomposition (SVD), where we use the hat notation ($\hat{U}$, $\hat{\Lambda}$ and $\hat{V}$) to indicate that they are estimated from data. Specifically, the following linear model with PC adjustment is commonly used in GWAS:

$$y = \sum_{k=1}^{q} \hat{U}_k b_k + Z_j a_j + \xi,$$

where $b_k$ is the corresponding coefficients corresponding to $\hat{U}_k$. In real GWAS data analysis, $q = 10$ or 20 PCs are often used.

Noting that $q \ll n$, and here we only consider two PCs without loss of generality. We assume that the underlying true model relating phenotype $y$ with the genotypes and PC scores is given as

$$y = U_1\beta_1 + U_2\beta_2 + Z\alpha + \epsilon,$$

where $U_1$ and $U_2$ are the underlying true PC scores rather than their estimates ($\hat{U}_1$ and $\hat{U}_2$). Clearly, this model is a natural extension of model (S.29). To account for population stratification, the following model incorporating PC adjustment is used accordingly

$$y = \hat{U}_1 b_1 + \hat{U}_2 b_2 + Z_j a_j + \xi, \tag{S.31}$$

where two PCs ($\hat{U}_1$ and $\hat{U}_2$) and one SNP are included, and $b_1$, $b_2$, $a_j$ are the corresponding regression coefficients to be estimated. This approach is often referred to as conditional screening in the statistical community [3].

Based on model (S.31), $a_j$ can be estimated as

$$\hat{a}_j = e^T(\hat{W}^T\hat{W})^{-1}\hat{W}^T y,$$

where $e = [0, 0, 1]^T$ and $\hat{W} = [\hat{U}_1, \hat{U}_2, Z_j]$. Accordingly, let $W = [U_1, U_2, Z]$ collect the underlying true PC scores and the genotype matrix. With these notations, we have

$$\begin{aligned} \hat{a}_j &= e^T(\hat{W}^T\hat{W})^{-1}\hat{W}^T(U_1\beta_1 + U_2\beta_2 + Z\alpha + \epsilon) \\ &= e^T(\hat{W}^T\hat{W})^{-1}\hat{W}^T W \begin{bmatrix} \beta_1 \\ \beta_2 \\ \alpha \end{bmatrix} + e^T(\hat{W}^T\hat{W})^{-1}\hat{W}^T\epsilon. \end{aligned} \tag{S.32}$$

To illustrate the asymptotic normality of $\hat{a}_j$ based on Eq. [S.32], we assume that $Z$ satisfies a linear structure, namely, $Z = TX$, where $T \in \mathbb{R}^{n\times n}$ is deterministic and $X \in \mathbb{R}^{n\times p}$ is random with independent mean 0 and variance $\frac{1}{p}$ variables. Further, we assume that $\Sigma = TT^T$ admits a spiked structure, namely, $\Sigma = I + d_1U_1U_1^T + d_2U_2U_2^T$ with orthonormal $U_1$ and $U_2$. We further denote by $\hat{d}_i$ the $i$-th largest eigenvalue of $ZZ^T = TXX^TT^T$ and set $\hat{U}_i$ the corresponding $\ell^2$ normalized eigenvector. Let $Z = \hat{U}\hat{\Lambda}\hat{V}^T$ be the SVD of $Z$, where $\hat{\Lambda}$ collects the singular values $\hat{\lambda}_1 = \sqrt{\hat{d}_1}$, $\hat{\lambda}_2 = \sqrt{\hat{d}_2}$ and etc. In Random Matrix Theory, it is well known that when $d_1$ and $d_2$ are sufficiently large, $\hat{U}_i$ will favor the direction of $U_i$, and does not favor any other direction. More precisely, when $\frac{p}{n} = \tau$ and $n \to \infty$, for any given unit vector $w \in S^{n-1}$, we have $|\hat{U}_i^T w|^2 = \frac{d_i^2 - n/p}{d_i(d_i + n/p)}|U_i^T w|^2 + O_p(n^{-1/2})$, see Theorem 2.5 of [2] for instance. Under our model assumption, by a leave-one-out argument, one can easily show that $\hat{U}_i$ is almost independent of

$Z_j$. Further, $Z_j$ does not favor the direction $U_i$. By the above estimate of $|\hat{U}_i^T w|^2$, it is easy to show that $|\hat{U}_i^T Z_j|^2$ is negligible. When $\frac{p}{n} = \tau$ and $n \to \infty$, we can actually estimate all entries in $\hat{W}^T\hat{W}$ as

$$\begin{aligned}\hat{W}^T\hat{W} &= \begin{bmatrix} \hat{U}_1^T\hat{U}_1 & \hat{U}_1^T\hat{U}_2 & \hat{U}_1^T Z_j \\ \hat{U}_2^T\hat{U}_1 & \hat{U}_2^T\hat{U}_2 & \hat{U}_2^T Z_j \\ Z_j^T\hat{U}_1 & Z_j^T\hat{U}_2 & Z_j^T Z_j \end{bmatrix} = \begin{bmatrix} 1 & 0 & \hat{U}_1^T Z_j \\ 0 & 1 & \hat{U}_2^T Z_j \\ Z_j^T\hat{U}_1 & Z_j^T\hat{U}_2 & \frac{n}{p} \end{bmatrix} \\ &= \begin{pmatrix} 1 & 0 & 0 \\ 0 & 1 & 0 \\ 0 & 0 & n/p \end{pmatrix} + \text{negligible error.}\end{aligned} \tag{S.33}$$

By the estimate of $|\hat{U}_i^T w|^2$, we have

$$\begin{aligned}\hat{W}^T W &= \begin{pmatrix} \hat{U}_1 U_1 & \hat{U}_1^T U_2 & \hat{U}_1^T Z \\ \hat{U}_2 U_1 & \hat{U}_2^T U_2 & \hat{U}_2^T Z \\ Z_j^T U_1 & Z_j^T U_2 & Z_j^T Z \end{pmatrix} \\ &= \begin{pmatrix} \left(\frac{d_1^2 - n/p}{d_1(d_1+n/p)}\right)^{\frac{1}{2}} & 0 & \hat{\lambda}_1 \hat{V}^T \\ 0 & \left(\frac{d_2^2 - n/p}{d_2(d_2+n/p)}\right)^{\frac{1}{2}} & \hat{\lambda}_2 \hat{V}^T \\ 0 & 0 & Z_j^T Z \end{pmatrix} + \begin{pmatrix} O_p(n^{-1/2}) & 0 & 0 \\ 0 & O_p(n^{-1/2}) & 0 \\ 0 & 0 & 0 \end{pmatrix}.\end{aligned} \tag{S.34}$$

Now we look at the first term of Eq. [S.32]. Using Eq. [S.33] and Eq. [S.34], we have

$$e^T(\hat{W}^T\hat{W})^{-1}\hat{W}^T W \begin{bmatrix} \beta_1 \\ \beta_2 \\ \alpha \end{bmatrix} \to \frac{p}{n} Z_j^T Z\alpha = \sum_{l=1}^{p} Z_j^T Z_l \alpha_l.$$

To see the normality of $\sum_{l=1}^p Z_j^T Z_l \alpha_l$, we can condition on $Z_j$, and then $Z_j^T Z_l$ is a linear combination of $Z_\ell$ entries. By CLT, $Z_j^T Z_l | Z_j \to \mathcal{N}(0, \frac{1}{p}\|Z_j\|^2)$ and further we have $Z_j^T Z_l \to \mathcal{N}(0, \frac{n}{p^2})$ by the concentration of $\|Z_j\|^2$. Further, $\sum_{l=1}^p Z_j^T Z_l \alpha_l | Z_j$ is a linear combination of asymptotically normal variables. By a further application of CLT, we see that $\sum_{l=1}^p Z_j^T Z_l \alpha_l | Z_j$ is asymptotically normal, and the limiting normal distribution does not depend on $Z_j$. Hence, $\sum_{l=1}^p Z_j^T Z_l \alpha_l$ itself is asymptotically normal $\mathcal{N}(0, \frac{n}{p^2}\|\alpha\|_2^2)$. Therefore, we show that the first term in Eq. [S.32] indeed is asymptotically normal.

Next we check the second term of of Eq. [S.32]. Because $\epsilon$ is a vector of independent errors with a normal distribution, we have

$$e^T(\hat{W}^T\hat{W})^{-1}\hat{W}^T\epsilon \to \mathcal{N}(0, e^T M_1^{-1} e\, \sigma_\epsilon^2) = \mathcal{N}(0, \frac{p}{n}\sigma_\epsilon^2),$$

where $M_1 = \begin{pmatrix} 1 & 0 & 0 \\ 0 & 1 & 0 \\ 0 & 0 & n/p \end{pmatrix}$ is given in Eq. [S.33]. From the above discussion, it is easy to see that the first term and the second term in the RHS of (S.32) are asymptotically independent, due to (S.33). Hence, we can conclude that $\hat{a}_j$ given in Eq. [S.32] is indeed asymptotically normal.

The real data $Z$ may not satisfy the assumption of our toy model $Z = TX$. Nevertheless, the above argument can be potentially extended to a more general model such as the separable model $Z = TXS$ where both column- and row-dependence are allowed (see [13] for instance). The justification of the asymptotic normality for $\hat{a}_j$ without any model assumption is out of reach for this moment. It will be certainly an interesting direction for theoretical study in the future.

Finally, according to our experience, the publicly available summary statistics have been generated with PC adjustment. However, it does not mean that sample structure is no longer an issue after PC adjustment. As demonstrated by the LDSC method [7] and several other recent works (e.g., sample structure driven by socioeconomic status [1] or geographic structure [17]), confounding bias still remains as a severe issue for downstream analysis of using GWAS summary statistics. This fact motivates us to develop a statistical method to simultaneously correct pleiotropy and sample structure in MR analysis.

### 2 Related methods

#### 2.1 Background

*Classical assumptions on valid IVs.* Let $X$ be the exposure and $Y$ be the outcome. As most MR methods apply LD clumping to make SNPs nearly independent, here we assume that there are $p$ independent SNPs represented by mutually independent random variables $G_1, G_2, ..., G_p$. Then we consider the following individual-level model:

$$X = f(G_1, ..., G_p, U, E_X),$$

$$Y = g(X, G_1, ..., G_p, U, E_Y),$$

$$U = h(G_1, ..., G_p, E_U),$$

where $U$ is the unmeasured confounder and $E_X, E_Y, E_U$ are mutually independent random noises which satisfy $(E_X, E_Y) \perp\!\!\!\perp (G_1, ..., G_p, U)$, $E_U \perp\!\!\!\perp (G_1, ..., G_p)$.

A variable $G_j$ is called a valid IV if it satisfies the following three assumptions:

**(A-I)**. Relevance: $G_j \not\perp\!\!\!\perp X|U$;

**(A-II)**. Effective random assignment: $G_j \perp\!\!\!\perp U$;

**(A-III)**. Exclusion restriction: $G_j \perp\!\!\!\perp Y|X, U$.

*Linear model for MR.* Many MR methods assume the **linearity** of $f, g, h$, i.e, functions $f, g, h$ are linear in their arguments. Under this assumption, the linear model for MR is written as:

$$X = \sum_{j=1}^{p} \gamma_j^* G_j + \eta_X U + E_X, \tag{S.35}$$

$$Y = \beta X + \sum_{j=1}^{p} \alpha_j G_j + \eta_Y U + E_Y, \tag{S.36}$$

$$U = \sum_{j=1}^{p} \psi_j G_j + E_U, \tag{S.37}$$

where $\beta$ is the causal effect of exposure $X$ on outcome $Y$. Based on the linear model for MR, the IV $G_j$ is valid if

**(A-I)**. Relevance: $\gamma_j^* \neq 0$;

**(A-II)**. Effective random assignment: $\psi_j = 0$;

**(A-III)**. Exclusion restriction: $\alpha_j = 0$.

*Assumptions on IVs recently proposed in the literature.* In the literature of MR, several assumptions were recently introduced to relax assumptions **(A-II)** and **(A-III)**:

**InSIDE** under **(A-II)**. The InSIDE assumption relaxes the exclusion restriction assumption: The direct effect $\alpha_j$ of the IV $G_j$ on the outcome $Y$ can be nonzero (violation of **(A-III)**), but the Instrument Strength must be Independent of the Direct Effect, i.e., $\gamma_j^* \perp\!\!\!\perp \alpha_j$.

**Majority valid**. The majority valid assumption allows for possible violation of **(A-II)** and **(A-III)**, but it requires that more than 50% of the IVs being used are valid IVs.

**Plurality valid**. The plurality valid assumption also allows possible violation of **(A-II)** and **(A-III)**, and it is weaker than the majority valid assumption. It requires that out of all groups of IVs having the same asymptotic ratio estimates of the causal effect, the largest group is the group of valid IVs. The difference of the majority valid assumption and the plurality valid assumption can be seen from the following example. Suppose there are three groups of IVs, with group proportions 30%, 30%, and 40%. The first group does not satisfy **(A-II)**, the second group does not satisfy **(A-III)**, and the third group satisfies **(A-I)**, **(A-II)**, and **(A-III)**. In this case, the plurality valid assumption holds while the majority valid assumption does not hold.

**NOME**. The assumption NOME refers to the NO Measurement Error assumption. It assumes that the variances of IV-exposure association estimates are negligible in summary-level MR methods.

#### 2.2 Review of summary-level MR methods

All of the compared summary-level MR methods, including dIVW [38] and RAPS [41] from the statistical community, assume the linearity of functions $f, g, h$ to derive the model for GWAS summary-level data. The linear model for MR in Eqs. [S.35], [S.36], [S.37] is equivalent to the following model:

$$X = \sum_{j=1}^{p} (\gamma_j^* + \eta_X \psi_j) G_j + E'_X,\ E'_X = \eta_X E_U + E_X,$$

$$Y = \sum_{j=1}^{p} [\beta(\gamma_j^* + \eta_X \psi_j) + \alpha_j + \eta_Y \psi_j] G_j + E'_Y,\ E'_Y = \beta(\eta_X E_U + E_X) + (\eta_Y E_U + E_Y).$$

For the sake of clarity, we do not consider the influence of linkage disequilibrium in this model, although we have carefully addressed the issue in our proposed MR-APSS.

Now we can define

$$\gamma_j = \gamma_j^* + \eta_X \psi_j, \tag{S.38}$$

$$\Gamma_j = \beta(\gamma_j^* + \eta_X \psi_j) + \alpha_j + \eta_Y \psi_j = \beta\gamma_j + \alpha_j + \eta_Y \psi_j, \tag{S.39}$$

where $\gamma_j$ and $\Gamma_j$ are the underlying true marginal effect sizes of $G_j$ on exposure $X$ and outcome $Y$. The estimated effect sizes and their standard errors, denoted as $(\hat{\gamma}_j, \hat{s}_{X,j})$ and $(\hat{\Gamma}_j, \hat{s}_{Y,j})$, are available in the released GWAS summary statistics. From Eqs. [S.38], [S.39], if $G_j$ is a valid

IV, i.e., **(A-I)** ensures $\gamma_j^* \neq 0$, **(A-II)** ensures $\psi_j = 0$, **(A-III)** ensures $\alpha_j = 0$, then

$$\gamma_j = \gamma_j^*,\ \Gamma_j = \beta\gamma_j^* = \beta\gamma_j. \tag{S.40}$$

Therefore, summary-level MR methods essentially use ratio estimates to perform causal inference. However, the assumptions on IVs, especially **(A-II)** and **(A-III)**, are often violated, and different summary-level MR methods are proposed to address the challenge. To summarize the efforts made in the development of summary-level MR methods, we roughly divide the related methods into three groups.

*Group 1: methods which require that all IVs are valid.*

IVW is a standard approach in two-sample summary-level MR studies under the strict condition that all IVs are valid, i.e., all IVs satisfy the relationship $\Gamma_j = \beta\gamma_j$. IVW forms a meta analysis of single causal estimates $\hat{\beta}_j = \hat{\Gamma}_j/\hat{\gamma}_j$. By further requiring NOME, IVW reports the causal effect estimate by taking the inverse-variance weighted ($w_j = \text{Var}(\hat{\beta}_j)^{-1} = (\hat{s}_{Y,j}^2/\hat{\gamma}_j^2)^{-1}$) mean of $\hat{\beta}_j$, leading to a simple estimator as $\frac{\sum_{j=1}^p \hat{\beta}_j w_j}{\sum_{j=1}^p w_j}$.

*Group 2: methods which addresses the possible violation of **(A-III)**.*

The MR methods in Group 2 are developed to relax **(A-III)**. These methods, including Egger [5], RAPS [41], and dIVW [38], still require assumptions **(A-I)** and **(A-II)**, but relax **(A-III)** by allowing for the presence of direct effect of IVs on the outcome ($\alpha_j \neq 0$). In this case, combining Eqs. [S.38], [S.39], and the condition $\psi_j = 0$ ensured by **(A-II)**, MR methods in Group 2 rely on the following relationship:

$$\gamma_j = \gamma_j^*, \quad \Gamma_j = \beta\gamma_j^* + \alpha_j = \beta\gamma_j + \alpha_j. \tag{S.41}$$

To account for the existence of non-zero $\alpha_j$, methods in this group further require that direct effects $\alpha_j$ of IVs on the outcome are independent of instrument strength $\gamma_j$ between IVs and the exposure, which is referred to as the InSIDE condition. Under this condition, direct effects $\alpha_j$'s, which are also referred to as horizontal pleiotropic effects in the literature of MR, can be viewed as independent random noises. Hence, Eq. [S.41] adopted by MR methods in this group can be viewed as the noisy version of Eq. [S.40] adopted by IVW in Group 1. Methods in this group make different assumptions on the distribution of $\alpha_j$ and use different strategies to construct estimators for the causal effect. Egger assumes that all IVs are affected by directional pleiotropy, i.e., $\mathbb{E}(\alpha_j) = \mu$, and it extends IVW estimator by further introducing an intercept term to capture the possible existence of non-zero $\mu$. Despite this improvement over IVW, Egger provides conservative results for causal inference, as known in literature. Different from MR-PRESSO and Egger, two MR methods, RAPS and dIVW, specify a distribution for $\alpha_j$. RAPS and dIVW assume that $\alpha_j$'s are independent and identically distributed random variables that follow normal distribution with mean zero. Additionally, they carefully account for the bias induced by the usage of many weak IVs by making use of estimation errors. These two methods adopt different strategies to estimate the causal effect under the similar assumptions. To have a robust estimate of the causal effect when $\alpha_j$ are deviated from the assumed distribution, RAPS modifies the profile likelihood. The recently proposed method dIVW extends IVW by modifying the weights. The resulting dIVW estimator is shown to be consistent and asymptotically normal in the presence of balanced pleiotropy ($\alpha_j \sim \mathcal{N}(0, \tau^2)$). Although much efforts have been made by developing MR methods in Group 2, the InSIDE condition may be violated due to correlated pleiotropy. The usage of MR methods in Group 2 may be limited in the presence of correlated pleiotropy.

*Group 3: methods which address the possible violation of assumptions **(A-II)** and **(A-III)**.*

MR Methods in Group 3 improve over MR methods in Groups 1 and 2 by allowing for IVs violating assumptions **(A-II)** and **(A-III)**. Eqs. [S.38] and [S.39] summarize the relationship of effect sizes from invalid IVs, where $\alpha_j, \psi_j, \eta_X$, and $\eta_Y$ are possibly non-zero. In this group, we summarize six recent works, including weighted-median [6], weighted-mode [16], MRMix [30], cML-MA [35], CAUSE[26] and our proposed MR-APSS. These MR methods require extra but weaker assumptions to relax **(A-II)** and **(A-III)**. According to their assumptions, we divide these six methods into three subgroups.

- **Subgroup 1**: MR methods in subgroup 1 require the majority valid assumption. Among all the IVs used for causal inference, more than 50% of them satisfy assumptions **(A-I)**, **(A-II)**, **(A-III)**, and thus the relationship $\Gamma_j = \beta\gamma_j$ holds for the majority of IVs. The method weighted-median is an MR approach of this kind. It combines the IVW estimator and the simple median estimator to provide a weighted median estimator which is consistent under the condition that at least 50% of the weight comes from valid IVs. Although progress has been made by methods in subgroup 1 to provide robust estimators, the validity of their required assumption is still hard to verify.

- **Subgroup 2**: MR methods in subgroup 2, including weighted-mode, MRMix, cML-MA, are developed based on plurality valid assumption. Under this assumption, IVs satisfying **(A-I)**, **(A-II)**, and **(A-III)** can form the largest group among all groups of IVs having the same asymptotic ratio estimates of the causal effect. Clearly, this assumption is weaker than the majority valid assumption. With the plurality valid assumption, MR methods in this subgroup can extend the simple mode estimator to perform causal inference. The weighted-mode method modifies the simple mode estimator by using a new weighting mechanism. MRMix is a model-based MR approach that leverages normal-mixture model to capture the mode of valid IVs. The cML-MA method uses the constrained likelihood approach to provide causal effect estimates where the $L_0$ penalty is introduced to select valid IVs among all IVs. We evaluated these methods using simulation and real data analysis. We found that the weighted-mode method is often very conservative. We also find that MRMix and cML-MA tend to provide inflated type I errors in the presence of population stratification.

- **Subgroup 3**: MR methods in subgroup 3, including CAUSE and our proposed MR-APSS, allow all IVs to be possibly invalid. The CAUSE model distinguishes two types of pleiotropy: correlated pleiotropy and uncorrelated pleiotropy. To account for the two types of pleiotropy, CAUSE uses the following model to relate effect size on exposure ($\gamma_j$) and outcome ($\Gamma_j$)

  $$\Gamma_j = \beta\gamma_j + Z_j\eta\gamma_j + \alpha_j,$$

  where $\beta$ is the causal effect of interest, $\eta$ is the correlated pleiotropic effect, $\alpha_j$ is uncorrelated pleiotropy, and $Z_j \in \{0, 1\}$ indicates whether correlated pleiotropy exists. To make the above model identifiable, CAUSE assumes that the proportion of IVs affected by correlated pleiotropy should be less than 50% and uncorrelated pleiotropy

$\alpha_j \sim N(0, \tau^2)$. In this sense, CAUSE tries to combine the majority valid assumption and balanced pleiotropy. Despite this conceptual advance, a closer examination of CAUSE (details presented in our supplementary note) shows that CAUSE tends to treat the causal effect as correlated pleiotropy during the model fitting, leading to very conservative performance for detecting causal effects.

In contrast to CAUSE, MR-APSS relaxes **(A-II)** and **(A-III)** by imposing the LDSC assumptions in its background model and the InSIDE condition in the foreground model. By integrating the background model and the foreground model using a mixture model, MR-APSS not only accounts for two types of pleiotropy but also accounts for **sample structure** (population stratification, cryptic relatedness, and sample overlap). To the best of our knowledge, however, sample structure is largely ignored in the literature of summary-level MR methods. Furthermore, MR-APSS allows incorporation of IVs with moderate effects to improve statistical power. To do so, MR-APSS accounts for **selection bias** (which is also referred to as winner's curse in the GWAS context) to avoid bias due to the IV selection. Among all compared summary-level MR methods, MR-APSS and recently developed dIVW are the only two methods that correct for selection bias. This correction is critical to improve power and avoid inflated type I errors.

#### 2.3 Review of individual-level MR methods

Different from summary-level MR methods which take GWAS summary statistics and a reference genome as inputs, individual-level MR methods can access individual-level samples, including genotypes $G$, phenotypes of exposure trait $X$ and outcome trait $Y$, and covariates $Z$. Here we mainly focus on individual-level MR methods which aim to relax assumptions **(A-II)** and **(A-III)**. As a supplement, we also summarize whether the compared individual-level MR methods assume the linearity for MR model or not in Table S1 to have a better comparison with summary-level MR approaches which require linearity for MR model. To summarize the progress made by individual-level MR studies, we roughly divide the related methods into three groups according to the key assumptions that they required.

*Group 1: methods which require all IVs to be valid.*

Two-stage least squares (TSLS) and Limited information maximum likelihood (LIML)[11] are two methods for performing causal inference based on the strict assumption that all IVs are valid. TSLS relies on the linear MR model and it is a two-stage sequential regression method. In the first stage, TSLS regresses exposure $X$ on IVs $G$ to obtain fitted values of the exposure as $\hat{X}|G$. In the second stage, it then regresses outcome $Y$ on the fitted values of the exposure $\hat{X}|G$. The obtained coefficient in the second stages serves as the causal effect estimate for TSLS. LIML extends TSLS by combining the two-stage regressions into a unified likelihood-based method. LIML often improves over TSLS as it avoids overfitting and reduces the impact of many weak instruments bias compared to TSLS.

*Group 2: methods which address the possible violation of **(A-III)**.*

The MR approach MBTSLS [19] is a representative method which belongs to group 2. MBTSLS allows a direct effect $\alpha_j$ on outcome $Y$ in the MR model by imposing the InSIDE condition, i.e., direct effects $\alpha_j$ of IVs on the outcome are independent of instrument strength $\gamma_j^*$ between IVs and the exposure. Although methods in group 2 improve over methods in group 1, they are not satisfactory for performing causal inference as the InSIDE assumption and **(A-II)** may be violated.

*Group 3: methods which addresses the possible violation of* ***(A-II)*** *and* ***(A-III)****.*

Representative methods in this groups include sisVIVE [18], Adaptive Lasso [34], TSHT [15], GENIUS [31], GENIUS-MAWII [37], and MR-MiSTERI [21]. Here we roughly divide these methods into three subgroups according to the key assumptions that they required.

- **Subgroup 1**: MR methods in subgroup 1 require the assumption that more than 50% of IVs being used are valid IVs satisfying **(A-I)**, **(A-II)**, and **(A-III)**, which is known as the majority valid assumption. Two methods, sisVIVE and Adaptive Lasso, are developed under the majority valid assumption to relax **(A-II)** and **(A-III)**. The method sisVIVE is an $L_1$ penalized regression approach based on the majority valid assumption, where the $L_1$ penalty is introduced to account for the sparsity of pleiotropic effects of IVs. Compared with sisVIVE, Adaptive Lasso can obtain a consistent estimator for causal effects under weaker conditions.

- **Subgroup 2**: MR methods in subgroup 2 are based on the plurality valid assumption. TSHT is an MR method of this kind. TSHT is a two-stage hard thresholding approach. In the first stage, it identifies the set of IVs that satisfy **(A-I)** by thresholding the strength of associations between IVs and the exposure. In the second stage, TSHT constructs multiple estimators for pleiotropic effects as $(\widehat{\Gamma_j - \beta\gamma_j})^{[k]} = \hat{\Gamma}_j - \frac{\hat{\Gamma}_k}{\hat{\gamma}_k}\hat{\gamma}_j$, where the $k$-th estimator is built upon the ratio estimate obtained using the $k$-th IV. It then performs thresholding on these estimates of pleiotropic effects with voting to select valid IVs. The resulting causal effect estimate is proved to be consistent under the plurality valid condition. It is worthwhile mentioning that TSHT uses both individual-level data and summary statistics in its two-stage thresholding procedure. So it belongs to individual-level MR methods.

- **Subgroup 3**: MR methods in subgroup 3 allow all IVs to be possibly invalid. Pervasive pleiotropy can lead to the violation of majority valid assumption and plurality valid assumption. Two individual-level MR methods, GENIUS and GENIUS-MAWII, are thus developed to provide robust estimate for causal effect even all IVs are invalid. Unlike existing methods, GENIUS leverages heteroscedasticity of the exposure for a robust estimator of causal effect. To see the key idea of GENIUS, we consider a simple exposure-outcome model:

$$X = \gamma(G) + U, \quad Y = \beta X + \alpha(G) + U,$$

where $G \perp\!\!\!\perp U$, $\alpha(G)$ represents the influence of pleiotropy and thus **(A-III)** is violated. Based on this simple model, it is easy to see that

$$\frac{E\{[G-\mathbb{E}(G)]Y\}}{E\{[G-\mathbb{E}(G)]X\}} = \beta + \frac{E\{[G-\mathbb{E}(G)]\alpha(G)\}}{E\{[G-\mathbb{E}(G)]X\}} + \underbrace{\frac{E\{[G-\mathbb{E}(G)]U\}}{E\{[G-\mathbb{E}(G)]X\}}}_{=0 \text{ because } G\perp\!\!\!\perp U}.$$

Therefore, using ratio $\frac{E\{[G-\mathbb{E}(G)]Y\}}{E\{[G-\mathbb{E}(G)]X\}}$ to obtain the causal effect $\beta$ only works when $E\{[G-\mathbb{E}(G)]\alpha(G)\} = 0$. To eliminate the influence of $\alpha(G)$, GENIUS uses the exposure residual term $X - \mathbb{E}(X|G)$ because $\mathbb{E}\{[X - \mathbb{E}(X|G)]\alpha(G)\} = 0$.

Instead of working with ratio $\frac{E\{[G-\mathbb{E}(G)]Y\}}{E\{[G-\mathbb{E}(G)]X\}}$, GENIUS considers the following relationship,

$$\begin{aligned}\frac{\mathbb{E}\{[G-\mathbb{E}(G)][X-\mathbb{E}(X|G)]Y\}}{\mathbb{E}\{[G-\mathbb{E}(G)][X-\mathbb{E}(X|G)]X\}}=\beta+&\frac{\mathbb{E}\{[G-\mathbb{E}(G)][X-\mathbb{E}(X|G)]\alpha(G)\}}{\mathbb{E}\{[G-\mathbb{E}(G)][X-\mathbb{E}[X|G]]X\}}\\+&\frac{\mathbb{E}\{[G-\mathbb{E}(G)][X-\mathbb{E}(X|G)]U\}}{\mathbb{E}\{[G-\mathbb{E}(G)][X-\mathbb{E}(X|G)]X\}},\end{aligned} \tag{S.42}$$

where the second term on the right hand side is zero because $\mathbb{E}\{[G-\mathbb{E}(G)][X-\mathbb{E}(X|G)]\alpha(G)\}=0$ and the third term is zero by assumption $G\perp\!\!\!\perp U$. Therefore, the causal effect can be obtained by $\frac{\mathbb{E}\{[G-\mathbb{E}(G)][X-\mathbb{E}(X|G)]Y\}}{\mathbb{E}\{[G-\mathbb{E}(G)][X-\mathbb{E}(X|G)]X\}}$, where $\mathbb{E}\{[G-\mathbb{E}(G)][X-\mathbb{E}(X|G)]X\}=\mathrm{Cov}[G,\mathrm{var}(X|G)]\neq 0$ is the key assumption which requires heteroscedasticity of the exposure. In the GENIUS paper, the authors consider a more general model than what we consider here,

$$\mathbb{E}(X|G,U)=\gamma(G,U)+\xi_x(U),\quad \mathbb{E}(Y|X,G,U)=\beta X+\alpha(G,U)+\xi_y(U),$$

where $\gamma(G,U)$, $\alpha(G,U)$, and $\xi_y(U)$ are some unknown functions satisfying $\gamma(0,U)=\alpha(0,U)=0$ and the orthogonality conditions:

$$\mathrm{Cov}(\alpha(G,U),\gamma(G,U)|G)=\mathrm{Cov}(\alpha(G,U),\xi_x(U)|G)=\mathrm{Cov}(\xi_y(U),\gamma(G,U)|G)=0.$$

The assumption $G\perp\!\!\!\perp U$ can be further relaxed by a weaker second-order condition $\mathrm{Cov}(\xi_x(U),\xi_y(U)|G)=\rho$, where $\rho$ is a constant. With these key assumptions, GENIUS can relax **(A-II)** and **(A-III)**. GENIUS-MAWII further extends GENIUS by allowing for incorporation of many weak IVs. GENIUS-MAWII proposes a continuous updating estimator of the causal effect and establishes its consistency and asymptotic normality. Very recently, a new MR method MR-MiSTERI [21] has been proposed by requiring heteroscedasticity of the outcome.

#### 2.4 Discussion on CAUSE

To gain insight into the assumptions and the properties of CAUSE, we first provide a review on CAUSE and then discuss its model. CAUSE is proposed to distinguish two types of pleiotropy. The first type of pleiotropy is uncorrelated pleiotropy where the direct effects of SNPs on the outcome are not correlated with the SNP effects on the exposure. The second type of pleiotropy is correlated pleiotropy. It occurs when the SNPs affect both the exposure and the outcome through shared pathways. The InSIDE condition no longer holds when correlated pleiotropy arises. To account for uncorrelated pleiotropy and correlated pleiotropy simultaneously, CAUSE models the relationship of SNP effects between the exposure and the outcome as follows:

$$\Gamma_i=\beta\gamma_i+Z_i\eta\gamma_i+\theta_i,$$

where $\beta$ is the causal effect of interest, $\eta$ is the correlated pleiotropic effect, $\theta_i$ is an uncorrelated pleiotropic effect, and $Z_i$ is indicators for a valid IV ($Z_i=0$) or not ($Z_i=1$). To successfully identify causal effect, CAUSE assumes that the proportion of IVs affected by correlated pleiotropy should be less than 50%, $q=\Pr(Z_i=1)<0.5$ (this is very similar to the majority valid assumption). With the prior on true SNP effects on the exposure and the outcome $\gamma_i\sim$

$\mathcal{N}(0,\sigma^2), \Gamma_i \sim \mathcal{N}(0,\tau^2)$ and the variance of estimation errors $\mathbf{S}_i(\rho) = \begin{pmatrix} s_{X,i}^2 & \rho s_{X,i}s_{Y,i} \\ \rho s_{X,i}s_{Y,i} & s_{Y,i}^2 \end{pmatrix}$, the CAUSE model is written as

$$p(\hat{\gamma}_i, \hat{\Gamma}_i|\beta, \eta, \sigma^2, \tau^2, \mathbf{S}_i) = q\mathcal{N}\left(\begin{pmatrix} \hat{\gamma}_i \\ \hat{\Gamma}_i \end{pmatrix} \middle| \mathbf{0}, \begin{pmatrix} \sigma^2 & (\beta+\eta)\sigma^2 \\ (\beta+\eta)\sigma^2 & (\beta+\eta)^2\sigma^2+\tau^2 \end{pmatrix} + \mathbf{S}_i(\rho)\right) + (1-q)\mathcal{N}\left(\begin{pmatrix} \hat{\gamma}_i \\ \hat{\Gamma}_i \end{pmatrix} \middle| \mathbf{0}, \begin{pmatrix} \sigma^2 & \beta\sigma^2 \\ \beta\sigma^2 & \beta^2\sigma^2+\tau^2 \end{pmatrix} + \mathbf{S}_i(\rho)\right).$$

On the right hand side of the above equation, the first term is related to IVs affected by correlated pleiotropic effects and the second term characterizes IVs that are only affected by uncorrelated pleiotropy.

To perform causal inference with this model, CAUSE proposes the following workflow:

- (step 1) Fix $\beta = 0$, $\eta = 0$, and estimate $\sigma^2, \tau^2$ (parameters in priors) and $\rho$ (impact of sample overlapping) using genome-wide summary statistics.
- (step 2) Fit the null model: fix $\beta = 0$, and estimate $\eta, q$ using selected IVs ($p$-value $\leq$ IV threshold).
- (step 3) Fit the CAUSE model using selected IVs.
- (step 4) Compute the expected log pointwise posterior density (ELPD) test statistics by comparing the results between the fitted null model and the fitted CAUSE model.

The problem occurs in step 2 of CAUSE. When fixing $\beta = 0$, the model becomes

$$p(\hat{\gamma}_i, \hat{\Gamma}_i|\beta = 0, \eta, \sigma^2, \tau^2, \mathbf{S}_i) = q\mathcal{N}\left(\begin{pmatrix} \hat{\gamma}_i \\ \hat{\Gamma}_i \end{pmatrix} \middle| \mathbf{0}, \begin{pmatrix} \sigma^2 & \eta\sigma^2 \\ \eta\sigma^2 & \eta^2\sigma^2+\tau^2 \end{pmatrix} + \mathbf{S}_i(\rho)\right) + (1-q)\mathcal{N}\left(\begin{pmatrix} \hat{\gamma}_i \\ \hat{\Gamma}_i \end{pmatrix} \middle| \mathbf{0}, \begin{pmatrix} \sigma^2 & 0 \\ 0 & \tau^2 \end{pmatrix} + \mathbf{S}_i(\rho)\right).$$

Therefore, the underlying causal effect can be absorbed into the estimated $\eta$ (in the first term of the right hand side). As a result, the causal estimate given by CAUSE is biased to the null ($\beta = 0$) even in simulations where data generation matches the CAUSE model. The $p$-value obtained through computing ELPD by comparing the null model and the CAUSE model will be deflated, leading to lower statistical power.

#### 2.5 Measures of the IV strength in literature

For clarity, we use a model of summary-level MR methods in groups 1 and 2 as an example to illustrate the notions of IV strength in the literature of summary-level MR methods. These methods rely on the following linear model for MR:

$$X = \sum_j \gamma_j G_j + \eta_X U + E_X,\ Y = \beta X + \sum_j \alpha_j G_j + \eta_Y U + E_Y,$$

where subscript $j$ denotes the $j$-th SNP, $X$ is a phenotype vector of the exposure, $Y$ is a phenotype vector of the outcome, $G_j$ is a genotype vector of the $j$-th SNP, $X, Y, G_j$ are standardized to have zero mean and variance one, $U$ is the unmeasured confounder, $E_X, E_Y$ are independent random noises. Here we do not consider the influence of linkage disequilibrium (LD) in this model to avoid unnecessary confusion. By regressing $X$ and $Y$ on $G_j$, we can obtain the estimated effect sizes of the $j$-th SNP and their standard errors $(\hat{\gamma}_j, \hat{s}_{X,j}), (\hat{\Gamma}_j, \hat{s}_{Y,j})$, respectively. In this setting, we typically obtain $\hat{s}^2_{X,j} \approx 1/N_1, \hat{s}^2_{X,j} \approx 1/N_2$ because the genotypes and phenotypes are assumed to be standardized, where $N_1$ and $N_2$ are the GWAS sample sizes of the exposure and the outcome, respectively. We denote the corresponding true effect sizes as $\gamma_j$ and $\Gamma_j$. Then, MR methods in groups 1 and 2 (Main text, Table 1) are closely related to the fitting of the following errors-in-variables regression of $\hat{\Gamma}_j$ on $\hat{\gamma}_j$:

$$\begin{aligned} &\begin{pmatrix} \hat{\gamma}_j \\ \hat{\Gamma}_j \end{pmatrix} | \gamma_j, \alpha_j \sim \mathcal{N}\left( \begin{pmatrix} \gamma_j \\ \beta\gamma_j + \alpha_j \end{pmatrix}, \begin{pmatrix} \hat{s}^2_{X,j} & 0 \\ 0 & \hat{s}^2_{Y,j} \end{pmatrix} \right), \forall j = 1, ..., M_t, \\ &\alpha_j = 0, \text{ or } \alpha_j \text{ follows a specified distribution,} \end{aligned} \tag{S.43}$$

where subscript $t$ corresponds to the IV selection criterion $(|\hat{\gamma}_j/\hat{s}_{X,j}| \geq t)$, $M_t$ represents the number of selected IVs for the given threshold $t$, and $\{\gamma_j\}_{j=1,...,M_t}$ are regarded as nuisance parameters. The $\{\hat{\gamma}_j\}_{j=1,...,M_t}$ serve as predictors in this errors-in-variables regression, and the $\{\gamma_j\}_{j=1,...,M_t}$ are the underlying true effect sizes with strengths $|\gamma_j|, j = 1, ..., M_t$. In the literature of MR, the collective IV strength [41] is defined as

$$\text{Collective IV strength} := \sum_{j=1}^{M_t} \gamma_j^2 = \|\boldsymbol{\gamma}\|_2^2.$$

Another notion related to the IV strength is the average IV strength [38, 41]:

$$\text{Average IV strength} := \frac{1}{M_t} \sum_{j=1}^{M_t} \frac{\gamma_j^2}{\hat{s}^2_{X,j}}. \tag{S.44}$$

Next, we discuss our definition of the IV strength. Recall that the MR-APSS model is given as:

$$\begin{pmatrix} \hat{\gamma}_j \\ \hat{\Gamma}_j \end{pmatrix} = Z_j \underbrace{\underset{\text{Uncorrelated pleiotropy}}{\begin{pmatrix} \gamma_j \\ \beta\gamma_j + \alpha_j \end{pmatrix}}}_{\text{Foreground}} + \underbrace{\underset{\substack{\text{Polygenicity} \\ \text{Correlated pleiotropy}}}{\begin{pmatrix} u_j \\ v_j \end{pmatrix}} + \underset{\substack{\text{Sample structure} \\ \text{(Population stratification,} \\ \text{cryptic relatedness,} \\ \text{sample overlap, etc.)}}}{\begin{pmatrix} e_j \\ \xi_j \end{pmatrix}}}_{\text{Background}}, \quad j = 1, \ldots M_t, \tag{S.45}$$

where the background model is designed to account for polygenicity, correlated pleiotropy, and sample structure, and the foreground model aims to identify informative instruments and account for uncorrelated pleiotropy to perform causal inference. By assuming the covariance matrices of $(u_j, v_j)^T$ and $(e_j, \xi_j)^T$, the MR-APSS model can be written as:

$$\begin{pmatrix} \hat{\gamma}_j \\ \hat{\Gamma}_j \end{pmatrix} | Z_j, \gamma_j, \alpha_j \sim \mathcal{N}\left( Z_j \begin{pmatrix} \gamma_j \\ \beta\gamma_j + \alpha_j \end{pmatrix}, \begin{pmatrix} \sigma_u^2 & r_g\sigma_u\sigma_v \\ r_g\sigma_u\sigma_v & \sigma_v^2 \end{pmatrix} + \begin{pmatrix} c_1\hat{s}^2_{X,j} & c_{12}\hat{s}_{X,j}\hat{s}_{Y,j} \\ c_{12}\hat{s}_{X,j}\hat{s}_{Y,j} & c_2\hat{s}^2_{Y,j} \end{pmatrix} \right). \tag{S.46}$$

Comparing Eq. [S.46] with Eq. [S.43], MR-APSS performs MR analysis based on the foreground component ($Z_j = 1$). The core term which captures the causal relationship is $\begin{pmatrix} \gamma_j \\ \beta\gamma_j + \alpha_j \end{pmatrix}$ with $Z_j = 1$. As $\gamma_j$, $Z_j$, and $M_t$ are random variables in the MR-APSS model, we define the total IV strength of MR-APSS using its expectation as:

$$\text{Total IV strength for MR-APSS} := \mathbb{E}\left(\sum_{j\in\{1,\ldots,M_t\}\text{ s.t. } Z_j=1} \gamma_j^2 \Big| t\right) = \mathbb{E}\left(\sum_{j=1}^{M_t} Z_j\gamma_j^2 \Big| t\right). \quad \text{(S.47)}$$

Correspondingly, we define the average IV strength of the $M_t$ IVs selected based on threshold $t$ ($|\hat{\gamma}_j/\hat{s}_{X,j}| \geq t$) by:

$$\text{Average IV strength for MR-APSS} = \mathbb{E}\left(\frac{1}{M_t}\sum_{j=1}^{M_t} Z_j\gamma_j^2 \Big| t\right). \quad \text{(S.48)}$$

Next, we need to find connection between our definitions and the definitions given in MR literature (e.g., Eq. [S.44]). Please be noted that $\sigma^2_{X_j} = 1/N_1$ in our setting because genotypes and phenotypes are assumed to be standardized. Therefore, Eq. [S.44] can be further written as

$$\text{Average IV strength} := \frac{1}{M_t}\sum_{j=1}^{M_t} \frac{\gamma_j^2}{\hat{s}^2_{X,j}} = N_1\|\boldsymbol{\gamma}\|_2^2/M_t.$$

In this sense, our definitions is closely related to the definitions of the IV strength in the literature except that we have an additional variable $Z_j$ to indicate whether the $j$-th SNP is a valid IV. As our definitions only involves the foreground effect $\gamma_j$ of the $j$-th IV with $Z_j = 1$, it naturally excludes the direct effect $\alpha_j$, the polygenic effect $u_j$ and estimation error $e_j$ because they are affected by uncorrelated pleiotropy, correlated pleiotropy and sample structure (see Eq. [S.45]), respectively.

So far, we have mainly discussed the definitions of the IV strengths for the summary-level MR methods. Here we would like to use GENIUS-MAWII as an example to discuss the IV strength defined by the individual-level methods. Different from the summary-level MR methods which use SNP effect sizes to define the IV strengths, both GENIUS and GENIUS-MAWII leverages heteroscedasticity of the exposure to perform causal inference. GENIUS-MAWII further extends GENIUS to account for the utility of many weak IVs. To see the key idea of GENIUS-MAWII, we consider a simple model:

$$X = \gamma(G) + U,\ Y = \beta X + \alpha(G) + U,$$

where $G$ and $U$ are independent, and $\alpha(G)$ represents the influence of pleiotropy. Pleiotropy $\alpha(G)$ can bias the causal effect estimate as it induces the violation of (A-III). In this case, GENIUS-MAWII makes use of the following relationship to address the challenge:

$$\frac{\mathbb{E}\{[G-\mathbb{E}(G)][X-\mathbb{E}(X|G)]Y\}}{\mathbb{E}\{[G-\mathbb{E}(G)][X-\mathbb{E}(X|G)]X\}} = \beta,$$

where $\mathbb{E}\{[G-\mathbb{E}(G)][X-\mathbb{E}(X|G)]X\} = \text{Cov}[G, \text{Var}(X|G)] \neq 0$ (heteroscedasticity of the exposure) holds. The above equation illustrates that GENIUS-MAWII regards $\text{Cov}[G, \text{Var}(X|G)]$

as "valid IVs" to perform causal inference. Correspondingly, GENIUS-MAWII essentially makes use of quantities $|\text{Cov}[G, \text{Var}(X|G)]|$ to define measure of weak identification. Here we can see that GENIUS-MAWII uses a different type of information as the IV strength. Therefore, the proposed MR-APSS method and GENIUS-MAWII are quite complementary to each other. Comparison results of these methods have been included in SI Appendix Figs. S17-S21.

#### 2.6 Theoretical analysis of the IVW and dIVW estimators under the MR-APSS model

##### 2.6.1 The IVW estimator

We show that the IVW estimator is asymptotically biased under the MR-APSS model in the presence of pleiotropy and sample structure. To do this, we first assume that all the selected $M_t$ IVs carry both background and foreground components. Without loss of the key idea of MR-APSS, this assumption is helpful to simplify the theoretical derivation. With this assumption, we have the following MR-APSS model,

$$\begin{pmatrix}\hat{\Gamma}_j\\ \hat{\gamma}_j\end{pmatrix} \sim \mathcal{N}\left(\begin{pmatrix}v_j+\beta\gamma_j+\alpha_j\\ u_j+\gamma_j\end{pmatrix}, \begin{pmatrix}c_2\hat{s}^2_{Y,j} & c_{12}\hat{s}_{X,j}\hat{s}_{Y,j}\\ c_{12}\hat{s}_{X,j}\hat{s}_{Y,j} & c_1\hat{s}^2_{X,j}\end{pmatrix}\right), j=1,2,...,M_t,$$

where $u_j$ and $v_j$ are polygenic effects of the $j$-th IV on the outcome and the exposure traits, effects $\beta\gamma_j+\alpha_j, \gamma_j$ are the foreground components, and the variance-covariance matrix is related to the influence of sample structure.

To facilitate the analysis of asymptotic properties of the IVW estimator under MR-APSS, we follow the setting of theoretical analysis in dIVW [38] and RAPS [41]. We consider the case that all the underlying effects $v_j, u_j, \gamma_j, \alpha_j$ have been realized and fixed. We denote

$$\begin{pmatrix}\mu_{1j}\\ \mu_{2j}\end{pmatrix} = \begin{pmatrix}v_j+\beta\gamma_j+\alpha_j\\ u_j+\gamma_j\end{pmatrix}, \quad \begin{pmatrix}S_{11j} & S_{12j}\\ S_{12j} & S_{22j}\end{pmatrix} = \begin{pmatrix}c_2\hat{s}^2_{Y,j} & c_{12}\hat{s}_{X,j}\hat{s}_{Y,j}\\ c_{12}\hat{s}_{X,j}\hat{s}_{Y,j} & c_1\hat{s}^2_{X,j}\end{pmatrix}.$$

Then we have

$$\begin{pmatrix}\hat{\Gamma}_j\\ \hat{\gamma}_j\end{pmatrix} \sim \mathcal{N}\left(\begin{pmatrix}\mu_{1j}\\ \mu_{2j}\end{pmatrix}, \begin{pmatrix}S_{11j} & S_{12j}\\ S_{12j} & S_{22j}\end{pmatrix}\right), \quad j=1,2,...,M_t.$$

By the definition of the IVW estimator, we have

$$\hat{\beta}_{IVW} - \beta = \frac{\sum_{j=1}^{M_t}\left(\hat{\Gamma}_j\hat{\gamma}_j - \beta\hat{\gamma}_j^2\right)\hat{s}^{-2}_{Y,j}}{\sum_{j=1}^{M_t}\hat{\gamma}_j^2\hat{s}^{-2}_{Y,j}}. \tag{S.49}$$

For every $j$, we have

$$\mathbb{E}(\hat{\gamma}_j^2\hat{s}^{-2}_{Y,j}) = (\mu^2_{2j}+S_{22j})\hat{s}^{-2}_{Y,j},$$
$$\text{Var}(\hat{\gamma}_j^2\hat{s}^{-2}_{Y,j}) = [\mathbb{E}(\hat{\gamma}_j^4)-\mathbb{E}^2(\hat{\gamma}_j^2)]\hat{s}^{-4}_{Y,j} = (4\mu^2_{2j}+2S_{22j})S_{22j}.$$

To simply notation, we define

$$w_j = \mu^2_{2j}\hat{s}^{-2}_{Y,j},\ v_j = S_{22j}\hat{s}^{-2}_{Y,j},\ \kappa = \frac{1}{M_t}\sum_{j=1}^{M_t}\frac{\mu^2_{2j}}{S_{22j}} = \frac{1}{M_t}\sum_{j=1}^{M_t}\frac{(u_j+\gamma_j)^2}{c_1^2\hat{s}^2_{X,j}}.$$

With these notations, we have

$$\mathbb{E}(\hat{\gamma}_j^2 \hat{s}_{Y,j}^{-2}) = w_j + v_j,\ \mathrm{Var}(\hat{\gamma}_j^2 \hat{s}_{Y,j}^{-2}) = (4w_j + 2v_j)v_j,\ \kappa = \frac{1}{M_t}\sum_{j=1}^{M_t} \frac{w_j}{v_j}.$$

By the definition of $v_j = \frac{c_1 \hat{s}_{X,j}^2}{\hat{s}_{Y,j}^2}$, it is reasonable to require that $v_j$ is bounded when $M_t \to \infty$, because this assumption holds when the sample sizes of exposure $X$ and outcome $Y$ diverge in the same order. Hence, we obtain

$$\frac{\mathrm{Var}(\hat{\gamma}_j^2 \hat{s}_{Y,j}^{-2})}{[\sum_{j=1}^{M_t}(w_j + v_j)]^2} = \frac{\sum_{j=1}^{M_t}(4w_j + 2v_j)v_j}{[\sum_{j=1}^{M_t}(w_j + v_j)]^2} \le \frac{4[\sum_{j=1}^{M_t}(w_j + v_j)]\max_j v_j}{[\sum_{j=1}^{M_t}(w_j + v_j)]^2} = O(\frac{1}{\kappa M_t + M_t}) = o(1),$$

as $\kappa M_t + M_t \to \infty$. By Markov's inequality, we have

$$\frac{\sum_{j=1}^{M_t} \hat{\gamma}_j^2 \hat{s}_{Y,j}^{-2}}{\sum_{j=1}^{M_t}(w_j + v_j)} \xrightarrow{p} 1. \tag{S.50}$$

Next, we evaluate mean and variance of the numerator given in Eq. [S.49]. We have the following expression for every $j$,

$$\mathbb{E}[(\hat{\Gamma}_j\hat{\gamma}_j - \beta\hat{\gamma}_j^2)\hat{s}_{Y,j}^{-2}] = (\mu_{1j}\mu_{2j} + S_{12j} - \beta\mu_{2j}^2 - \beta S_{22j})\hat{s}_{Y,j}^{-2}.$$

$$\begin{aligned}
&\mathrm{Var}[(\hat{\Gamma}_j\hat{\gamma}_j - \beta\hat{\gamma}_j^2)\hat{s}_{Y,j}^{-2}] \\
&= [\mathbb{E}(\hat{\Gamma}_j^2\hat{\gamma}_j^2) - 2\beta\,\mathbb{E}(\hat{\Gamma}_j\hat{\gamma}_j^3) + \beta^2\,\mathbb{E}(\hat{\gamma}_j^4) - \mathbb{E}^2(\hat{\Gamma}_j\hat{\gamma}_j - \beta\hat{\gamma}_j^2)]\hat{s}_{Y,j}^{-4} \\
&= \frac{\beta^2(4\mu_{2j}^2 S_{22j} + 2S_{22j}^2) - \beta(4S_{12j}S_{22j} + 4\mu_{2j}^2 S_{12j} + 4\mu_{1j}\mu_{2j}S_{22j}) + S_{12j}^2 + 2\mu_{1j}\mu_{2j}S_{12j} + \mu_{1j}^2 S_{22j} + \mu_{2j}^2 S_{11j} + S_{11j}S_{22j}}{\hat{s}_{Y,j}^4}.
\end{aligned}$$

To simplify the notation, we denote

$$b_j = \mathbb{E}[(\hat{\Gamma}_j\hat{\gamma}_j - \beta\hat{\gamma}_j^2)\hat{s}_{Y,j}^{-2}],\ a_j^2 = \mathrm{Var}[(\hat{\Gamma}_j\hat{\gamma}_j - \beta\hat{\gamma}_j^2)\hat{s}_{Y,j}^{-2}],$$

and further define

$$K_j = \frac{(\hat{\Gamma}_j\hat{\gamma}_j - \beta\hat{\gamma}_j^2)\hat{s}_{Y,j}^{-2} - b_j}{a_j},\ \sigma_p^2 = \sum_{j=1}^{M_t} a_j^2.$$

Then, under the condition that $\max_j(a_j^2/\sigma_p^2) = o(1)$ as $M_t \to \infty$, for any $\epsilon > 0$, we have

$$\sum_{j=1}^{M_t} \mathbb{E}\left[\frac{a_j^2 K_j^2}{\sigma_p^2} I_{\{a_j|K_j| > \epsilon\sigma_p\}}\right] \le \sum_{j=1}^{M_t} \frac{a_j^2}{\sigma_p^2} \max_j \mathbb{E}[K_j^2 I_{\{a_j|K_j|>\epsilon\sigma_p\}}] = \max_j \mathbb{E}[K_j^2 I_{\{a_j|K_j|>\epsilon\sigma_p\}}] = o(1), \tag{S.51}$$

as $M_t \to \infty$. Inequality (S.51) verifies Lindeberg's condition. Hence, by Lindeberg central limit theorem, as $M_t \to \infty$,

$$\frac{\sum_{j=1}^{M_t}(\hat{\Gamma}_j\hat{\gamma}_j - \beta\hat{\gamma}_j^2)\hat{s}_{Y,j}^{-2} - \sum_{j=1}^{M_t} b_j}{(\sum_{j=1}^{M_t} a_j^2)^{1/2}} \xrightarrow{d} \mathcal{N}(0,1). \tag{S.52}$$

Now we can define the bias term and the variance term as

$$\mathrm{bias}_{IVW} = \frac{\sum_{j=1}^{M_t} b_j}{\sum_{j=1}^{M_t}(w_j + v_j)}, V_{IVW} = \frac{\sum_{j=1}^{M_t} a_j^2}{[\sum_{j=1}^{M_t}(w_j + v_j)]^2}.$$

Combining Eqs. [S.50] and [S.52], we have the following result by Slutsky's theorem,

$$V_{IVW}^{-1/2}(\hat{\beta}_{IVW} - \beta - \text{bias}_{IVW}) \xrightarrow{d} \mathcal{N}(0, 1).$$

To see the asymptotic bias of $\hat{\beta}_{IVW}$, we need to check the limit of the following term as $M_t \to \infty$,

$$\frac{\text{bias}_{IVW}}{V_{IVW}^{1/2}} = \frac{\frac{\sum_{j=1}^{M_t} b_j}{\sum_{j=1}^{M_t}(w_j+v_j)}}{\sqrt{\frac{\sum_{j=1}^{M_t} a_j^2}{[\sum_{j=1}^{M_t}(w_j+v_j)]^2}}} = \frac{\sum_{j=1}^{M_t} b_j}{\sqrt{\sum_{j=1}^{M_t} a_j^2}}$$

$$= \frac{\sum_{j=1}^{M_t}[(\mu_{1j}\mu_{2j} + S_{12j} - \beta\mu_{2j}^2 - \beta S_{22j})\hat{s}_{Y,j}^{-2}]}{\{\sum_{j=1}^{M_t}[\beta^2(4\mu_{2j}^2 S_{22j} + 2S_{22j}^2) - \beta(4S_{12j}S_{22j} + 4\mu_{2j}^2 S_{12j} + 4\mu_{1j}\mu_{2j}S_{22j}) + S_{12j}^2 + 2\mu_{1j}\mu_{2j}S_{12j} + \mu_{1j}^2 S_{22j} + \mu_{2j}^2 S_{11j} + S_{11j}S_{22j}]\hat{s}_{Y,j}^{-4}\}^{1/2}}. \tag{S.53}$$

Now we consider the case of using strong IVs, where $\mu_{1j}^2\hat{s}_{Y,j}^{-2}$, $\mu_{1j}\mu_{2j}\hat{s}_{Y,j}^{-2}$, $\mu_{2j}^2\hat{s}_{Y,j}^{-2}$ are higher-order terms compared with $S_{11j}\hat{s}_{Y,j}^{-2}$, $S_{12j}\hat{s}_{Y,j}^{-2}$, $S_{22j}\hat{s}_{Y,j}^{-2}$, as $M_t \to \infty$. In such case, the dominant term in $\text{bias}_{IVW}/V_{IVW}^{1/2}$ is roughly

$$\frac{\sum_{j=1}^{M_t}(\mu_{1j}\mu_{2j} - \beta\mu_{2j}^2)\hat{s}_{Y,j}^{-2}}{[\sum_{j=1}^{M_t}(4\beta^2\mu_{2j}^2 S_{22j} - 4\beta\mu_{2j}^2 S_{12j} - 4\beta\mu_{1j}\mu_{2j}S_{22j} + 2\mu_{1j}\mu_{2j}S_{12j} + \mu_{1j}^2 S_{22j} + \mu_{2j}^2 S_{11j})\hat{s}_{Y,j}^{-4}]^{1/2}}.$$

Hence, noting that $\mu_{1j} = v_j + \beta\gamma_j + \alpha_j$, $\mu_{2j} = u_j + \gamma_j$, the asymptotic bias of the IVW estimator can be induced by the correlation of polygenic effects $u_j, v_j$ due to the presence of correlated pleiotropy.

In the case of using many weak IVs where the influence of terms $S_{11j}\hat{s}_{Y,j}^{-2}$, $S_{12j}\hat{s}_{Y,j}^{-2}$, $S_{22j}\hat{s}_{Y,j}^{-2}$ can not be neglected compared to that of $\mu_{1j}^2\hat{s}_{Y,j}^{-2}$, $\mu_{1j}\mu_{2j}\hat{s}_{Y,j}^{-2}$, $\mu_{2j}^2\hat{s}_{Y,j}^{-2}$ as $M_t \to \infty$. As indicated by Eq. [S.53], the non-zero $c_{12}$ in $S_{12j} = c_{12}\hat{s}_{X,j}\hat{s}_{Y,j}$ due to sample structure can also induce the asymptotic bias of the IVW estimator.

##### 2.6.2 The dIVW estimator

We show that the dIVW estimator is asymptotically biased under the MR-APSS model in the presence of pleiotropy and sample structure. Let $\hat{\Gamma}_j, \hat{\gamma}_j$ be the estimates of the $j$-th IV's effects $\Gamma_j, \gamma_j$ on the outcome and the exposure, respectively. Let $\hat{s}_{Y,j}, \hat{s}_{X,j}$ be the corresponding standard errors of the estimates. Because of the large sample size of GWAS, the uncertainty in estimating $\hat{s}_{Y,j}, \hat{s}_{X,j}$ can be ignored. The dIVW estimator is developed based on the following model:

$$\hat{\Gamma}_j|\gamma_j, \alpha_{0j} \sim \mathcal{N}(\beta_0\gamma_j + \alpha_{0j}, \hat{s}_{Y,j}^2),\ \alpha_{0j} \sim \mathcal{N}(0, \tau_0^2),\ \hat{\gamma}_j \sim \mathcal{N}(\gamma_j, \hat{s}_{X,j}^2),$$

where $\beta_0$ is causal effect, $\alpha_{0j}$ accounts for horizontal pleiotropy, and $\tau_0^2$ is the variance of horizontal pleiotropic effects. It is worthwhile to mention that the above dIVW model can be regarded as a simplified version of the RAPS model. The difference is that RAPS further robustly accounts for the potential existence of outliers in horizontal pleiotropic effects (for some $j$, $\alpha_{0j}$ can be much larger than what is predicted by $\alpha_{0j} \sim \mathcal{N}(0, \tau_0^2)$). By demonstrating the bias of dIVW under MR-APSS model, we are able to explain the key reason that causes the biases of MR methods in group 2 under our proposed MR-APSS model. For the sake of simplicity, in the following analysis, we do not include the discussion related to the selection of

IVs and the selection bias. In this case, the dIVW estimator is written as:

$$\hat{\beta}_{dIVW} = \frac{\sum_j \hat{\Gamma}_j \hat{\gamma}_j \hat{s}_{Y,j}^{-2}}{\sum_j (\hat{\gamma}_j^2 - \hat{s}_{X,j}^2)\hat{s}_{Y,j}^{-2}},\ \hat{\tau}_{dIVW}^2 = \frac{\sum_j [(\hat{\Gamma}_j - \hat{\beta}_{dIVW}\hat{\gamma}_j)^2 - \hat{s}_{Y,j}^2 - \hat{\beta}_{dIVW}^2 \hat{s}_{X,j}^2]\hat{s}_{Y,j}^{-2}}{\sum_j \hat{s}_{Y,j}^{-2}}.$$

As $\hat{\beta}_{dIVW}$ does not depend on $\hat{\tau}_{dIVW}^2$, we only focus on the analysis of $\hat{\beta}_{dIVW}$. We will show that $\hat{\beta}_{dIVW}$ is asymptotically biased under the MR-APSS model.

Following a similar argument in the theoretical justification for the bias of the IVW method, here we assume that all the selected $M_t$ IVs carry both background and foreground components to show dIVW is biased under the MR-APSS model,. With this assumption, we have the following MR-APSS model,

$$\begin{pmatrix} \hat{\Gamma}_j \\ \hat{\gamma}_j \end{pmatrix} \sim \mathcal{N}\left( \begin{pmatrix} v_j + \beta\gamma_j + \alpha_j \\ u_j + \gamma_j \end{pmatrix}, \begin{pmatrix} c_2 \hat{s}_{Y,j}^2 & c_{12}\hat{s}_{X,j}\hat{s}_{Y,j} \\ c_{12}\hat{s}_{X,j}\hat{s}_{Y,j} & c_1 \hat{s}_{X,j}^2 \end{pmatrix} \right), j = 1, 2, ..., M_t,$$

where $u_j, v_j$ are polygenic effects of the $j$-th IV on the outcome and the exposure traits, $\beta\gamma_j + \alpha_j, \gamma_j$ are the foreground effects, and the variance-covariance matrix is related to the influence of sample structure. Following the setting of theoretical analysis in dIVW [38], we consider the case that all the underlying effects $v_j, u_j, \gamma_j, \alpha_j$ have been realized and fixed. We denote

$$\begin{pmatrix} \mu_{1j} \\ \mu_{2j} \end{pmatrix} = \begin{pmatrix} v_j + \beta\gamma_j + \alpha_j \\ u_j + \gamma_j \end{pmatrix}, \quad \begin{pmatrix} S_{11j} & S_{12j} \\ S_{12j} & S_{22j} \end{pmatrix} = \begin{pmatrix} c_2 \hat{s}_{Y,j}^2 & c_{12}\hat{s}_{X,j}\hat{s}_{Y,j} \\ c_{12}\hat{s}_{X,j}\hat{s}_{Y,j} & c_1 \hat{s}_{X,j}^2 \end{pmatrix}.$$

Then we have

$$\begin{pmatrix} \hat{\Gamma}_j \\ \hat{\gamma}_j \end{pmatrix} \sim \mathcal{N}\left( \begin{pmatrix} \mu_{1j} \\ \mu_{2j} \end{pmatrix}, \begin{pmatrix} S_{11j} & S_{12j} \\ S_{12j} & S_{22j} \end{pmatrix} \right), \quad j = 1, 2, ..., M_t.$$

By the definition of the dIVW estimator, we have

$$\hat{\beta}_{dIVW} - \beta = \frac{\sum_{j=1}^{M_t} \left( \hat{\Gamma}_j \hat{\gamma}_j - \beta\hat{\gamma}_j^2 + \beta\hat{s}_{X,j}^2 \right) \hat{s}_{Y,j}^{-2}}{\sum_{j=1}^{M_t} (\hat{\gamma}_j^2 - \hat{s}_{X,j}^2)\hat{s}_{Y,j}^{-2}}. \tag{S.54}$$

For every $j$, we have

$$\mathbb{E}[(\hat{\gamma}_j^2 - \hat{s}_{X,j}^2)\hat{s}_{Y,j}^{-2}] = (\mu_{2j}^2 + S_{22j} - \hat{s}_{X,j}^2)\hat{s}_{Y,j}^{-2},$$
$$\mathrm{Var}[(\hat{\gamma}_j^2 - \hat{s}_{X,j}^2)\hat{s}_{Y,j}^{-2}] = (4\mu_{2j}^2 + 2S_{22j})S_{22j}.$$

Note that it is sufficient to show that the dIVW estimator is biased under the MR-APSS model with $c_1 = c_2 = 1$ as it is a special case of the MR-APSS model. To simplify the theoretical derivation, we then consider the case $S_{11j} = \hat{s}_{Y,j}^2, S_{22j} = \hat{s}_{X,j}^2$. By further defining

$$w_j = \mu_{2j}^2 \hat{s}_{Y,j}^{-2},\ v_j = S_{22j}\hat{s}_{Y,j}^{-2},\ \kappa = \frac{1}{M_t}\sum_{j=1}^{M_t} \frac{\mu_{2j}^2}{S_{22j}} = \frac{1}{M_t}\sum_{j=1}^{M_t} \frac{(u_j + \gamma_j)^2}{\hat{s}_{X,j}^2},$$

we have

$$\mathbb{E}[(\hat{\gamma}_j^2 - \hat{s}_{X,j}^2)\hat{s}_{Y,j}^{-2}] = w_j,\ \mathrm{Var}(\hat{\gamma}_j^2 \hat{s}_{Y,j}^{-2}) = (4w_j + 2v_j)v_j,\ \kappa = \frac{1}{M_t}\sum_{j=1}^{M_t} \frac{w_j}{v_j}.$$

By the definition of $v_j = \frac{\hat{s}^2_{X,j}}{\hat{s}^2_{Y,j}}$, it is reasonable to require that $v_j$ is bounded when $M_t \to \infty$, because this assumption holds when the sample sizes of exposure $X$ and outcome $Y$ diverge in the same order. As this assumption often holds in real applications, it is reasonable to assume that there exists a constant $C > 0$ such that $C^{-1} \leq v_j \leq C$, $\forall j$. Hence, we obtain

$$C^{-1}\sum_{j=1}^{M_t} w_j \leq \kappa M_t = \sum_{j=1}^{M_t} \frac{w_j}{v_j} \leq C \sum_{j=1}^{M_t} w_j,$$

therefore,

$$\frac{\mathrm{Var}[(\hat{\gamma}_j^2 - \hat{s}^2_{X,j})\hat{s}^{-2}_{Y,j}]}{(\sum_{j=1}^{M_t} w_j)^2} \leq \frac{4C\sum_{j=1}^{M_t} w_j + 2M_t C^2}{(\sum_{j=1}^{M_t} w_j)^2} = \frac{4C^2\kappa M_t + 2C^2 M_t}{C^{-2}\kappa^2 M_t^2} = O(\frac{1}{\kappa M_t} + \frac{1}{\kappa^2 M_t}).$$

Following the same condition, $\kappa\sqrt{M_t} \to \infty$ as $M_t \to \infty$, required by theoretical analysis in dIVW, we obtain

$$\frac{\mathrm{Var}[(\hat{\gamma}_j^2 - \hat{s}^2_{X,j})\hat{s}^{-2}_{Y,j}]}{(\sum_{j=1}^{M_t} w_j)^2} \leq o(1), \text{ as } M_t \to \infty.$$

By Markov's inequality, we have

$$\frac{\sum_{j=1}^{M_t}(\hat{\gamma}_j^2 - \hat{s}^2_{X,j})\hat{s}^{-2}_{Y,j}}{\sum_{j=1}^{M_t} w_j} \xrightarrow{p} 1. \tag{S.55}$$

Next, we evaluate mean and variance of the numerator given in Eq. [S.54]. For every $j$,

$$\mathbb{E}[(\hat{\Gamma}_j\hat{\gamma}_j - \beta\hat{\gamma}_j^2 + \beta\hat{s}^2_{X,j})\hat{s}^{-2}_{Y,j}] = (\mu_{1j}\mu_{2j} + S_{12j} - \beta\mu_{2j}^2)\hat{s}^{-2}_{Y,j}.$$

$$\mathrm{Var}[(\hat{\Gamma}_j\hat{\gamma}_j - \beta\hat{\gamma}_j^2 + \beta\hat{s}^2_{X,j})\hat{s}^{-2}_{Y,j}]$$
$$= \frac{\beta^2(4\mu_{2j}^2 S_{22j} + 2S_{22j}^2) - \beta(4S_{12j}S_{22j} + 4\mu_{2j}^2 S_{12j} + 4\mu_{1j}\mu_{2j}S_{22j}) + S_{12j}^2 + 2\mu_{1j}\mu_{2j}S_{12j} + \mu_{1j}^2 S_{22j} + \mu_{2j}^2 S_{11j} + S_{11j}S_{22j}}{\hat{s}^4_{Y,j}}.$$

We denote

$$b_j = \mathbb{E}[(\hat{\Gamma}_j\hat{\gamma}_j - \beta\hat{\gamma}_j^2 + \beta\hat{s}^2_{X,j})\hat{s}^{-2}_{Y,j}],\ a_j^2 = \mathrm{Var}[(\hat{\Gamma}_j\hat{\gamma}_j - \beta\hat{\gamma}_j^2 + \beta\hat{s}^2_{X,j})\hat{s}^{-2}_{Y,j}],$$

and

$$K_j = \frac{(\hat{\Gamma}_j\hat{\gamma}_j - \beta\hat{\gamma}_j^2 + \beta\hat{s}^2_{X,j})\hat{s}^{-2}_{Y,j} - b_j}{a_j},\ \sigma_p^2 = \sum_{j=1}^{M_t} a_j^2.$$

Then, under the condition that $\max_j(a_j^2/\sigma_p^2) = o(1)$ as $M_t \to \infty$, for any $\epsilon > 0$, we have

$$\sum_{j=1}^{M_t}\mathbb{E}\left[\frac{a_j^2 K_j^2}{\sigma_p^2} I_{\{a_j|K_j|>\epsilon\sigma_p\}}\right] \leq \sum_{j=1}^{M_t}\frac{a_j^2}{\sigma_p^2}\max_j \mathbb{E}[K_j^2 I_{\{a_j|K_j|>\epsilon\sigma_p\}}] = \max_j \mathbb{E}[K_j^2 I_{\{a_j|K_j|>\epsilon\sigma_p\}}] = o(1), \tag{S.56}$$

as $M_t \to \infty$. Inequality (S.56) verifies Lindeberg's condition. Hence, by Lindeberg central limit theorem, as $M_t \to \infty$,

$$\frac{\sum_{j=1}^{M_t}(\hat{\Gamma}_j\hat{\gamma}_j - \beta\hat{\gamma}_j^2 + \beta\hat{s}^2_{X,j})\hat{s}^{-2}_{Y,j} - \sum_{j=1}^{M_t} b_j}{(\sum_{j=1}^{M_t} a_j^2)^{1/2}} \xrightarrow{d} \mathcal{N}(0,1). \tag{S.57}$$

Define

$$\text{bias}_{IVW} = \frac{\sum_{j=1}^{M_t} b_j}{\sum_{j=1}^{M_t} w_j}, V_{dIVW} = \frac{\sum_{j=1}^{M_t} a_j^2}{(\sum_{j=1}^{M_t} w_j)^2}.$$

Combining Eqs. [S.55] and [S.57], we have the following result by Slutsky's theorem,

$$V_{dIVW}^{-1/2}(\hat{\beta}_{dIVW} - \beta - \text{bias}_{IVW}) \xrightarrow{d} \mathcal{N}(0, 1).$$

To see the asymptotic bias of $\hat{\beta}_{IVW}$, we need to check the limit of the following term as $M_t \to \infty$,

$$\frac{\text{bias}_{IVW}}{V_{dIVW}^{1/2}} = \frac{\sum_{j=1}^{M_t} b_j}{\sqrt{\sum_{j=1}^{M_t} a_j^2}}$$
$$= \frac{\sum_{j=1}^{M_t}[(\mu_{1j}\mu_{2j} + S_{12j} - \beta\mu_{2j}^2)\hat{s}_{Y,j}^{-2}]}{\{\sum_{j=1}^{M_t}[\beta^2(4\mu_{2j}^2 S_{22j} + 2S_{22j}^2) - \beta(4S_{12j}S_{22j} + 4\mu_{2j}^2 S_{12j} + 4\mu_{1j}\mu_{2j}S_{22j}) + S_{12j}^2 + 2\mu_{1j}\mu_{2j}S_{12j} + \mu_{1j}^2 S_{22j} + \mu_{2j}^2 S_{11j} + S_{11j}S_{22j}]\hat{s}_{Y,j}^{-4}\}^{1/2}}. \tag{S.58}$$

Compared to the asymptotic bias of the IVW estimator, the numerator in the asymptotic bias of the dIVW estimator corrects for the bias induced by the term $-\beta S_{22j}\hat{s}_{Y,j}^{-2}$. This is because the dIVW estimator has taken the uncertainty $\hat{s}_{X,j}^2$ of $\hat{\gamma}_j$ into account and thus can eliminate the bias due to the usage of many weak IVs to perform MR analysis. According to Eq. [S.58], however, dIVW is still biased due to its neglect of correlated pleiotropy and sample structure. Specifically, we observe that $\mu_{1j} = v_j + \beta\gamma_j + \alpha_j$, $\mu_{2j} = u_j + \gamma_j$. The asymptotic bias of the dIVW estimator can be induced by the correlation of polygenic effects $u_j, v_j$ due to the presence of correlated pleiotropy. Besides the influence of correlated pleiotropy, the influence of terms $S_{11j}\hat{s}_{Y,j}^{-2}$, $S_{12j}\hat{s}_{Y,j}^{-2}$, $S_{22j}\hat{s}_{Y,j}^{-2}$ can not be neglected in the case of using many weak IVs . As indicated by Eq. [S.58], the non-zero $c_{12}$ in $S_{12j} = c_{12}\hat{s}_{X,j}\hat{s}_{Y,j}$ due to sample structure can also induce the asymptotic bias of the dIVW estimator.

### 3 Simulation studies

#### 3.1 Simulations under the MR-APSS model

MR-APSS assumes that the effects of SNP $j$ on the exposure and the outcome have the following relationship:

$$\tilde{\gamma}_j = Z_j\gamma_j + u_j, \Gamma_j = Z_j(\beta\gamma_j + \alpha_j) + v_j,$$

where $\beta$ is causal effect of interest, $u_j, v_j$ are background signals, $\gamma_j, \alpha_j$ are foreground signals, and $Z_j$ indicates whether SNP $j$ carries foreground signals ($Z_j = 1$) or not ($Z_j = 0$). In the simulation, we assumed that the background component accounted for a total heritability of $h^2 = 0.5$ for each trait. Given that there were $M = 47,049$ SNPs, we set the background variances to be $\sigma_u^2 = \tau_v^2 = \frac{h^2}{M}$. Then the background signals were sampled by

$$\begin{pmatrix} u_j \\ v_j \end{pmatrix} \sim \mathcal{N}\left(\begin{pmatrix} 0 \\ 0 \end{pmatrix}, \begin{pmatrix} \sigma_u^2 & r_g\sigma_u\tau_v \\ r_g\sigma_u\tau_v & \tau_v^2 \end{pmatrix}\right),$$

with varying genetic correlation $r_g \in \{0.1, 0.2\}$. Additionally, we randomly chose 500 out of 47,049 SNPs to carry foreground signals. Specifically, $Z_j = 1$ were randomly assigned on 500

SNPs while $Z_j = 0$ were assigned on the remaining 46,549 SNPs. For those 500 SNPs which carried foreground signals, we assumed that the foreground-background variance ratios for exposure and outcome were $\sigma^2 : \sigma_u^2 \in \{10, 20, 40\}$ and $\tau^2 : \tau_v^2 = 1$. Then the foreground effects of the 500 SNPs were sampled by

$$\begin{pmatrix} \gamma_j \\ \alpha_j \end{pmatrix} \sim \mathcal{N}\left( \begin{pmatrix} 0 \\ 0 \end{pmatrix}, \begin{pmatrix} \sigma^2 & 0 \\ 0 & \tau^2 \end{pmatrix} \right).$$

We generated phenotypes based on simulated SNP effects and real genotypes from UKBB. To simulate scenarios with or without sample structure, we used 0 or 10,000 overlapped samples in exposure and outcome studies. If individual $i$ was shared in the exposure and outcome studies, then the environmental noises were simulated by

$$\begin{pmatrix} \epsilon_{x,i} \\ \epsilon_{y,i} \end{pmatrix} \sim \mathcal{N}\left( \begin{pmatrix} 0 \\ 0 \end{pmatrix}, \begin{pmatrix} 1 - h^2 & r_e(1 - h^2) \\ r_e(1 - h^2) & 1 - h^2 \end{pmatrix} \right),$$

where $r_e \in \{0.3, 0.6\}$. If individual $i$ was not shared, the noise terms were independently generated as

$$\epsilon_{x,i} \sim \mathcal{N}(0, 1 - h^2), \text{ and } \epsilon_{y,i} \sim \mathcal{N}(0, 1 - h^2).$$

To simulate phenotype vectors, we standardized the genotype matrices from UKBB (i.e., the standardized genotypes on each SNP to have zero mean and unit variance) and denoted them as $\mathbf{G}_x$ and $\mathbf{G}_y$, respectively. Let $\boldsymbol{\epsilon}_x$ and $\boldsymbol{\epsilon}_y$ be the random noises of exposure and outcome traits, respectively. Then, the phenotype vectors of exposure and outcome traits were simulated as

$$\mathbf{x} = \mathbf{G}_x\tilde{\boldsymbol{\gamma}} + \boldsymbol{\epsilon}_x, \ \mathbf{y} = \mathbf{G}_y\boldsymbol{\Gamma} + \boldsymbol{\epsilon}_y.$$

#### 3.2 Simulations under the CAUSE model

Following the CAUSE model, we simulated effects $(\gamma_j, \Gamma_j)$ of SNP $j$ on exposure $X$ and outcome $Y$ based on following relationship:

$$\Gamma_j = \beta\gamma_j + Z_j\eta\gamma_j + \theta_j,$$

where $\beta$ is causal effect of interest, $\eta$ is the correlated pleiotropic effect, $\theta_j$ is uncorrelated pleiotropic effect, and $Z_j$ indicates whether the SNP is affected by correlated pleiotropy ($Z_j = 1$) or not ($Z_j = 0$). CAUSE assumes sparsity for direct effects $(\gamma_j, \theta_j)$. Here we randomly assigned $P_1 = 10,000$ out of 47,049 SNPs with non-zero effects on exposure $X$. To be specific, the effects of $P_1 = 10,000$ SNPs on exposure $X$ were sampled by $\gamma_j \sim \mathcal{N}(0, \frac{h^2}{P_1})$, where $h^2 = 0.5$ is the heritability of exposure $X$. The remaining $36,049$ SNPs were not associated with exposure $X$. To simulate $Y$, we followed the assumption from CAUSE that the proportion of IVs affected by correlated pleiotropy should be lower than 50%, i.e., we sampled $Z_j$ by $Z_j \sim \text{Bern}(q)$, where $q = \Pr(Z_j = 1) < 0.5$. Here we varied $q$ as $q \in \{0.2, 0.4\}$. To ensure that the heritability of $Y$ was $h^2 = 0.5$, we then randomly chose $P_2 = 10,000$ out of $37,049$ SNPs and assigned non-zero effects as $\theta_j \sim \mathcal{N}(0, \frac{(1-\beta^2-q\eta^2)h^2}{P_2})$. Similar to the simulations based on MR-APSS model, we simulated phenotype vectors for $X$ and $Y$ by

$$\mathbf{x} = \mathbf{G}_x\boldsymbol{\gamma} + \boldsymbol{\epsilon}_x, \ \mathbf{y} = \mathbf{G}_y\boldsymbol{\Gamma} + \boldsymbol{\epsilon}_y,$$

where $\mathbf{G}_x, \mathbf{G}_y$ were standardized genotype matrices, $\boldsymbol{\epsilon}_x, \boldsymbol{\epsilon}_y$ were noise vectors sampled based on

$$\begin{pmatrix} \epsilon_{x,i} \\ \epsilon_{y,i} \end{pmatrix} \sim \mathcal{N}\left( \begin{pmatrix} 0 \\ 0 \end{pmatrix}, \begin{pmatrix} 1-h^2 & r_e(1-h^2) \\ r_e(1-h^2) & 1-h^2 \end{pmatrix} \right),$$

if individual $i$ was shared in the exposure and outcome studies, and

$$\epsilon_{x,i} \sim \mathcal{N}(0, 1-h^2), \quad \epsilon_{y,i} \sim \mathcal{N}(0, 1-h^2),$$

otherwise.

##### 3.3 Evaluation of the performance of individual-level MR methods in simulation studies

We evaluated the performance of four individual-level MR methods, including TSLS, TSHT, GENIUS, and GENIUS-MAWII, based on simulations under the MR-APSS model and the CAUSE model. We first evaluated the type I errors of these methods. Fig. S1 (A) and Fig. S6 (A) show the QQ-plots of $-\log(p)$-values produced by these methods under different settings in the MR-APSS model and the CAUSE model, respectively. Clearly, TSLS, TSHT produced inflated $p$-values in the presence of pleiotropy and sample structure. When influence of sample structure was small ($c_{12} = 0.075$ under MR-APSS model, $r_e = 0.2$ under CAUSE model), the $p$-values produced by GENIUS are well-calibrated, but its $p$-values tended to be slightly inflated when the influence of sample structure became larger ($c_{12} = 0.15$ under MR-APSS model, $r_e = 0.6$ under CAUSE model). The $p$-values of GENIUS-MAWII remained to be calibrated in all the settings. Besides GENIUS-MAWII, we also observed that the $p$-values produced by Egger, Weighted-mode, and MR-APSS were well-calibrated. Next we evaluated the power of GENIUS and GENIUS-MAWII as they provided satisfactory type I error control under the null ($\beta = 0$). We compared these two methods with MR-APSS, Egger, and CAUSE. Fig. S1 (C) and Fig. S6 (C) show that MR-APSS had a higher power than GENIUS, GENIUS-MAWII, Weighted-mode, and Egger in the simulations under the MR-APSS model as well as under the CAUSE model.

#### 4 Real data analysis

##### 4.1 GWAS summary statistics and pre-processing

For all GWAS summary datasets, we used SNPs in the set of HapMap 3 list with minor allele frequency >0.05. We further excluded SNPs in the complex Major Histocompatibility Region (Chromosome 6, 26Mb−34Mb). Following the process in LDSC [8], we checked the $\chi^2$ statistic of each SNP and excluded SNPs with $\chi^2 > \max\{80, N/1000\}$ to prevent the outliers that may unduly affect the results. For a pair of exposure and outcome traits, we took the overlapped SNPs from their GWAS summary statistics and aligned the sign of effect sizes for those SNPs to the same allele. Then we applied bivariate LDSC to estimate the $\boldsymbol{\Omega}$ and $\mathbf{C}$ using genome-wide summary statistics. After this step, we selected SNPs as IVs using an IV threshold (with the default $p$-value $5 \times 10^{-5}$), and applied PLINK clumping ($r^2 < 0.001$, window size 1Mb) to obtain nearly independent IVs. Finally, we fitted the proposed foreground-background model to infer the causal effect.

#### 4.2 Illustrative examples for the IV strength

We consider Height (GIANT) and Height (UKBB) as exposures to examine the influence of sample size on the IV strength. The sample sizes for Height (GIANT) and Height (UKBB) are 253,288 and 385,748, respectively. The $p$-value threshold for IV selection varied from $5 \times 10^{-8}$ to $5 \times 10^{-5}$. We evaluated the number of IVs, average IV strength, and total IV strength when we considered Height (GIANT) and Height (UKBB) as exposures, and each of the remaining 24 traits as an outcome (The information of these traits is listed in SI Appendix, Table S1). Based on our MR-APSS model, we define the number of valid IVs as $\pi_t M_t$, where $\pi_t = p(Z_j = 1 \big| |\hat{\gamma}_j / \hat{s}_{X_j}| \geq t)$ is the proportion of IVs with foreground signal given in Eq. [7] of main text, and $M_t$ is the number of selected IVs based on the threshold $t$. As shown in Fig. S14A and Fig. S14B, given the same IV threshold, the number selected IVs as well as the number of valid IVs of Height (UKBB) are larger than those of Height (GIANT) because Height (UKBB) has a larger sample size. Fig. S14C and Fig. S14D show the estimated average IV strength defined in Eq. [11] and total IV strength defined in Eq. [12] of the main text. The larger sample size of Height (UKBB) allows us to select more SNPs with moderate effects as IVs. Therefore, given the same IV threshold, the average IV strength of Height (UKBB) is weaker than that of Height (GAINT) but the total IV strength of Height (UKBB) is stronger. To further examine the influence of the IV threshold on estimating causal effects, we compared the estimated causal effects of Height (UKBB) and Height (GIANT) on the outcome traits. As shown in Fig. S15, despite their different sample sizes, the causal inference results of Height (UKBB) and Height (GIANT) agree with each other for different IV thresholds. Of note, the standard errors of Height (UKBB) are smaller than those of Height (GIANT).

#### 4.3 The default IV threshold for MR-APSS in real applications

Regarding the IV selection threshold, we have shown that the type I error rate of MR-APSS is insensitive to the choice of IV threshold, and the statistical power of MR-APSS can be improved by including SNPs with moderate effects using a looser IV threshold (Fig. 5 of main text). Practically, we recommend using $5 \times 10^{-5}$ as the default IV threshold. There are two major reasons. First, for most of exposure traits, we have observed that the proportion of valid IVs ($\pi_t$) decreases when the IV threshold $p$-value becomes looser, as shown in (Fig. 5A of main text). If the IV threshold becomes looser, the proportion of valid IVs can be very small as most selected SNPs belong to the background component. As we are working with a mixture model, we hope that $\pi_t$ should be bounded away from either 0 or 1. Second, perhaps more important, we have observed the selection bias due to the LD clumping procedure. To ensure that IVs are nearly independent, as a common practice, we applied LD clumping after using the IV threshold for SNP selection. Please be noted that the LD clumping procedure will retain SNPs with smaller $p$-values. When the IV threshold $p \leq 5 \times 10^{-5}$, we find the bias due to LD clumping is very small and can be corrected empirically, i.e., adjusting the IV threshold by the ratio of the median after the LD clumping to the median before LD clumping (see the details in the SI Appendix, section 1.6, Figs S9-S10). When the IV threshold becomes looser, say, $5 \times 10^{-3}$, all SNPs that survive after LD clumping will have a $p$-value much smaller than $5 \times 10^{-3}$. To our best knowledge, no method can analytically correct this bias due to the complicated process of LD clumping. Therefore, we would like to recommend using $5 \times 10^{-5}$ as the default IV threshold in real data analysis.

#### 4.4 Evaluation of the performance of individual-level MR methods in real data analysis

##### 4.4.1 Type I error control of individual-level MR methods

Different from summary-level MR methods, individual-level MR methods require that both exposure $X$ and outcome $Y$ have been measured for the individuals under consideration. To have a comparison, we use the individual-level data from UKBB. Again, we use the same five traits as the negative control outcomes. Among the 26 exposure traits used to compare the summary-level MR methods, we can extract 8 traits from UKBB, including Daytime sleepiness, Neuroticism, Angina, BMI, Height, HBP, Income, and Intelligence. In total, we have $8 \times 5 = 40$ pairs to evaluate individual-level methods. As a supplement, we also include the 10 summary-level MR methods (see Table 1 in main text) in comparison.

Regarding the data quality control (QC) when we handle individual-level datasets, we followed the QC criteria described in [12] to include individuals. In total, there are 337,209 samples satisfying these criteria. For genotypes, we only keep those SNPs in the Hapmap3 with minor allele frequency $> 0.01$, missing genotypes in less than 0.1 of the sample, and Hardy-Weinberg equilibrium (HWE) $p$-value $> 10^{-7}$. The IVs for the individual-level methods are selected using the IV threshold $p = 5 \times 10^{-8}$.

We evaluated the type I errors of four individual-level MR methods and ten summary-level MR methods based on the 40 trait pairs. Fig. S16 shows the QQ-plots of $-\log(p)$ -values of all MR methods. Clearly, TSLS and TSHT produced inflated $p$-values, while GENIUS and GENIUS-MAWII produced well-calibrated $p$-values. The obtained results suggest that the key assumption of GENIUS and GENIUS-MAWII (heteroscedasticity of the exposure) is robust in the presence of pleiotropy and sample structure. We also noticed that MR-APSS and Weighted-mode produced well-calibrated $p$-values among ten summary-level MR methods.

##### 4.4.2 MR-APSS is complementary to GENIUS and GENIUS-MAWII

So far, we have found that GENIUS, GENIUS-MAWII, MR-APSS, and Weighted-mode can produce well-calibrated $p$-values based on real data analysis using negative control outcomes. Next, we evaluated the estimation efficiency of MR methods using 8 exposure traits and negative control outcomes. Fig. S17 shows the causal effect estimates ($\hat{\beta}$) and their 95% confidence intervals (obtained as $2\times$s.e.($\hat{\beta}$)) obtained by the 14 MR methods where we used 8 exposure traits and one negative outcome trait (Hair_Blonde). The results of other four negative control outcomes are given in Fig. S18 - Fig. S21. The ground-truth of the causal effects should be zero (i.e., $\beta = 0$) because we are using the negative control outcomes. Let's first focus on the comparison between GENIUS (GENIUS-MAWII) and MR-APSS. In Fig. S17A, the 95% confidence intervals of GENIUS and GENIUS-MAWII were shorter than those of MR-APSS. Because all three methods can control the type I errors, GENIUS and GENIUS-MAWII were more efficient than MR-APSS. In Fig. S17B, the situation changed. MR-APSS had shorter 95% confidence intervals than those of GENIUS and GENIUS-MAWII. The above real data analysis can be explained by the fact that GENIUS (GENIUS-MAWII) and MR-APSS use different types of information for causal inference. The IV strength of GENIUS and GENIUS-MAWII is related to heteroscedasticity of the exposure while the IV strength of MR-APSS is related to the SNP effect sizes deviating from polygenic effects. When heteroscedasticity of the exposure is strong, GENIUS and GENIUS-MAWII can be very efficient estimators of the causal effect and they are also robust in the presence of pleiotropy and sample structure. For example, when obesity-related traits are considered as exposures, the heteroscedasticity assumption is often satisfied [33]. However, When some other traits are considered as exposures, the heteroscedasticity assumption may not hold. An example trait is height. According to Wang et al. (2019) [33], more than 1,000 independent loci have been identified by GWAS to be associated with height but no variance quantitative trait locus (vQTL) has been identified. This helps to explain why GENIUS and GENIUS are more efficient than MR-APSS when BMI is the exposure but less efficient than MR-APSS when Height is the exposure. In summary, MR-APSS is complementary to GENIUS and GENIUS-MAWII in terms of estimation efficiency.

#### 4.5 Analysis results of LCV

We have mainly focused on comparing MR-APSS with methods using IVs. We note that causal inference can be performed without using IVs, for example, a recently developed summary-level data based method: the latent causal variable (LCV) model [24]. Unlike exsisting summary-level MR methods which use instrument variables to infer the causal effect between trait pairs, LCV estimates the so-called genetic causality proportion (GCP) without using instrument variables. Under the LCV model, trait 1 is defined to be partially genetically causal for trait 2 ($0 < \text{GCP} < 1$) if part of the genetic component in trait 1 is causal for the trait 2, and trait 1 is defined to be fully genetically causal ($\text{GCP} = 1$) for trait 2 if the entire genetic component in trait 1 is causal for the trait 2. Notably, trait pairs with low GCP values have limited partial causality, and the large GCP estimate implies a plausible causal effect between traits. As suggested by the LCV paper, trait pairs with $\widehat{\text{GCP}} > 0.6$ are unlikely to be false positives. To facilitate comparison of MR-APSS with LCV, we applied LCV to the trait pairs between 26 complex traits and the five negative control outcomes. As shown in Fig. S32, LCV produced deflated $p$-values. We further applied LCV to the 320 trait pairs among 26 traits. As shown in supplementary Fig. S33, LCV identified only four trait pairs with $\widehat{\text{GCP}} > 0.6$ based on Bonferroni correction. Our results suggest that LCV tends to be conservative in real data analysis.

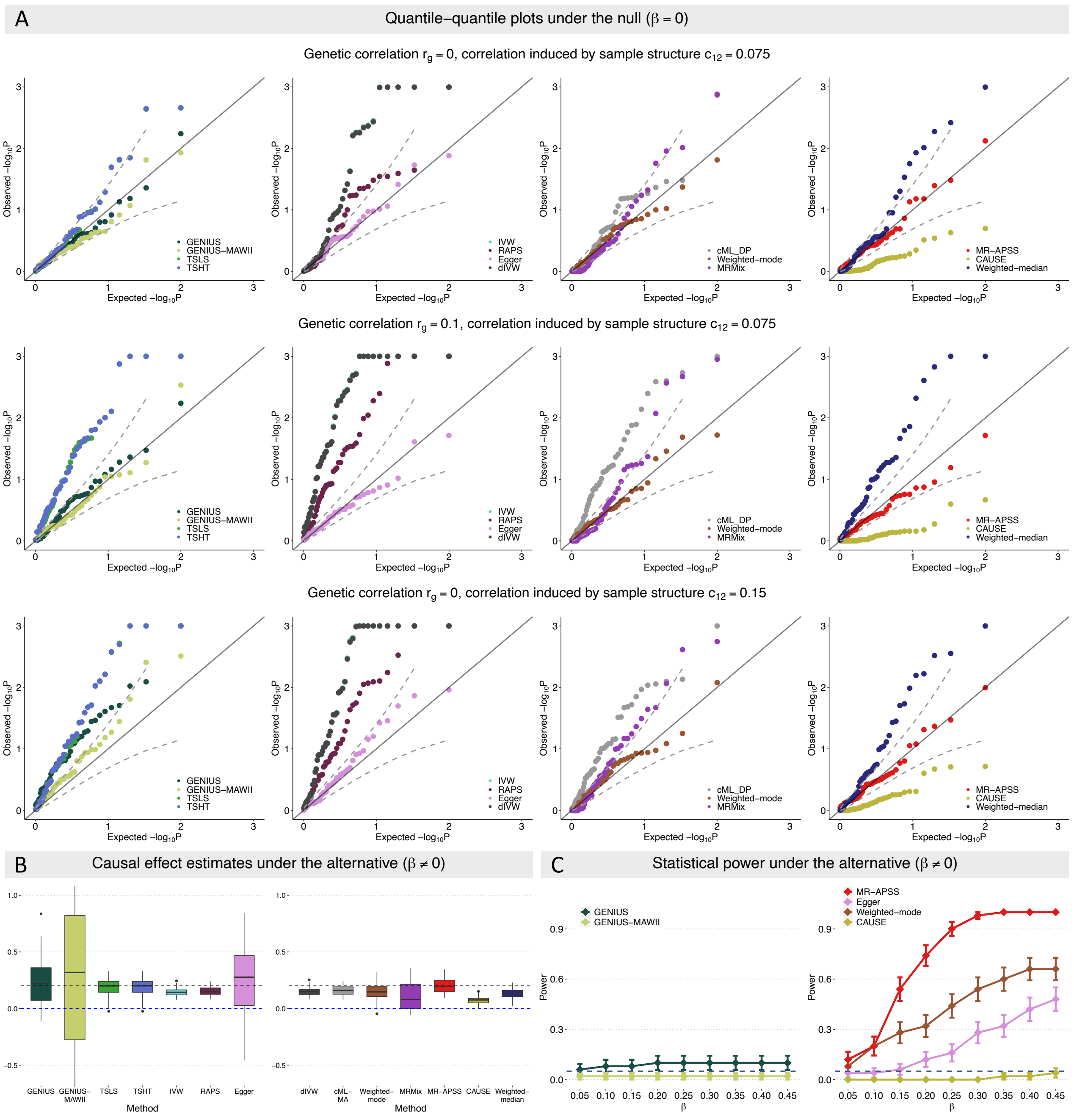

**Figure** S1: Comparison of 14 MR methods on simulated data based on the MR-APSS model. (A) Quantile-quantile plots of $-\log_{10}(p)$-values under null simulations with varying settings including (i) $r_g = 0.0, c_{12} = 0.075$, (ii) $r_g = 0.1, c_{12} = 0.075$, (iii) $r_g = 0.0, c_{12} = 0.15$. (B) Estimates of causal effect under the alternative simulations ($\beta = 0.2$). (C) Power in settings where the causal effect size $\beta$ varied from 0.05 to 0.45. The comparison of power was conducted among those methods whose type I errors were under controlled in the null simulations. The results were summarized from 50 replications.

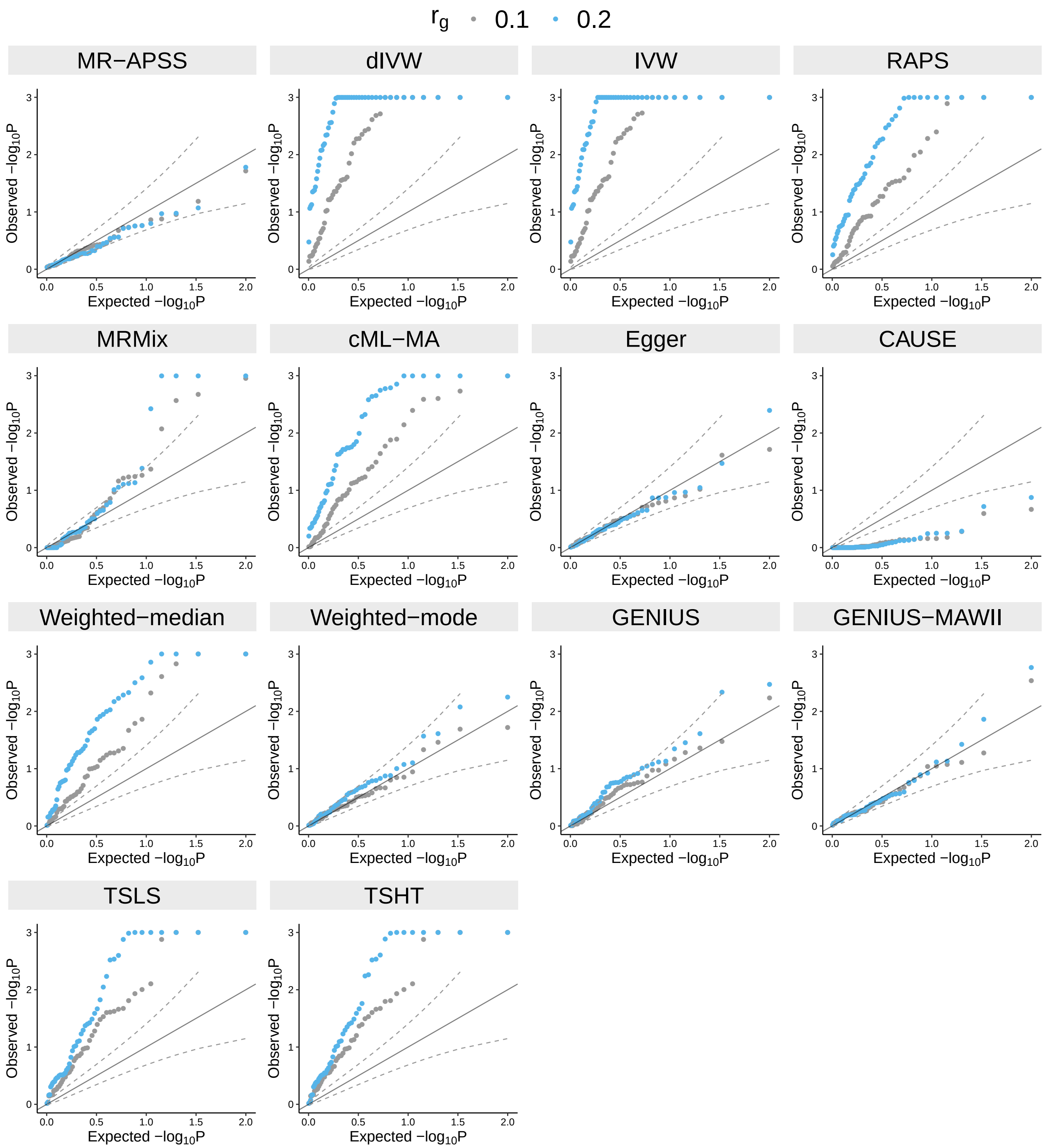

**Figure** S2: Type I error control of 14 MR methods in the presence of genetic correlation induced by pleiotropy under MR-APSS model. Quantile-quantile plots of $-\log_{10}(p)$-values under null simulations ($\beta = 0$) with varying genetic correlation $r_g \in \{0.1, 0.2\}$ and with fixed correlation in estimation errors ($c_{12} = 0.075$). The foreground-background variance ratio was set to be $\sigma^2 : \sigma_u^2 = 20, \tau^2 : \tau_v^2 = 1$. The results were summarized from 50 replications.

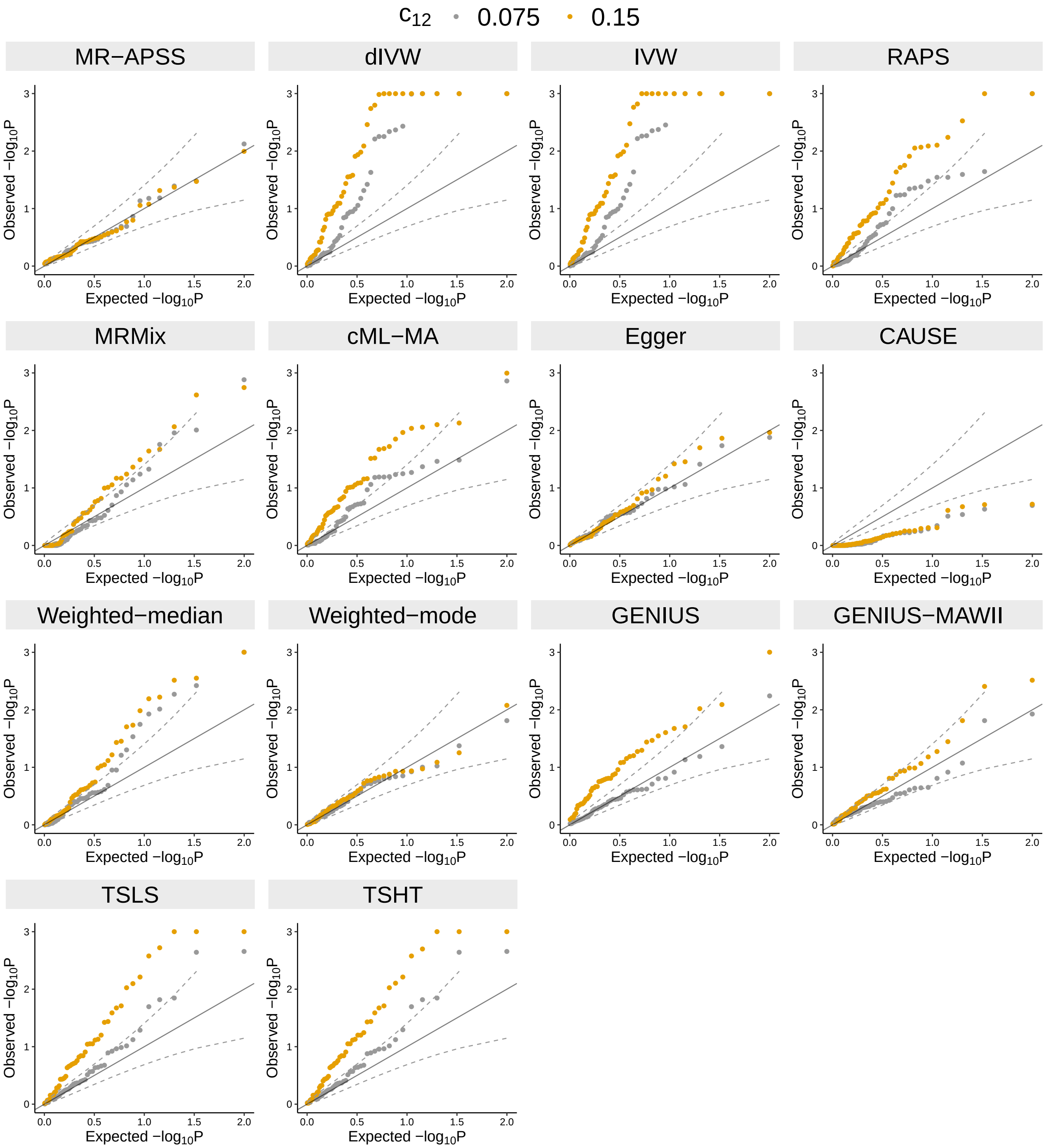

**Figure** S3: Type I error control of 14 MR methods in the presence of sample structure under MR-APSS model. Quantile-quantile plots of $-log_{10}(p)$-values under null simulations ($\beta = 0$) with genetic correlation $r_g = 0$ and with correlation in estimation errors $c_{12} \in \{0.075, 0.15\}$. The correlation in estimation errors was induced by 10,000 overlapped samples with correlation of environmental noises $r_e = 0.3$, 0.6. The foreground-background variance ratio was set to be $\sigma^2 : \sigma_u^2 = 20$, and $\tau^2 : \tau_v^2 = 1$. The results were summarized from 50 replications.

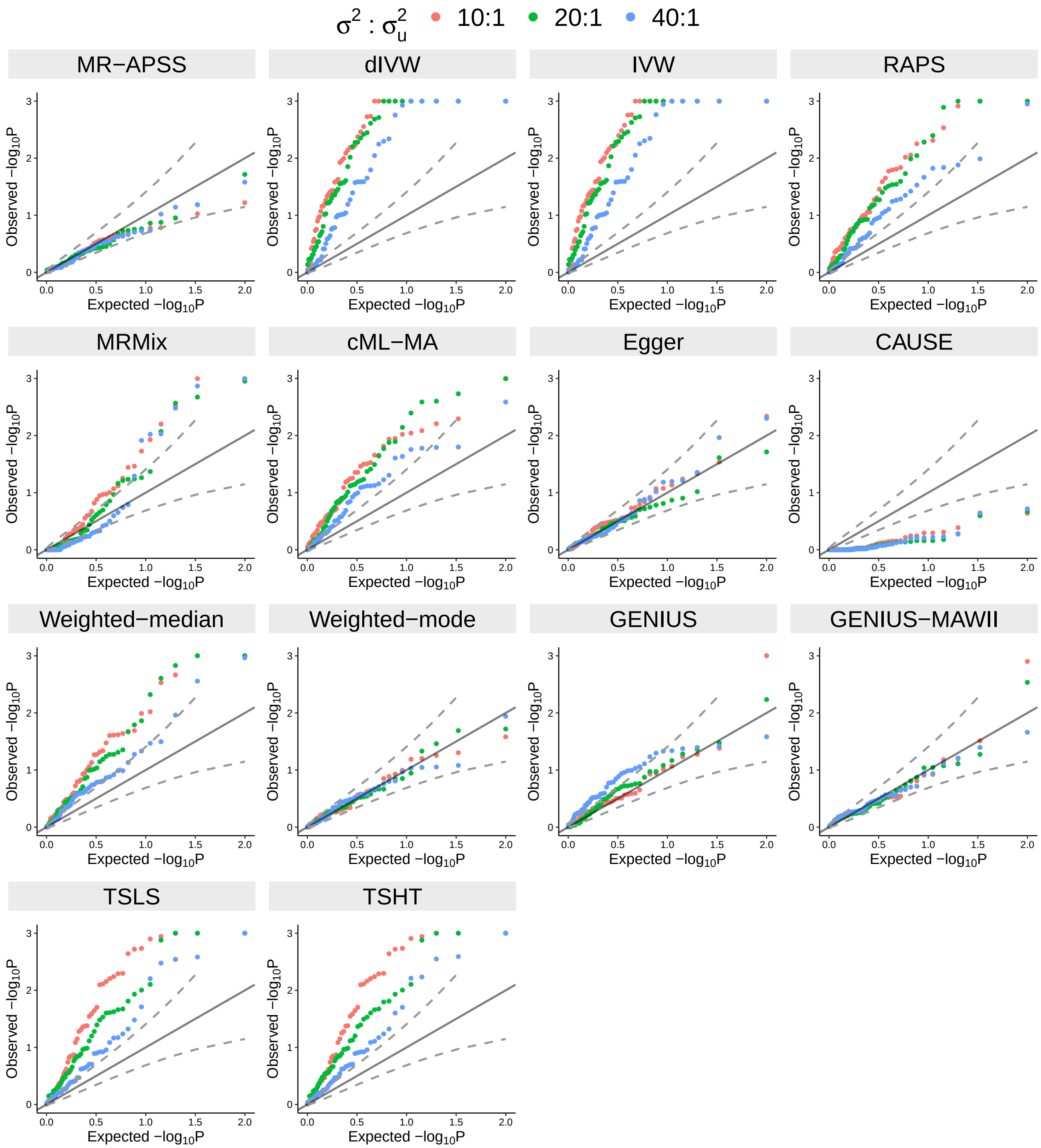

**Figure** S4: The type I error control of 14 MR methods in settings with varying foreground-background variance ratio ($\sigma^2 : \sigma_u^2$). Quantile-quantile plots of $-\log_{10}(p)$-values under null simulations ($\beta = 0$) with genetic correlation $r_g = 0.1$ and with correlation in estimation errors $c_{12} = 0.075$. The correlation in estimation errors was induced by 10,000 overlapped samples with correlation of environmental noises $r_e = 0.3$. The foreground-background variance ratio was varied as $\sigma^2 : \sigma_u^2 \in \{10, 20, 40\}$, $\tau^2 : \tau_v^2 = 1$. The results were summarized from 50 replications.

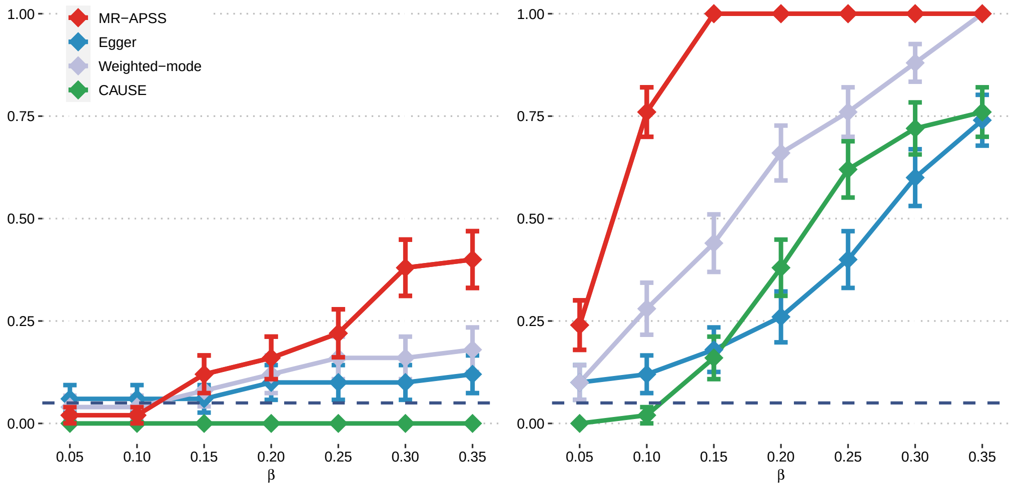

**Figure** S5: Power of MR-APSS, Egger, Weighted-mode, and CAUSE in settings with varying foreground-background variance ratio ($\sigma^2 : \sigma_u^2$). Power when causal effect size $\beta$ is varied from 0.05 to 0.35. The foreground-background variance ratio was varied at: $\sigma^2 : \sigma_u^2 = 10$ (Left), and $\sigma^2 : \sigma_u^2 = 40$ (Right). The results were summarized from 50 replications.

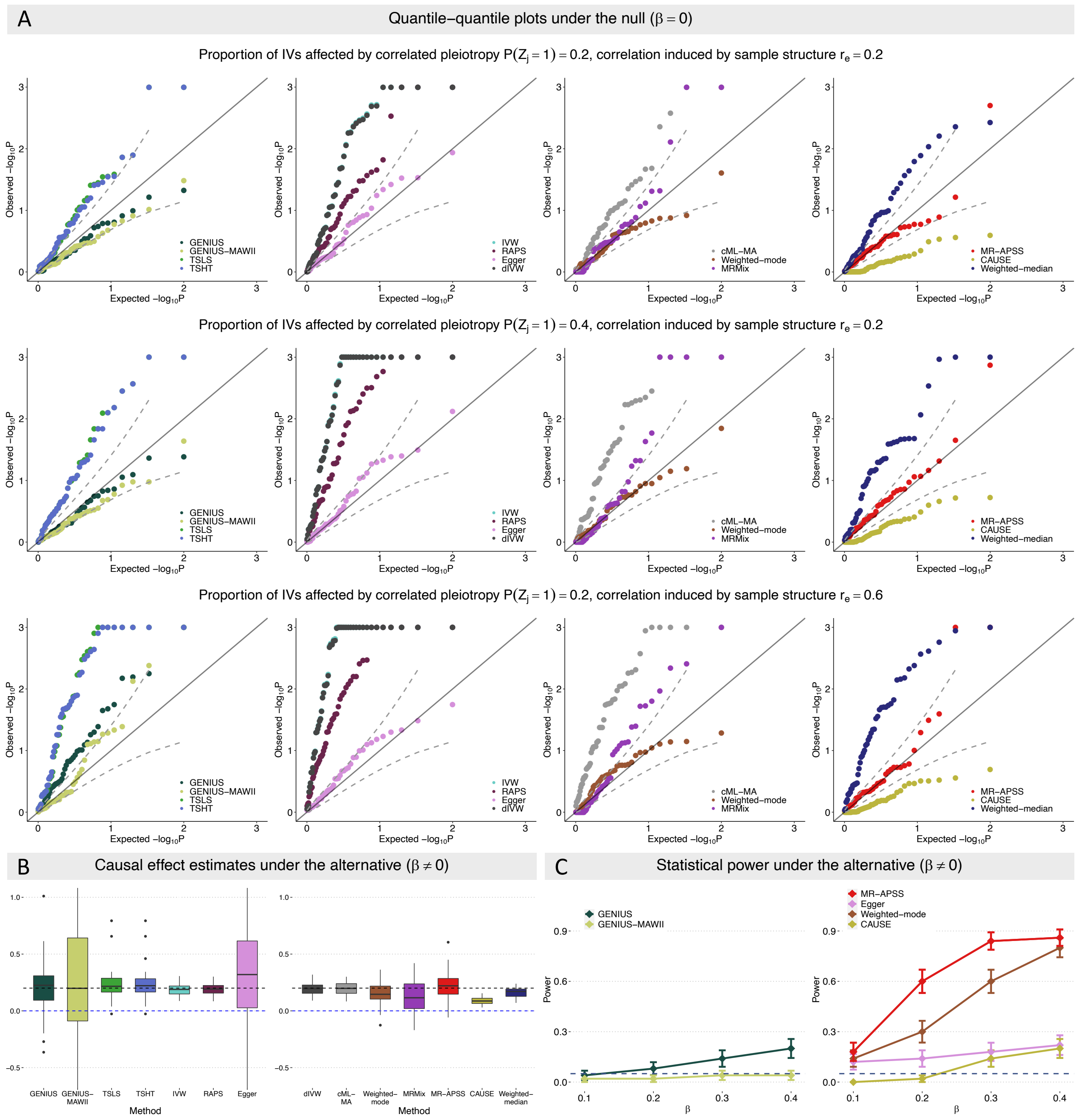

**Figure** S6: Comparison of 14 MR methods on simulated data based on the CAUSE model. (A) Quantile-quantile plots of $-\log_{10}(p)$-values under null simulations in settings including (i) $q = 0.2, r_e = 0.2$, (ii) $q = 0.4, r_e = 0.2$, (iii) $q = 0.2, r_e = 0.6$. (B) Estimates of causal effect under alternative simulations with $\beta = 0.2$. (C) Power under causal effect size $\beta$ varied from 0.1 to 0.4. The comparison of power was conducted among those methods whose type I errors are under controlled in the null simulations.

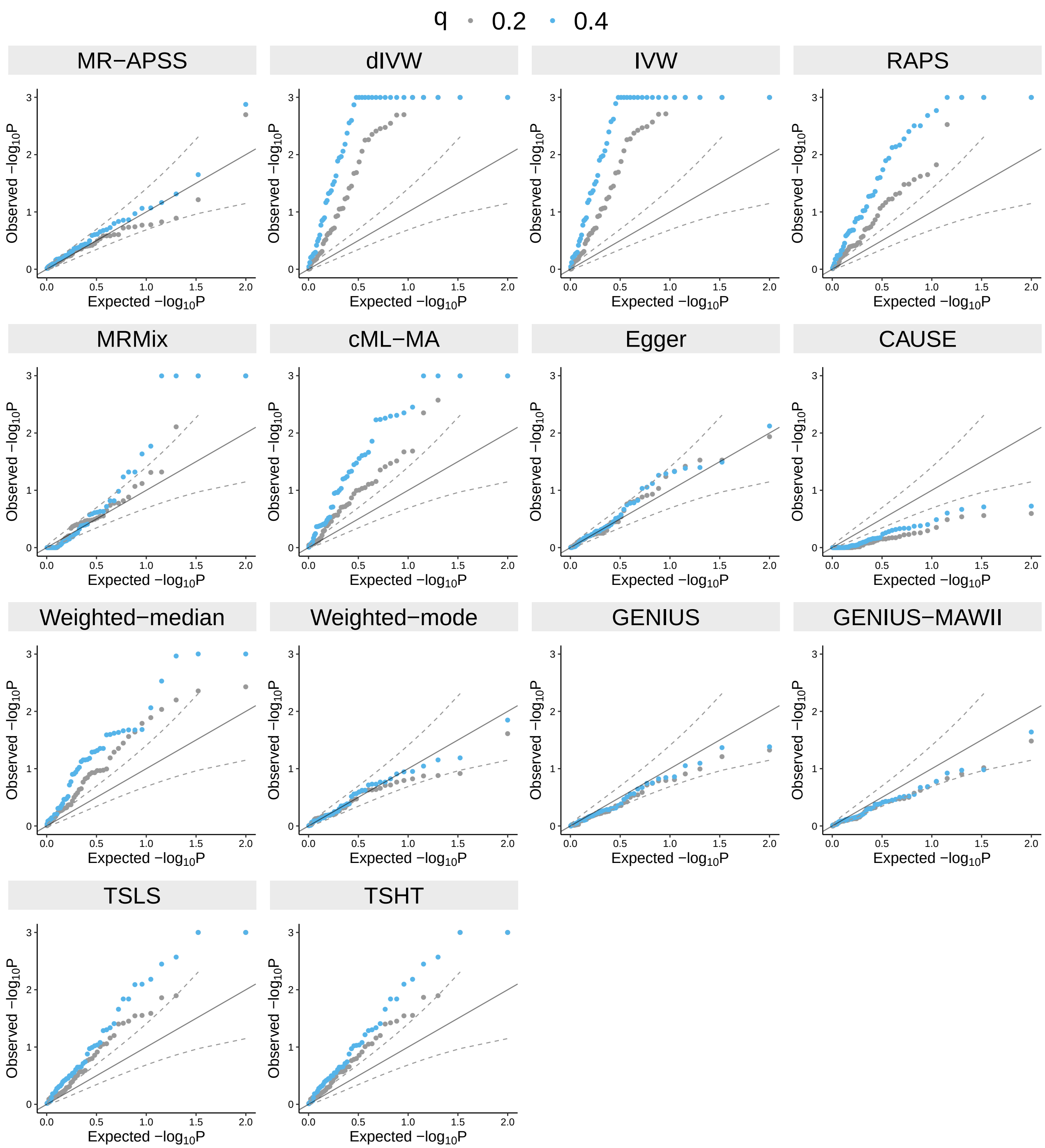

**Figure** S7: Type I error control of 14 MR methods on inferring causal effects in the presence of correlated pleiotropy under CAUSE model. Quantile-quantile plots of $-\log_{10}(p)$-values under null simulations ($\beta = 0$) with correlated pleiotropic effect $\eta = 0.2$. We varied the proportion of SNPs affected by correlated pleiotropy to be $q \in \{0.2, 0.4\}$. The results were summarized from 50 replications.

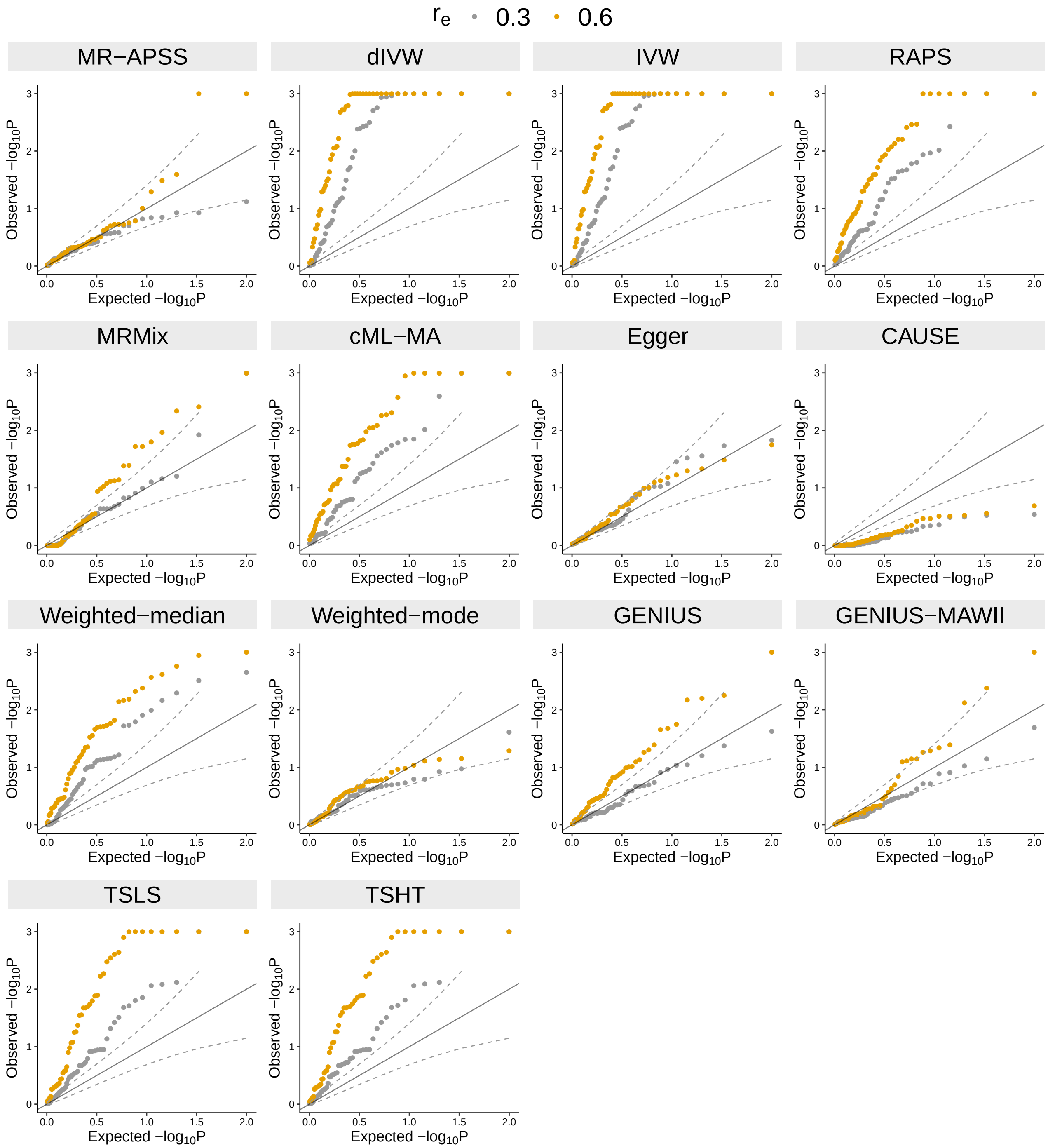

**Figure** S8: The type I error control of 14 MR methods on inferring causal effects under CAUSE model in the presence of sample structure under CAUSE model. Quantile-quantile plots of $-\log_{10}(p)$-values under null simulations ($\beta = 0$) with correlation between estimation errors $c_{12} \in \{0.075, 0.15\}$. The correlation in estimation errors was induced by 10,000 overlapped samples with correlation of environmental noises $r_e \in \{0.3, 0.6\}$. The results were summarized from 50 replications.

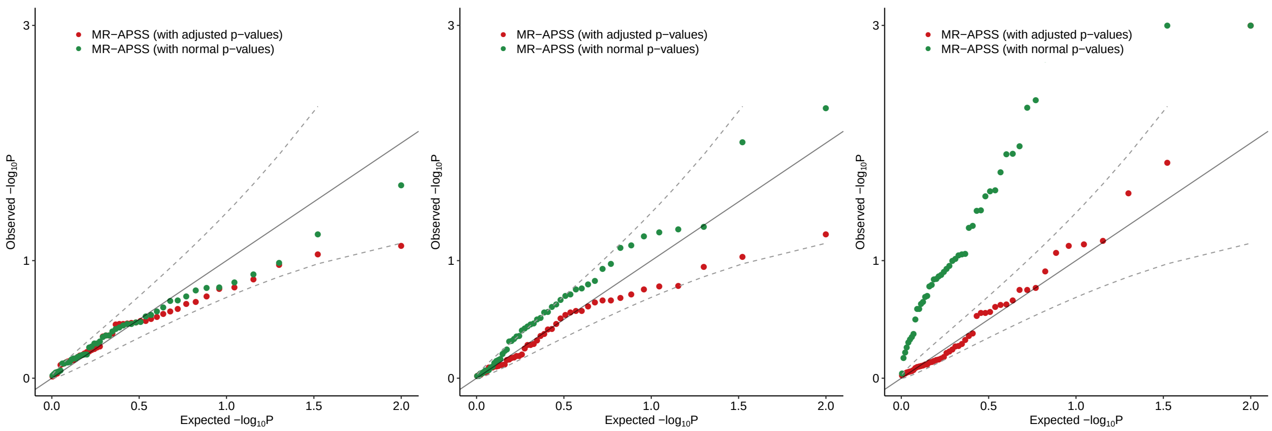

**Figure** S9: Type I error control of MR-APSS with/without adjustment for selection bias due to LD clumping. Quantile-quantile plots of $-\log_{10}(p)$-values under null simulations based on the MR-APSS model with genetic correlation $r_g = 0.1$ and with correlation in estimation errors $c_{12} = 0.075$. The correlation in estimation errors was induced by 10,000 overlapped samples with the correlation of environmental noises $r_e = 0.3$. MR-APSS with adjustment for selection bias arising from LD clumping is denoted as MR-APSS (with adjusted $p$-values). MR-APSS without adjustment for selection bias arising from LD clumping was denoted as MR-APSS (with normal $p$-values). We examined the type I error control of MR-APSS (with adjusted $p$-values) and MR-APSS (with normal $p$-values) with varying IV thresholds: $5 \times 10^{-5}$**(Left)**, $5 \times 10^{-4}$ **(Middle)**, $5 \times 10^{-3}$**(Right)**. The results were summarized from 50 replications.

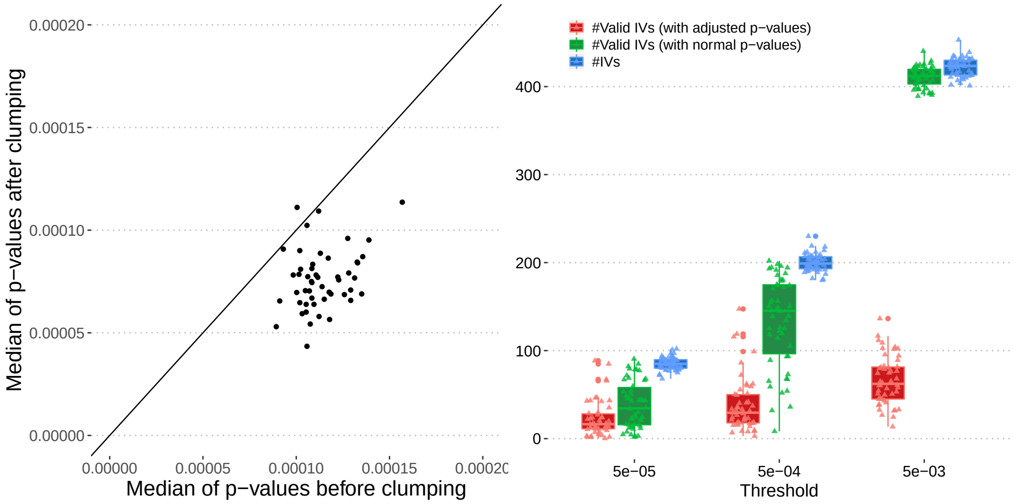

**Figure** S10: Comparison of causal inference by MR-APSS with/without accounting for selection bias arising from LD clumping. We performed null simulations based on the MR-APSS model with genetic correlation $r_g = 0.1$ and with correlation in estimation errors $c_{12} = 0.075$. The correlation in estimation errors was induced by 10,000 overlapped samples with correlation of environmental noises $r_e = 0.3$. **(Left panel)** Comparison of the median of selected IVs' $p$-values before / after LD clumping. As expected, the median of IVs' $p$-values after clumping were generally smaller than that of before clumping. This is because the default LD clumping procedure is designed to keep the independent SNPs with the most significant $p$-values; **(Right panel)** Boxplots comparing the number of selected IVs and the estimated number of valid IVs ($\hat{\pi}_t M_t$) which carry the foreground signals detected by MR-APSS. We examined the impact of the $p$-value adjustment for selection bias arising from LD clumping on the detection of valid IVs. To be specific, we compared the number of valid IVs detected by MR-APSS with the $p$-value adjustment and MR-APSS without the adjustment. The IV threshold was varied from $5 \times 10^{-5}$, $5 \times 10^{-4}$ to $5 \times 10^{-3}$. The results were summarized from 50 replications.

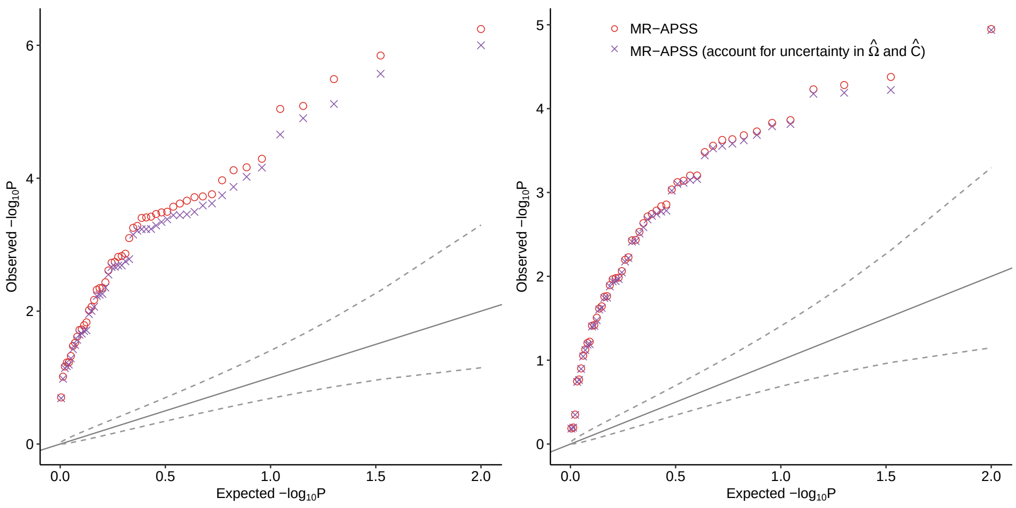

**Figure** S11: The influence of the estimation uncertainty in $\hat{\mathbf{\Omega}}$ and $\hat{\mathbf{C}}$ of the background model on MR-APSS. Quantile-quantile plots of $-\log_{10}(p)$-values under alternative simulations ($\beta = 0.2$) based on MR-APSS model with genetic correlation $r_g = 0.1$ and with correlation in estimation errors $c_{12} = 0075$. We compared MR-APSS and MR-APSS (accounting for uncertainty in $\hat{\mathbf{\Omega}}$ and $\hat{\mathbf{C}}$) with varying IV thresholds $5 \times 10^{-4}$ **(Left)**, and $5 \times 10^{-6}$ **(Right)**.

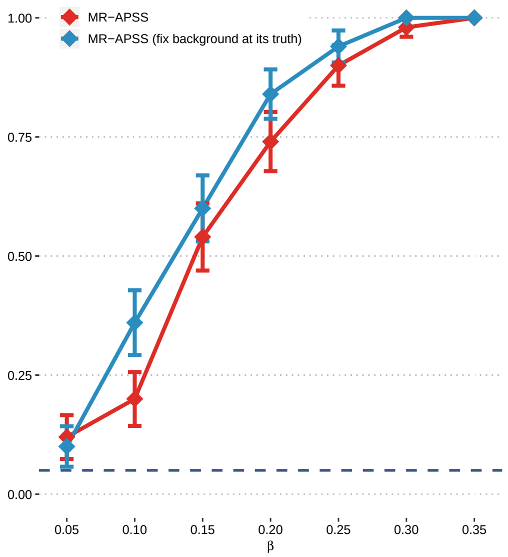

**Figure** S12: The influence of overestimation of $\boldsymbol{\Omega}$ on the power of MR-APSS. Power under alternative simulations based on the MR-APSS model with genetic correlation $r_g = 0.1$ and with correlation in estimation errors $c_{12} = 0.075$. Causal effect $\beta$ was varied from 0.05 to 0.35. We manually fixed the background components $\hat{\boldsymbol{\Omega}}$ and $\hat{\mathbf{C}}$ at their ground truth, denoted as MR-APSS (fix background at its truth), and compared its power to the power of MR-APSS. The results were summarized from 50 replications.

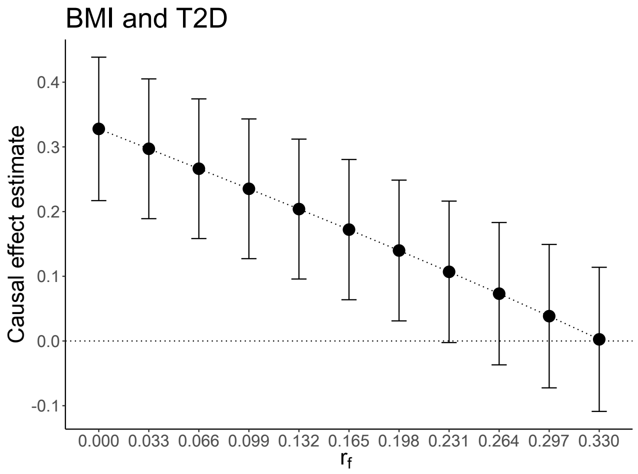

**Figure** S13: Sensitivity analysis result for BMI and T2D. Dots represent the causal effect estimates ($y$-axis) when changing $\gamma_f$ ($x$-axis), i.e., the correlation between IV strength ($\gamma_j$) and direct effect ($\alpha_j$) in the foreground model. Error bars represent their 95% confidence intervals.

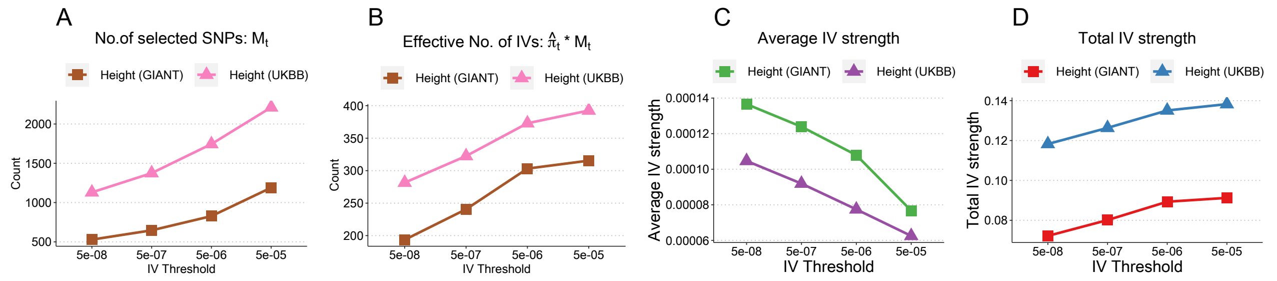

**Figure** S14: Two illustrative examples of exposures for measures of IV strength, including Height (GIANT) and Height (UKBB). (**A**) The number of selected IVs at different IV thresholds, (**B**) The estimated number of valid IVs ($\hat{\pi}_t M_t$) at different IV thresholds. (**C**) and (**D**) The estimated average and total IV strengths.

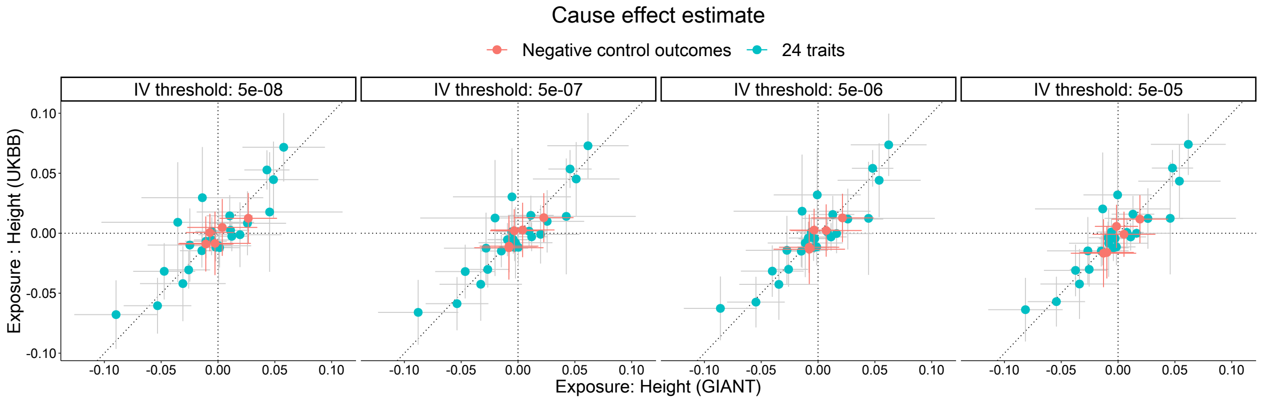

**Figure** S15: Evaluation of the causal effect estimates from MR-APSS when using Height (GIANT) ($x$-axis) and Height (UKBB) ($y$-axis) as exposures at different IV thresholds. The red dots represent the estimated causal effects ($\hat{\beta}$) of the two exposures on the five negative control outcomes, and the cyan dots represent the estimated causal effects ($\hat{\beta}$) on the 24 complex traits. The bars represent the 95% confidence intervals.

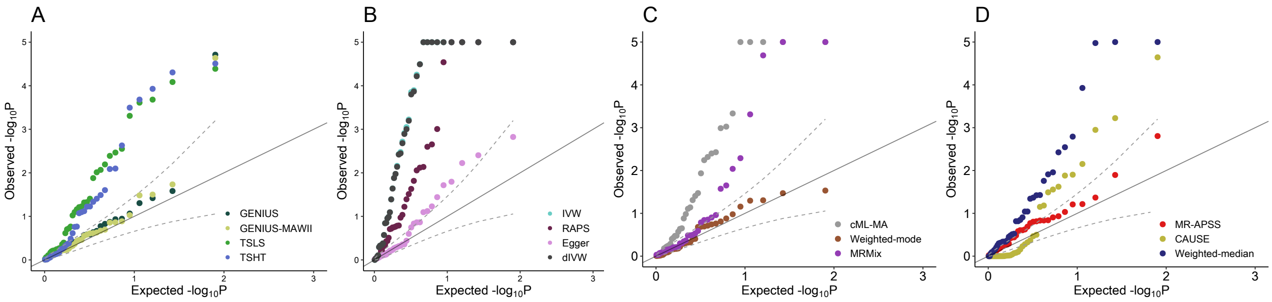

**Figure** S16: Quantile-quantile plots of $-\log_{10}(p)$-values for causal inference between eight complex traits and five negative control outcomes from fourteen MR methods, including four individual-level methods (**A**), ten summary-level methods (**B-D**).

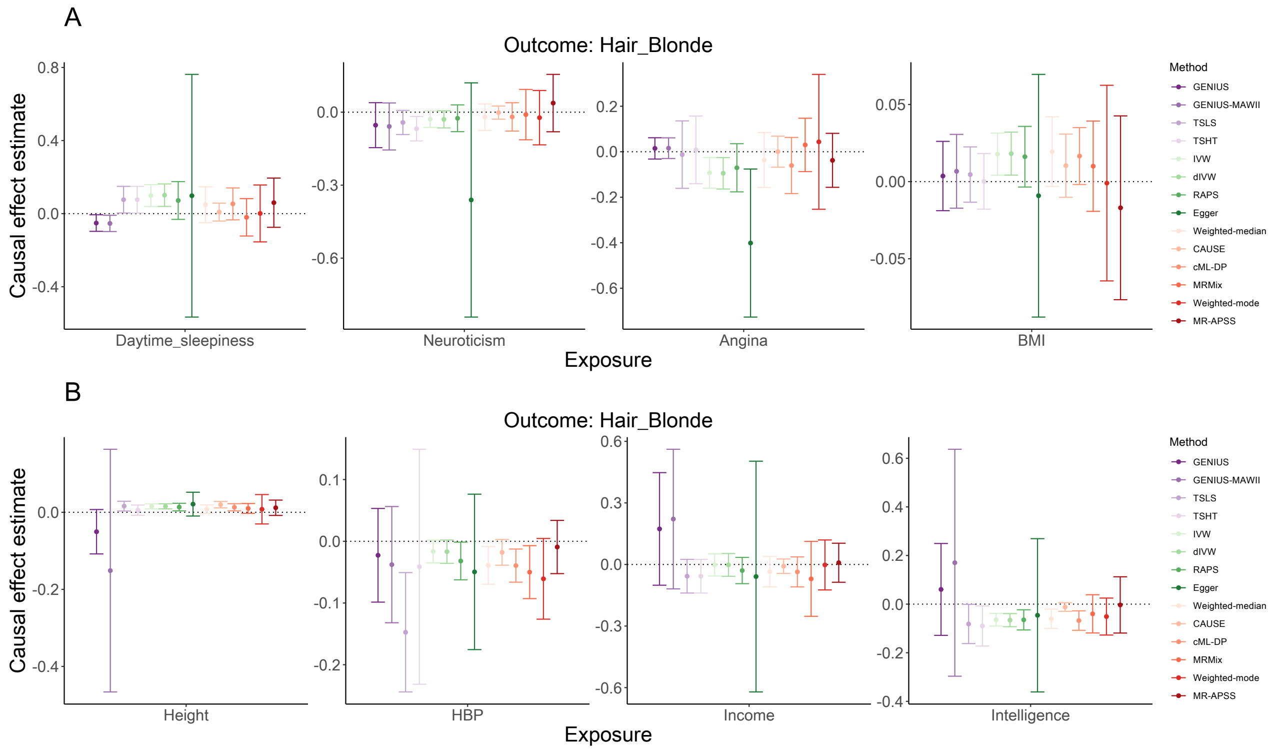

**Figure** S17: Causal effect estimates and their 95% confidence intervals from different MR methods between eight exposures and one negative control outcome (Hair colour: blonde).

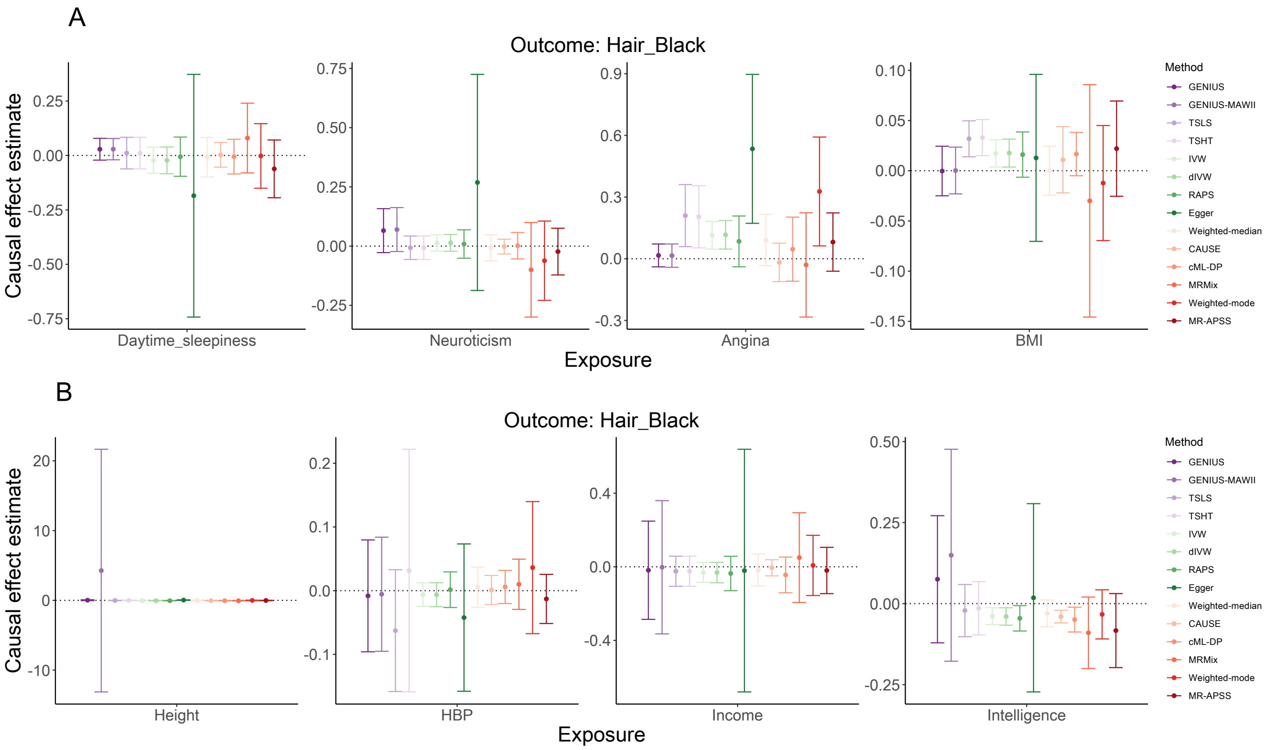

**Figure** S18: Causal effect estimates and their 95% confidence intervals from different MR methods between eight exposures and one negative control outcome (Hair colour: black).

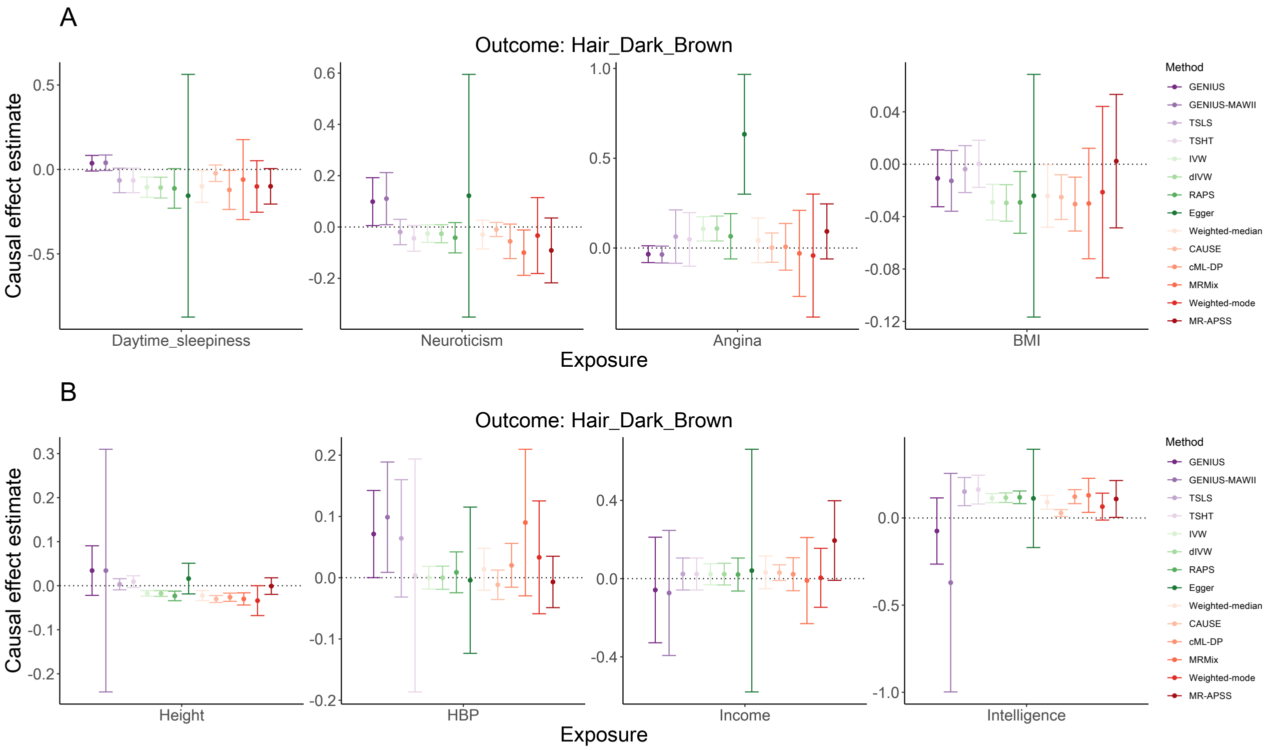

**Figure** S19: Causal effect estimates and their 95% confidence intervals from different MR methods between eight exposures and one negative control outcome (Hair colour: dark brown).

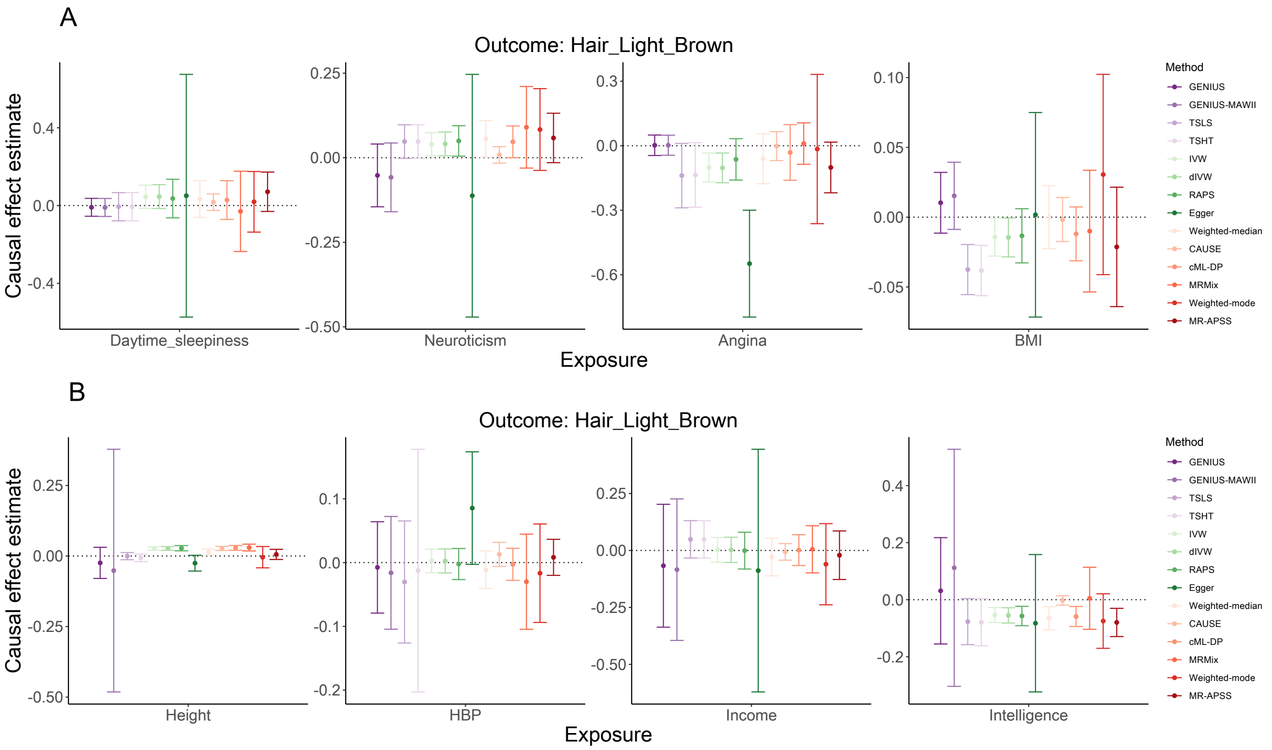

**Figure** S20: Causal effect estimates and their 95% confidence intervals from different MR methods between eight exposures and one negative control outcome (Hair colour: light brown).

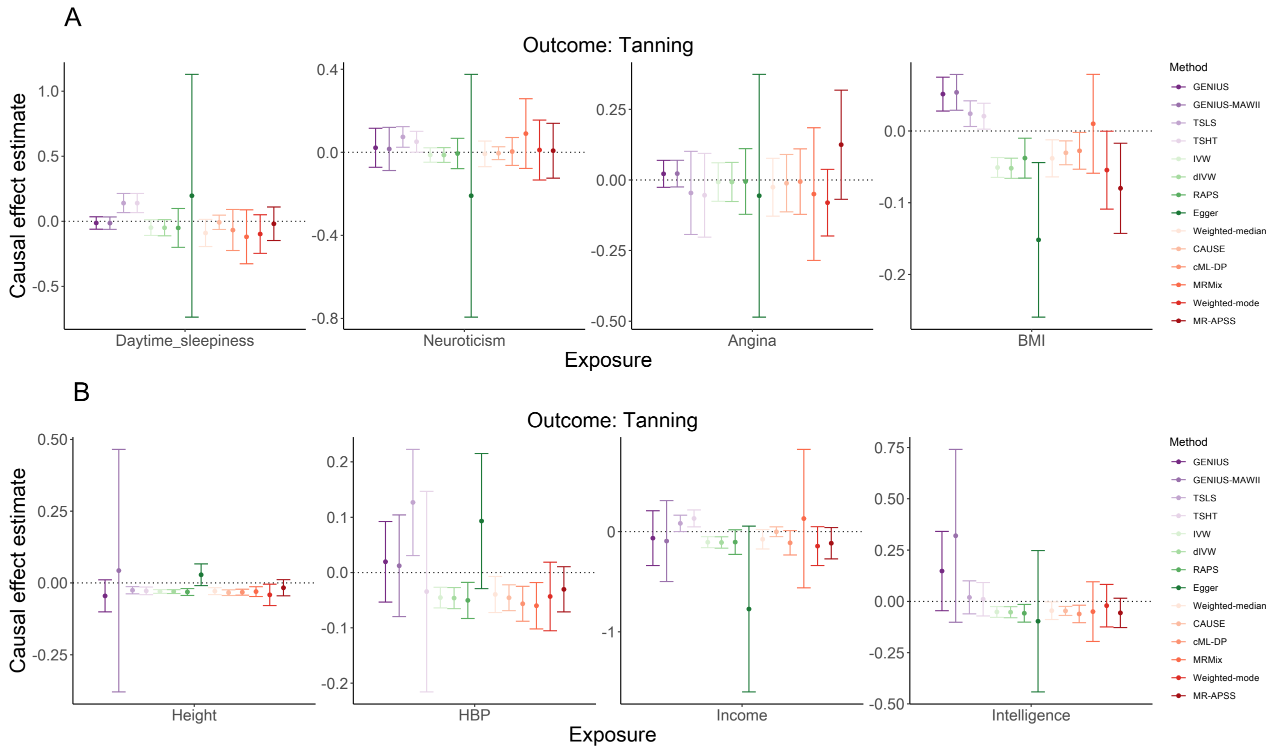

**Figure** S21: Causal effect estimates and their 95% confidence intervals from different MR methods between eight exposures and one negative control outcome (Tanning).

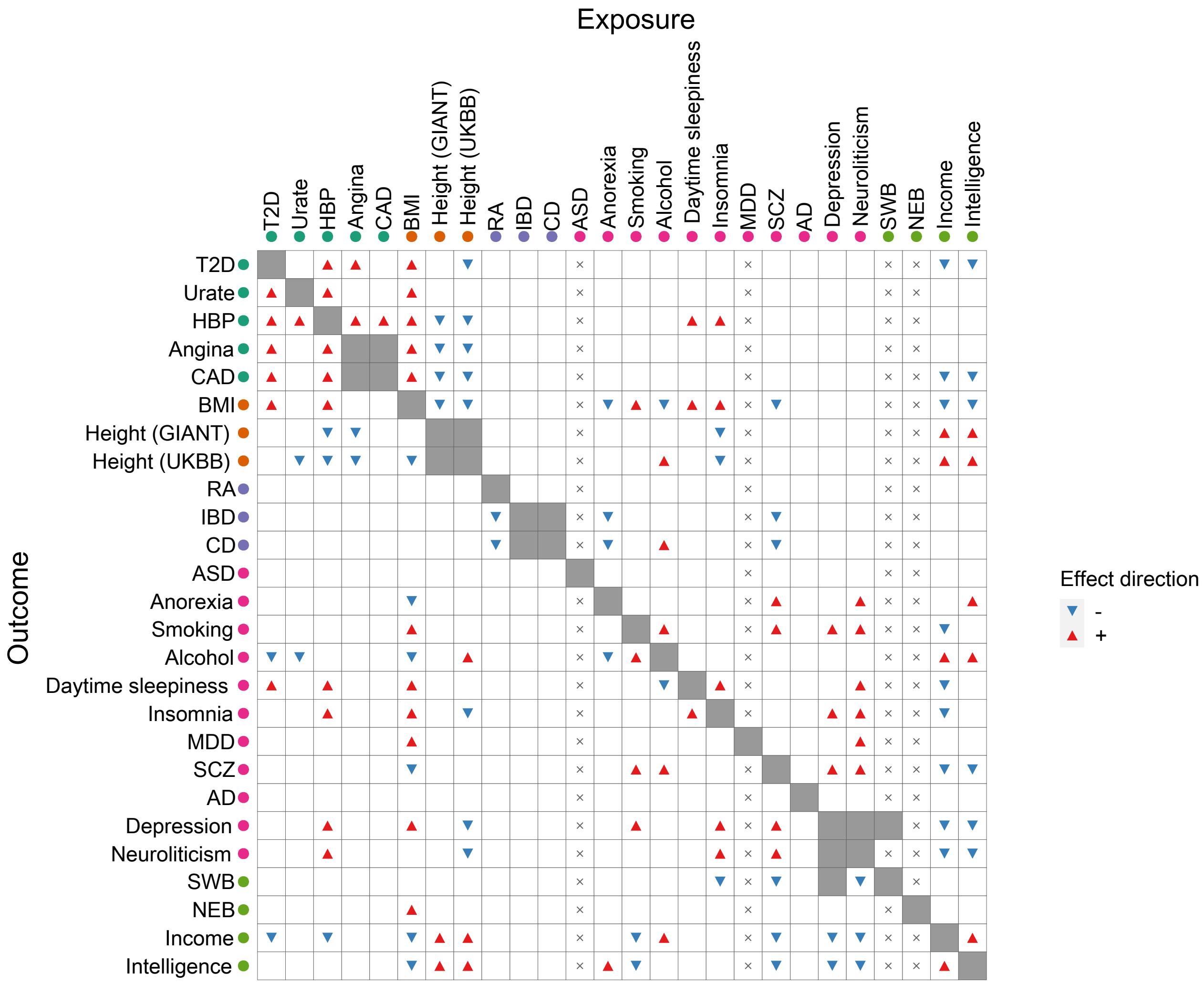

**Figure** S22: Causal relationships between 26 complex traits detected by IVW. The positive and negative estimates of causal effects of the exposure on the outcome are indicated by red up-pointing triangles and blue down-pointing triangles, respectively. Cells marked with $\times$ are trait pairs excluded in MR analysis due to insufficient number of IVs ($< 4$). Non-diagonal cells shaded with grey color are those with genetic correlation large than 0.75.

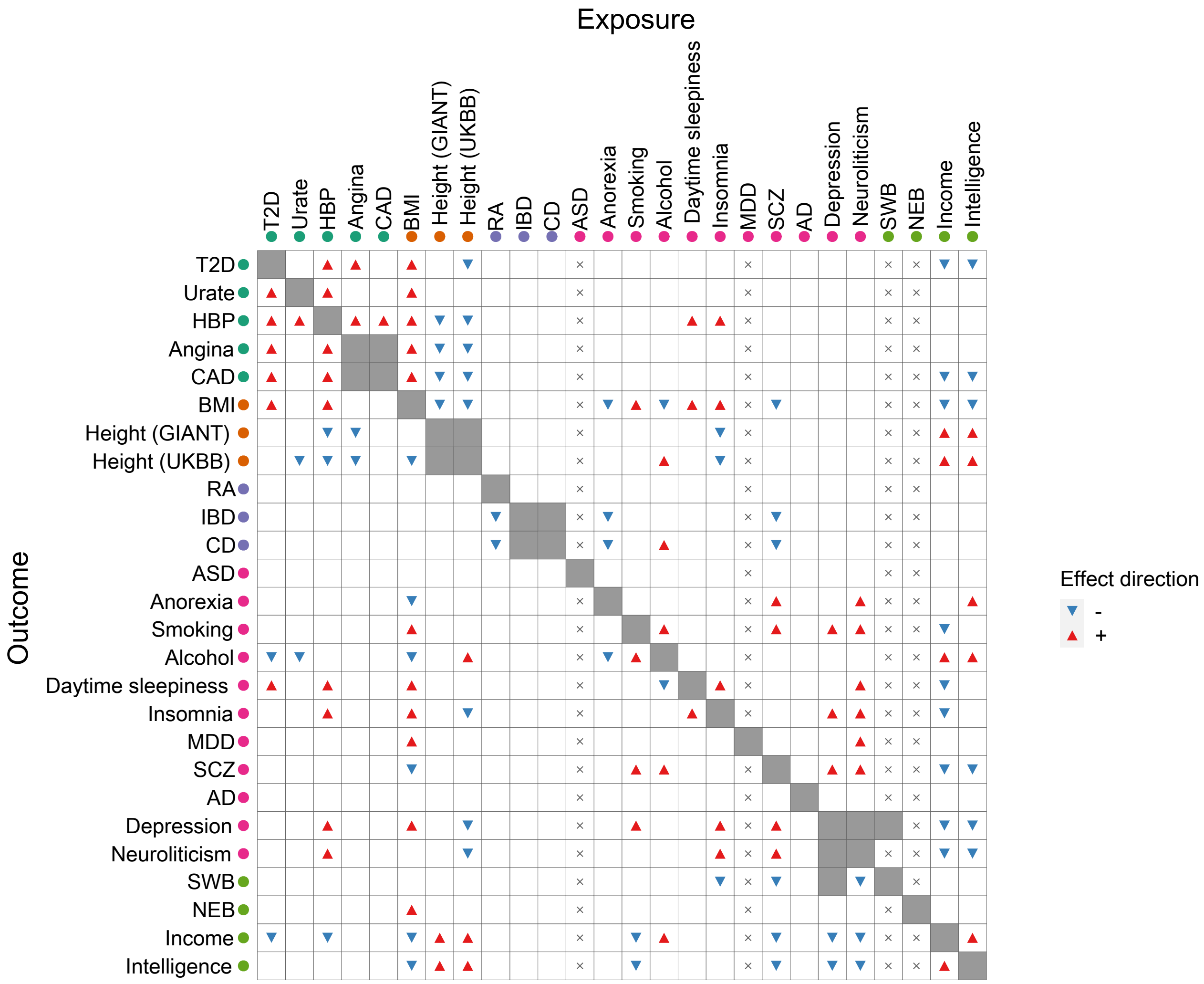

**Figure** S23: Causal relationships between 26 complex traits detected by dIVW. The positive and negative estimates of causal effects of the exposure on the outcome are indicated by red up-pointing triangles and blue down-pointing triangles, respectively. Cells marked with × are trait pairs excluded in MR analysis due to insufficient number of IVs (< 4). Non-diagonal cells shaded with grey color are those with genetic correlation large than 0.75.

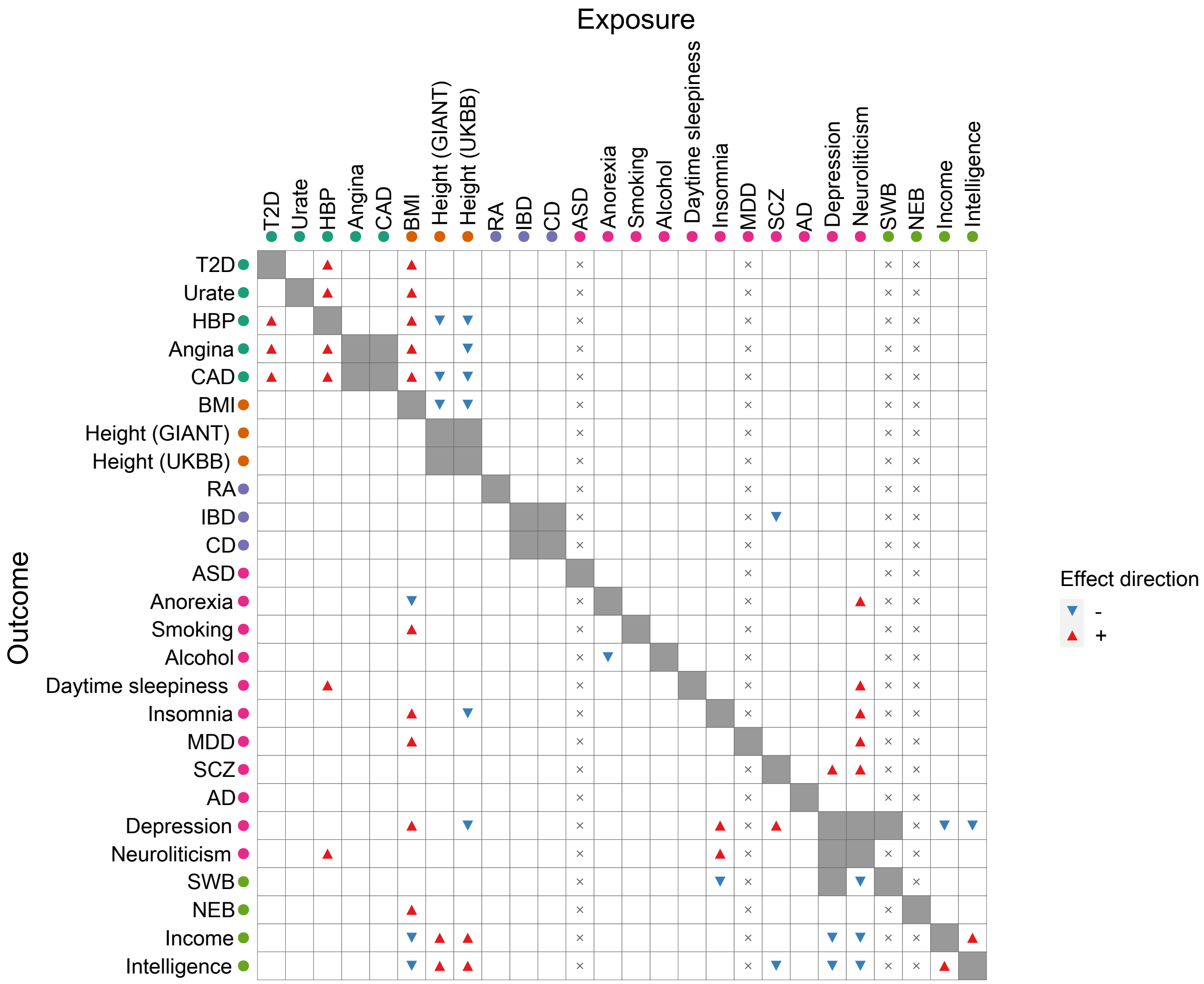

**Figure** S24: Causal relationships between 26 complex traits detected by RAPS. The positive and negative estimates of causal effects of the exposure on the outcome are indicated by red up-pointing triangles and blue down-pointing triangles, respectively. Cells marked with × are trait pairs excluded in MR analysis due to insufficient number of IVs ($< 4$). Non-diagonal cells shaded with grey color are those with genetic correlation large than 0.75.

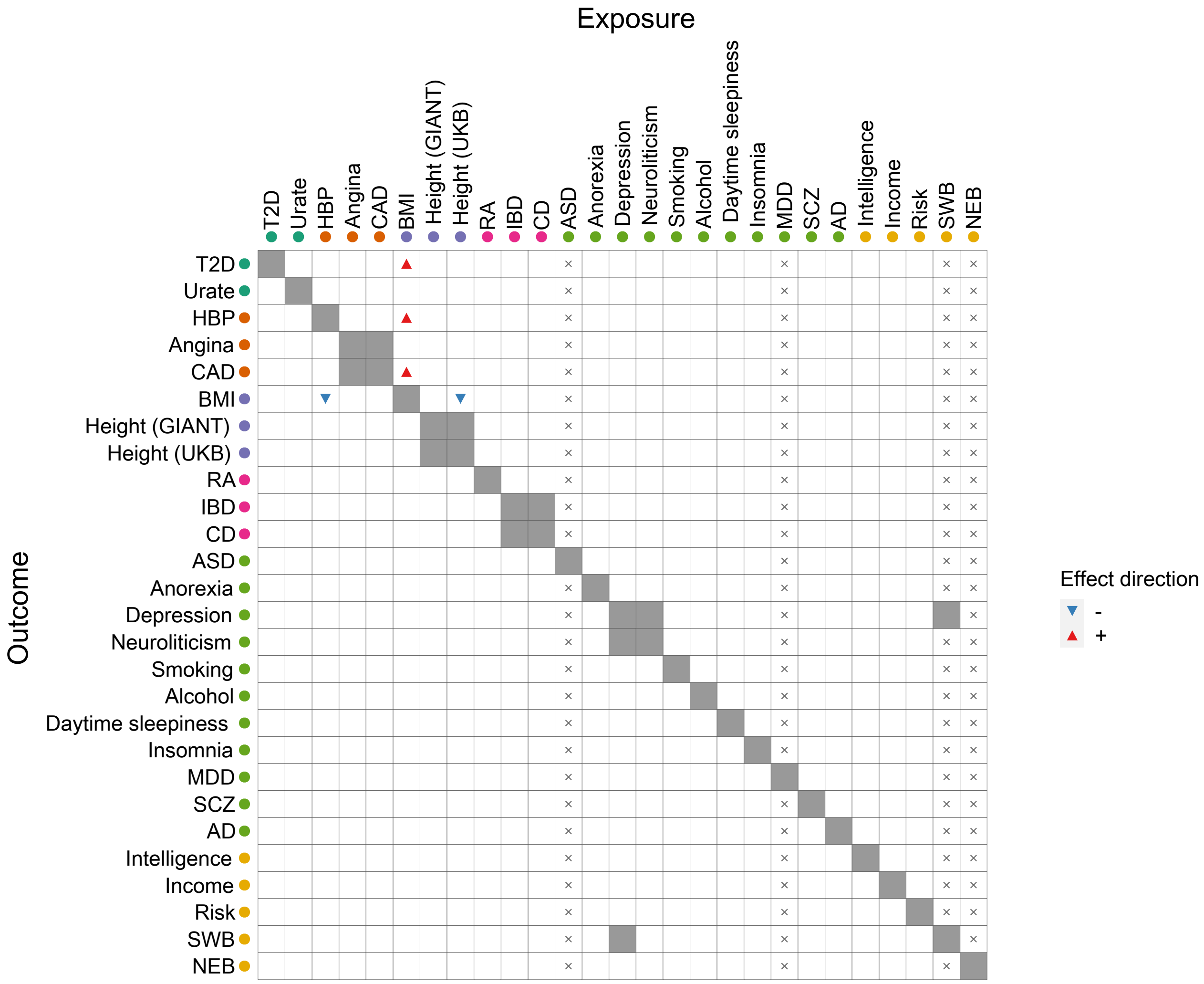

**Figure** S25: Causal relationships between 26 complex traits detected by Egger. The positive and negative estimates of causal effects of the exposure on the outcome are indicated by red up-pointing triangles and blue down-pointing triangles, respectively. Cells marked with × are trait pairs excluded in MR analysis due to insufficient number of IVs ($<4$). Non-diagonal cells shaded with grey color are those with genetic correlation large than 0.75.

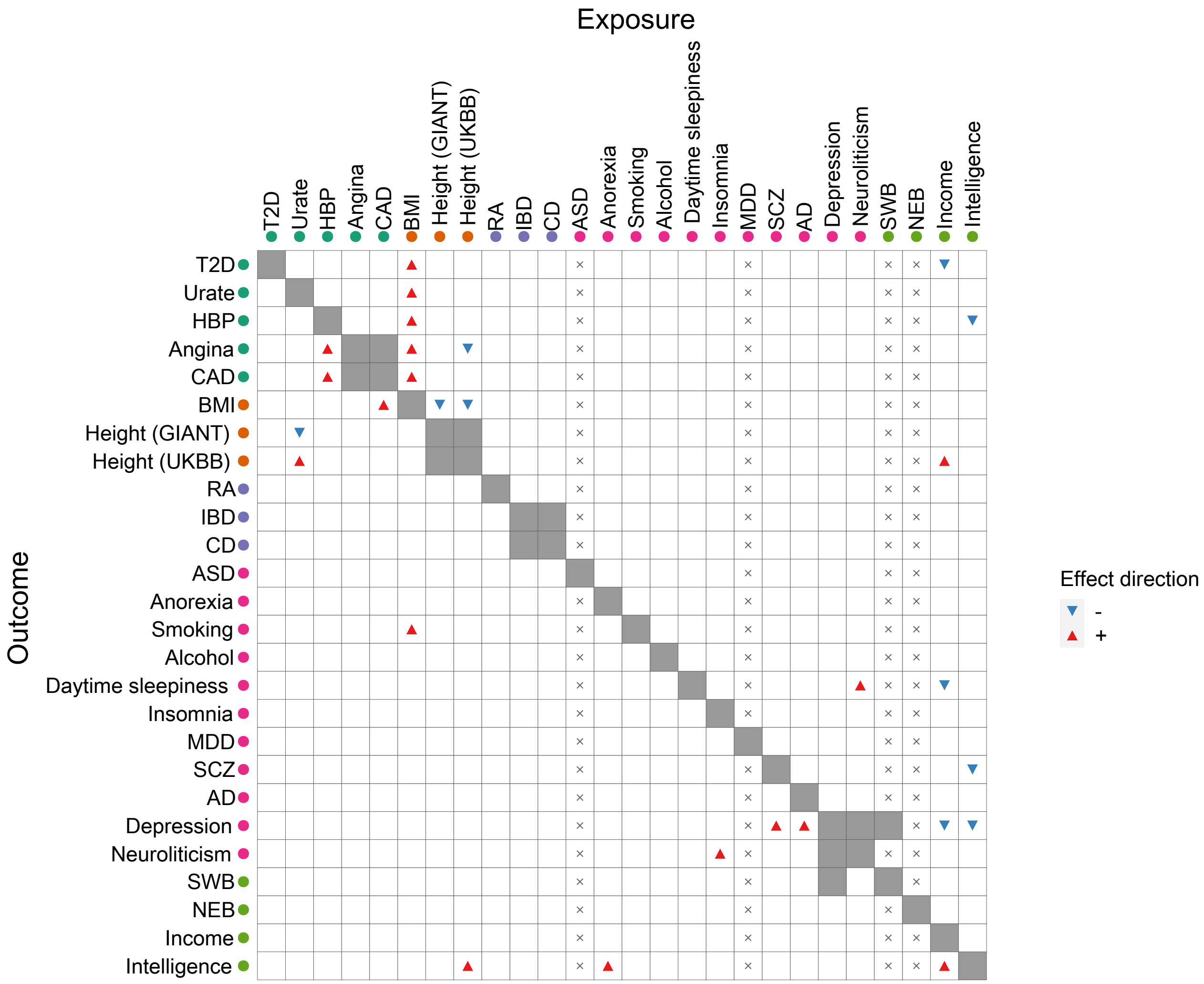

**Figure** S26: Causal relationships between 26 complex traits detected by MRMix. The positive and negative estimates of causal effects of the exposure on the outcome are indicated by red up-pointing triangles and blue down-pointing triangles, respectively. Cells marked with × are trait pairs excluded in MR analysis due to insufficient number of IVs ($< 4$). Non-diagonal cells shaded with grey color are those with genetic correlation large than 0.75.

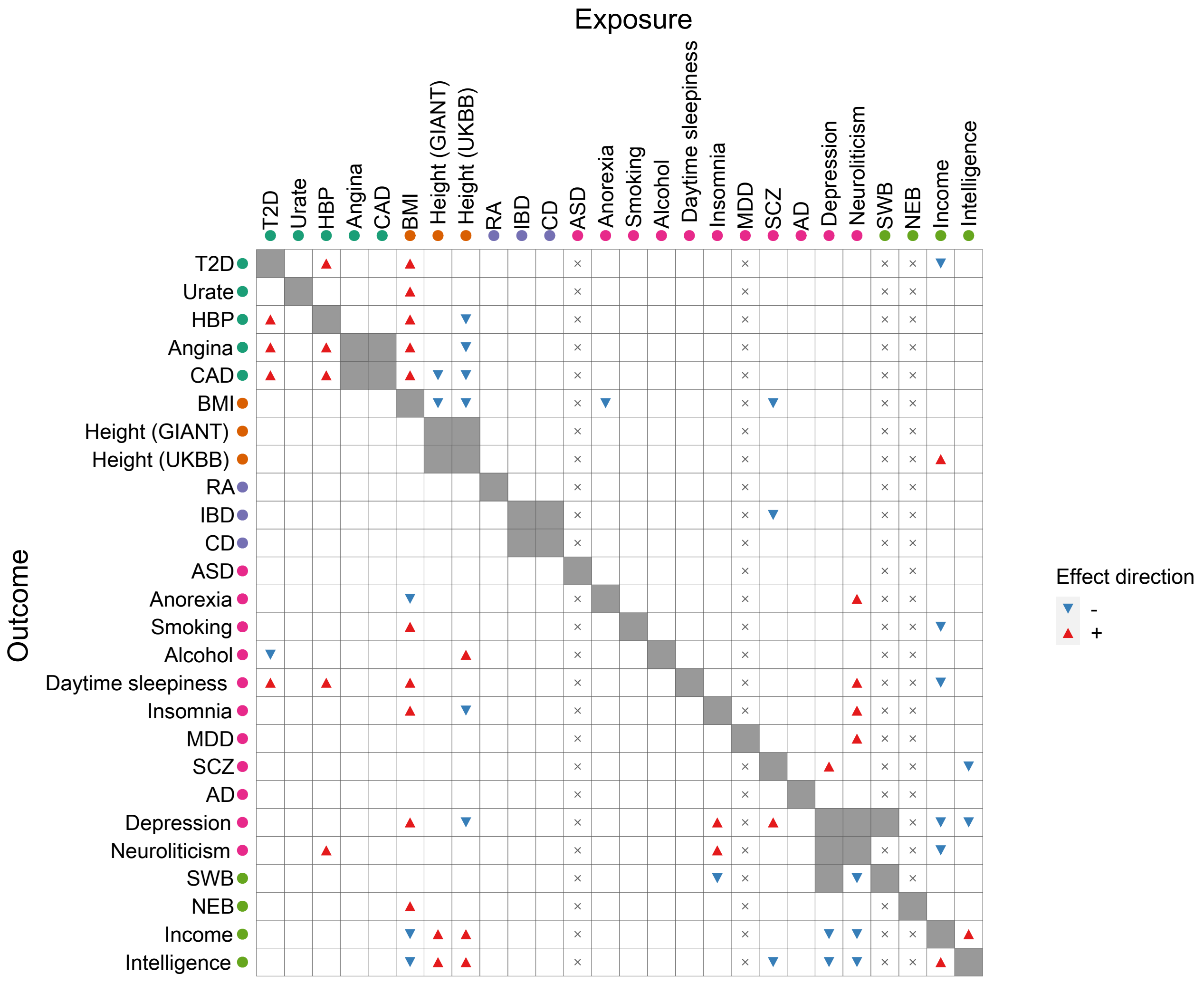

**Figure** S27: Causal relationships between 26 complex traits detected by cML-MA. The positive and negative estimates of causal effects of the exposure on the outcome are indicated by red up-pointing triangles and blue down-pointing triangles, respectively. Cells marked with × are trait pairs excluded in MR analysis due to insufficient number of IVs ($< 4$). Non-diagonal cells shaded with grey color are those with genetic correlation large than 0.75.

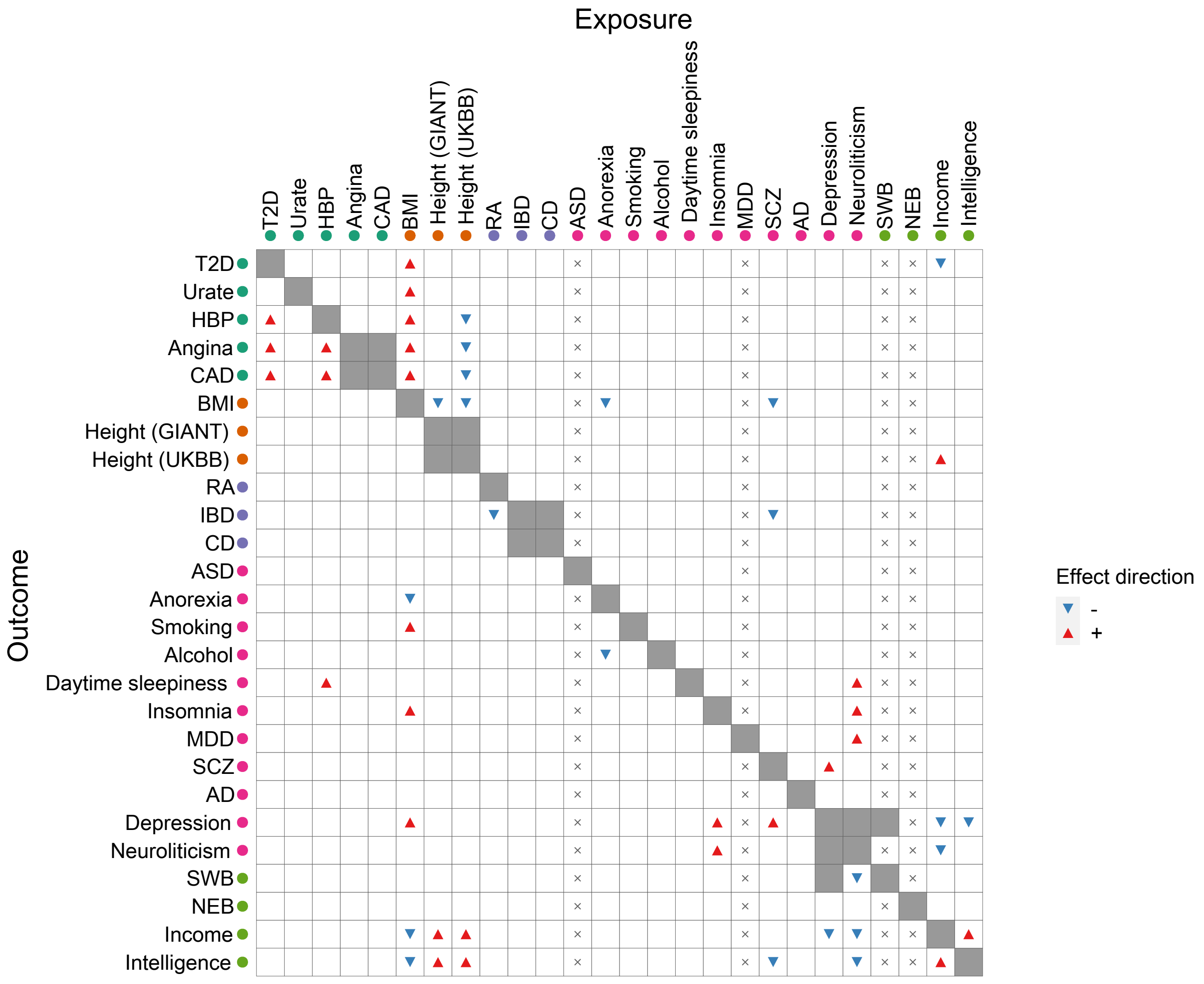

**Figure** S28: Causal relationships between 26 complex traits detected by Weighted-median. The positive and negative estimates of causal effects of the exposure on the outcome are indicated by red up-pointing triangles and blue down-pointing triangles, respectively. Cells marked with × are trait pairs excluded in MR analysis due to insufficient number of IVs ($< 4$). Non-diagonal cells shaded with grey color are those with genetic correlation large than 0.75.

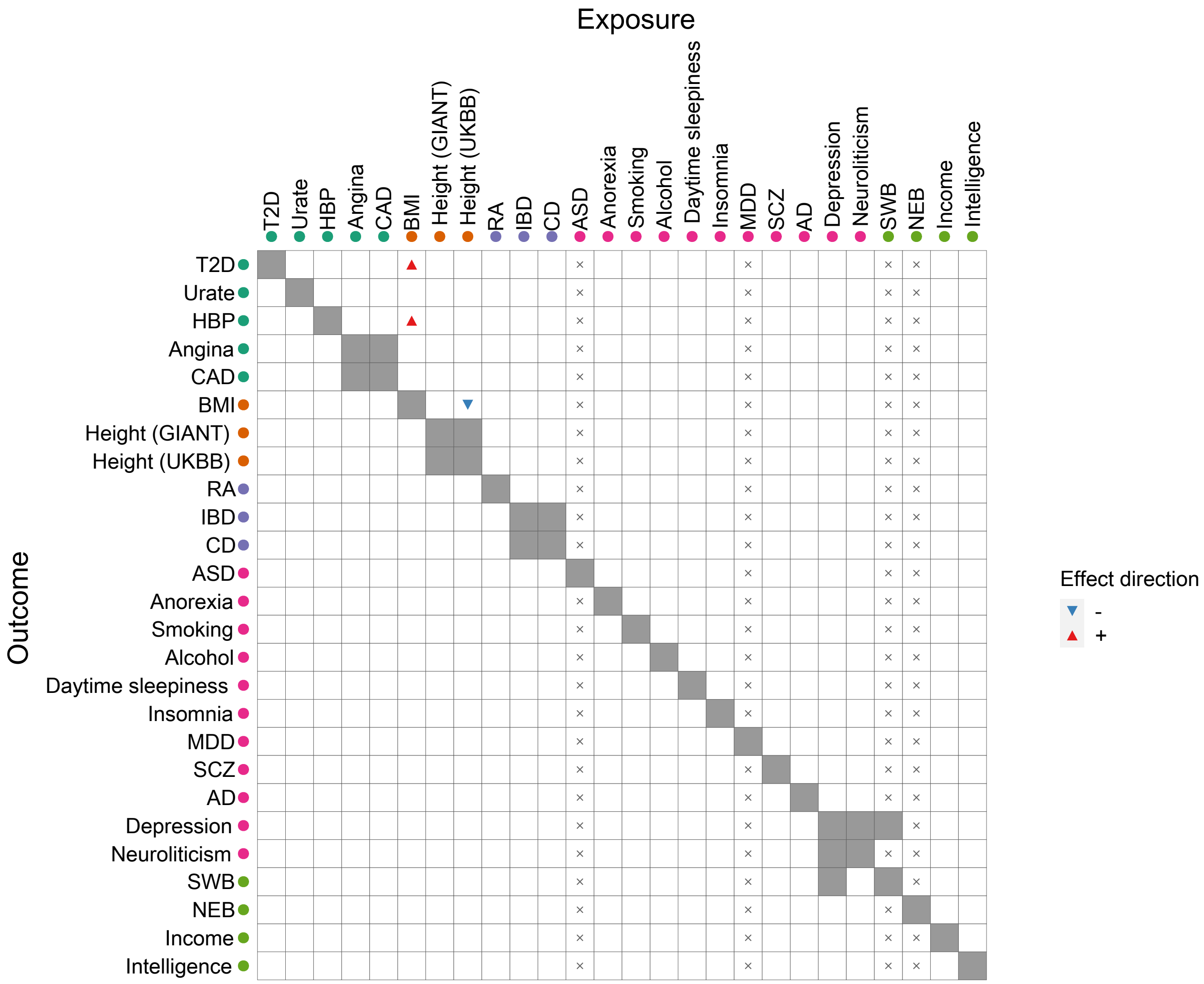

**Figure** S29: Causal relationships between 26 complex traits detected by Weighted-mode. The positive and negative estimates of causal effects of the exposure on the outcome are indicated by red up-pointing triangles and blue down-pointing triangles, respectively. Cells marked with $\times$ are trait pairs excluded in MR analysis due to insufficient number of IVs ($< 4$). Non-diagonal cells shaded with grey color are those with genetic correlation large than 0.75.

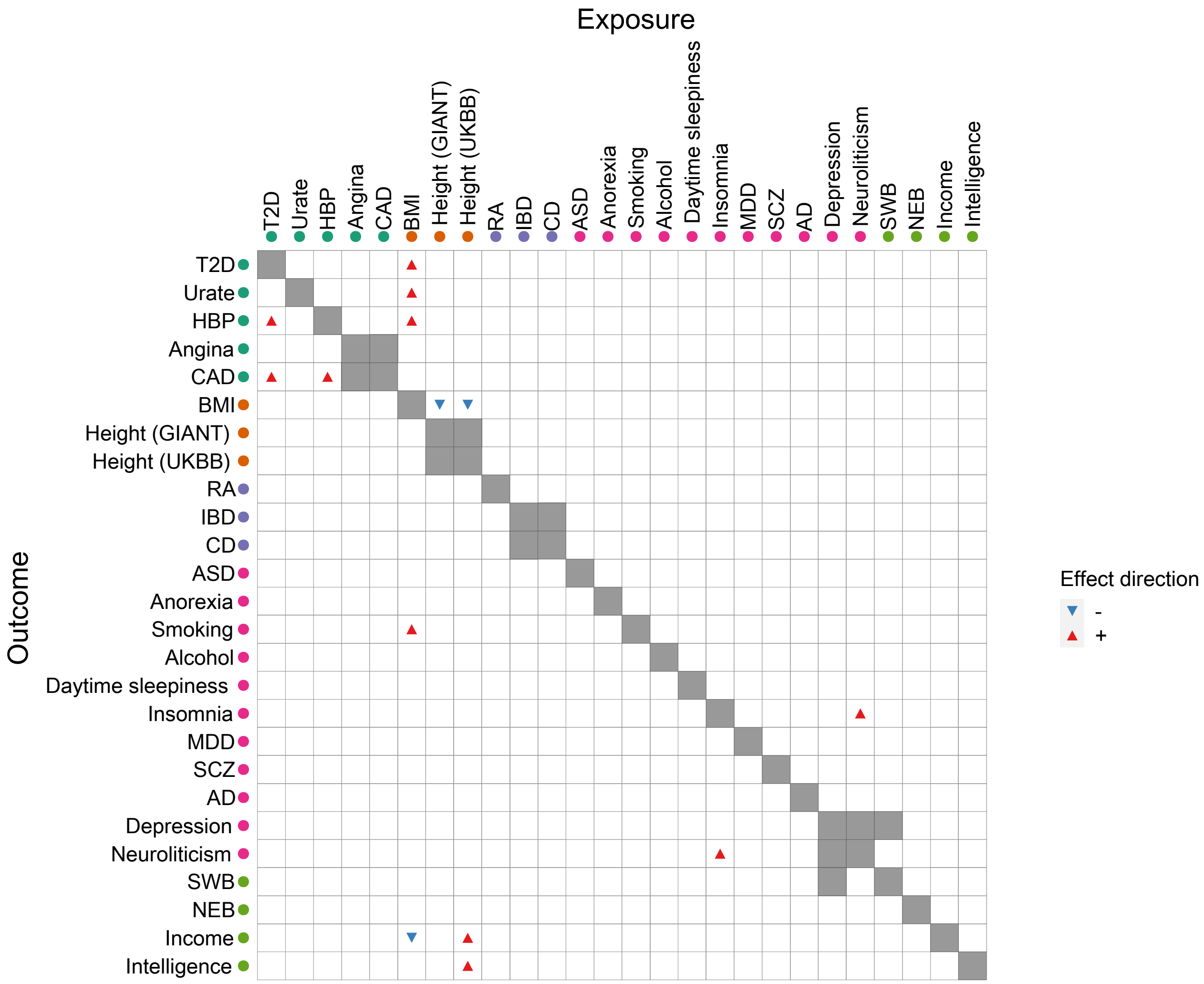

**Figure** S30: Causal relationships between 26 complex traits detected by CAUSE. The positive and negative estimates of causal effects of the exposure on the outcome are indicated by red up-pointing triangles and blue down-pointing triangles, respectively. Non-diagonal cells shaded with grey color are those with genetic correlation large than 0.75.

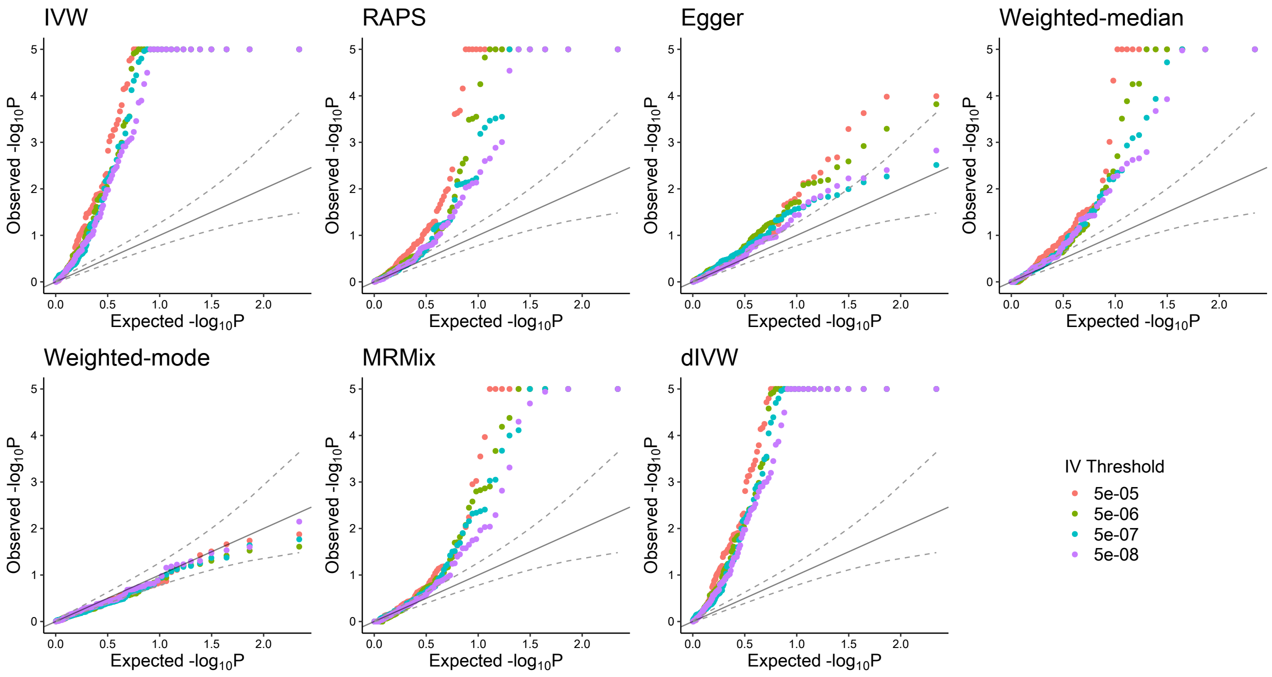

**Figure** S31: Quantile-quantile plots of $-\log(p)$-values produced by several summary-level MR methods for trait pairs between 26 complex traits and five negative control outcomes. We varied the IV threshold at $5 \times 10^{-5}$, $5 \times 10^{-6}$, $5 \times 10^{-7}$ and $5 \times 10^{-8}$ to test their performance.

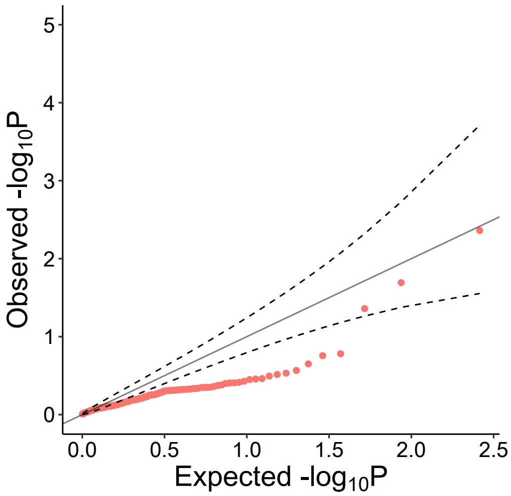

**Figure** S32: Quantile-quantile plots of $-\log 10(p)$-values from LCV for the test of partial causality between 26 complex traits and five negative control outcomes.

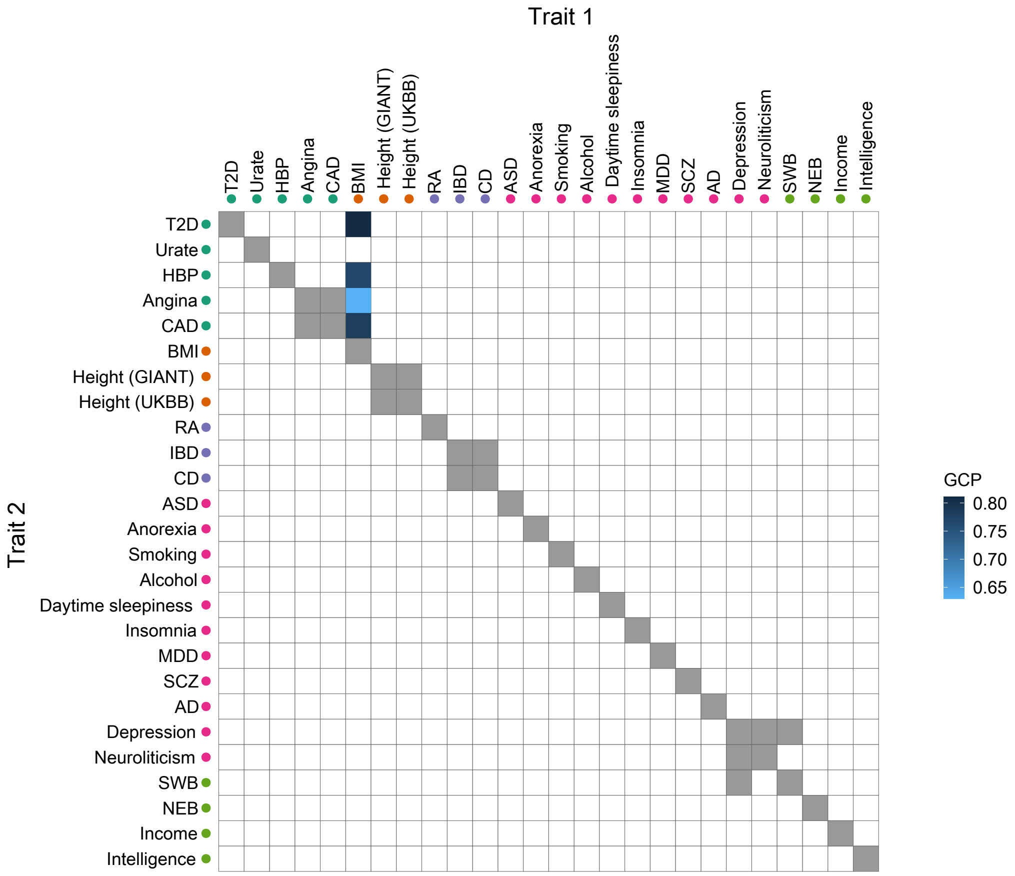

**Figure** S33: Partially or fully genetically causal relationships among 26 complex traits based on LCV. The blue shaded squares indicate significant partially or fully causal effect of trait 1 on trait 2 based on Bonferroni correction.

**Table** S1: Summary of individual-level MR methods.

| **Method** | **Linearity** | **(A-II)** | **(A-III)** | **Key assumptions** |
|---|---|---|---|---|
| TSLS | ✓ | ✓ | ✓ | All IVs are valid. |
| LIML [11] | ✓ | ✓ | ✓ | All IVs are valid. |
| MBTSLS [19] | ✓ | ✓ | × | InSIDE. |
| sisVIVE [18] | ✓ | × | × | Majority valid. |
| Adaptive lasso [34] | ✓ | × | × | Majority valid. |
| TSHT [15] | ✓ | × | × | Plurality valid. |
| GENIUS [31] | × | × | × | All IVs can be invalid.<br>Heteroscedasticity of the exposure. |
| GENIUS-MAWII [37] | × | × | × | All IVs can be invalid.<br>Heteroscedasticity of the exposure. |
| MR-MiSTERI [21] | × | × | × | All IVs can be invalid.<br>Heteroscedasticity of the outcome. |

IV: Instrumental Variable; Three IV assumptions: **(A-I)** IVs are associated with the exposure; **(A-II)** IVs are independent of confounders; and **(A-III)** IVs only affect the outcome through the exposure.

**Table** S2: GWAS sources

| Trait | Group | Description | N | Year | PMID |
|---|---|---|---|---|---|
| Tanning | Negative control outcome | Ease of skin tanning | 378,364 | 2019 | 31427789 |
| Hair: black | Negative control outcome | Hair colour (natural, before greying): Black | 385,603 | 2019 | 31427789 |
| Hair: blonde | Negative control outcome | Hair colour (natural, before greying): Blonde | 385,603 | 2019 | 31427789 |
| Hair: dark brown | Negative control outcome | Hair colour (natural, before greying): Dark brown | 385,603 | 2019 | 31427789 |
| Hair: light brown | Negative control outcome | Hair colour (natural, before greying): Light brown | 385,603 | 2019 | 31427789 |
| Height (GIANT) | Anthropometric | Standing Height | 253,288 | 2014 | 25282103 |
| BMI | Anthropometric | Body Mass Index | 385,336 | 2019 | 31427789 |
| Height (UKB) | Anthropometric | Standing Height | 385,748 | 2019 | 31427789 |
| CAD | Cardiovascular | Coronary Artery disease | 184,305 | 2015 | 26343387 |
| Angina | Cardiovascular | Non-cancer illness code, self-reported: angina | 289,307 | 2019 | 31427789 |
| HBP | Cardiovascular | High blood pressure | 385,699 | 2019 | 31427789 |
| CD | Immune | Crohn disease | 40,266 | 2018 | 28067908 |
| RA | Immune | rheumatoid arthritis | 58,284 | 2014 | 24390342 |
| IBD | Immune | Inflammatory Bowel Disease | 59,957 | 2017 | 28067908 |
| Urate | Metabolic | Serum Urate | 110,347 | 2012 | 23263486 |
| T2D | Metabolic | Type II diabetes | 898,130 | 2018 | 30297969 |
| ASD | Neurological/Psychiatric | Autism Spectrum disorser | 15,954 | 2017 | 28540026 |
| AD | Neurological/Psychiatric | Late-onset Alzheimer's Disease | 54,162 | 2013 | 24162737 |
| Anorexia | Neurological/Psychiatric | Anorexia Nervosa | 72,517 | 2019 | 31308545 |
| SCZ | Neurological/Psychiatric | Schizophrenia | 105,318 | 2018 | 29483656 |
| Intelligence | Neurological/Psychiatric | Fluid intelligence score | 125,935 | 2019 | 31427789 |
| MDD | Neurological/Psychiatric | Major Depressive Disorder | 244,890 | 2019 | 31427789 |
| Smoking | Neurological/Psychiatric | Ever smoked regularly(no ukb) | 249,171 | 2019 | 30643251 |
| Neuroliticism | Neurological/Psychiatric | Neroliticism | 312,740 | 2019 | 31427789 |
| Depression | Neurological/Psychiatric | Depressive Symptoms | 381,455 | 2018 | 29942085 |
| Alcohol | Neurological/Psychiatric | Drinks per week | 414,343 | 2019 | 30643258 |
| Daytime sleepiness | Neurological/Psychiatric | Daytime sleepiness | 452,071 | 2019 | 31409809 |
| Insomnia | Neurological/Psychiatric | Insomnia | 453,379 | 2019 | 30804566 |
| Income | Social | Income | 286,301 | 2019 | 31844048 |
| NEB | Social | Number of children ever born | 343,072 | 2016 | 27798627 |
| SWB | Social | Subject Well Being | 298,420 | 2016 | 27089181 |

**Table** S3: Analysis results from MR-APSS and RAPS for BMI and Insomnia

| Method | IV Threshold: $5 \times 10^{-5}$ | | | | IV Threshold: $5 \times 10^{-8}$ | | | |
|---|---|---|---|---|---|---|---|---|
| | # IV (#Valid IV) | $\hat{\beta}$ | s.e.($\hat{\beta}$) | $p$-value | # IV (#Valid IV) | $\hat{\beta}$ | s.e.($\hat{\beta}$) | $p$-value |
| MR-APSS | 1298 (219) | 0.0337 | 0.0221 | 0.128 | 400 (106) | 0.0284 | 0.0274 | 0.298 |
| MR-APSS($\mathbf{\Omega} = \mathbf{0}$) | 1298 (558) | 0.0698 | 0.0134 | 1.70e-07 | 400 (190) | 0.0544 | 0.0197 | 5.71e-03 |
| MR-APSS($\mathbf{C} = \mathbf{I}$) | 1298 (478) | 0.0629 | 0.0162 | 1.00e-04 | 400 (200) | 0.0519 | 0.0208 | 0.012 |
| MR-APSS($\mathbf{\Omega} = \mathbf{0}, \mathbf{C} = \mathbf{I}$) | 1298 (952) | 0.0854 | 0.0100 | 1.28e-17 | 400 (352) | 0.0739 | 0.0138 | 9.09e-08 |
| RAPS | 1298 | 0.0721 | 0.0074 | 1.54e-22 | 400 | 0.0702 | 0.0118 | 3.04e-09 |

The default IV threshold for MR-APSS is $5 \times 10^{-5}$; the default IV threshold for RAPS is $5 \times 10^{-8}$.

#### **Table** S4: The 36 trait pairs detected by MR-APSS, Egger, CAUSE or Weighted-mode

| Exposure | Outcome | $\hat{r}_g$ | $\hat{c}_{12}$ | $c_1$ | $c_2$ | # IVs: 5e-05 (# valid IVs) | # IVs: 5e-08 | MR-APSS | IVW | RAPS | Weighted-mode | Egger | CAUSE |
|---|---|---|---|---|---|---|---|---|---|---|---|---|---|
| BMI | Angina | 0.31 (0.03) | 0.08 (0.01) | 1.18 | 1.03 | 1301 (205) | 400 | **0.09 (3.03e-07)** | **0.10 (5.83e-39)** | **0.11 (6.06e-30)** | 0.10 (4.03e-03) | 0.08 (0.03) | 0.06 (1.20e-04) |
| BMI | CAD | 0.28 (0.03) | 0.01 (0.01) | 1.18 | 0.9 | 1300 (205) | 399 | **0.15 (5.55e-08)** | **0.14 (6.92e-42)** | **0.14 (1.47e-27)** | 0.14 (0.01) | **0.32 (4.99e-10)** | 0.06 (7.40e-04) |
| BMI | Depression | 0.22 (0.02) | 0.08 (0.01) | 1.18 | 1.02 | 1297 (212) | 399 | **0.07 (2.09e-05)** | **0.08 (1.03e-30)** | **0.08 (5.79e-16)** | 0.05 (0.09) | -0.01 (0.88) | 0.04 (6.61e-04) |
| BMI | HBP | 0.35 (0.02) | 0.29 (0.01) | 1.18 | 1.13 | 1301 (206) | 400 | **0.18 (2.76e-07)** | **0.24 (3.05e-255)** | **0.26 (2.63e-87)** | **0.25 (1.67e-17)** | **0.23 (2.53e-05)** | **0.12 (2.06e-13)** |
| BMI | Income | -0.26 (0.02) | -0.08 (0.01) | 1.18 | 1.05 | 1300 (222) | 400 | **−0.17 (1.83e-11)** | **−0.15 (7.87e-73)** | **−0.14 (4.02e-26)** | -0.14 (9.66e-05) | **−0.20 (7.42e-05)** | **−0.07 (4.19e-05)** |
| BMI | Smoking | 0.26 (0.02) | 0.02 (0.01) | 1.18 | 0.98 | 1284 (215) | 399 | **0.11 (1.36e-06)** | **0.11 (3.43e-37)** | **0.11 (1.45e-17)** | 0.06 (0.28) | 0.12 (0.01) | **0.06 (2.35e-05)** |
| BMI | T2D | 0.55 (0.02) | 0.14 (0.01) | 1.18 | 1.12 | 1296 (190) | 399 | **0.33 (6.77e-09)** | **0.42 (0.00e+00)** | **0.47 (6.06e-165)** | **0.47 (2.09e-26)** | **0.50 (4.16e-10)** | **0.15 (2.11e-12)** |
| BMI | Urate | 0.35 (0.03) | 0.02 (0.01) | 1.18 | 0.91 | 1278 (200) | 390 | 0.12 (0.15) | **0.20 (4.47e-50)** | **0.21 (1.26e-34)** | 0.15 (0.15) | 0.26 (1.82e-04) | **0.10 (6.74e-06)** |
| Depression | Insomnia | 0.45 (0.03) | 0.18 (0.01) | 1.02 | 1.03 | 197 (70) | 7 | **0.57 (4.38e-05)** | **0.39 (3.52e-12)** | 0.38 (1.97e-03) | 0.18 (0.09) | -1.16 (0.31) | 0.15 (4.50e-03) |
| HBP | Angina | 0.47 (0.04) | 0.08 (0.01) | 1.12 | 1.03 | 684 (189) | 197 | **0.15 (2.19e-13)** | **0.15 (3.87e-44)** | **0.14 (1.49e-22)** | 0.06 (0.22) | 0.09 (0.10) | 0.10 (9.39e-05) |
| HBP* | BMI | 0.35 (0.02) | 0.29 (0.01) | 1.13 | 1.18 | 683 (225) | 196 | 0.03 (0.40) | **0.10 (2.48e-24)** | 0.10 (1.14e-04) | 0.20 (0.20) | **−0.40 (2.80e-05)** | 0.10 (4.38e-04) |
| HBP | CAD | 0.46 (0.03) | 0.02 (0.01) | 1.12 | 0.9 | 683 (192) | 196 | **0.32 (4.92e-22)** | **0.28 (2.82e-96)** | **0.28 (4.89e-37)** | -0.06 (0.20) | 0.16 (0.06) | **0.15 (1.82e-08)** |
| HBP* | Urate | 0.30 (0.03) | 0.01 (0.01) | 1.12 | 0.9 | 677 (185) | 193 | 0.07 (0.07) | **0.12 (9.27e-12)** | **0.12 (3.74e-05)** | 0.79 (0.79) | **−0.43 (6.28e-05)** | 0.07 (8.60e-03) |
| Height (GIANT) | BMI | -0.11 (0.02) | -0.01 (0.01) | 1.35 | 1.19 | 1188 (320) | 527 | **−0.08 (1.08e-06)** | **−0.07 (6.65e-66)** | **−0.08 (1.83e-17)** | 0.19 (1.05e-03) | -0.13 (4.39e-04) | **−0.05 (2.24e-06)** |
| Height (GIANT) | CAD | -0.09 (0.02) | -0.04 (0.01) | 1.35 | 0.89 | 1200 (319) | 532 | **−0.05 (2.51e-05)** | **−0.05 (4.62e-17)** | **−0.05 (1.07e-09)** | 0.01 (0.80) | -0.02 (0.42) | -0.03 (2.14e-03) |
| Height (GIANT) | Income | 0.17 (0.02) | 0.02 (0.01) | 1.34 | 1.05 | 1201 (320) | 534 | **0.05 (2.35e-07)** | **0.05 (3.66e-22)** | **0.05 (1.45e-11)** | 0.04 (0.21) | 0.05 (0.06) | 0.03 (2.14e-04) |
| Height (UKBB) | Angina | -0.19 (0.03) | -0.07 (0.01) | 1.97 | 1.03 | 2227 (401) | 1136 | **−0.03 (3.99e-05)** | **−0.04 (5.58e-22)** | **−0.04 (1.66e-14)** | -0.04 (0.02) | -0.02 (0.08) | -0.02 (1.49e-03) |
| Height (UKBB) | BMI | -0.13 (0.02) | -0.10 (0.01) | 1.97 | 1.18 | 2226 (405) | 1136 | **−0.06 (2.48e-06)** | **−0.06 (2.01e-85)** | **−0.08 (2.41e-26)** | **−0.11 (1.64e-07)** | **−0.10 (6.78e-06)** | **−0.04 (5.09e-09)** |
| Height (UKBB) | CAD | -0.13 (0.02) | -0.03 (0.01) | 1.97 | 0.9 | 2224 (397) | 1136 | **−0.06 (7.36e-08)** | **−0.05 (3.75e-26)** | **−0.05 (5.95e-14)** | 0.03 (0.40) | -0.06 (2.01e-03) | -0.02 (1.76e-03) |
| Height (UKBB) | Income | 0.21 (0.02) | 0.11 (0.01) | 1.97 | 1.05 | 2226 (395) | 1136 | **0.05 (1.62e-12)** | **0.06 (3.85e-59)** | **0.06 (1.67e-31)** | 0.06 (5.60e-03) | 0.04 (0.01) | **0.04 (4.39e-07)** |
| Height (UKBB) | Intelligence | 0.16 (0.02) | 0.17 (0.01) | 1.97 | 1.11 | 2227 (393) | 1136 | **0.07 (1.34e-08)** | **0.09 (5.97e-61)** | **0.09 (5.66e-29)** | 0.06 (0.08) | 0.05 (0.03) | **0.06 (4.10e-07)** |
| Income | BMI | -0.26 (0.02) | -0.08 (0.01) | 1.05 | 1.18 | 260 (66) | 25 | **−0.47 (7.29e-05)** | **−0.22 (3.22e-16)** | -0.24 (7.46e-03) | -0.17 (0.37) | 0.22 (0.73) | -0.10 (0.03) |
| Income | Depression | -0.45 (0.03) | -0.11 (0.01) | 1.05 | 1.02 | 259 (73) | 25 | **−0.35 (4.07e-09)** | **−0.27 (1.19e-22)** | **−0.27 (7.17e-14)** | -0.23 (2.02e-03) | -0.44 (0.11) | -0.12 (4.95e-04) |
| Income | Intelligence | 0.58 (0.03) | 0.12 (0.01) | 1.05 | 1.11 | 260 (70) | 25 | **1.07 (1.01e-13)** | **0.79 (1.00e-59)** | **0.78 (2.30e-21)** | 0.53 (7.19e-03) | **2.31 (6.70e-05)** | 0.27 (2.12e-03) |
| Insomnia | Depression | 0.45 (0.03) | 0.18 (0.01) | 1.03 | 1.02 | 348 (80) | 37 | **0.25 (6.90e-05)** | **0.24 (5.88e-17)** | **0.24 (1.04e-09)** | 0.23 (8.99e-03) | 0.19 (0.60) | 0.14 (8.11e-04) |
| Insomnia | Neuroliticism | 0.42 (0.02) | 0.23 (0.01) | 1.04 | 1.05 | 348 (88) | 37 | **0.47 (6.75e-08)** | **0.43 (2.43e-43)** | **0.45 (2.80e-12)** | 0.55 (3.07e-04) | -0.46 (0.38) | **0.22 (4.42e-07)** |
| Intelligence | Income | 0.58 (0.03) | 0.12 (0.01) | 1.11 | 1.05 | 403 (62) | 47 | **0.36 (2.23e-09)** | **0.29 (7.38e-77)** | **0.28 (1.26e-19)** | 0.19 (4.13e-03) | 0.81 (7.13e-04) | 0.08 (1.53e-03) |
| Neuroticism | Anorexia | 0.28 (0.03) | 0.02 (0.01) | 1.04 | 1.02 | 427 (158) | 68 | **−0.40 (4.19e-07)** | **0.24 (1.96e-09)** | **0.25 (5.95e-06)** | 0.15 (0.21) | 0.21 (0.63) | 0.15 (3.75e-03) |
| Neuroticism | Insomnia | 0.42 (0.02) | 0.23 (0.01) | 1.05 | 1.04 | 463 (141) | 69 | **0.29 (2.70e-10)** | **0.27 (1.99e-64)** | **0.29 (8.01e-26)** | 0.23 (3.66e-03) | -0.42 (0.03) | **0.14 (3.89e-06)** |
| Neuroticism | MDD | 0.53 (0.05) | 0.13 (0.01) | 1.05 | 0.99 | 463 (151) | 69 | **0.18 (2.06e-05)** | **0.18 (5.65e-16)** | **0.18 (1.54e-09)** | 0.13 (0.08) | 0.02 (0.94) | 0.10 (0.05) |
| Neuroticism | SCZ | 0.21 (0.02) | 0.01 (0.01) | 1.04 | 1.1 | 457 (123) | 69 | **0.57 (7.02e-07)** | **0.32 (3.63e-21)** | **0.34 (4.65e-05)** | 0.17 (0.18) | -0.56 (0.38) | 0.15 (2.02e-03) |
| Neuroticism | SWB | -0.66 (0.04) | -0.05 (0.01) | 1.04 | 1 | 450 (141) | 68 | **−0.23 (5.05e-09)** | **−0.19 (1.04e-20)** | **−0.19 (1.11e-11)** | -0.02 (0.80) | -0.06 (0.76) | -0.10 (3.47e-03) |
| SCZ | Depression | 0.32 (0.03) | 0.01 (0.01) | 1.1 | 1.01 | 664 (163) | 121 | **0.08 (4.10e-05)** | **0.06 (3.05e-16)** | **0.07 (6.01e-09)** | 0.05 (0.23) | 0.12 (0.18) | 0.04 (8.73e-04) |
| T2D | Angina | 0.37 (0.04) | 0.06 (0.01) | 1.11 | 1.03 | 652 (199) | 187 | **0.06 (3.95e-05)** | **0.06 (1.03e-11)** | **0.06 (1.35e-06)** | 0.09 (0.03) | 0.04 (0.38) | 0.05 (0.03) |
| T2D | CAD | 0.39 (0.03) | 0.03 (0.01) | 1.1 | 0.89 | 660 (206) | 190 | **0.13 (1.43e-10)** | **0.13 (1.56e-36)** | **0.13 (8.48e-18)** | 0.10 (6.39e-03) | 0.05 (0.35) | **0.08 (1.15e-05)** |
| T2D | HBP | 0.43 (0.02) | 0.11 (0.01) | 1.11 | 1.12 | 652 (202) | 187 | **0.10 (8.69e-07)** | **0.13 (2.30e-70)** | **0.13 (1.61e-15)** | 0.06 (0.02) | -0.01 (0.81) | **0.11 (1.01e-07)** |

We also provided the results of IVW, the standardized MR method require all IVs are valid, and the results of RAPS, a method rely on InSIDE assumption in the table for comparison. Trait pairs detected by Egger but not by MR-APSS are marked by *.

### References

[1] Abdel Abdellaoui, David Hugh-Jones, Loic Yengo, Kathryn E. Kemper, and Peter M. Visscher. Genetic correlates of social stratification in great britain. *Nature Human Behaviour*, 3(12), 2019.

[2] Zhigang Bao, Xiucai Ding, Jingming Wang, and Ke Wang. Statistical inference for principal components of spiked covariance matrices. *arXiv preprint arXiv:2008.11903*, 2020.

[3] Emre Barut, Jianqing Fan, and Anneleen Verhasselt. Conditional sure independence screening. *Journal of the American Statistical Association*, 111(515):1266–1277, 2016.

[4] Andrew C. Berry. The Accuracy of the Gaussian Approximation to the Sum of Independent Variates. *Transactions of the American Mathematical Society*, 49(1):122–136, 1941.

[5] Jack Bowden, George Davey Smith, and Stephen Burgess. Mendelian randomization with invalid instruments: effect estimation and bias detection through Egger regression. *International journal of epidemiology*, 44(2):512–525, 2015.

[6] Jack Bowden, George Davey Smith, Philip C Haycock, and Stephen Burgess. Consistent estimation in mendelian randomization with some invalid instruments using a weighted median estimator. *Genetic epidemiology*, 40(4):304–314, 2016.

[7] Brendan K Bulik-Sullivan, Po-Ru Loh, Hilary K Finucane, Stephan Ripke, Jian Yang, Nick Patterson, Mark J Daly, Alkes L Price, and Benjamin M Neale. LD Score regression distinguishes confounding from polygenicity in genome-wide association studies. *Nature genetics*, 47(3):291–295, 2015.

[8] Brendan Buliksullivan, Hilary K Finucane, Verneri Anttila, Alexander Gusev, Felix R Day, Poru Loh, Laramie Duncan, John R B Perry, Nick Patterson, Elise B Robinson, et al. An atlas of genetic correlations across human diseases and traits. *Nature Genetics*, 47(11):1236–1241, 2015.

[9] Esseen Carl-Gustav. On the Liapunoff limit of error in the theory of probability. *Transactions of the American Mathematical Society*, A28:1–19, 1942.

[10] Bei-Hung Chang, Stuart Lipsitz, and Christine Waternaux. Logistic regression in meta-analysis using aggregate data. *Journal of Applied Statistics*, 27(4):411–424, 2000.

[11] John C Chao and Norman R Swanson. Consistent estimation with a large number of weak instruments. *Econometrica*, 73(5):1673–1692, 2005.

[12] Christopher DeBoever, Yosuke Tanigawa, Malene E Lindholm, Greg McInnes, Adam Lavertu, Erik Ingelsson, Chris Chang, Euan A Ashley, Carlos D Bustamante, Mark J Daly, et al. Medical relevance of protein-truncating variants across 337,205 individuals in the UK Biobank study. *Nature communications*, 9(1):1–10, 2018.

[13] Xiucai Ding and Fan Yang. Spiked separable covariance matrices and principal components. *The Annals of Statistics*, 49(2):1113 – 1138, 2021.

[14] Jianqing Fan and Jinchi Lv. Sure independence screening for ultrahigh dimensional feature space. *Journal of the Royal Statistical Society: Series B (Statistical Methodology)*, 70(5):849–911, 2008.

[15] Zijian Guo, Hyunseung Kang, T Tony Cai, and Dylan S Small. Confidence intervals for causal effects with invalid instruments by using two-stage hard thresholding with voting. *Journal of the Royal Statistical Society: Series B (Statistical Methodology)*, 80(4):793–815, 2018.

[16] Fernando Pires Hartwig, George Davey Smith, and Jack Bowden. Robust inference in summary data mendelian randomization via the zero modal pleiotropy assumption. *International Journal of Epidemiology*, (6):6, 2017.

[17] Simon Haworth, Ruth Mitchell, Laura Corbin, Kaitlin H. Wade, Tom Dudding, Ashley Budu-Aggrey, David Carslake, Gibran Hemani, Lavinia Paternoster, and George Davey and Smith. Apparent latent structure within the UK Biobank sample has implications for epidemiological analysis. *Nature Communications*, 10(1), 2019.

[18] Hyunseung Kang, Anru Zhang, T Tony Cai, and Dylan S Small. Instrumental variables estimation with some invalid instruments and its application to mendelian randomization. *Journal of the American statistical Association*, 111(513):132–144, 2016.

[19] Michal Kolesár, Raj Chetty, John Friedman, Edward Glaeser, and Guido W Imbens. Identification and inference with many invalid instruments. *Journal of Business & Economic Statistics*, 33(4):474–484, 2015.

[20] Sang Hong Lee, Naomi R Wray, Michael E Goddard, and Peter M Visscher. Estimating missing heritability for disease from genome-wide association studies. *American journal of human genetics*, 88(3):294–305, 03 2011.

[21] Zhonghua Liu, Ting Ye, Baoluo Sun, Mary Schooling, and Eric Tchetgen Tchetgen. On Mendelian Randomization Mixed-Scale Treatment Effect Robust Identification (MR MiSTERI) and Estimation for Causal Inference. *arXiv preprint arXiv:2009.14484*, 2020.

[22] Po-Ru Loh, George Tucker, Brendan K Bulik-Sullivan, Bjarni J Vilhjálmsson, Hilary K Finucane, Rany M Salem, Daniel I Chasman, Paul M Ridker, Benjamin M Neale, Bonnie Berger, Nick Patterson, and Alkes L Price. Efficient Bayesian mixed-model analysis increases association power in large cohorts. *Nature Genetics*, 47(3):284–290, 2015.

[23] Qiongshi Lu, Boyang Li, Derek Ou, Margret Erlendsdottir, Ryan L Powles, Tony Jiang, Yiming Hu, David Chang, Chentian Jin, Wei Dai, et al. A powerful approach to estimating annotation-stratified genetic covariance via GWAS summary statistics. *The American Journal of Human Genetics*, 101(6):939–964, 2017.

[24] Luke, J, O'Connor, Alkes, L, and Price. Distinguishing genetic correlation from causation across 52 diseases and complex traits. *Nature Genetics*, 2018.

[25] Teri A. Manolio, Francis S. Collins, Nancy J. Cox, David B. Goldstein, Lucia A. Hindorff, David J. Hunter, Mark I. McCarthy, Erin M. Ramos, Lon R. Cardon, Aravinda Chakravarti, Judy H. Cho, Alan E. Guttmacher, Augustine Kong, Leonid Kruglyak, Elaine Mardis, Charles N. Rotimi, Montgomery Slatkin, David Valle, Alice S. Whittemore, Michael Boehnke, Andrew G. Clark, Evan E. Eichler, Greg Gibson, Jonathan L. Haines, Trudy F. C. Mackay, Steven A. McCarroll, and Peter M. Visscher. Finding the missing heritability of complex diseases. *Nature*, 461(7265):747–753, 2009.

[26] Jean Morrison, Nicholas Knoblauch, Joseph H. Marcus, Matthew Stephens, and Xin He. Mendelian randomization accounting for correlated and uncorrelated pleiotropic effects using genome-wide summary statistics. *Nature Genetics*, 2020.

[27] Zheng Ning, Yudi Pawitan, and Xia Shen. High-definition likelihood inference of genetic correlations across human complex traits. *Nature genetics*, 52(8):859–864, 2020.

[28] Matti Pirinen, Peter Donnelly, and Chris CA Spencer. Efficient computation with a linear mixed model on large-scale data sets with applications to genetic studies. *The Annals of Applied Statistics*, pages 369–390, 2013.

[29] Alkes L Price, Nick J Patterson, Robert M Plenge, Michael E Weinblatt, Nancy A Shadick, and David Reich. Principal components analysis corrects for stratification in genome-wide association studies. *Nature Genetics*, 38(8):904–909, 2006.

[30] Guanghao Qi and Nilanjan Chatterjee. Mendelian Randomization Analysis Using Mixture Models (MRMix) for genetic effect-size-distribution leads to robust estimation of causal effects. *bioRxiv*, page 367821, 2018.

[31] Eric Tchetgen Tchetgen, BaoLuo Sun, and Stefan Walter. The GENIUS approach to robust Mendelian randomization inference. *Statistical Science*, 36(3):443–464, 2021.

[32] Peter M. Visscher, Loic Yengo, Nancy J. Cox, and Naomi R. Wray. Discovery and implications of polygenicity of common diseases. *Science*, 373(6562):1468–1473, 2021.

[33] Huanwei Wang, Futao Zhang, Jian Zeng, Yang Wu, Kathryn E Kemper, Angli Xue, Min Zhang, Joseph E Powell, Michael E Goddard, Naomi R Wray, et al. Genotype-by-environment interactions inferred from genetic effects on phenotypic variability in the UK Biobank. *Science advances*, 5(8):eaaw3538, 2019.

[34] Frank Windmeijer, Helmut Farbmacher, Neil Davies, and George Davey Smith. On the use of the lasso for instrumental variables estimation with some invalid instruments. *Journal of the American Statistical Association*, 114(527):1339–1350, 2019.

[35] Haoran Xue, Xiaotong Shen, and Wei Pan. Constrained maximum likelihood-based Mendelian randomization robust to both correlated and uncorrelated pleiotropic effects. *The American Journal of Human Genetics*, 108(7):1251–1269, 2021.

[36] Jian Yang, Beben Benyamin, Brian P McEvoy, Scott Gordon, Anjali K Henders, Dale R Nyholt, Pamela A Madden, Andrew C Heath, Nicholas G Martin, Grant W Montgomery,

et al. Common snps explain a large proportion of the heritability for human height. *Nature genetics*, 42(7):565–569, 2010.

[37] Ting Ye, Zhonghua Liu, Baoluo Sun, and Eric Tchetgen Tchetgen. GENIUS-MAWII: For Robust Mendelian Randomization with Many Weak Invalid Instruments. *arXiv preprint arXiv:2107.06238*, 2021.

[38] Ting Ye, Jun Shao, and Hyunseung Kang. Debiased inverse-variance weighted estimator in two-sample summary-data mendelian randomization. *The Annals of Statistics*, 49(4):2079–2100, 2021.

[39] Loic Yengo, Jian Yang, and Peter M. Visscher. Expectation of the intercept from bivariate LD score regression in the presence of population stratification. *bioRxiv*, 2018.

[40] Yiliang Zhang, Youshu Cheng, Wei Jiang, Yixuan Ye, Qiongshi Lu, and Hongyu Zhao. Comparison of methods for estimating genetic correlation between complex traits using GWAS summary statistics. *Briefings in bioinformatics*, 22(5):bbaa442, 2021.

[41] Qingyuan Zhao, Jingshu Wang, Gibran Hemani, Jack Bowden, and Dylan S Small. Statistical inference in two-sample summary-data mendelian randomization using robust adjusted profile score. *The Annals of Statistics*, 48(3):1742–1769, 2020.

[42] Hua Zhou, Liuyi Hu, Jin Zhou, and Kenneth Lange. MM algorithms for variance components models. *Journal of Computational and Graphical Statistics*, 28(2):350–361, 2019.

[43] Wei Zhou, Jonas B Nielsen, Lars G Fritsche, Rounak Dey, Maiken E Gabrielsen, Brooke N Wolford, Jonathon LeFaive, Peter VandeHaar, Sarah A Gagliano, Aliya Gifford, et al. Efficiently controlling for case-control imbalance and sample relatedness in large-scale genetic association studies. *Nature genetics*, 50(9):1335–1341, 2018.

[44] Xiang Zhou, Peter Carbonetto, and Matthew Stephens. Polygenic Modeling with Bayesian Sparse Linear Mixed Models. *PLOS Genetics*, 9:1–14, 02 2013.

[45] Xiang Zhu and Matthew Stephens. Bayesian large-scale multiple regression with summary statistics from genome-wide association studies. *The Annals of Applied Statistics*, 11(3):1561–1592, 2017.

[46] Xiang Zhu and Matthew Stephens. Large-scale genome-wide enrichment analyses identify new trait-associated genes and pathways across 31 human phenotypes. *Nature Communications*, 9(1):4361, 2018.
